# A Network-centric Framework for the Evaluation of Mutual Exclusivity Tests on Cancer Drivers

**DOI:** 10.1101/2021.08.05.455220

**Authors:** Rafsan Ahmed, Cesim Erten, Aissa Houdjedj, Hilal Kazan, Cansu Yalcin

## Abstract

One of the key concepts employed in cancer driver gene identification is that of mutual exclusivity (ME); a driver mutation is less likely to occur in case of an earlier mutation that has common functionality in the same molecular pathway. Several ME tests have been proposed recently, however the current protocols to evaluate ME tests have two main limitations. Firstly the evaluations are mostly with respect to simulated data and secondly the evaluation metrics lack a network-centric view. The latter is especially crucial as the notion of common functionality can be achieved through searching for interaction patterns in relevant networks. We propose a network-centric framework to evaluate the pairwise significances found by statistical ME tests. It has three main components. The first component consists of metrics employed in the network-centric ME evaluations. Such metrics are designed so that network knowledge and the reference set of known cancer genes are incorporated in ME evaluations under a careful definition of proper control groups. The other two components are designed as further mechanisms to avoid confounders inherent in ME detection on top of the network-centric view. To this end, our second objective is to dissect the side effects caused by mutation load artifacts where mutations driving tumor subtypes with low mutation load might be incorrectly diagnosed as mutually exclusive. Finally, as part of the third main component, the confounding issue stemming from the use of nonspecific interaction networks generated as combinations of interactions from different tissues is resolved through the creation and use of tissue-specific networks in the proposed framework. The data, the source code and useful scripts are available at: https://github.com/abu-compbio/NetCentric.

## 1 INTRODUCTION

Cancer is a disease caused mostly due to a gradual accumulation of somatic alterations that give rise to pathway dysregulation through alterations in copy number, DNA methylation, gene expression, and molecular function. An important challenge in cancer genomics is to distinguish driver mutations from passenger mutations. The former are those determined to be causal for cancer progression, whereas the latter are characterized as those not leading to any selective advantage. Several computational methods have been proposed for the identification of cancer driver genes or driver modules of genes by integrating mutations data with various other types of genetic data; see Dimitrakopoulos and Beerenwinkel [2017], Zhang and Zhang [2018], Bailey et al. [2018], Tokheim et al. [2016] for recent comprehensive evaluations and surveys on the topic.

A phenomenon observed frequently in the data pertaining to the alterations that the tumors acquire is *mutual exclusivity (ME)*; a driver mutation is less likely to occur in case of an earlier mutation that has common functionality in the same molecular pathway [Leiserson et al., 2016, van de Haar et al., 2019, Thomas et al., 2007, Yeang and Levine, 2008]. Therefore several driver gene or module identification approaches employ ME detection as part of their problem definitions and optimization goals [Babur et al., 2015, Ciriello et al., 2012, Leiserson et al., 2013b, Kim et al., 2015, Ahmed et al., 2019, Baali et al., 2020]. Such a central role in driver gene and module identification has led to the design of many different approaches for defining and computing mutual exclusivity. Some of these approaches are based on combinatorial definitions of mutual exclusivity [Vandin et al., 2012, Leiserson et al., 2013a, Basso et al., 2019, Ahmed et al., 2019, Song et al., 2020, Baali et al., 2020]. In most cases the combinatorial definitions are incorporated and tested within a driver gene or module identification framework, rather than as stand-alone ME tests. On the other hand, the vast majority of the ME detection approaches are based on statistical tests [Ciriello et al., 2012, Szczurek and Beerenwinkel, 2014, Leiserson et al., 2015a, Constantinescu et al., 2015, Hua et al., 2016, Canisius et al., 2016, Leiserson et al., 2016, Kim et al., 2017, Liu et al., 2020, Zhang et al., 2020] and in most cases for such approaches the specific goal is to provide ME significance results. Therefore the focus of the proposed framework is the evaluation of the latter set of approaches consisting of the statistical ME tests.

Among such approaches, MEMo builds a graph based on gene similarities and extracts cliques from this graph. To determine whether each clique has significant mutual exclusivity, it then proposes a null model generated by randomly permuting the set of genomic events, while preserving the overall distribution of observed alterations across both genes and samples, and introduces a Markov Chain Monte Carlo (MCMC) permutation strategy based on random network generation models [Ciriello et al., 2012]. Szczurek and Beerenwinkel [2014] propose a probabilistic, generative model of mutual exclusivity, explicitly taking coverage, impurity, and error rates into account. Based on such a model, they provide a statistical test of mutual exclusivity by comparing its likelihood to the null model that assumes independent gene alterations. Mutex [Constantinescu et al., 2015] defines the alteration of two genes to be mutually exclusive if their overlap in samples is significantly less than expected by chance, where the statistical significance of the overlaps are calculated using a hypergeometric test with the assumption of a uniform alteration frequency among samples. This may not always be the case as in many data sources there are hyper-mutated samples. The problem is resolved partially by simply excluding such samples from the analysis. CoMEt [Leiserson et al., 2015a] on the other hand provides an exact statistical test for mutual exclusivity conditional on the observed frequency of each alteration with the goal of introducing less bias towards high frequency alterations. Based on this it provides a tail enumeration procedure to compute the exact test, as well as a binomial approximation. DISCOVER provides a statistical independence test that makes no assumption of identical gene alteration probabilities across tumors [Canisius et al., 2016]. The alteration probabilities are estimated by solving a constrained optimization problem guaranteeing the probabilities are consistent with both the observed number of alterations per gene and the observed number of alterations per tumor. The tumor-specific gene alteration probabilities are then used to compute the probability of concurrent alterations which in turn are used to decide whether the number of tumors altered in both genes deviates from the expectation through an analytical test based on the Poisson-binomial distribution. WeXT provides a weighted exact test that conditions simultaneously on the number of samples with a mutation and the per-event, per-sample mutation probabilities [Leiserson et al., 2016]. A recursive formulation to compute P-values for this weighted test exactly and a saddle-point approximation of the test are proposed. WeSMe provides a permutation-based test and an approximation of significance through a weighted sampling technique that enables further improvements in running time spent for sampling and a way to obtain a better precision without increasing the computational time significantly [Kim et al., 2017]. Two recently suggested ME tests are FSME [Zhang et al., 2020] and MEScan [Liu et al., 2020]. The former proposes a *seed-and-extend* strategy to alleviate the computational cost of a permutation-based test. The seed pairs are constructed by a combinatorial formulation incorporating both ME and the coverage of the pair. The seeds are then grown with new genes by employing an independence test. MESCan provides a test statistic that incorporates a patient and gene-specific background mutation rate in the calculation to adjust for the background noise, and that includes a gene-specific weight to down-weigh genes with high mutation rates. Such a statistic is then employed in an MCMC algorithm followed by a false discovery rate control.

We propose a network-centric framework to evaluate the pairwise significances found by statistical ME tests. It is important to make a distinction between the network-centric view of the current study and that of the previous studies employing both network data and the concept of ME [Ciriello et al., 2012, Leiserson et al., 2013b, Kim et al., 2015, Ahmed et al., 2019, Baali et al., 2020]. The latter are network-centric in the sense that the proposed ME tests are applied on interacting pairs or subnetworks as part of a more general goal of identifying cancer driver genes/modules. Thus due to the nature of the set objectives their evaluations focus on the success of output genes/modules matching reference cancer-related drivers/pathways. The proposed study takes on an approach in the opposite direction; we assume the interaction network and the reference cancer-related drivers to be inputs to our framework which evaluates the success of various ME tests. The focus of the proposed framework is on pairwise significances since one of the major application areas where ME tests are commonly employed is knowledge-based cancer driver identification where pairwise ME significances are of major essence. In terms of the general objectives our work is most similar to that of Deng et al. [2017], where a framework for performance comparisons of statistical ME detection approaches is proposed and executed on six such tests. An important distinction is that the performance analysis of Deng *et al.* is based on experiments with simulated data and the framework does not suggest any mechanism to avoid confounders inherent in ME detection. One such confounder is due to the alterations specific to cancer subtypes [Deng et al., 2017, van de Haar et al., 2019]. Alterations in different subtypes may be incorrectly diagnosed with ME, although the alterations are not due to any natural root causes of ME such as redundant functionality. Our network-centric view aims to recognize such false positives by constructing reference sets based on known drivers gathered from neighborhoods of interaction networks. Furthermore, inspired by the *mutation load confounding* concept of van de Haar et al. [2019], we extend our network-centric framework to dissect side effects caused by mutation load artifacts; mutations that drive tumor subtypes with low mutation load might be incorrectly diagnosed as mutually exclusive. A possible drawback of the proposed network-centric evaluation framework would be due to the use of nonspecific interaction networks that are generated as combinations of interactions from different tissues and are thus suboptimal in resolving confounding issues of mutual exclusivity. In order to detect whether there exists such discrepancies or to limit their effect if they do, we therefore refine the network-centric approach by designing further tests on *tissue-specific networks (TSN)* we construct based on gene co-expression.

## 2 METHODS

The overall network-centric ME evaluations framework has three main components. The first one consists of definitions of the metrics employed in the network-centric ME evaluations. Such metrics are designed so that network knowledge and the reference set of known cancer genes are incorporated in ME evaluations under a careful definition of proper control groups. The second component detects whether the use of the interactome information provides similar advantages in ME corrections of pairwise mutual exclusivity findings as the subtype-stratification idea suggested by van de Haar et al. [2019]. Finally, the third component extends our framework to incorporate tissue-specific networks with the aim of reducing the possible side effects of using nonspecific interaction networks.

### 2.1 Metrics for the Network-centric ME Evaluations

Assuming that cancer driver genes in the same pathway are more likely to show mutually exclusive mutation profiles, we utilize the interactome to devise a strategy for evaluating the ME methods and the effects of the interactome information on quantifying ME. Let 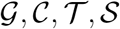, *p_t_, c* denote respectively the input Protein-Protein Interaction (PPI) network, the employed cohort, the statistical ME test undergoing the network-centric ME evaluations, the golden standard reference gene set of known cancer drivers, the p-value threshold for significance, and the type of the control group to be employed. Let 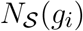 denote the set of genes from 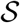 that are in the neighborhood of the node corresponding to gene *g_i_* in the PPI network 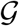. For a gene 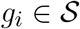, corresponding to each neighbor 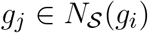, we randomly select a gene *g_r_* from a control group 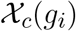, and compute *T P^cur^*, *F P^cur^*, based on the – log-transformed p-values *p_i,j_* and *p_i,r_* as computed by the ME test 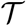. Here *p_i,j_* denotes the significance of the mutual exclusivity of the pair *g_i_, g_j_* for 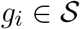 and 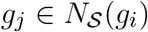, and *p_i,r_* denotes the significance of the mutual exclusivity of the pair *g_i_, g_r_* for a random gene *g_r_* from the control group. Based on the premise that cancer driver genes interacting in the PPI network are likely to exhibit ME, a pair *g_i_, g_j_* belongs to the set of True Positives if *p_i,j_* is significant and a pair *g_i_, g_r_* belongs to the set of False Positives if *p_i,r_* is significant. To obtain robust results, the selection of the random genes from the control group is repeated 100 times. Finally the medians of these 100 instances are summed over all genes 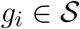 to provide the necessary statistics *T P, F P*. Thus precision, sensitivity, and the F1 scores are computed based on these 4 statistics. The number of condition positives used in the sensitivity calculations corresponds to the number of pairs *g_i_*, 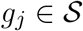 where *g_i_, g_j_* interact in 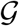.

We note that limiting our focus solely on these conventionally formed *T P, F P* classes may be misleading as each one considers the significance of *p_i,j_* and *p_i,r_* individually. A more detailed inspection with a simultaneous consideration of their values could prove more insightful in certain cases since they both involve a common gene *g_i_*. Towards this aim we introduce the *strict* versions of these conventional classes. More specifically *T P_strict_* consists of *g_i_, g_j_* pairs where *p_i,j_* is significant not only with respect to the given threshold but also as compared to the p-value of the control pair *g_i_, g_r_*. Similarly *F P_strict_* consists of the control pairs *g_i_, g_r_*, where *p_i,r_* is more significant than both the threshold value and *p_i,j_*. Based on these strict classes we can compute three metrics: precision_*strict*_, sensitivity_*strict*_, and F1_*strict*_. Such a consideration is especially convenient in reducing any potential bias inherent in genes like TP53 which have large mutation frequencies almost exclusively in tumors with small numbers of mutations; both *p_i,j_* and *p_i,r_* are likely to be significant in such a scenario giving rise to vagueness in the conventional F1 score.

A comparison of F1_*strict*_ values based on the two statistics simultaneous by their nature, *TP_strict_* and *FP_strict_* provides a more rigorous evaluation in such cases.

For the network-centric ME evaluations we employ two different definitions for the control groups. For the first one, the control group *χ*_1_(*g_i_*) consists of genes in 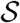 that do not interact with *g_i_* in the PPI network. For the second one, *χ*_2_(*g_i_*) consists of neighbors of *g_i_* in the PPI network that are not in 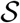. In the latter case only the genes 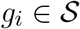 for which the number of neighbors not in 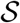 is larger than or equal to the number of neighbors in 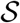 are taken into account.

### 2.2 Network-centric ME Corrections in Relation to MLA

Some statistical mutual exclusivity tests are based on the assumption that genes’ alterations across tumors are identically distributed. Among the approaches considered in this study Fisher’s Exact Test and MEGSA belong to this category. However, it has been observed that the number of alterations per tumor can vary quite considerably, even in tumors of the same type; colorectal tumors with microsatellite stability have a median of 66 non-synonymous mutations, but colorectal tumors with microsatellite instability have a median of 777 mutations [Vogelstein et al., 2013, Leiserson et al., 2016]. It has been shown that under such settings the mutual exclusivity tests relying on identical alteration probabilities across tumors may lead to reduced sensitivity for mutual exclusivity analysis [Canisius et al., 2016]. The effects of varying alteration probabilities on pairwise mutual exclusivity calculations have been formalized within the context of the so-called *mutation load confounding (MLC)* in a recent study by van de Haar et al. [2019]. MLC is a correlation between the number of statistically significant mutual exclusivity findings and the *mutation load association (MLA)* of a gene, where logistic regression is used to compute MLA as a standardized score of association between the mutation likelihood of each gene and the *mutation load*, that is the genome-wide number of somatic mutations observed in a tumor. Note that negative MLA values correspond to higher mutation frequencies in tumors with low mutation loads, whereas positive values correspond to higher mutation frequencies in tumors with high mutation loads. Strong negative correlations between the MLA of a gene and the number of statistically significant pairwise mutual exclusivities have been observed, implicating the finding that the more negative a gene’s MLA, the higher the number of other genes that show mutual exclusivity with that particular gene [van de Haar et al., 2019]. However, such a negative correlation does not always imply true ME since a gene that exclusively shows large mutation frequency in tumors with low mutation loads, naturally has a better chance of forming mutually exclusive pairs with other genes. Thus extra sources of information are necessary to filter out the pairs with true ME relations among a set of statistically significant pairwise mutual exclusivities postulated by some exclusivity test. van de Haar et al. [2019] make use of the subtype information for such a purpose and show that MLC can be reduced by correcting via tumor subtype stratification. Such a correction greatly reduces the number of gene pairs reported to show mutual exclusivity, especially for pairs that include genes with low MLA. A major drawback is the absence of subtype information for many tumors. As part of our network-centric ME framework, we suggest that such a correction can be efficiently done with the interaction network data, rather than or better yet on top of the subtype information. For this purpose we calculate the correlation between the number of statistically significant pairwise ME findings and the MLA for two settings; one where pairwise mutual exclusivities are sought between a gene in 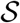 and all other genes in 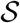, and the other where a gene in 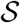 is checked against only its PPI neighbors that are in 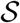. The computations of the two settings are repeated with the subtype-stratified data as well, to see the added value of the network-centric ME corrections on top of the subtype-based corrections on statistically significant pairwise MEs.

### 2.3 Network-centric ME Evaluations in Relation to TSN

Rather than using a common nonspecific network for all the cancer types, in this component of our evaluation framework we employ TSN based on the tissue in which the tumor develops. To construct the TSN for a particular tissue, we start with the original PPI network and remove the edges between the pairs of genes that are not co-expressed in the corresponding tissue. For this purpose, we download RNA-seq datasets from GTEX portal [GTEXConsortium, 2020]. See Supplementary Table 49 for the total number of available samples for each tissue. To determine the co-expressed genes, we follow the procedure described in Luck et al. [2020]. For each pair of genes that have an edge in the original PPI network, we identify the number of samples where both genes have Transcripts Per Kilobase Million (TPM) values ≥ 1. We then divide this number with the total number of samples where either gene has a TPM value ≥ 1. The resulting value is called the *co-expression ratio*. Gene pairs interacting in the original network are included in the TSN_*cor*_ if the *co-expression ratio* is ≥ *cor*, for a given threshold *cor*.

In addition to applying the network-centric metrics introduced in Section 2.1 on the constructed TSNs, we also propose a more detailed evaluation in terms of ROC analysis based on tissue-specificity. For this purpose, we define the gene pairs with co-expression ratio value of 1 as *tissue-specific gene pairs*. Similarly, the gene pairs with co-expression ratio values ≤ 0.5 are called *non-tissue-specific gene pairs*. To test whether a specific ME test identifies stronger mutual exclusivities for the tissue-specific gene pairs in 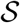, we rank the gene pairs in 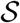 in increasing order of p-values. To construct the control group, we rank the same number of random samples of gene pairs not in 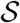 with respect to the p-values making sure that the sizes of the positive (or negative) sets of gene pairs not in 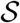 are exactly the same as those that are found for the gene pairs in 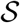. For both gene pairs in 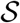 and gene pairs not in 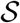, the set of positives consists of the tissue-specific gene pairs, whereas non-tissue-specific gene pairs are labelled as negatives. We then compute the True Positive Rate (TPR) and the False Positive Rate (FPR) for each case. Note that for robustness considerations the control group computations are repeated 100 times and the median TPR and FPR values are reported.

## 3 RESULTS

### 3.1 Input Data and Parameter Settings

The somatic mutation data from TCGA was preprocessed and provided by van de Haar et al., 2019. The 8 different cancer types and their corresponding tumor samples within the dataset is as follows: BLCA (411), BRCA (1026), COADREAD (498), LUAD (568), LUSC (485), SKCM (468), STAD (438) and UCEC (531). The preprocessing step involves the removal of all mutations with ‘variant classification’ of ‘Silent’, ‘3’UTR’, ‘Intron’, ‘5’UTR’, ‘RNA’, ‘3’Flank’ and ‘5’Flank’ from the TCGA data. The input data is then further filtered by mutation frequency threshold, *t*, to include genes with > *t* mutations across the cohort. More specifically, with *t* = 20 we include the genes that are mutated in more than 20 samples within the cancer type under study. Regarding subtypes, we download subtype information for BRCA from the cBioPortal [Cerami et al., 2012, Gao et al., 2013] and the CMS stratification for COADREAD from [Guinney et al., 2015]. We use the COSMIC Cancer Gene Census database to compile the set of known cancer genes [Sondka et al., 2018]. For the results presented in the main document we employ the IntAct PPI network as it is a comprehensive and well-characterized database [Orchard et al., 2014]. As a preprocessing step, we remove duplicate edges and edges below the confidence threshold of 0.35 from the network. The final network contains 15,079 nodes and 103,520 edges. For the gene expression data employed in the construction of TSNs, we download RNA-Seq data from the Genotype-Tissue Expression (GTEx) portal [GTEXConsortium, 2020] (05-06-2017).

For the comparative evaluations of our network-centric framework described in the previous section, we choose six popular statistical mutual exclusivity methods: DISCOVER [Canisius et al., 2016], DISCOVER Strat [Canisius et al., 2016, van de Haar et al., 2019], Fisher’s Exact Test, WeXT [Leiserson et al., 2016],MEMo [Ciriello et al., 2012] and MEGSA [Hua et al., 2016]. Among these, MEMo and MEGSA are originally designed to output p-values for a set of genes with size > 2. For MEMo, we re-implement the first part of the algorithm where pairwise ME p-values are estimated. We use *Q* = 100 and and *N* = 10, 000 as suggested by the original paper [Ciriello et al., 2012]. For MEGSA, pairwise ME p-values are calculated by applying chi-square cumulative probability less than or equal to the value of the log likelihood calculated by the *funestimate* function. We should note that all the employed methods output multiple testing corrected p-values. With regards to the parameter settings of our proposed framework, we employ the values of 5 and 20 for *t*.

### 3.2 ME Evaluations Based on Defined Metrics

Table 1 and 2 show the results of evaluating the six ME detection methods on COADREAD data where *t* = 20 and we use the data from 498 patients for which subtype information is available. We use *χ*_1_ and *χ*_2_ as the control group in Table 1 and 2, respectively. We first discuss the results of *χ*_1_. We observe that DISCOVER Strat gives the highest precision and precision_*strict*_ values. The ranking of the other methods from best to worst in terms of precision or precision_*strict*_ is as follows: WeXT, DISCOVER, MEMo, MEGSA and Fisher’s Exact Test. A comparison of the precision and precision_*strict*_ values distinguishes two groups of ME methods; for DISCOVER, DISCOVER Strat, Fisher’s Exact Test, and WexT the precision_*strict*_ values are greater than or equal to the precision values, whereas the exact opposite is observed for MEGSA and MEMo. This suggests that the performance of the methods in the latter group gets worse when random control gene pair is considered simultaneously in the precision calculation, that is precision_*strict*_. Compared to the precision, we observe much larger differences among the sensitivity or the sensitivity_*strict*_ values output by the employed methods. We can group the methods into two where the first group contains WeXT, MEMo and DISCOVER, and the second group contains the rest of the methods. The first group of methods give much larger sensitivity or sensitivity_*strict*_ values than the second. For instance, the sensitivity value obtained with WeXT is an order of magnitude larger than that of Fisher’s Exact Test. This also shows that the second group of methods are more conservative than the first group of methods. WeXT is the least conservative approach based on its high sensitivity value. Even though WexT predicts many significant p-values, it still has a competitive precision value which is slightly lower than the maximum observed value (0.725 vs 0.727). Accordingly, WeXT obtains the best F1 score and F1_*strict*_ score which is followed by MEMo and DISCOVER. The remaining three methods give much smaller F1 scores and they rank as follows from highest to lowest: MEGSA, DISCOVER Strat and Fisher’s Exact Test. Comparing the conventional F1 score with the F1_*strict*_ score of each ME method, the largest difference is observed for MEMo indicating that the consideration of the random pair as a control affects its performance dramatically. Another interesting observation is the lower performance of DISCOVER Strat compared to DISCOVER which suggests that the use of subtype information is not useful for COADREAD. Table 2 shows the results where *χ*_2_ is used as the control group. Since *χ*_2_(*g_i_*) is defined as the non-CGC neighbors of *g_i_* in the PPI network, we can only consider the CGC genes that have more non-CGC neighbors than CGC neighbors. As such, the number of pairs included in this analysis is much smaller than that of Table 1 (107 vs 196). The ranking of the methods in Table 2 with respect to F1 score and sensitivity remain the same as Table 1. However, there are differences in the ranking with respect to other metrics. For instance, WeXT ranks best in terms of precision whereas the best ranking method in Table 1, DISCOVER Strat, ranks the fifth. Compared to Table 1, the precision values of all the methods are smaller in Table 2. We see the opposite trend for sensitivity values. These changes are in parallel with the increase in percent significant p-values output by the methods. For instance, the percentage of significant p-values output by DISCOVER is 12% in Table 1 and 18% in Table 2. We also observe differences between the conventional and the strict versions of the employed metrics. WeXT and DISCOVER have increased precision_*strict*_ values compared to precision whereas we observe the opposite trend for the rest of the methods. Additionally, the ranking of the methods with respect to F1 score and F1_*strict*_ score is different. Namely, MEMo’s ranking decreases from second highest to third highest when we switch from F1 score to F1_*strict*_ score. Accordingly, DISCOVER’s ranking improves from third highest to second highest based on F1 score. This increases the confidence of DISCOVER results as F1_*strict*_ requires a stricter definition of true and false positives.

**Table 1.**
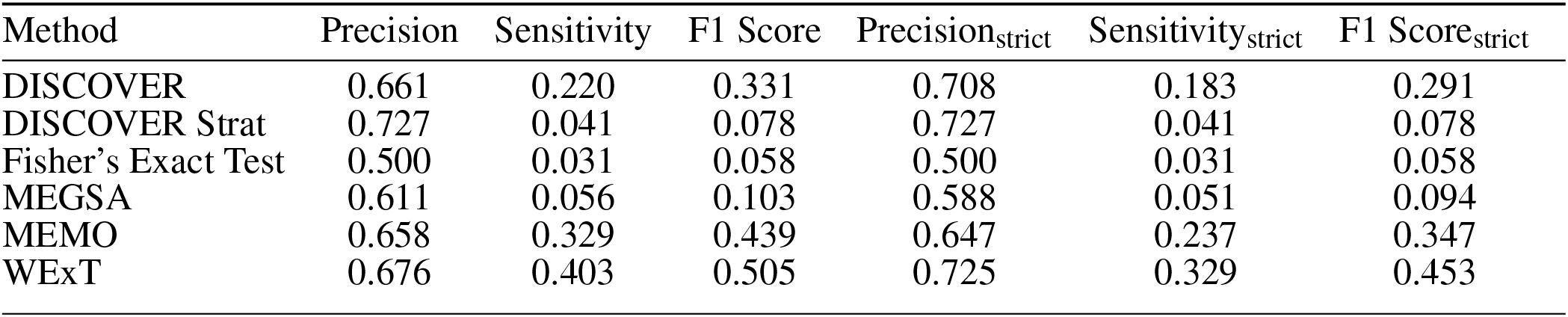
Results of network-centric ME evaluation framework with control group *χ*_1_ COADREAD t20 (498 samples, 196 CGC-CGC pairs)

**Table 2.**
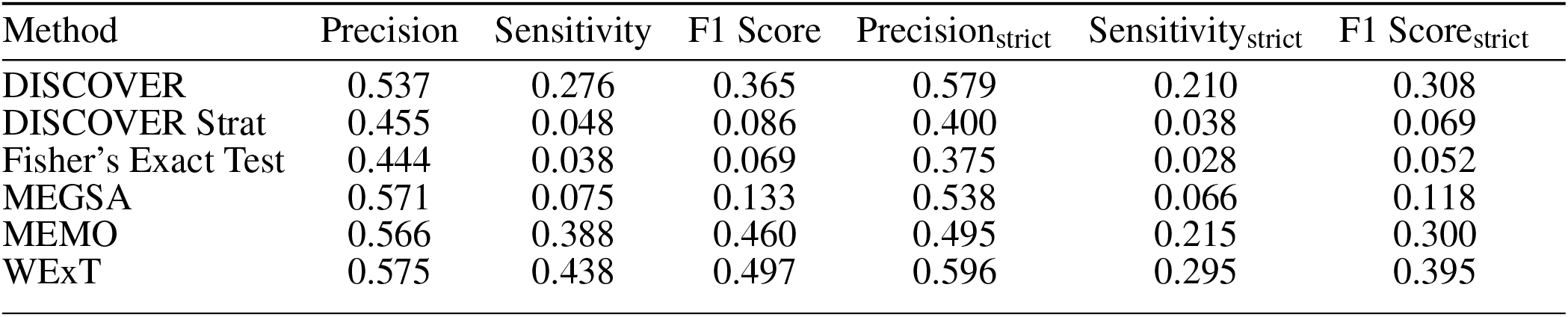
Results of network-centric ME evaluation framework with control group *χ*_2_ COADREAD t20 (498 samples, 107 CGC-CGC pairs)

Table S1 show the results with *χ*_1_ control group and *t* = 20 filtering for the other cancer types. We observe that the methods report a small number of significant p-values for BLCA data (Table S1-a). In line with this, we observe smaller sensitivity values for BLCA data compared to the results on COADREAD data. For BRCA dataset, we observe that the top F1 score is obtained by DISCOVER Strat indicating the benefit of considering subtype information for BRCA (Table S1-b). When we compare the values of the conventional and strict versions of the metrics, we observe that DISCOVER Strat’s precision_*strict*_ is significantly larger than its precision value whereas we observe negligible changes for the other methods. Accordingly, DISCOVER Strat’s F1_*strict*_ score is slightly lower than its F1 score whereas we observe significant drops for the other methods. We also observe that MEGSA and Fisher’s Exact Test perform significantly worse than the other methods. Interestingly MEGSA ranks the best in terms of F1 score for LUSC dataset (Table S1-e). However, we should point out that the methods report a small number of significant p-values similar to the BLCA dataset. WeXT shows a dramatically better performance on SKCM dataset (Table S1-f). For UCEC dataset, we observe that Fisher’s Exact Test and MEGSA perform poorly due to their conservative calculation of p-values when compared to DISCOVER and WeXT (Table S1-h). Table S13 show the analogous results when the control group is defined as *χ*_2_. We observe that the analysis includes less than 50 pairs for BLCA, BRCA and LUSC. Again, we observe that Fisher’s Exact Test and MEGSA report zero or very few number of significant p-values across the majority of the the cancer types. For LUAD, SKCM, STAD and UCEC, WeXT gives the largest F1-values. Similar to the results obtained with *χ*_1_, MEGSA ranks the top in terms of F1-score on LUSC dataset. MEGSA and MEMo results are not available for some cancer types since we are unable to run these methods due to memory issues.

Table 3 and 4 show the COADREAD results of *t* = 5 setting with *c* = *X*_1_ and *c* = *X*_2_, respectively. Using a lower value for *t* increases the number of gene pairs tested in our analysis. When we compare these results with the results we obtained when *t* = 20, we observe few differences. Though the number of tested gene pairs is larger, the percentage of significant p-values obtained by the methods decreases. For instance, the percentage of significant p-values output by WeXT for COADREAD data decreases from 42% to 14% when *t* is changed from 20 to 5. This is likely related to the larger inclusion of low mutation frequency genes when *t* = 5. An interesting observation for *t* = 5 results is the decrease in DISCOVER Strat’s performance. For COADREAD, DISCOVER Strat’s precision and precision_*strict*_ value is the highest for *t* = 20 when _1_ is used as the control group. However, when *t* = 5, we observe that it ranks after WeXT and DISCOVER in terms of precision/precision_*strict*_ value. Similarly, for BRCA dataset, DISCOVER Strat ranks after WeXT for both control groups *χ*_1_ and *χ*_2_ (Table S25-b, Table S37-b).

**Table 3.**
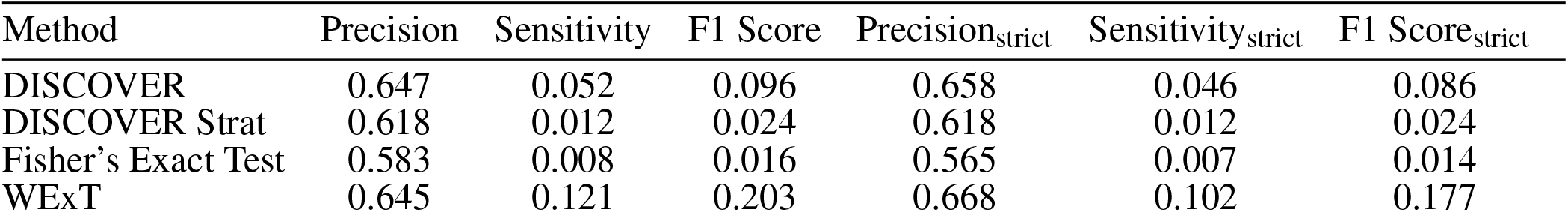
Results of network-centric ME evaluation framework with control group *χ*_1_ COADREAD t5 (498 samples, 1748 CGC-CGC pairs)

**Table 4.**
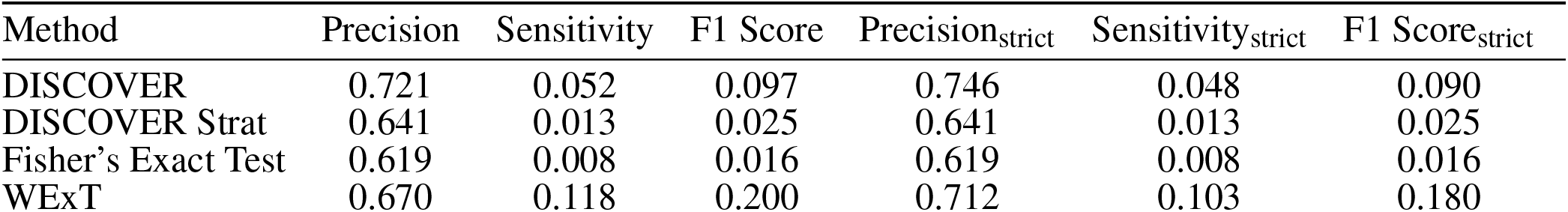
Results of network-centric ME evaluation framework with control group *χ*_2_ COADREAD t5 (498 samples, 1625 CGC-CGC pairs)

### 3.3 Robustness Analysis of Evaluations Based on Defined Metrics

We also investigate the robustness of our results with respect to *robustness iterations* value, the p-value significance threshold value, the reference gene set and the employed PPI network. For *robustness iterations*, we try the values 300 and 500. For p-value significance threshold, we try a more stringent significance threshold of 0.01. Since the employed ME methods output multiple testing corrected p-values, we also try a less stringent significance threshold: 0.1. Regarding the reference gene set, we try using a subset of CGC genes to include only those which have SNV type of mutations in cancer (378 out of 723 genes). To this end, we filter out the genes where the *mutation type* column consists of only A (amplification), D (large deletion) or T (translocation). Additionally, we use an alternative source named *IntoGen* [Martínez-Jiménez et al., 2020] to compile reference cancer genes. We download *Unfiltered driver results* 05*.tsv* file (2020-02-02 release) and include the genes where FILTER column is *PASS*, which results in 503 genes. For the PPI network, we try different confidence threshold values for filtering IntAct. 0.45 threshold value is commonly used in the literature to filter out the interactions with low confidence [Sügis et al., 2019, Porras et al., 2020]. Additionally, to observe the effect of using a lower threshold than the current one, we also tried filtering IntAct with a threshold value of 0.25. Lastly, we utilize two alternative networks in our analyses: *HINT* + *HI*2012 [Das and Yu, 2012, Yu et al., 2011, Leiserson et al., 2015b] and STRING network [Szklarczyk D et al., 2018]. For the latter, we download the file *9606.protein.physical.links.v11.0.txt* and only use physical interactions with scores greater than 700. This results in 9,524 nodes and 146,120 edges.

The results with these different settings are available in Table S2-S9 for *t* = 20 and *c* = *X*_1_. Table S2 and S3 show the results with *robustness iterations* value 300 and 500, respectively. When *robustness iterations* value is increased from 100 to 300, the ranking of the ME methods based on F1 score remains the same for all the cancer types except for BRCA (Table S2). Similarly, when the *robustness iterations* value is increased from 100 to 500, the ranking of the ME methods based on F1 score changes only for BLCA and BRCA (Table S3). Table S4 shows the results where p-value significance threshold is decreased from 0.05 to 0.01. We observe that the 0.01 threshold is too strict for most of the methods as evident from low sensitivity values for most cancer types. In fact, Fisher’s Exact Test predicts no mutually exclusive cases for six out of eight cancer types. When we use the p-value threshold of 0.1, we observe an overall improvement in F1 scores for all cancer types and for all the methods except for MEGSA which shows decreased F1 scores for four cancer types (Table S5). The observed increase in F1 scores for the majority of the cases is due to the large increase in sensitivity values. Additionally, for some cases precision values also increase with this less stringent p-value threshold. In particular, we observe a dramatic increase in F1/F1_*strict*_ scores for LUAD cancer type. In terms of ranking of the methods, we observe a difference in BLCA and BRCA types. For BLCA, WeXT’s ranking improves from fourth place to first place whereas MEMO’s ranking decreases from first place to third place. We observe a similar switch in ranking of the methods for BRCA. When CGC_*SNV*_ is used as the reference gene set, F1 score-based ranking of the ME methods changes for three cancer types: BLCA, BRCA and LUSC (Table S6). Though, among these three cancer types, the top ranking method remains the same for BRCA and LUSC. When *IntoGen* is used as the reference set, we only observe a change in ranking of the ME methods for BLCA and BRCA types (Table S7). Next, for the experiments where we try different confidence thresholds for IntAct or switch to the *HINT* + *HI*2012 network, the only cancer types where the ranking of the ME methods change are BLCA and BRCA (Table S8-S10). When we use the STRING network, we observe a decrease in F1/F1_*strict*_ scores of all the methods except for WeXT for BRCA and UCEC cancer types as compared to the results obtained with the IntAct network (Table S11). For COADREAD, LUAD and STAD we observe both increases and decreases in F1/ F1_*strict*_ scores. Lastly, all the methods show decreased F1/ F1_*strict*_ scores in LUSC and SKCM. The ranking of the methods based on F1 score and F1_*strict*_ score differs for BRCA and COADREAD datasets. For BRCA, WeXT and DISCOVER Strat switch positions in the ranking based on F1/ F1_*strict*_ scores. For COADREAD, MEMO’s ranking decreases from first position to third position showing that its top performance is not preserved with stricter versions of evaluation metrics. Additionally, the ranking of the methods change in five out of eight cancer types as compared to the results obtained with IntAct network suggesting that the input PPI plays a role in the performance of ME methods.

Results of robustness analyses for *t* = 20 and *c* = *X*_2_ setting is available in Table S13-S20. When *robustness iterations* is increased to 300 or to 500, the ranking of the ME methods remains the same for all the cancer types. Using a stricter p-value significance threshold of 0.01 results in too few mutual exclusivities predicted for many of the cancer types (Table S15). For instance, for LUAD, the sensitivity values are zero for all the methods. Similarly, for BLCA, SKCM and STAD cancer types the maximum sensitivity value across the ME methods is 0.1. When p-value threshold is switched to 0.1, we observe changes in both directions for different cancer types (Table S16). For BRCA, F1/F1_*strict*_ scores of all the methods either remain the same or decrease. On the other hand, all the methods have increased F1/F1_*strict*_ scores for COADREAD. We also observe a change in the ranking of the methods when we change the p-value threshold from 0.05 to 0.1. For LUAD, SKCM and UCEC, we observe significant increase in F1/F1_*strict*_ scores of all the methods. For LUSC and STAD, we see both increases and decreases in F1/F1_*strict*_ scores. In particular, WeXT and DISCOVER show dramatic increases in F1/F1_*strict*_ scores for STAD type. Overall, we can conclude that the p-value threshold of 0.1 leads to a better performance across the employed methods. When the reference gene set is changed to CGC_*SNV*_ or to *IntoGen*, the F1 score-based ranking of the methods only change for BLCA (Table S18-S19). When we use a version of IntAct filtered with confidence value 0.25, the ranking remains the same for all the cancer types except for BLCA, BRCA and LUSC (Table S20). When the confidence value threshold is increased to 0.45, the number of considered CGC-CGC gene pairs decreases to values that are < 20 for all the cancer types except for SKCM and UCEC (Table S21). For SKCM and UCEC, we observe the same ranking as in our original parameter settings. Switching to *HINT* + *HI*2012 network also leads to the inclusion of too few gene pairs for many cancer types (Table S22). For the rest, the ranking of the ME methods is in accordance with our original results. When we use the STRING network as the input PPI, we observe very few CGC-CGC pairs for BLCA, BRCA and LUSC. Comparing these results with those obtained with the IntAct network, for COADREAD, MEMo and WeXT are the only methods that show decreased F1/F1_*strict*_ scores where the magnitude of change is much larger for MEMo. However, these changes do not lead to a difference in the ranking of the methods. For LUAD, we observe large improvements in F1/F1_*strict*_ scores of all the methods. Lastly, for SKCM, STAD and UCEC we observe smaller F1/F1_*strict*_ scores in almost all the cases (Table S23). These results reveal that the change of the network leads to distinct changes in different cancer types.

Robustness analyses for *t* = 5 setting is available in Table S25-S35 and in Table S37-S47 for *c* = *X*_1_ and *c* = *X*_2_, respectively. We observe similar patterns compared to the *t* = 20 setting. For instance, changing the p-value threshold to 0.01 decreases the F1/F1_*strict*_ scores overall whereas increasing the threshold to 0.1 also increases the F1/F1_*strict*_ scores. Overall, changing the different settings of the analysis do not lead a change in the ranking of the methods.

We also assess whether the F1 scores improve or worsen when different confidence thresholds are used to filter the IntAct network. Increasing the interaction confidence threshold increases the support of both the reference pair and the control pair with respect to *X*_2_, since they are both interacting pairs. In parallel with this, for the *t* = 5 setting where we have adequate number of gene pairs for all the cancer types, we observe an increased F1 score for all the methods for all the cancer types except for BRCA and LUAD. For these two cancer types, we observe both increases and decreases in F1 scores. On the other hand, it is difficult to propose a similar argument for *c* = *X*_1_ setting, since increasing the interaction confidence threshold increases the support of the reference pair but decreases that of the control pair. This is due to the possibility of considering a random reference cancer gene as a non-neighbor with the 0.45 threshold even though it would considered as a neighbor with a lower confidence threshold e.g. 0.35.

### 3.4 ME Evaluations Based on Corrections via MLA

Having compared the ME tests with respect to our novel network-centric evaluation framework, we now assess whether including network knowledge reduces the mutation load confounding (MLC) problem introduced by van de Haar et al. [2019]. van de Haar et al identified a strong negative correlation between the MLAs of genes and their percent significant findings in mutual exclusivity tests. In van de Haar et al. [2019], these statistics are computed for a set of 341 genes from an established cancer gene panel [Cheng et al., 2015] where, for each gene, mutual exclusivity tests are performed with all the other genes in the panel. Here, we first perform a similar analysis where we use the COSMIC CGC database [Forbes et al., 2017] to define the reference cancer gene set as it is more comprehensive and up to date.

Figure 1-A shows the MLA of the reference cancer genes vs the percent significant findings in mutual exclusivity tests performed with DISCOVER for the TCGA COADREAD cohort (498 tumors). We observe a strong negative correlation between MLA values and percent significant findings in mutual exclusivity tests (Pearson correlation −0.88, p-value 3.0*e* 25) similar to van de Haar et al. [2019]. In Figure 1-B, we take into account the PPI information to calculate percent significant findings. Namely, for each CGC gene, we perform mutual exclusivity tests only with its PPI neighbors that are also in CGC. Note that CGC genes which do not have any CGC neighbors are excluded from this analysis. To make a fair comparison between Figure 1-A and 1-B, only the CGC genes that have CGC neighbors are shown in Figure 1-A. We also ensure that the mutual exclusivity of a gene of interest is checked with same sized group of genes in both Figure 1-A and 1-B. To achieve this in Figure 1-A, for each gene, we compute mutual exclusivity with a random subsample of the CGC reference set, the same size as the set of CGC neighbors of that gene. We repeat this random sampling 100 times and plot the mean percent significant findings value. For reference, Supplementary Figure 3-A and 3-D contains versions of Figure 1-A and 1-C, where all CGC genes (i.e., with and without CGC neighbors) are plotted and mutual exclusivities are checked between all CGC pairs, as it was done in van de Haar et al. [2019].

**Figure 1.**
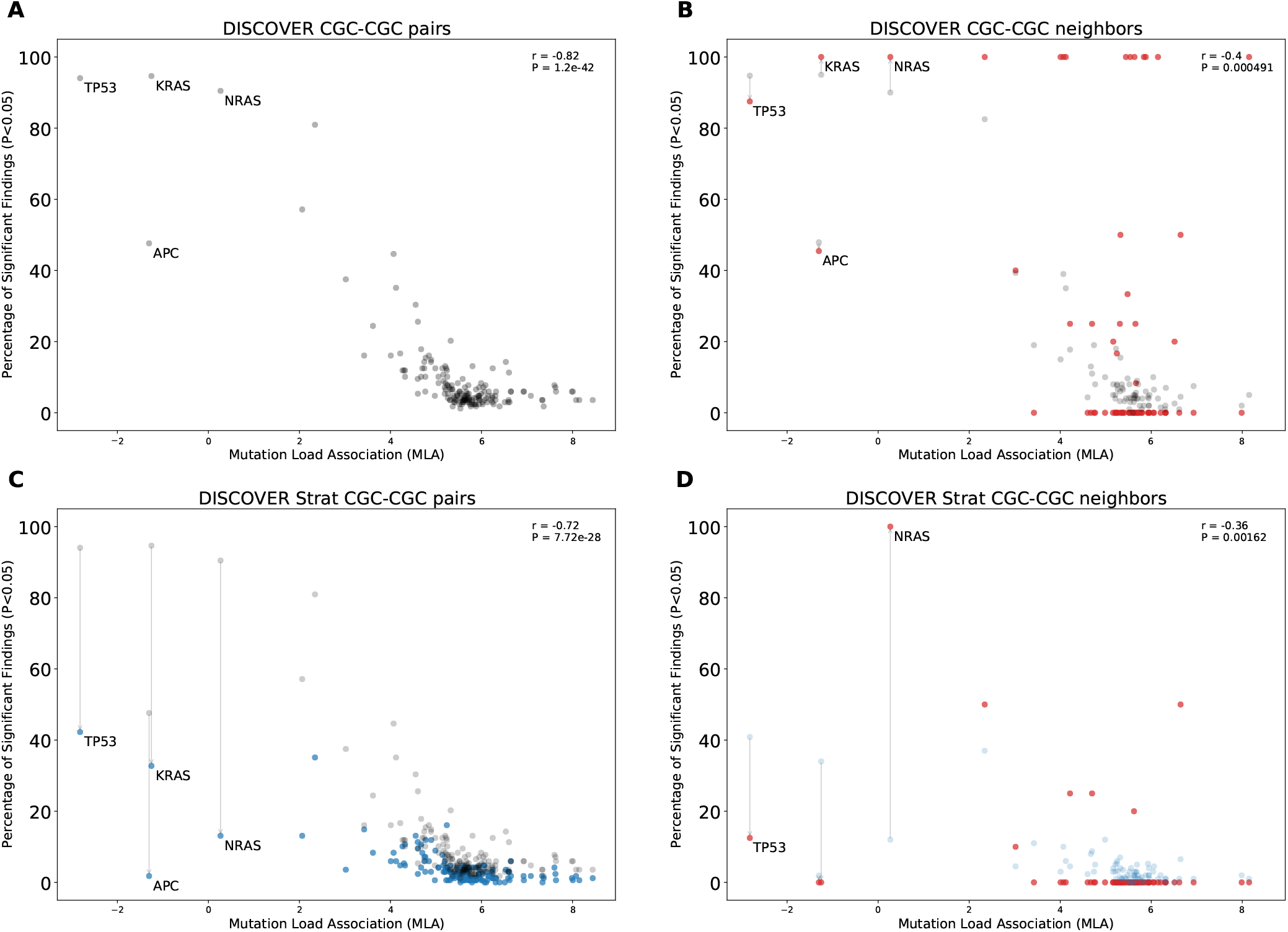
Comparison of mutual exclusivity results of DISCOVER and DISCOVER Strat on COADREAD cohort (498 samples) (A) The scatterplot of percentage significance of mutual exclusivity runs (p-value¡0.05) of DISCOVER on COADREAD data where tests are performed between a CGC gene and all other CGC genes. (B) The scatter plot of percentage significance of mutual exclusivity runs of DISCOVER where tests are performed between a CGC gene and its PPI neighbors that are in CGC (red) compared with (A) in gray. (C) The scatterplot of percentage significance of mutual exclusivity runs of DISCOVER Strat where where tests are performed between a CGC gene and all other CGC genes (blue) compared with the results from (A) in gray. (D) The scatterplot of percentage significance of mutual exclusivity runs of DISCOVER Strat where tests are performed between a CGC gene and its PPI neighbors that are in CGC (red) compared with (C) in blue.

In Figure 1-B, we observe a reduced correlation when network information is included (Pearson correlation −0.4, p-value 4.91*e* 4). We also run DISCOVER Strat where stratification is based on CMS subtypes [Guinney et al., 2015]. We plot these results in Fig 1-C where we again ensure comparability with Fig 1-D where both subtype and network information are considered. Comparing Figure 1-A and Figure 1-C, we verify the findings of van de Haar *et al.*, although with less significance in correlation difference (Pearson correlation −0.73, p-value 1.1*e* 13). It should be noted that the subtype stratification inherently causes an overall decrease in percent significant findings, not specific to genes with low MLA. On the contrary the idea of ME corrections through network incorporation, materialized in the comparison of Figure 1-A and Figure 1-B, inherently leads to an increase in percent significant findings. Most of the decreases occur in genes with small number of CGC neighbors. When we compare Fig 1-D to Fig 1-B, the decrease in correlation from −0.4 to −0.36 indicates that including subtype information is still useful when used on top of network-based corrections we propose.

Supplementary Figure 2 shows the results of the same analysis repeated for BRCA. Similar to the results that we obtain for COADREAD data, including network information reduces the correlation between MLA and ME detection rate (Supp Fig 2-A vs 2-B). The magnitude of reduction is even more significant than what we observe for COADREAD data (Pearson correlation −0.93 vs −0.27). Interestingly, including subtype information results in a very slight decrease in correlation coefficient (−0.93 to −0.92)(Supp Fig 2-A vs Supp Fig 2-C) as opposed to what we observe for COADREAD. We again observe that including subtype information on top of network information results in a negligible decrease in correlation. (Supp Fig 2-B vs 2-D). This difference in the effect of including subtype information for BRCA and COADREAD datasets could be related to the average tumor mutation load of subtypes. BRCA subtypes have comparable average TML values (Her2: 146, LumA:65, LumB: 71, Normal: 55) whereas the CMS1 subtype in COADREAD has a dramatically larger average TML value compared to the other subtypes of COADREAD (CMS1: 1387, CMS2:93, CMS3: 272, CMS4: 212) We repeat the same analysis with the other ME detection methods as well as for other cancer types when *t* is set to 20 (Supplementary Figures 1-8). We observe that the percent significant finding values can vary remarkably across the tumor types. Compared to other cancer types, we observe smaller percent significant findings for LUSC (Supplementary Figs 5 A,D,G). Similarly, very few pairs have percent significance value ≥ 20 when we consider network information in LUSC (Supplementary Figs 5 C,F,I). On the contrary, we observe many pairs with large percent significant values for CGC-CGC neighbors in UCEC data. This is particularly true for DISCOVER and WeXT results (Supplementary Figs 8 C-I).

When we consider the correlation between MLA and percent significant values, we observe that adding network information decreases the correlation coefficient values for all cancer types and for all ME detection methods except for Fisher’s Exact Test. Fisher’s Exact Test results show an increased correlation with the addition of network information for LUSC and SKCM (Supplementary Fig. 5-6 D-F). Also, the correlation coefficient can not be computed for LUAD and STAD since Fisher’s Exact Test gives a value of 0 for the percent significant findings of all considered genes (Supplementary Fig. 4-7 D-F). Another interesting observation is the variance in magnitude of decrease in correlation values across different tumor types. In particular, we observe a smaller decrease in correlation values for LUAD compared to other cancer types. The analogous results are also available for *t* = 5 setting (Supplementary Figs 9-15). For all the cancer types, the correlation between MLA values and percent significant findings decreases and becomes non-significant for most cases.

We should also note that the majority of CGC genes have only one neighbor within the data setting of the cancer type under consideration. This leads to percentage significant findings of either 0 or 1 in many cases simply because these are the only possible values; for COADREAD see 1-B and 1-D where 41 out of 74 genes under study have only one CGC neighbor in the COADREAD data settings. To avoid any such possible biases, we repeat the same evaluations after filtering out those CGC genes with only one neighbor. The evaluations still provide significant decreases in correlation coefficient values analogous to the decreases observed in 1-B as compared to 1-A and 1-D as compared to 1-C. For detailed results, see Supplementary Figures 17-24 for *t* = 20 and Supplementary Figures 25-32 for *t* = 5.

Individual genes of interest are those that have increased percent significant findings when network neigborhood information is incorporated while at the same have significant number of CGC neighbors. More specifically, for the former constraint, we identify the CGC genes with at least 0.1 increase in percentage of significant findings value of WeXT, DISCOVER and MEMo when the network information is included as opposed to the scenario when it is not (e.g. for COADREAD, Figure 1-A vs Figure 1-B). We choose these three ME methods since they are top performers based on the defined metrics in Section 2.1. For STAD, SKCM and UCEC, since MEMo results are unavailable, we only consider WeXT and DISCOVER results. For the second constraint, we include the CGC genes with at least 3 CGC neighbors. For COADREAD, this selection procedure results in four genes: EP300, CREBBP, NCOA2 and NCOR2. Among these, EP300 is a well-known tumor suppressor in epithelial cancer types including COADREAD [Gayther et al., 2000]. For BRCA, the only identified gene is PIK3R1. PIK3R1 is found to be significantly mutually exclusive with PIK3CA and SPEN based on both WeXT, DISCOVER and MEMo results. PIK3R1 and PIK3CA are members of the PI3K pathway and their mutual exclusivity has been previously established in the literature [Chen et al., 2018]. For LUAD, PTPRB is the only identified gene and is found to be mutually exclusive with EGFR, a well-known oncogene in non-small cell lung cancer [Bethune et al., 2010]. The set of identified genes for STAD are NCOA2, NCOR2 and CREBBP; all of which are found to be mutually exclusive with TP53. For SKCM, we identify ERBB4, RAC1, EP300 and ITK. ERBB4 is a well-known oncogene in skin cancer and found to be mutually exclusive with ERBB2 [Prickett et al., 2009, Nielsen et al., 2014]. ERBB2 and ERBB4 indeed belong to the same family (i.e. ErbB family of receptor tyrosine kinases) and form a heterodimer receptor for Heparin-binding EGF-like growth factor (HB-EGF) [Iwamoto et al., 2017]. RAC1 mutation P29S is an established driver in melanoma [Jiang et al., 2018]. RAC1 is found to be mutually exclusive with MYH9, a tumor suppressor in melanoma [Singh et al., 2020]. Lastly, ITK has been shown to be an oncogene in melanoma [Carson et al., 2015]. For UCEC, we identify 33 genes in total. Among these, KIT and PTEN have established roles in UCEC cancer development [Chang et al., 2015, Wang et al., 2020]. Moreover, PTEN is found to be strongly mutually exclusive with SPOP, whose mutations are also associated with endometrial cancer [Clark and Burleson, 2020]. Lastly, for BLCA and LUSC, no gene satisfies the abovementioned criteria. Overall these results suggest that the CGC genes that show increased ME with network incorporation as well as their mutually exclusive partner genes often have established roles in the development of the particular cancer type.

### 3.5 ME Evaluations Based on Corrections via TSN

We first provide our ME evaluations with respect to the metrics defined in Section 2.1 by replacing the non-specific networks with TSNs. We provide two types of comparisons; one where we compare TSN_0.5_ with the original non-tissue specific Intact network and one where results of TSN_0.5_ are compared against TSN_0_. We do the latter to avoid artifacts that may be introduced due to the fact that some genes in the original Intact network might be simply missing from even TSN_0_ since they may be nonexistent in the GTEX database. For the BLCA dataset, comparing the F1 scores of the ME methods under TSN_0_ and TSN_0.5_ settings, we observe that the scores of all methods are higher for the latter network. The largest percent increase of 10% is observed for WeXT when the control group is *χ*_1_. Similarly, the largest percent increase of 12% is observed for MEMo when the control group is *χ*_2_. On the other hand, when we compare the scores of TSN_0_ against the original network, the differences are negligible. The next largest difference between the F1 scores obtained under under TSN_0.5_ as compared to TSN_0_ is observed in STAD where we see a 7% increase in DISCOVER’s score for *χ*_1_, and a 10% increase in WeXT’s score for *χ*_2_. For the rest of the cancer types under study, for LUSC and UCEC we observe slight increase in performances of all the ME methods comparing the metrics under TSN_0.5_ against TSN_0_. For COADREAD, BRCA and SKCM we observe both increases and decreases in performances but the differences are almost negligible; see Supplementary Tables 50-81 for detailed results.

Figure 2 compares the ROC curves of CGC gene pairs and non-CGC gene pairs for COADREAD data where mutual exclusivities are estimated with DISCOVER, DISCOVER Strat, Fisher’s Exact Test, MEGSA, MEMo and WeXT with t = 20. We observe that all the ME methods estimate stronger mutual exclusivities for tissue-specific CGC gene pairs compared to non-tissue-specific CGC gene pairs since AUROCs are greater than 0.5. Additionally, we observe much smaller AUROCs for the control group where we repeat the same analysis with non-CGC gene pairs. Analogous results are available for the other cancer types where both the positive and negative set contains at least 10 number of pairs when t is set to 20. (Supplementary Figs. 33-35). We observe a similar result for SKCM where CGC pairs result in larger AUROCs compared to non-CGC pairs for all ME methods (Supplementary Fig. 34). We observe a steep increase in the ROC curves plotted for MEGSA results. This is due to the utilized likelihood ratio test that results in a p-value of 0.5 when the likelihood values are equal to each other. For UCEC, we see a significant difference between the ROC curves of CGC-pairs vs non-CGC pairs for Fisher’s Exact Test and MEGSA; whereas the corresponding difference is negligible for DISCOVER and WeXT.

**Figure 2.**
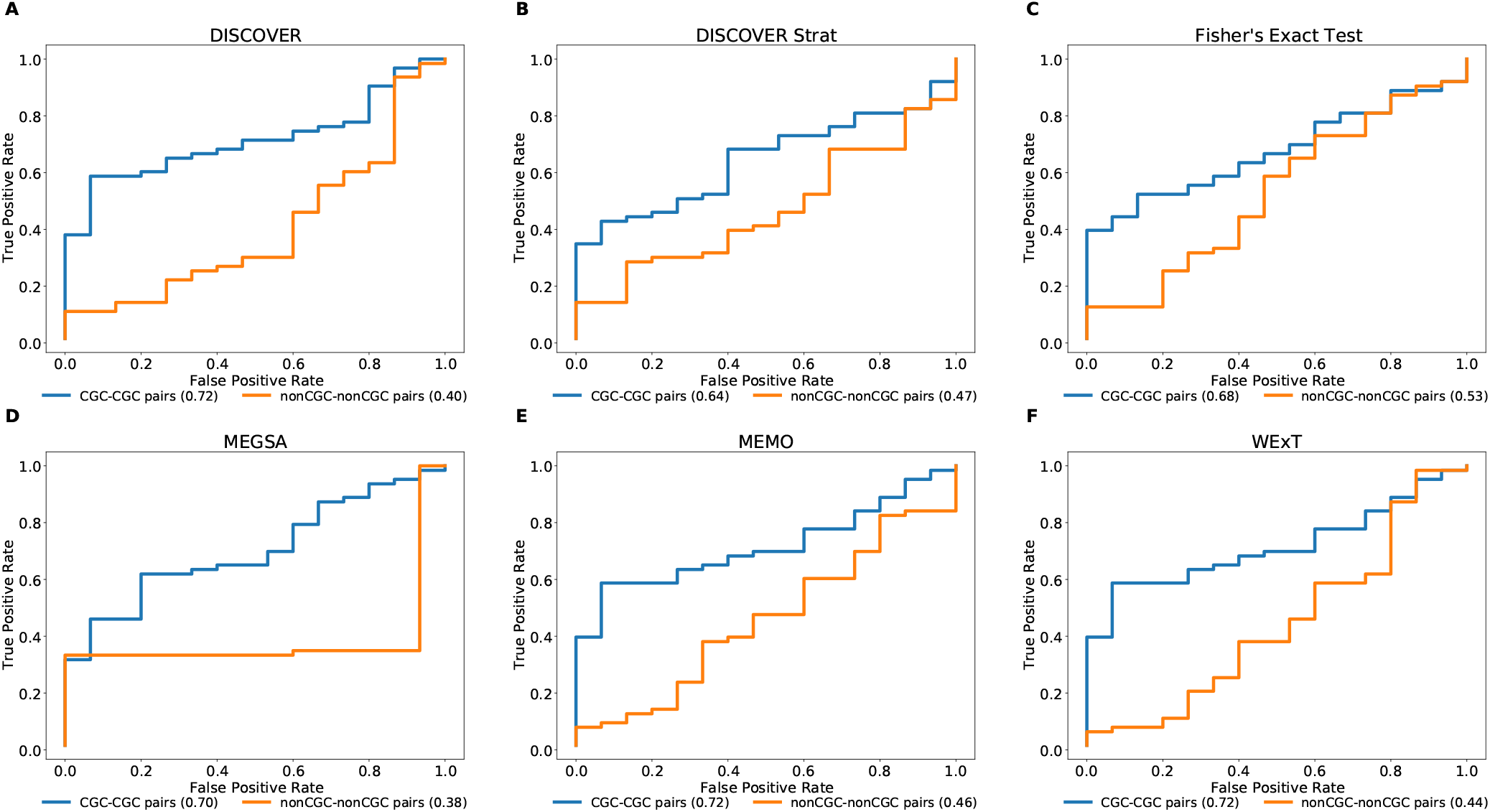
Performance of selected ME tests in terms of discriminating TSN and non-TSN gene pairs based on estimated ME p-values on COADREAD data. Blue curve is plotted with CGC gene pairs and red curve is plotted with non-CGC gene pairs. Mutual exclusivities are estimated with a) DISCOVER, b) DISCOVER Strat, c) Fisher’s Exact Test, d) MEGSA, e) MEMo and g) WeXT respectively.

## 4 CASE STUDY

Apart from the defined network-centric ME evaluation framework, we discuss a case study where we assess whether mutual exclusivities estimated by the considered ME methods improve the performance of driver identification methods that utilize mutual exclusivity information. To this end, we compare the original version of MEXCOwalk with its alternatives where mutual exclusivity estimates are provided by the employed ME methods. Assuming that *g_i_* and *g_j_* genes are mutated in patient sets *S_i_* and *S_j_*, respectively; MEXCOWalk simply computes the mutual exclusivity between these two genes with the following formula: |*S_i_ ∪ S_j_|/*(|*S_i_*| + |*S_j_*|). MEXCOwalk uses the estimated mutual exclusivity values as part of edge weights. As such, to utilize the p-values output by ME detection methods in MEXCOwalk, we first compute −log(p-value) and then convert the resulting values between 0 and 1. To this end, we replace all −log(p-value)’s larger than 10 with 1. We then find the maximum −log(p-value) less than 10 and divide all other −log(p-value)’s with this value. The reason why we set a threshold for finding the maximum is the large differences across the smallest p-values output by different ME methods. For instance, WeXT outputs a very large range of p-values and if we use the smallest p-value to scale, all other −log(p-value)s will be converted to values that are very close to 0. In the original MEXCOWalk study, a threshold of 0.7 is applied to ME values such that all values ≤0.7 are clamped to 0. This conversion is equivalent to removing those edges from the network since the edge weights include a multiplicative term for ME values. We find that the removal of these edges correspond to a 0.035 percent reduction in graph density. For the current analysis, we determine the threshold value for each ME detection method to achieve the same percent density reduction in the graph. Figure 3 shows the number of recovered CGC genes for fixed output gene sizes from 100 to 2500 as a ROC curve for original MEXCOwalk as well as for versions of MEXCOwalk where mutual exclusivity values are estimated with DISCOVER, Fisher’s Exact Test and WeXT, respectively. We observe that MEXCOwalk with WeXT’s ME values results in the best AUROC value for COADREAD. Supplementary Figure 36 shows the analogous results for the other cancer types. For, LUSC, STAD and UCEC, MEXCOwalk with DISCOVER gives the best AUROC whereas for BLCA, LUAD and SKCM MEXCOwalk with Fisher’s Exact Test performs the best. An important observation is the worse performance of MEXCOwalk with Fisher’s Exact Test compared to the original MEXCOwalk for COADREAD, STAD and UCEC. As such, using Fisher’s Exact Test in place of MEXCOwalk’s original ME values does have the potential to decrease the performance whereas for the other ME methods we do not observe such a risk. Note that for these analysis we employ *t* = 5 since *t* = 20 filtering does not provide enough number of genes to be evaluated.

**Figure 3.**
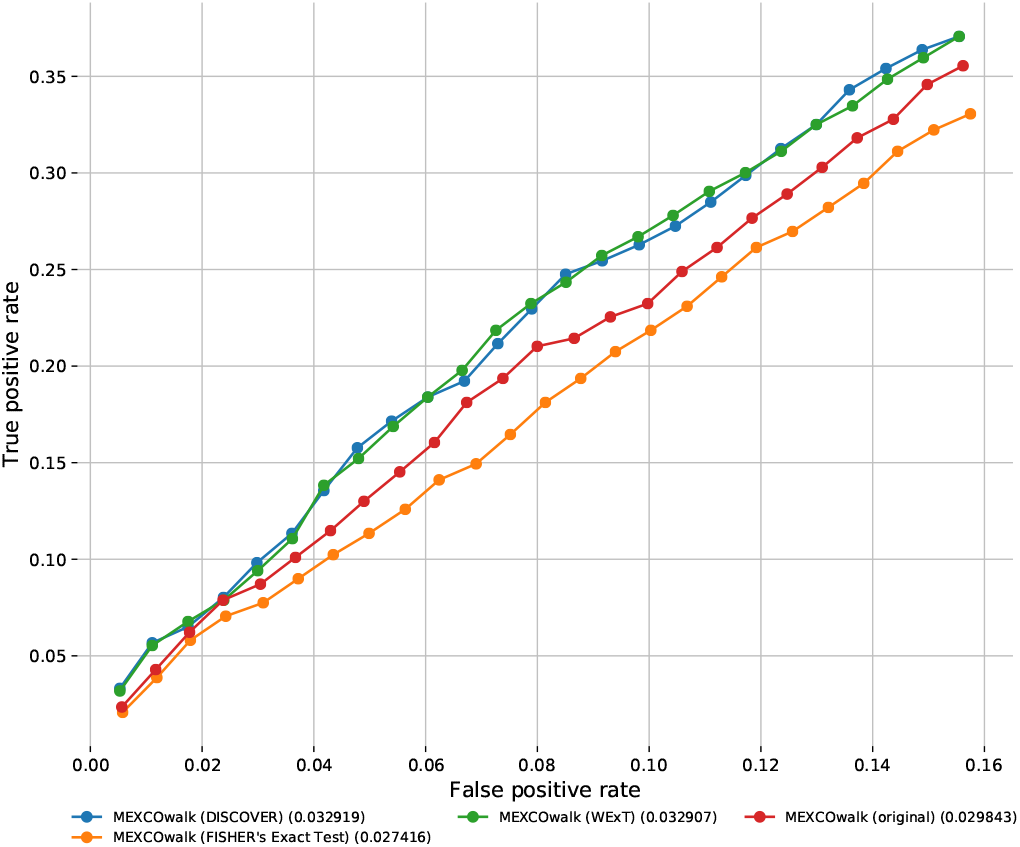
The number of recovered CGC genes for the original MEXCOwalk as well as for its modified versions where mutual exclusivity values are estimated with DISCOVER, Fisher’s Exact Test and WeXT. COADREAD dataset is used with *t* = 5 setting. The numbers in parentheses indicate the area under the ROC curve for the corresponding curve.

## 5 DISCUSSION

It is important to investigate whether the employment of an interaction network within our ME evaluation framework causes any ascertainment bias in the findings and to elaborate on how any such potential bias is mediated within the framework. It is established that known cancer genes have larger number of interactions compared to other genes in the network [Hou and Ma, 2014]. This implies a potential bias that needs to be resolved in cancer driver gene identification methods employing interaction network data. Such a bias is less of a problem for the current study, since our aim is not to identify novel cancer driver genes but to utilize the interaction network and known cancer genes to form a ground truth of mutually exclusive interactions for evaluating existing ME methods. On the contrary, the fact that most known cancer genes have well-characterized interactions in the network provides a benefit for our work as it supports the confidence of our true positive examples. Additionally, our framework makes use of not only genes from the reference set 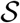 but also genes not in 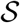 to create random controls. Nevertheless, the fact that some known cancer genes have significantly larger number of interactions compared to other known cancer genes could lead to a bias. For instance, for our analysis of the COADREAD data (*t* = 20, 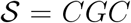), there are 74 CGC genes among which five CGC genes have more than ten CGC neighbors whereas 41 have exactly one CGC neighbor. This could lead to a bias as CGC genes with large number of CGC neighbors contribute to the aggregate statistics and metrics much more than those CGC genes with small number of CGC neighbors. To mediate for this bias, our framework includes additional results where all the statistics and the traditional measures such as the F1 score are calculated in a degree-normalized way for each gene and the gene-level results are then aggregated by taking an average across the genes. These results are available in the Supplementary Document; see Table S12, S24, S36, and S48. To summarize, the degree-normalized results are in agreement with those of the previous settings in almost all the cases in terms of ranking based on F1 score.

Another important point worth emphasizing is that apart from the aggregate statistics provided in the previous sections as part of the metrics for the network-centric ME evaluations, our proposed framework also provides analogous statistics at the gene-level as well. Such statistics may in fact be of more interest to cancer biologists than the aggregate statistics in certain cases. Several interesting observations can be made through an inspection of these gene-level evaluations, especially for the settings where the conventionally defined F1 score fails in quantifying ME. Genes with low MLA comprise an example setting, where TP53 is a leading member. Consider the case of TP53 in COADREAD evaluations for instance. With respect to the degree-normalized setting, the values of precision, sensitivity, precision_*strict*_ and sensitivity_*strict*_ for WeXT are respectively 0.5, 1, 0.25, 0.25 which gives rise to an F1 score of 0.66 and F1_*strict*_ score of 0.25. On the other hand, MEMo provides the same precision, sensitivity and F1 scores as WeXT whereas its precision_*strict*_, sensitivity_*strict*_ and F1_*strict*_ scores are all 0. To summarize, although the inspection of the F1 scores does not provide a distinction between the two results, an inspection of the F1_*strict*_ scores establishes that MEMo is worse than WeXT in this setting. We note that the advantages of inspections based on the strict definitions of the metrics rather than the conventional ones are also apparent in the aggregate analysis as well. In addition to the COADREAD evaluations shown in Table2, BRCA also contains an example instance where the conventional and the strict versions of the metrics provide different conclusions; see Table S3-b. In terms of the F1 scores WeXT and DISCOVER Strat obtain very close values with WeXT providing slightly better results. However comparing F1_*strict*_ scores reveals DISCOVER Strat a better ME test candidate for this instance. Also, overall we observe that MEMo’s performance gets severely affected when the strict versions of the metrics are employed.

Lastly, our robustness analysis results reveal some suggestions for potential users of our framework. We recommend using a p-value threshold greater than 0.05 as lower threshold values are too stringent and lead to too few predicted positives. Regarding *robustness iterations*, we tested values both smaller than and higher than the default value of 100 for COADREAD evaluations: 5, 50, 100, 300 and 500. We repeated each experiment 20 times and calculated the standard deviation of the obtained set of F1 and F1_*strict*_ scores. For the majority of the cases, we observe a large decrease in the standard deviation values when *robustness iterations* is increased from 5 to 50. (Table S82). This analysis suggests that the *robustness iterations* should be set to a at least 50. Lastly, we observe that different PPI networks can lead to large differences in both the F1/F1_*strict*_ scores and the ranking of the methods. As such, exploring different PPI sources would be beneficial.

## 6 CONCLUSION

We propose a network-centric framework to evaluate pairwise mutual exclusivity findings reported by different ME algorithms. The first component of our framework consists of useful definitions of statistics employed in the network-centric ME evaluations. We observe that for the majority of the cancer types under study WeXT outperforms the other methods in terms of F1 score measured with respect to appropriately defined control groups. In half of the cancer types DISCOVER and in the other half MEMo perform as the second best methods. When comparing different cancer types we observe that BRCA and COADREAD are among the top two types leading to maximum F1 scores with at least one of the ME methods providing a score greater than 0.5. We note that DISCOVER Strat is only applicable in two cancer types among a total of eight since these are the only cancer types with well-defined subtypes. Furthermore, among these two cancer types, DISCOVER Strat outperforms original DISCOVER algorithm in BRCA, whereas it is the second worst method after Fisher’s Exact Test in COADREAD. This is noteworthy since van de Haar *et al.* propose subtype stratification as employed by DISCOVER Strat as a way to emphasize true mutual exclusivity by reducing mutation load confounding [van de Haar et al., 2019]. We also observe that Fisher’s exact test and MEGSA are more conservative compared to DISCOVER and WeXT, where from the latter group, WeXT outputs notably larger number of significant p-values. The second component of our framework evaluates ME tests by comparing two types of measures obtained with and without network information. First measure is with respect to the percent significant findings of mutually exclusive gene pairs, whereas the second is based on MLC values. In most of the cancer types and for most of the genes we observe an increase with respect to the former whereas a decrease with respect to the latter measure. Finally, we repeat the same analysis by considering TSNs in the network-centric framework. Considerable improvements achieved due to the use of TSNs as opposed tissue nonspecific interaction network are only observed for BLCA and STAD datasets. A more detailed analysis in terms of comparing ROCs of CGC gene pairs and non-CGC gene pairs on cancer types with considerable number of tissue-specific gene pairs indicate the advantages of employing tissue specificity in detecting mutual exclusivity in COADREAD, SKCM, and UCEC. Finally we extend out network-centric evaluation framework to assess whether including network knowledge reduces the mutation load confounding problem.

As noted earlier the proposed framework is intended for the network-centric evaluations of mutual exclusivities of pairs of genes rather than groups of genes. Such a choice stems form the fact that the mutual exclusivities are commonly made use of in driver gene/module identification algorithms which mostly employ pairwise mutual exclusivities. Furthermore the extensive evaluation settings proposed, the number of ME methods under study and their own computational requirements, and the potentially exponential computational complexity inherent in handling groups of genes limits the scope of the current study to evaluations of pairwise ME scorings. Nonetheless most statistical ME methods are capable of providing ME results for groups of genes as well. Regarding the ME tests considered in this study, the main ME test provided by DISCOVER is based on a pairwise test definition but it also extends the definition for possible use in quantifying the ME of a group of genes, although the experiments involving the latter are based only on simulation data. The remaining tests MEGSA, MEMo, and WeXT are all ME tests specifically designed for groups of genes. An important direction for future work is to design a suitable extension of the proposed network-centric framework to evaluate the results of ME tests on groups of genes. Design choices relevant for such an extension would involve an appropriate and computationally efficient definition of the reference groups of genes analogous to a pair of interacting genes from the set S in the current setting and the definitions of control groups analogous to *χ*_1_ and *χ*_2_.

## 7 ADDITIONAL REQUIREMENTS

For additional requirements for specific article types and further information please refer to Author Guidelines.

## AUTHOR CONTRIBUTIONS

Authors names are written in alphabetical order. C. E. and H.K. conceived the idea and supervised the study. R. A implemented the code and performed the initial experiments. C. Y. and A.H. repeated the experiments for other cancer types. All authors contributed to the preparation of the manuscript. All authors read and approved the manuscript.

## ACKNOWLEDGMENTS

We thank Joris van de Haar for sharing us the data used in van de Haar et al. [2019].

## SUPPLEMENTAL DATA

Supplementary Material should be uploaded separately on submission, if there are Supplementary Figures, please include the caption in the same file as the figure. LaTeX Supplementary Material templates can be found in the Frontiers LaTeX folder.

## DATA AVAILABILITY STATEMENT

The datasets analyzed for this study can be found on the following repository: https://github.com/abu-compbio/NetCentric.

