## Supplementary Material for "A Network-centric Framework for the Evaluation of Mutual Exclusivity Tests on Cancer Drivers"

Table 1: **Results of network-centric ME evaluation framework with $\mathcal{G}$ = Intact (w conf. threshold 0.35), $\mathcal{S}$ = CGC, c= $X_1$, $p_t$ = 0.05, t=20, robustness_iterations = 100**

(a) Metrics for BLCA data. (411 samples | 56 CGC-CGC pairs)

| Method | Precision | Sensitivity | F1 Score | $\text{Precision}_{\text{strict}}$ | $\text{Sensitivity}_{\text{strict}}$ | $\text{F1 Score}_{\text{strict}}$ |
|---|---|---|---|---|---|---|
| DISCOVER | 0.800 | 0.075 | 0.137 | 0.800 | 0.075 | 0.137 |
| Fisher's Exact Test | 1.000 | 0.036 | 0.069 | 1.000 | 0.036 | 0.069 |
| MEGSA | 1.000 | 0.074 | 0.138 | 1.000 | 0.074 | 0.138 |
| MEMO | 0.800 | 0.079 | 0.144 | 0.800 | 0.079 | 0.144 |
| WExT | 0.571 | 0.077 | 0.136 | 0.571 | 0.077 | 0.136 |

(b) Metrics for BRCA data. (1026 samples | 34 CGC-CGC pairs)

| Method | Precision | Sensitivity | F1 Score | $\text{Precision}_{\text{strict}}$ | $\text{Sensitivity}_{\text{strict}}$ | $\text{F1 Score}_{\text{strict}}$ |
|---|---|---|---|---|---|---|
| DISCOVER | 0.609 | 0.452 | 0.519 | 0.579 | 0.355 | 0.440 |
| DISCOVER Strat | 0.744 | 0.492 | 0.593 | 0.811 | 0.462 | 0.589 |
| Fisher's Exact Test | 1.000 | 0.061 | 0.114 | 1.000 | 0.061 | 0.115 |
| MEGSA | 1.000 | 0.059 | 0.111 | 1.000 | 0.059 | 0.111 |
| MEMO | 0.700 | 0.483 | 0.571 | 0.706 | 0.414 | 0.522 |
| WExT | 0.612 | 0.526 | 0.566 | 0.615 | 0.421 | 0.500 |

(c) Metrics for COADREAD data. (498 samples | 196 CGC-CGC pairs)

| Method | Precision | Sensitivity | F1 Score | $\text{Precision}_{\text{strict}}$ | $\text{Sensitivity}_{\text{strict}}$ | $\text{F1 Score}_{\text{strict}}$ |
|---|---|---|---|---|---|---|
| DISCOVER | 0.661 | 0.220 | 0.331 | 0.708 | 0.183 | 0.291 |
| DISCOVER Strat | 0.727 | 0.041 | 0.078 | 0.727 | 0.041 | 0.078 |
| Fisher's Exact Test | 0.500 | 0.031 | 0.058 | 0.500 | 0.031 | 0.058 |
| MEGSA | 0.611 | 0.056 | 0.103 | 0.588 | 0.051 | 0.094 |
| MEMO | 0.658 | 0.329 | 0.439 | 0.647 | 0.237 | 0.347 |
| WExT | 0.676 | 0.403 | 0.505 | 0.725 | 0.329 | 0.453 |

(d) Metrics for LUAD data. (568 samples | 92 CGC-CGC pairs)

| Method | Precision | Sensitivity | F1 Score | $\text{Precision}_{\text{strict}}$ | $\text{Sensitivity}_{\text{strict}}$ | $\text{F1 Score}_{\text{strict}}$ |
|---|---|---|---|---|---|---|
| DISCOVER | 0.773 | 0.099 | 0.176 | 0.789 | 0.088 | 0.158 |
| Fisher's Exact Test | 0.000 | 0.000 | NaN | 0.000 | 0.000 | NaN |
| MEGSA | 0.667 | 0.022 | 0.043 | 0.667 | 0.022 | 0.043 |
| MEMO | 0.722 | 0.149 | 0.248 | 0.733 | 0.126 | 0.215 |
| WExT | 0.625 | 0.174 | 0.273 | 0.667 | 0.163 | 0.262 |

(e) Metrics for LUSC data. (485 samples | 38 CGC-CGC pairs)

| Method | Precision | Sensitivity | F1 Score | $\text{Precision}_{\text{strict}}$ | $\text{Sensitivity}_{\text{strict}}$ | $\text{F1 Score}_{\text{strict}}$ |
|---|---|---|---|---|---|---|
| DISCOVER | 1.0 | 0.054 | 0.103 | 1.0 | 0.054 | 0.102 |
| Fisher's Exact Test | 1.0 | 0.053 | 0.100 | 1.0 | 0.053 | 0.101 |
| MEGSA | 1.0 | 0.158 | 0.273 | 1.0 | 0.158 | 0.273 |
| MEMO | 1.0 | 0.108 | 0.195 | 1.0 | 0.108 | 0.195 |
| WExT | 1.0 | 0.114 | 0.205 | 1.0 | 0.114 | 0.205 |

(f) Metrics for SKCM data. (468 samples | 458 CGC-CGC pairs)

| Method | Precision | Sensitivity | F1 Score | $\text{Precision}_{\text{strict}}$ | $\text{Sensitivity}_{\text{strict}}$ | $\text{F1 Score}_{\text{strict}}$ |
|---|---|---|---|---|---|---|
| DISCOVER | 0.800 | 0.045 | 0.084 | 0.833 | 0.045 | 0.085 |
| Fisher's Exact Test | 1.000 | 0.004 | 0.009 | 1.000 | 0.004 | 0.008 |
| MEGSA | 0.889 | 0.018 | 0.034 | 0.889 | 0.018 | 0.035 |
| WExT | 0.717 | 0.121 | 0.207 | 0.730 | 0.116 | 0.200 |

(g) Metrics for STAD data. (438 samples | 140 CGC-CGC pairs)

| Method | Precision | Sensitivity | F1 Score | $\text{Precision}_{\text{strict}}$ | $\text{Sensitivity}_{\text{strict}}$ | $\text{F1 Score}_{\text{strict}}$ |
|---|---|---|---|---|---|---|
| DISCOVER | 0.667 | 0.122 | 0.206 | 0.684 | 0.099 | 0.173 |
| Fisher's Exact Test | 0.000 | 0.000 | NaN | 0.000 | 0.000 | NaN |
| MEGSA | 0.667 | 0.014 | 0.028 | 0.667 | 0.014 | 0.027 |
| WExT | 0.634 | 0.190 | 0.292 | 0.636 | 0.153 | 0.247 |

(h) Metrics for UCEC data. (531 samples | 1356 CGC-CGC pairs)

| Method | Precision | Sensitivity | F1 Score | $\text{Precision}_{\text{strict}}$ | $\text{Sensitivity}_{\text{strict}}$ | $\text{F1 Score}_{\text{strict}}$ |
|---|---|---|---|---|---|---|
| DISCOVER | 0.651 | 0.177 | 0.279 | 0.711 | 0.143 | 0.238 |
| Fisher's Exact Test | 0.833 | 0.007 | 0.015 | 0.833 | 0.007 | 0.014 |
| MEGSA | 0.786 | 0.008 | 0.016 | 0.786 | 0.008 | 0.016 |
| WExT | 0.616 | 0.278 | 0.383 | 0.665 | 0.227 | 0.338 |

Table 2: **Results of network-centric ME evaluation framework with $\mathcal{G}$ = Intact (w conf. threshold 0.35), $\mathcal{S}$ = CGC, c= $X_1$, $p_t$ = 0.05, t=20, robustness_iterations = 300**

(a) Metrics for BLCA data. (411 samples | 56 CGC-CGC pairs)

| Method | Precision | Sensitivity | F1 Score | $\text{Precision}_{\text{strict}}$ | $\text{Sensitivity}_{\text{strict}}$ | $\text{F1 Score}_{\text{strict}}$ |
|---|---|---|---|---|---|---|
| DISCOVER | 0.800 | 0.075 | 0.138 | 0.800 | 0.075 | 0.137 |
| Fisher's Exact Test | 1.000 | 0.037 | 0.071 | 1.000 | 0.037 | 0.071 |
| MEGSA | 1.000 | 0.073 | 0.136 | 1.000 | 0.073 | 0.136 |
| MEMO | 0.667 | 0.078 | 0.140 | 0.667 | 0.078 | 0.140 |
| WExT | 0.571 | 0.075 | 0.133 | 0.571 | 0.075 | 0.133 |

(b) Metrics for BRCA data. (1026 samples | 34 CGC-CGC pairs)

| Method | Precision | Sensitivity | F1 Score | $\text{Precision}_{\text{strict}}$ | $\text{Sensitivity}_{\text{strict}}$ | $\text{F1 Score}_{\text{strict}}$ |
|---|---|---|---|---|---|---|
| DISCOVER | 0.652 | 0.484 | 0.556 | 0.632 | 0.387 | 0.480 |
| DISCOVER Strat | 0.762 | 0.516 | 0.615 | 0.833 | 0.484 | 0.612 |
| Fisher's Exact Test | 1.000 | 0.062 | 0.118 | 1.000 | 0.062 | 0.117 |
| MEGSA | 1.000 | 0.059 | 0.111 | 1.000 | 0.059 | 0.111 |
| MEMO | 0.667 | 0.467 | 0.549 | 0.667 | 0.400 | 0.500 |
| WExT | 0.600 | 0.536 | 0.566 | 0.632 | 0.429 | 0.511 |

(c) Metrics for COADREAD data. (498 samples | 196 CGC-CGC pairs)

| Method | Precision | Sensitivity | F1 Score | $\text{Precision}_{\text{strict}}$ | $\text{Sensitivity}_{\text{strict}}$ | $\text{F1 Score}_{\text{strict}}$ |
|---|---|---|---|---|---|---|
| DISCOVER | 0.659 | 0.218 | 0.327 | 0.707 | 0.184 | 0.292 |
| DISCOVER Strat | 0.727 | 0.042 | 0.079 | 0.727 | 0.042 | 0.079 |
| Fisher's Exact Test | 0.545 | 0.031 | 0.059 | 0.545 | 0.031 | 0.059 |
| MEGSA | 0.611 | 0.056 | 0.103 | 0.588 | 0.051 | 0.094 |
| MEMO | 0.660 | 0.338 | 0.447 | 0.662 | 0.247 | 0.360 |
| WExT | 0.684 | 0.406 | 0.510 | 0.724 | 0.328 | 0.451 |

(d) Metrics for LUAD data. (568 samples | 92 CGC-CGC pairs)

| Method | Precision | Sensitivity | F1 Score | $\text{Precision}_{\text{strict}}$ | $\text{Sensitivity}_{\text{strict}}$ | $\text{F1 Score}_{\text{strict}}$ |
|---|---|---|---|---|---|---|
| DISCOVER | 0.750 | 0.103 | 0.182 | 0.700 | 0.080 | 0.144 |
| Fisher's Exact Test | 0.000 | 0.000 | NaN | 0.000 | 0.000 | NaN |
| MEGSA | 0.667 | 0.022 | 0.043 | 0.667 | 0.022 | 0.043 |
| MEMO | 0.722 | 0.149 | 0.248 | 0.733 | 0.126 | 0.215 |
| WExT | 0.667 | 0.179 | 0.282 | 0.700 | 0.156 | 0.255 |

(e) Metrics for LUSC data. (485 samples | 38 CGC-CGC pairs)

| Method | Precision | Sensitivity | F1 Score | $\text{Precision}_{\text{strict}}$ | $\text{Sensitivity}_{\text{strict}}$ | $\text{F1 Score}_{\text{strict}}$ |
|---|---|---|---|---|---|---|
| DISCOVER | 1.0 | 0.054 | 0.103 | 1.0 | 0.054 | 0.102 |
| Fisher's Exact Test | 1.0 | 0.053 | 0.100 | 1.0 | 0.053 | 0.101 |
| MEGSA | 1.0 | 0.158 | 0.273 | 1.0 | 0.158 | 0.273 |
| MEMO | 1.0 | 0.111 | 0.200 | 1.0 | 0.111 | 0.200 |
| WExT | 1.0 | 0.111 | 0.200 | 1.0 | 0.111 | 0.200 |

(f) Metrics for SKCM data. (468 samples | 458 CGC-CGC pairs)

| Method | Precision | Sensitivity | F1 Score | $\text{Precision}_{\text{strict}}$ | $\text{Sensitivity}_{\text{strict}}$ | $\text{F1 Score}_{\text{strict}}$ |
|---|---|---|---|---|---|---|
| DISCOVER | 0.800 | 0.045 | 0.084 | 0.800 | 0.045 | 0.085 |
| Fisher's Exact Test | 1.000 | 0.004 | 0.009 | 1.000 | 0.004 | 0.008 |
| MEGSA | 0.889 | 0.017 | 0.034 | 0.889 | 0.017 | 0.033 |
| WExT | 0.708 | 0.117 | 0.201 | 0.721 | 0.113 | 0.195 |

(g) Metrics for STAD data. (438 samples | 140 CGC-CGC pairs)

| Method | Precision | Sensitivity | F1 Score | $\text{Precision}_{\text{strict}}$ | $\text{Sensitivity}_{\text{strict}}$ | $\text{F1 Score}_{\text{strict}}$ |
|---|---|---|---|---|---|---|
| DISCOVER | 0.682 | 0.112 | 0.192 | 0.667 | 0.090 | 0.159 |
| Fisher's Exact Test | 0.000 | 0.000 | NaN | 0.000 | 0.000 | NaN |
| MEGSA | 0.667 | 0.014 | 0.028 | 0.667 | 0.014 | 0.027 |
| WExT | 0.634 | 0.191 | 0.294 | 0.636 | 0.154 | 0.248 |

(h) Metrics for UCEC data. (531 samples | 1356 CGC-CGC pairs)

| Method | Precision | Sensitivity | F1 Score | $\text{Precision}_{\text{strict}}$ | $\text{Sensitivity}_{\text{strict}}$ | $\text{F1 Score}_{\text{strict}}$ |
|---|---|---|---|---|---|---|
| DISCOVER | 0.652 | 0.178 | 0.279 | 0.712 | 0.143 | 0.238 |
| Fisher's Exact Test | 0.833 | 0.007 | 0.015 | 0.833 | 0.007 | 0.014 |
| MEGSA | 0.786 | 0.008 | 0.016 | 0.786 | 0.008 | 0.016 |
| WExT | 0.615 | 0.277 | 0.382 | 0.663 | 0.226 | 0.337 |

Table 3: **Results of network-centric ME evaluation framework with $\mathcal{G}$ = Intact (w conf. threshold 0.35), $\mathcal{S}$ = CGC, c= $X_1$, $p_t$ = 0.05, t=20, robustness_iterations = 500**

(a) Metrics for BLCA data. (411 samples | 56 CGC-CGC pairs)

| Method | Precision | Sensitivity | F1 Score | $\text{Precision}_{\text{strict}}$ | $\text{Sensitivity}_{\text{strict}}$ | $\text{F1 Score}_{\text{strict}}$ |
|---|---|---|---|---|---|---|
| DISCOVER | 0.800 | 0.075 | 0.138 | 0.800 | 0.075 | 0.137 |
| Fisher's Exact Test | 1.000 | 0.037 | 0.071 | 1.000 | 0.037 | 0.071 |
| MEGSA | 1.000 | 0.073 | 0.136 | 1.000 | 0.073 | 0.136 |
| MEMO | 0.667 | 0.078 | 0.140 | 0.667 | 0.078 | 0.140 |
| WExT | 0.571 | 0.075 | 0.133 | 0.571 | 0.075 | 0.133 |

(b) Metrics for BRCA data. (1026 samples | 34 CGC-CGC pairs)

| Method | Precision | Sensitivity | F1 Score | $\text{Precision}_{\text{strict}}$ | $\text{Sensitivity}_{\text{strict}}$ | $\text{F1 Score}_{\text{strict}}$ |
|---|---|---|---|---|---|---|
| DISCOVER | 0.652 | 0.469 | 0.545 | 0.632 | 0.375 | 0.471 |
| DISCOVER Strat | 0.711 | 0.478 | 0.571 | 0.769 | 0.448 | 0.566 |
| Fisher's Exact Test | 1.000 | 0.061 | 0.114 | 1.000 | 0.061 | 0.115 |
| MEGSA | 1.000 | 0.059 | 0.111 | 1.000 | 0.059 | 0.111 |
| MEMO | 0.644 | 0.475 | 0.547 | 0.622 | 0.377 | 0.469 |
| WExT | 0.625 | 0.536 | 0.577 | 0.684 | 0.464 | 0.553 |

(c) Metrics for COADREAD data. (498 samples | 196 CGC-CGC pairs)

| Method | Precision | Sensitivity | F1 Score | $\text{Precision}_{\text{strict}}$ | $\text{Sensitivity}_{\text{strict}}$ | $\text{F1 Score}_{\text{strict}}$ |
|---|---|---|---|---|---|---|
| DISCOVER | 0.662 | 0.224 | 0.335 | 0.706 | 0.188 | 0.297 |
| DISCOVER Strat | 0.727 | 0.042 | 0.080 | 0.727 | 0.042 | 0.079 |
| Fisher's Exact Test | 0.538 | 0.036 | 0.067 | 0.538 | 0.036 | 0.067 |
| MEGSA | 0.611 | 0.056 | 0.103 | 0.588 | 0.051 | 0.094 |
| MEMO | 0.660 | 0.339 | 0.448 | 0.657 | 0.243 | 0.355 |
| WExT | 0.691 | 0.412 | 0.516 | 0.744 | 0.342 | 0.469 |

(d) Metrics for LUAD data. (568 samples | 92 CGC-CGC pairs)

| Method | Precision | Sensitivity | F1 Score | $\text{Precision}_{\text{strict}}$ | $\text{Sensitivity}_{\text{strict}}$ | $\text{F1 Score}_{\text{strict}}$ |
|---|---|---|---|---|---|---|
| DISCOVER | 0.714 | 0.114 | 0.197 | 0.727 | 0.091 | 0.162 |
| Fisher's Exact Test | 0.000 | 0.000 | NaN | 0.000 | 0.000 | NaN |
| MEGSA | 0.667 | 0.022 | 0.043 | 0.667 | 0.022 | 0.043 |
| MEMO | 0.722 | 0.149 | 0.248 | 0.733 | 0.126 | 0.215 |
| WExT | 0.640 | 0.179 | 0.279 | 0.667 | 0.156 | 0.253 |

(e) Metrics for LUSC data. (485 samples | 38 CGC-CGC pairs)

| Method | Precision | Sensitivity | F1 Score | $\text{Precision}_{\text{strict}}$ | $\text{Sensitivity}_{\text{strict}}$ | $\text{F1 Score}_{\text{strict}}$ |
|---|---|---|---|---|---|---|
| DISCOVER | 1.0 | 0.054 | 0.103 | 1.0 | 0.054 | 0.102 |
| Fisher's Exact Test | 1.0 | 0.053 | 0.100 | 1.0 | 0.053 | 0.101 |
| MEGSA | 1.0 | 0.158 | 0.273 | 1.0 | 0.158 | 0.273 |
| MEMO | 1.0 | 0.111 | 0.200 | 1.0 | 0.111 | 0.200 |
| WExT | 1.0 | 0.111 | 0.200 | 1.0 | 0.111 | 0.200 |

(f) Metrics for SKCM data. (468 samples | 458 CGC-CGC pairs)

| Method | Precision | Sensitivity | F1 Score | $\text{Precision}_{\text{strict}}$ | $\text{Sensitivity}_{\text{strict}}$ | $\text{F1 Score}_{\text{strict}}$ |
|---|---|---|---|---|---|---|
| DISCOVER | 0.833 | 0.044 | 0.084 | 0.833 | 0.044 | 0.084 |
| Fisher's Exact Test | 1.000 | 0.004 | 0.009 | 1.000 | 0.004 | 0.008 |
| MEGSA | 0.889 | 0.018 | 0.034 | 0.889 | 0.018 | 0.035 |
| WExT | 0.697 | 0.121 | 0.206 | 0.708 | 0.116 | 0.199 |

(g) Metrics for STAD data. (438 samples | 140 CGC-CGC pairs)

| Method | Precision | Sensitivity | F1 Score | $\text{Precision}_{\text{strict}}$ | $\text{Sensitivity}_{\text{strict}}$ | $\text{F1 Score}_{\text{strict}}$ |
|---|---|---|---|---|---|---|
| DISCOVER | 0.667 | 0.119 | 0.201 | 0.684 | 0.096 | 0.168 |
| Fisher's Exact Test | 0.000 | 0.000 | NaN | 0.000 | 0.000 | NaN |
| MEGSA | 0.667 | 0.014 | 0.028 | 0.667 | 0.014 | 0.027 |
| WExT | 0.634 | 0.191 | 0.294 | 0.636 | 0.154 | 0.248 |

(h) Metrics for UCEC data. (531 samples | 1356 CGC-CGC pairs)

| Method | Precision | Sensitivity | F1 Score | $\text{Precision}_{\text{strict}}$ | $\text{Sensitivity}_{\text{strict}}$ | $\text{F1 Score}_{\text{strict}}$ |
|---|---|---|---|---|---|---|
| DISCOVER | 0.655 | 0.177 | 0.278 | 0.716 | 0.143 | 0.238 |
| Fisher's Exact Test | 0.833 | 0.007 | 0.015 | 0.833 | 0.007 | 0.014 |
| MEGSA | 0.786 | 0.008 | 0.016 | 0.786 | 0.008 | 0.016 |
| WExT | 0.617 | 0.277 | 0.383 | 0.669 | 0.227 | 0.339 |

Table 4: **Results of network-centric ME evaluation framework with $\mathcal{G}$ = Intact (w conf. threshold 0.35), $\mathcal{S}$ = CGC, c= $X_1$, $p_t$ = 0.01, t=20, robustness_iterations = 100**

(a) Metrics for BLCA data. (411 samples | 56 CGC-CGC pairs)

| Method | Precision | Sensitivity | F1 Score | $\text{Precision}_{\text{strict}}$ | $\text{Sensitivity}_{\text{strict}}$ | $\text{F1 Score}_{\text{strict}}$ |
|---|---|---|---|---|---|---|
| DISCOVER | 1.000 | 0.036 | 0.07 | 1.000 | 0.036 | 0.069 |
| Fisher's Exact Test | NaN | 0.000 | NaN | NaN | 0.000 | NaN |
| MEGSA | NaN | 0.000 | NaN | NaN | 0.000 | NaN |
| MEMO | 0.667 | 0.037 | 0.07 | 0.667 | 0.037 | 0.070 |
| WExT | 0.667 | 0.037 | 0.07 | 0.667 | 0.037 | 0.070 |

(b) Metrics for BRCA data. (1026 samples | 34 CGC-CGC pairs)

| Method | Precision | Sensitivity | F1 Score | $\text{Precision}_{\text{strict}}$ | $\text{Sensitivity}_{\text{strict}}$ | $\text{F1 Score}_{\text{strict}}$ |
|---|---|---|---|---|---|---|
| DISCOVER | 0.714 | 0.156 | 0.256 | 0.714 | 0.156 | 0.256 |
| DISCOVER Strat | 0.857 | 0.176 | 0.293 | 0.857 | 0.176 | 0.292 |
| Fisher's Exact Test | NaN | 0.000 | NaN | NaN | 0.000 | NaN |
| MEGSA | NaN | 0.000 | NaN | NaN | 0.000 | NaN |
| MEMO | 0.625 | 0.147 | 0.238 | 0.625 | 0.147 | 0.238 |
| WExT | 0.556 | 0.152 | 0.238 | 0.556 | 0.152 | 0.239 |

(c) Metrics for COADREAD data. (498 samples | 196 CGC-CGC pairs)

| Method | Precision | Sensitivity | F1 Score | $\text{Precision}_{\text{strict}}$ | $\text{Sensitivity}_{\text{strict}}$ | $\text{F1 Score}_{\text{strict}}$ |
|---|---|---|---|---|---|---|
| DISCOVER | 0.686 | 0.125 | 0.211 | 0.690 | 0.104 | 0.181 |
| DISCOVER Strat | 0.000 | 0.000 | NaN | 0.000 | 0.000 | NaN |
| Fisher's Exact Test | 0.000 | 0.000 | NaN | 0.000 | 0.000 | NaN |
| MEGSA | 0.400 | 0.010 | 0.020 | 0.400 | 0.010 | 0.020 |
| MEMO | 0.689 | 0.265 | 0.383 | 0.655 | 0.187 | 0.291 |
| WExT | 0.679 | 0.280 | 0.396 | 0.724 | 0.222 | 0.340 |

(d) Metrics for LUAD data. (568 samples | 92 CGC-CGC pairs)

| Method | Precision | Sensitivity | F1 Score | $\text{Precision}_{\text{strict}}$ | $\text{Sensitivity}_{\text{strict}}$ | $\text{F1 Score}_{\text{strict}}$ |
|---|---|---|---|---|---|---|
| DISCOVER | 0.667 | 0.044 | 0.082 | 0.667 | 0.044 | 0.083 |
| Fisher's Exact Test | NaN | 0.000 | NaN | NaN | 0.000 | NaN |
| MEGSA | 0.667 | 0.022 | 0.042 | 0.667 | 0.022 | 0.043 |
| MEMO | 0.700 | 0.078 | 0.141 | 0.667 | 0.067 | 0.122 |
| WExT | 0.750 | 0.105 | 0.184 | 0.700 | 0.081 | 0.145 |

(e) Metrics for LUSC data. (485 samples | 38 CGC-CGC pairs)

| Method | Precision | Sensitivity | F1 Score | $\text{Precision}_{\text{strict}}$ | $\text{Sensitivity}_{\text{strict}}$ | $\text{F1 Score}_{\text{strict}}$ |
|---|---|---|---|---|---|---|
| DISCOVER | 1.0 | 0.053 | 0.100 | 1.0 | 0.053 | 0.101 |
| Fisher's Exact Test | NaN | 0.000 | NaN | NaN | 0.000 | NaN |
| MEGSA | NaN | 0.000 | NaN | NaN | 0.000 | NaN |
| MEMO | 1.0 | 0.056 | 0.105 | 1.0 | 0.056 | 0.106 |
| WExT | 1.0 | 0.053 | 0.100 | 1.0 | 0.053 | 0.101 |

(f) Metrics for SKCM data. (468 samples | 458 CGC-CGC pairs)

| Method | Precision | Sensitivity | F1 Score | $\text{Precision}_{\text{strict}}$ | $\text{Sensitivity}_{\text{strict}}$ | $\text{F1 Score}_{\text{strict}}$ |
|---|---|---|---|---|---|---|
| DISCOVER | 1.000 | 0.013 | 0.026 | 1.00 | 0.013 | 0.026 |
| Fisher's Exact Test | 1.000 | 0.004 | 0.009 | 1.00 | 0.004 | 0.008 |
| MEGSA | 1.000 | 0.004 | 0.009 | 1.00 | 0.004 | 0.008 |
| WExT | 0.731 | 0.042 | 0.080 | 0.75 | 0.040 | 0.076 |

(g) Metrics for STAD data. (438 samples | 140 CGC-CGC pairs)

| Method | Precision | Sensitivity | F1 Score | $\text{Precision}_{\text{strict}}$ | $\text{Sensitivity}_{\text{strict}}$ | $\text{F1 Score}_{\text{strict}}$ |
|---|---|---|---|---|---|---|
| DISCOVER | 0.667 | 0.059 | 0.108 | 0.70 | 0.051 | 0.095 |
| Fisher's Exact Test | NaN | 0.000 | NaN | NaN | 0.000 | NaN |
| MEGSA | NaN | 0.000 | NaN | NaN | 0.000 | NaN |
| WExT | 0.622 | 0.084 | 0.148 | 0.63 | 0.062 | 0.113 |

(h) Metrics for UCEC data. (531 samples | 1356 CGC-CGC pairs)

| Method | Precision | Sensitivity | F1 Score | $\text{Precision}_{\text{strict}}$ | $\text{Sensitivity}_{\text{strict}}$ | $\text{F1 Score}_{\text{strict}}$ |
|---|---|---|---|---|---|---|
| DISCOVER | 0.654 | 0.120 | 0.203 | 0.718 | 0.100 | 0.176 |
| Fisher's Exact Test | 0.857 | 0.004 | 0.009 | 0.857 | 0.004 | 0.008 |
| MEGSA | 0.857 | 0.004 | 0.009 | 0.857 | 0.004 | 0.008 |
| WExT | 0.636 | 0.199 | 0.303 | 0.699 | 0.158 | 0.258 |

Table 5: **Results of network-centric ME evaluation framework with $\mathcal{G}$ = Intact (w conf. threshold 0.35), $\mathcal{S}$ = CGC, c= $X_1$, $p_t$ = 0.1, t=20, robustness_iterations = 100**

(a) Metrics for BLCA data. (411 samples | 56 CGC-CGC pairs)

| Method | Precision | Sensitivity | F1 Score | $\text{Precision}_{\text{strict}}$ | $\text{Sensitivity}_{\text{strict}}$ | $\text{F1 Score}_{\text{strict}}$ |
|---|---|---|---|---|---|---|
| DISCOVER | 0.75 | 0.111 | 0.194 | 0.750 | 0.111 | 0.193 |
| Fisher's Exact Test | 1.00 | 0.036 | 0.070 | 1.000 | 0.036 | 0.069 |
| MEGSA | 0.80 | 0.074 | 0.136 | 0.800 | 0.074 | 0.135 |
| MEMO | 0.60 | 0.113 | 0.190 | 0.600 | 0.113 | 0.190 |
| WExT | 0.60 | 0.117 | 0.195 | 0.667 | 0.117 | 0.199 |

(b) Metrics for BRCA data. (1026 samples | 34 CGC-CGC pairs)

| Method | Precision | Sensitivity | F1 Score | $\text{Precision}_{\text{strict}}$ | $\text{Sensitivity}_{\text{strict}}$ | $\text{F1 Score}_{\text{strict}}$ |
|---|---|---|---|---|---|---|
| DISCOVER | 0.660 | 0.600 | 0.629 | 0.692 | 0.491 | 0.574 |
| DISCOVER Strat | 0.667 | 0.600 | 0.632 | 0.696 | 0.533 | 0.604 |
| Fisher's Exact Test | 1.000 | 0.062 | 0.118 | 1.000 | 0.062 | 0.117 |
| MEGSA | 0.667 | 0.059 | 0.108 | 0.667 | 0.059 | 0.108 |
| MEMO | 0.635 | 0.600 | 0.617 | 0.667 | 0.473 | 0.553 |
| WExT | 0.638 | 0.698 | 0.667 | 0.707 | 0.547 | 0.617 |

(c) Metrics for COADREAD data. (498 samples | 196 CGC-CGC pairs)

| Method | Precision | Sensitivity | F1 Score | $\text{Precision}_{\text{strict}}$ | $\text{Sensitivity}_{\text{strict}}$ | $\text{F1 Score}_{\text{strict}}$ |
|---|---|---|---|---|---|---|
| DISCOVER | 0.649 | 0.291 | 0.401 | 0.702 | 0.245 | 0.363 |
| DISCOVER Strat | 0.738 | 0.127 | 0.217 | 0.737 | 0.111 | 0.193 |
| Fisher's Exact Test | 0.708 | 0.087 | 0.156 | 0.667 | 0.072 | 0.130 |
| MEGSA | 0.714 | 0.103 | 0.179 | 0.688 | 0.085 | 0.151 |
| MEMO | 0.670 | 0.423 | 0.519 | 0.678 | 0.324 | 0.438 |
| WExT | 0.640 | 0.421 | 0.508 | 0.703 | 0.347 | 0.465 |

(d) Metrics for LUAD data. (568 samples | 92 CGC-CGC pairs)

| Method | Precision | Sensitivity | F1 Score | $\text{Precision}_{\text{strict}}$ | $\text{Sensitivity}_{\text{strict}}$ | $\text{F1 Score}_{\text{strict}}$ |
|---|---|---|---|---|---|---|
| DISCOVER | 0.698 | 0.173 | 0.278 | 0.703 | 0.150 | 0.247 |
| Fisher's Exact Test | 0.667 | 0.022 | 0.043 | 0.667 | 0.022 | 0.043 |
| MEGSA | 0.667 | 0.066 | 0.120 | 0.667 | 0.066 | 0.120 |
| MEMO | 0.645 | 0.220 | 0.328 | 0.654 | 0.187 | 0.291 |
| WExT | 0.659 | 0.326 | 0.436 | 0.667 | 0.270 | 0.384 |

(e) Metrics for LUSC data. (485 samples | 38 CGC-CGC pairs)

| Method | Precision | Sensitivity | F1 Score | $\text{Precision}_{\text{strict}}$ | $\text{Sensitivity}_{\text{strict}}$ | $\text{F1 Score}_{\text{strict}}$ |
|---|---|---|---|---|---|---|
| DISCOVER | 1.000 | 0.111 | 0.200 | 1.000 | 0.111 | 0.200 |
| Fisher's Exact Test | 1.000 | 0.056 | 0.105 | 1.000 | 0.056 | 0.106 |
| MEGSA | 0.857 | 0.154 | 0.261 | 0.857 | 0.154 | 0.261 |
| MEMO | 0.800 | 0.118 | 0.205 | 0.800 | 0.118 | 0.206 |
| WExT | 0.750 | 0.176 | 0.286 | 0.750 | 0.176 | 0.285 |

(f) Metrics for SKCM data. (468 samples | 458 CGC-CGC pairs)

| Method | Precision | Sensitivity | F1 Score | $\text{Precision}_{\text{strict}}$ | $\text{Sensitivity}_{\text{strict}}$ | $\text{F1 Score}_{\text{strict}}$ |
|---|---|---|---|---|---|---|
| DISCOVER | 0.740 | 0.105 | 0.184 | 0.756 | 0.101 | 0.178 |
| Fisher's Exact Test | 1.000 | 0.004 | 0.009 | 1.000 | 0.004 | 0.008 |
| MEGSA | 0.800 | 0.026 | 0.051 | 0.800 | 0.026 | 0.050 |
| WExT | 0.622 | 0.170 | 0.267 | 0.634 | 0.163 | 0.259 |

(g) Metrics for STAD data. (438 samples | 140 CGC-CGC pairs)

| Method | Precision | Sensitivity | F1 Score | $\text{Precision}_{\text{strict}}$ | $\text{Sensitivity}_{\text{strict}}$ | $\text{F1 Score}_{\text{strict}}$ |
|---|---|---|---|---|---|---|
| DISCOVER | 0.641 | 0.183 | 0.285 | 0.625 | 0.147 | 0.238 |
| Fisher's Exact Test | 0.667 | 0.014 | 0.028 | 0.667 | 0.014 | 0.027 |
| MEGSA | 0.500 | 0.014 | 0.028 | 0.500 | 0.014 | 0.027 |
| WExT | 0.632 | 0.224 | 0.331 | 0.623 | 0.179 | 0.278 |

(h) Metrics for UCEC data. (531 samples | 1356 CGC-CGC pairs)

| Method | Precision | Sensitivity | F1 Score | $\text{Precision}_{\text{strict}}$ | $\text{Sensitivity}_{\text{strict}}$ | $\text{F1 Score}_{\text{strict}}$ |
|---|---|---|---|---|---|---|
| DISCOVER | 0.621 | 0.219 | 0.324 | 0.672 | 0.176 | 0.279 |
| Fisher's Exact Test | 0.750 | 0.013 | 0.026 | 0.773 | 0.013 | 0.026 |
| MEGSA | 0.806 | 0.018 | 0.036 | 0.821 | 0.017 | 0.033 |
| WExT | 0.594 | 0.344 | 0.436 | 0.640 | 0.279 | 0.389 |

Table 6: **Results of network-centric ME evaluation framework with $\mathcal{G}$ = Intact (w conf. threshold 0.35), $\mathcal{S} = CGC_{SNV}$ , c= $X_1$, $p_t$ = 0.05, t=20, robustness_iterations = 100**

(a) Metrics for BLCA data. (411 samples | 36 $CGC_{SNV}$-$CGC_{SNV}$ pairs)

| Method | Precision | Sensitivity | F1 Score | $\text{Precision}_{\text{strict}}$ | $\text{Sensitivity}_{\text{strict}}$ | $\text{F1 Score}_{\text{strict}}$ |
|---|---|---|---|---|---|---|
| DISCOVER | 0.8 | 0.118 | 0.205 | 0.8 | 0.118 | 0.206 |
| Fisher's Exact Test | 1.0 | 0.056 | 0.105 | 1.0 | 0.056 | 0.106 |
| MEGSA | 1.0 | 0.113 | 0.203 | 1.0 | 0.113 | 0.203 |
| MEMO | 0.8 | 0.121 | 0.211 | 0.8 | 0.121 | 0.210 |
| WExT | 0.8 | 0.129 | 0.222 | 0.8 | 0.129 | 0.222 |

(b) Metrics for BRCA data. (1026 samples | 28 $CGC_{SNV}$-$CGC_{SNV}$ pairs)

| Method | Precision | Sensitivity | F1 Score | $\text{Precision}_{\text{strict}}$ | $\text{Sensitivity}_{\text{strict}}$ | $\text{F1 Score}_{\text{strict}}$ |
|---|---|---|---|---|---|---|
| DISCOVER | 0.636 | 0.519 | 0.571 | 0.647 | 0.407 | 0.500 |
| DISCOVER Strat | 0.737 | 0.560 | 0.636 | 0.812 | 0.520 | 0.634 |
| Fisher's Exact Test | 1.000 | 0.073 | 0.136 | 1.000 | 0.073 | 0.136 |
| MEGSA | 1.000 | 0.071 | 0.133 | 1.000 | 0.071 | 0.133 |
| MEMO | 0.667 | 0.480 | 0.558 | 0.667 | 0.400 | 0.500 |
| WExT | 0.625 | 0.556 | 0.588 | 0.645 | 0.444 | 0.526 |

(c) Metrics for COADREAD data. (498 samples | 146 $CGC_{SNV}$-$CGC_{SNV}$ pairs)

| Method | Precision | Sensitivity | F1 Score | $\text{Precision}_{\text{strict}}$ | $\text{Sensitivity}_{\text{strict}}$ | $\text{F1 Score}_{\text{strict}}$ |
|---|---|---|---|---|---|---|
| DISCOVER | 0.635 | 0.237 | 0.345 | 0.675 | 0.194 | 0.301 |
| DISCOVER Strat | 0.750 | 0.042 | 0.079 | 0.750 | 0.042 | 0.080 |
| Fisher's Exact Test | 0.600 | 0.041 | 0.077 | 0.600 | 0.041 | 0.077 |
| MEGSA | 0.667 | 0.068 | 0.123 | 0.643 | 0.061 | 0.111 |
| MEMO | 0.634 | 0.330 | 0.434 | 0.627 | 0.234 | 0.341 |
| WExT | 0.656 | 0.380 | 0.482 | 0.706 | 0.304 | 0.425 |

(d) Metrics for LUAD data. (568 samples | 70 $CGC_{SNV}$-$CGC_{SNV}$ pairs)

| Method | Precision | Sensitivity | F1 Score | $\text{Precision}_{\text{strict}}$ | $\text{Sensitivity}_{\text{strict}}$ | $\text{F1 Score}_{\text{strict}}$ |
|---|---|---|---|---|---|---|
| DISCOVER | 0.727 | 0.121 | 0.208 | 0.667 | 0.091 | 0.160 |
| Fisher's Exact Test | 0.000 | 0.000 | NaN | 0.000 | 0.000 | NaN |
| MEGSA | 0.500 | 0.029 | 0.055 | 0.500 | 0.029 | 0.055 |
| MEMO | 0.686 | 0.185 | 0.291 | 0.690 | 0.154 | 0.252 |
| WExT | 0.609 | 0.211 | 0.313 | 0.600 | 0.180 | 0.277 |

(e) Metrics for LUSC data. (485 samples | 30 $CGC_{SNV}$-$CGC_{SNV}$ pairs)

| Method | Precision | Sensitivity | F1 Score | $\text{Precision}_{\text{strict}}$ | $\text{Sensitivity}_{\text{strict}}$ | $\text{F1 Score}_{\text{strict}}$ |
|---|---|---|---|---|---|---|
| DISCOVER | 1.0 | 0.067 | 0.125 | 1.0 | 0.067 | 0.126 |
| Fisher's Exact Test | 1.0 | 0.067 | 0.125 | 1.0 | 0.067 | 0.126 |
| MEGSA | 1.0 | 0.200 | 0.333 | 1.0 | 0.200 | 0.333 |
| MEMO | 1.0 | 0.138 | 0.242 | 1.0 | 0.138 | 0.243 |
| WExT | 1.0 | 0.140 | 0.246 | 1.0 | 0.140 | 0.246 |

(f) Metrics for SKCM data. (468 samples | 288 $CGC_{SNV}$-$CGC_{SNV}$ pairs)

| Method | Precision | Sensitivity | F1 Score | $\text{Precision}_{\text{strict}}$ | $\text{Sensitivity}_{\text{strict}}$ | $\text{F1 Score}_{\text{strict}}$ |
|---|---|---|---|---|---|---|
| DISCOVER | 0.714 | 0.036 | 0.068 | 0.714 | 0.036 | 0.069 |
| Fisher's Exact Test | 1.000 | 0.007 | 0.014 | 1.000 | 0.007 | 0.014 |
| MEGSA | 0.800 | 0.014 | 0.027 | 0.800 | 0.014 | 0.028 |
| WExT | 0.651 | 0.131 | 0.218 | 0.660 | 0.122 | 0.206 |

(g) Metrics for STAD data. (438 samples | 112 $CGC_{SNV}$-$CGC_{SNV}$ pairs)

| Method | Precision | Sensitivity | F1 Score | $\text{Precision}_{\text{strict}}$ | $\text{Sensitivity}_{\text{strict}}$ | $\text{F1 Score}_{\text{strict}}$ |
|---|---|---|---|---|---|---|
| DISCOVER | 0.600 | 0.111 | 0.187 | 0.562 | 0.083 | 0.145 |
| Fisher's Exact Test | 0.000 | 0.000 | NaN | 0.000 | 0.000 | NaN |
| MEGSA | 0.667 | 0.018 | 0.034 | 0.667 | 0.018 | 0.035 |
| WExT | 0.609 | 0.182 | 0.281 | 0.588 | 0.140 | 0.226 |

(h) Metrics for UCEC data. (531 samples | 776 $CGC_{SNV}$-$CGC_{SNV}$ pairs)

| Method | Precision | Sensitivity | F1 Score | $\text{Precision}_{\text{strict}}$ | $\text{Sensitivity}_{\text{strict}}$ | $\text{F1 Score}_{\text{strict}}$ |
|---|---|---|---|---|---|---|
| DISCOVER | 0.637 | 0.210 | 0.316 | 0.681 | 0.166 | 0.267 |
| Fisher's Exact Test | 0.833 | 0.013 | 0.025 | 0.833 | 0.013 | 0.026 |
| MEGSA | 0.846 | 0.014 | 0.028 | 0.846 | 0.014 | 0.028 |
| WExT | 0.599 | 0.318 | 0.415 | 0.630 | 0.250 | 0.358 |

Table 7: **Results of network-centric ME evaluation framework with $\mathcal{G}$ = Intact (w conf. threshold 0.35), $\mathcal{S}$ = Intogen , c= $X_1$, $p_t$ = 0.05, t=20, robustness_iterations = 100**

(a) Metrics for BLCA data. (411 samples | 56 Intogen-Intogen pairs)

| Method | Precision | Sensitivity | F1 Score | $\text{Precision}_{\text{strict}}$ | $\text{Sensitivity}_{\text{strict}}$ | $\text{F1 Score}_{\text{strict}}$ |
|---|---|---|---|---|---|---|
| DISCOVER | 0.889 | 0.145 | 0.250 | 0.889 | 0.145 | 0.249 |
| Fisher's Exact Test | 1.000 | 0.036 | 0.070 | 1.000 | 0.036 | 0.069 |
| MEGSA | 1.000 | 0.109 | 0.197 | 1.000 | 0.109 | 0.197 |
| MEMO | 0.800 | 0.150 | 0.252 | 0.800 | 0.150 | 0.253 |
| WExT | 0.727 | 0.154 | 0.254 | 0.727 | 0.154 | 0.254 |

(b) Metrics for BRCA data. (1026 samples | 36 Intogen-Intogen pairs)

| Method | Precision | Sensitivity | F1 Score | $\text{Precision}_{\text{strict}}$ | $\text{Sensitivity}_{\text{strict}}$ | $\text{F1 Score}_{\text{strict}}$ |
|---|---|---|---|---|---|---|
| DISCOVER | 0.654 | 0.500 | 0.567 | 0.619 | 0.382 | 0.472 |
| DISCOVER Strat | 0.700 | 0.444 | 0.544 | 0.722 | 0.413 | 0.525 |
| Fisher's Exact Test | 1.000 | 0.057 | 0.108 | 1.000 | 0.057 | 0.108 |
| MEGSA | 1.000 | 0.056 | 0.105 | 1.000 | 0.056 | 0.106 |
| MEMO | 0.667 | 0.500 | 0.571 | 0.650 | 0.406 | 0.500 |
| WExT | 0.654 | 0.567 | 0.607 | 0.667 | 0.467 | 0.549 |

(c) Metrics for COADREAD data. (498 samples | 206 Intogen-Intogen pairs)

| Method | Precision | Sensitivity | F1 Score | $\text{Precision}_{\text{strict}}$ | $\text{Sensitivity}_{\text{strict}}$ | $\text{F1 Score}_{\text{strict}}$ |
|---|---|---|---|---|---|---|
| DISCOVER | 0.657 | 0.224 | 0.335 | 0.685 | 0.189 | 0.296 |
| DISCOVER Strat | 0.679 | 0.048 | 0.090 | 0.679 | 0.048 | 0.090 |
| Fisher's Exact Test | 0.538 | 0.034 | 0.065 | 0.538 | 0.034 | 0.064 |
| MEGSA | 0.611 | 0.054 | 0.099 | 0.588 | 0.049 | 0.090 |
| MEMO | 0.652 | 0.339 | 0.446 | 0.638 | 0.245 | 0.354 |
| WExT | 0.678 | 0.416 | 0.515 | 0.722 | 0.338 | 0.460 |

(d) Metrics for LUAD data. (568 samples | 96 Intogen-Intogen pairs)

| Method | Precision | Sensitivity | F1 Score | $\text{Precision}_{\text{strict}}$ | $\text{Sensitivity}_{\text{strict}}$ | $\text{F1 Score}_{\text{strict}}$ |
|---|---|---|---|---|---|---|
| DISCOVER | 0.800 | 0.091 | 0.163 | 0.778 | 0.080 | 0.145 |
| Fisher's Exact Test | 0.000 | 0.000 | NaN | 0.000 | 0.000 | NaN |
| MEGSA | 0.667 | 0.021 | 0.041 | 0.667 | 0.021 | 0.041 |
| MEMO | 0.765 | 0.152 | 0.254 | 0.786 | 0.129 | 0.222 |
| WExT | 0.667 | 0.184 | 0.288 | 0.700 | 0.161 | 0.262 |

(e) Metrics for LUSC data. (485 samples | 50 Intogen-Intogen pairs)

| Method | Precision | Sensitivity | F1 Score | $\text{Precision}_{\text{strict}}$ | $\text{Sensitivity}_{\text{strict}}$ | $\text{F1 Score}_{\text{strict}}$ |
|---|---|---|---|---|---|---|
| DISCOVER | 1.000 | 0.081 | 0.150 | 1.000 | 0.081 | 0.150 |
| Fisher's Exact Test | 1.000 | 0.040 | 0.077 | 1.000 | 0.040 | 0.077 |
| MEGSA | 1.000 | 0.167 | 0.286 | 1.000 | 0.167 | 0.286 |
| MEMO | 1.000 | 0.128 | 0.226 | 1.000 | 0.128 | 0.227 |
| WExT | 0.857 | 0.129 | 0.224 | 0.857 | 0.129 | 0.224 |

(f) Metrics for SKCM data. (468 samples | 464 Intogen-Intogen pairs)

| Method | Precision | Sensitivity | F1 Score | $\text{Precision}_{\text{strict}}$ | $\text{Sensitivity}_{\text{strict}}$ | $\text{F1 Score}_{\text{strict}}$ |
|---|---|---|---|---|---|---|
| DISCOVER | 0.750 | 0.040 | 0.075 | 0.783 | 0.040 | 0.076 |
| Fisher's Exact Test | 1.000 | 0.004 | 0.009 | 1.000 | 0.004 | 0.008 |
| MEGSA | 0.857 | 0.013 | 0.026 | 0.857 | 0.013 | 0.026 |
| WExT | 0.671 | 0.115 | 0.197 | 0.681 | 0.111 | 0.191 |

(g) Metrics for STAD data. (438 samples | 162 Intogen-Intogen pairs)

| Method | Precision | Sensitivity | F1 Score | $\text{Precision}_{\text{strict}}$ | $\text{Sensitivity}_{\text{strict}}$ | $\text{F1 Score}_{\text{strict}}$ |
|---|---|---|---|---|---|---|
| DISCOVER | 0.625 | 0.095 | 0.164 | 0.600 | 0.076 | 0.135 |
| Fisher's Exact Test | 0.000 | 0.000 | NaN | 0.000 | 0.000 | NaN |
| MEGSA | 0.667 | 0.012 | 0.024 | 0.667 | 0.012 | 0.024 |
| WExT | 0.673 | 0.214 | 0.325 | 0.650 | 0.169 | 0.268 |

(h) Metrics for UCEC data. (531 samples | 1240 Intogen-Intogen pairs)

| Method | Precision | Sensitivity | F1 Score | $\text{Precision}_{\text{strict}}$ | $\text{Sensitivity}_{\text{strict}}$ | $\text{F1 Score}_{\text{strict}}$ |
|---|---|---|---|---|---|---|
| DISCOVER | 0.636 | 0.210 | 0.315 | 0.687 | 0.167 | 0.269 |
| Fisher's Exact Test | 0.778 | 0.011 | 0.022 | 0.765 | 0.011 | 0.022 |
| MEGSA | 0.727 | 0.013 | 0.025 | 0.750 | 0.012 | 0.024 |
| WExT | 0.586 | 0.309 | 0.405 | 0.621 | 0.245 | 0.351 |

Table 8: **Results of network-centric ME evaluation framework with $\mathcal{G}$ = Intact (w conf. threshold 0.25), $\mathcal{S}$ = CGC, c= $X_1$, $p_t$ = 0.05, t=20, robustness_iterations = 100**

(a) Metrics for BLCA data. (411 samples | 92 CGC-CGC pairs)

| Method | Precision | Sensitivity | F1 Score | $\text{Precision}_{\text{strict}}$ | $\text{Sensitivity}_{\text{strict}}$ | $\text{F1 Score}_{\text{strict}}$ |
|---|---|---|---|---|---|---|
| DISCOVER | 0.833 | 0.114 | 0.200 | 0.833 | 0.114 | 0.201 |
| Fisher's Exact Test | 1.000 | 0.066 | 0.124 | 1.000 | 0.066 | 0.124 |
| MEGSA | 1.000 | 0.109 | 0.197 | 1.000 | 0.109 | 0.197 |
| MEMO | 0.750 | 0.103 | 0.182 | 0.750 | 0.103 | 0.181 |
| WExT | 0.737 | 0.163 | 0.267 | 0.778 | 0.163 | 0.270 |

(b) Metrics for BRCA data. (1026 samples | 54 CGC-CGC pairs)

| Method | Precision | Sensitivity | F1 Score | $\text{Precision}_{\text{strict}}$ | $\text{Sensitivity}_{\text{strict}}$ | $\text{F1 Score}_{\text{strict}}$ |
|---|---|---|---|---|---|---|
| DISCOVER | 0.712 | 0.485 | 0.577 | 0.702 | 0.412 | 0.519 |
| DISCOVER Strat | 0.782 | 0.473 | 0.589 | 0.796 | 0.429 | 0.558 |
| Fisher's Exact Test | 0.667 | 0.038 | 0.071 | 0.667 | 0.038 | 0.072 |
| MEGSA | 0.667 | 0.037 | 0.070 | 0.667 | 0.037 | 0.070 |
| MEMO | 0.706 | 0.485 | 0.575 | 0.700 | 0.424 | 0.528 |
| WExT | 0.630 | 0.580 | 0.604 | 0.639 | 0.460 | 0.535 |

(c) Metrics for COADREAD data. (498 samples | 264 CGC-CGC pairs)

| Method | Precision | Sensitivity | F1 Score | $\text{Precision}_{\text{strict}}$ | $\text{Sensitivity}_{\text{strict}}$ | $\text{F1 Score}_{\text{strict}}$ |
|---|---|---|---|---|---|---|
| DISCOVER | 0.659 | 0.216 | 0.325 | 0.713 | 0.181 | 0.289 |
| DISCOVER Strat | 0.667 | 0.054 | 0.100 | 0.684 | 0.050 | 0.093 |
| Fisher's Exact Test | 0.583 | 0.053 | 0.097 | 0.591 | 0.049 | 0.090 |
| MEGSA | 0.667 | 0.072 | 0.131 | 0.667 | 0.065 | 0.118 |
| MEMO | 0.648 | 0.318 | 0.427 | 0.615 | 0.220 | 0.324 |
| WExT | 0.681 | 0.372 | 0.481 | 0.731 | 0.301 | 0.426 |

(d) Metrics for LUAD data. (568 samples | 138 CGC-CGC pairs)

| Method | Precision | Sensitivity | F1 Score | $\text{Precision}_{\text{strict}}$ | $\text{Sensitivity}_{\text{strict}}$ | $\text{F1 Score}_{\text{strict}}$ |
|---|---|---|---|---|---|---|
| DISCOVER | 0.755 | 0.153 | 0.254 | 0.800 | 0.137 | 0.234 |
| Fisher's Exact Test | 0.800 | 0.029 | 0.056 | 0.800 | 0.029 | 0.056 |
| MEGSA | 0.727 | 0.058 | 0.107 | 0.727 | 0.058 | 0.107 |
| MEMO | 0.735 | 0.191 | 0.303 | 0.767 | 0.176 | 0.286 |
| WExT | 0.686 | 0.225 | 0.339 | 0.753 | 0.210 | 0.328 |

(e) Metrics for LUSC data. (485 samples | 70 CGC-CGC pairs)

| Method | Precision | Sensitivity | F1 Score | $\text{Precision}_{\text{strict}}$ | $\text{Sensitivity}_{\text{strict}}$ | $\text{F1 Score}_{\text{strict}}$ |
|---|---|---|---|---|---|---|
| DISCOVER | 1.000 | 0.029 | 0.056 | 1.000 | 0.029 | 0.056 |
| Fisher's Exact Test | 1.000 | 0.029 | 0.056 | 1.000 | 0.029 | 0.056 |
| MEGSA | 0.857 | 0.087 | 0.158 | 0.857 | 0.087 | 0.158 |
| MEMO | 0.800 | 0.059 | 0.110 | 0.800 | 0.059 | 0.110 |
| WExT | 0.571 | 0.058 | 0.106 | 0.571 | 0.058 | 0.105 |

(f) Metrics for SKCM data. (468 samples | 640 CGC-CGC pairs)

| Method | Precision | Sensitivity | F1 Score | $\text{Precision}_{\text{strict}}$ | $\text{Sensitivity}_{\text{strict}}$ | $\text{F1 Score}_{\text{strict}}$ |
|---|---|---|---|---|---|---|
| DISCOVER | 0.845 | 0.048 | 0.091 | 0.870 | 0.048 | 0.091 |
| Fisher's Exact Test | 1.000 | 0.003 | 0.006 | 1.000 | 0.003 | 0.006 |
| MEGSA | 0.889 | 0.013 | 0.025 | 0.889 | 0.013 | 0.026 |
| WExT | 0.714 | 0.110 | 0.190 | 0.721 | 0.105 | 0.183 |

(g) Metrics for STAD data. (438 samples | 194 CGC-CGC pairs)

| Method | Precision | Sensitivity | F1 Score | $\text{Precision}_{\text{strict}}$ | $\text{Sensitivity}_{\text{strict}}$ | $\text{F1 Score}_{\text{strict}}$ |
|---|---|---|---|---|---|---|
| DISCOVER | 0.675 | 0.142 | 0.235 | 0.710 | 0.116 | 0.199 |
| Fisher's Exact Test | 0.667 | 0.010 | 0.020 | 0.667 | 0.010 | 0.020 |
| MEGSA | 0.667 | 0.021 | 0.040 | 0.667 | 0.021 | 0.041 |
| WExT | 0.661 | 0.201 | 0.308 | 0.698 | 0.163 | 0.264 |

(h) Metrics for UCEC data. (531 samples | 1916 CGC-CGC pairs)

| Method | Precision | Sensitivity | F1 Score | $\text{Precision}_{\text{strict}}$ | $\text{Sensitivity}_{\text{strict}}$ | $\text{F1 Score}_{\text{strict}}$ |
|---|---|---|---|---|---|---|
| DISCOVER | 0.652 | 0.163 | 0.261 | 0.708 | 0.132 | 0.223 |
| Fisher's Exact Test | 0.771 | 0.007 | 0.014 | 0.771 | 0.007 | 0.014 |
| MEGSA | 0.773 | 0.009 | 0.018 | 0.773 | 0.009 | 0.018 |
| WExT | 0.610 | 0.267 | 0.371 | 0.656 | 0.220 | 0.329 |

Table 9: **Results of network-centric ME evaluation framework with $\mathcal{G}$ = Intact (w conf. threshold 0.45), $\mathcal{S}$ = CGC, c= $X_1$, $p_t$ = 0.05, t=20, robustness_iterations = 100**

(a) Metrics for BLCA data. (411 samples | 34 CGC-CGC pairs)

| Method | Precision | Sensitivity | F1 Score | $\text{Precision}_{\text{strict}}$ | $\text{Sensitivity}_{\text{strict}}$ | $\text{F1 Score}_{\text{strict}}$ |
|---|---|---|---|---|---|---|
| DISCOVER | 1.0 | 0.127 | 0.225 | 1.0 | 0.127 | 0.225 |
| Fisher's Exact Test | 1.0 | 0.060 | 0.113 | 1.0 | 0.060 | 0.113 |
| MEGSA | 1.0 | 0.125 | 0.222 | 1.0 | 0.125 | 0.222 |
| MEMO | 0.8 | 0.129 | 0.222 | 0.8 | 0.129 | 0.222 |
| WExT | 0.8 | 0.127 | 0.219 | 0.8 | 0.127 | 0.219 |

(b) Metrics for BRCA data. (1026 samples | 18 CGC-CGC pairs)

| Method | Precision | Sensitivity | F1 Score | $\text{Precision}_{\text{strict}}$ | $\text{Sensitivity}_{\text{strict}}$ | $\text{F1 Score}_{\text{strict}}$ |
|---|---|---|---|---|---|---|
| DISCOVER | 0.667 | 0.457 | 0.542 | 0.700 | 0.400 | 0.509 |
| DISCOVER Strat | 0.778 | 0.438 | 0.560 | 0.778 | 0.438 | 0.560 |
| Fisher's Exact Test | 1.000 | 0.125 | 0.222 | 1.000 | 0.125 | 0.222 |
| MEGSA | 1.000 | 0.114 | 0.205 | 1.000 | 0.114 | 0.205 |
| MEMO | 0.636 | 0.467 | 0.538 | 0.667 | 0.400 | 0.500 |
| WExT | 0.583 | 0.467 | 0.519 | 0.667 | 0.400 | 0.500 |

(c) Metrics for COADREAD data. (498 samples | 100 CGC-CGC pairs)

| Method | Precision | Sensitivity | F1 Score | $\text{Precision}_{\text{strict}}$ | $\text{Sensitivity}_{\text{strict}}$ | $\text{F1 Score}_{\text{strict}}$ |
|---|---|---|---|---|---|---|
| DISCOVER | 0.731 | 0.261 | 0.384 | 0.837 | 0.218 | 0.346 |
| DISCOVER Strat | 0.750 | 0.062 | 0.114 | 0.750 | 0.062 | 0.115 |
| Fisher's Exact Test | 0.600 | 0.030 | 0.058 | 0.600 | 0.030 | 0.057 |
| MEGSA | 0.727 | 0.080 | 0.144 | 0.700 | 0.070 | 0.127 |
| MEMO | 0.700 | 0.311 | 0.431 | 0.724 | 0.233 | 0.353 |
| WExT | 0.660 | 0.368 | 0.473 | 0.732 | 0.316 | 0.441 |

(d) Metrics for LUAD data. (568 samples | 46 CGC-CGC pairs)

| Method | Precision | Sensitivity | F1 Score | $\text{Precision}_{\text{strict}}$ | $\text{Sensitivity}_{\text{strict}}$ | $\text{F1 Score}_{\text{strict}}$ |
|---|---|---|---|---|---|---|
| DISCOVER | 0.800 | 0.089 | 0.160 | 0.750 | 0.067 | 0.123 |
| Fisher's Exact Test | NaN | 0.000 | NaN | NaN | 0.000 | NaN |
| MEGSA | 1.000 | 0.044 | 0.085 | 1.000 | 0.044 | 0.084 |
| MEMO | 0.800 | 0.143 | 0.242 | 0.769 | 0.119 | 0.206 |
| WExT | 0.727 | 0.184 | 0.294 | 0.778 | 0.161 | 0.267 |

(e) Metrics for LUSC data. (485 samples | 22 CGC-CGC pairs)

| Method | Precision | Sensitivity | F1 Score | $\text{Precision}_{\text{strict}}$ | $\text{Sensitivity}_{\text{strict}}$ | $\text{F1 Score}_{\text{strict}}$ |
|---|---|---|---|---|---|---|
| DISCOVER | 1.0 | 0.095 | 0.174 | 1.0 | 0.095 | 0.174 |
| Fisher's Exact Test | 1.0 | 0.091 | 0.167 | 1.0 | 0.091 | 0.167 |
| MEGSA | 1.0 | 0.273 | 0.429 | 1.0 | 0.273 | 0.429 |
| MEMO | 1.0 | 0.195 | 0.327 | 1.0 | 0.195 | 0.326 |
| WExT | 1.0 | 0.182 | 0.308 | 1.0 | 0.182 | 0.308 |

(f) Metrics for SKCM data. (468 samples | 194 CGC-CGC pairs)

| Method | Precision | Sensitivity | F1 Score | $\text{Precision}_{\text{strict}}$ | $\text{Sensitivity}_{\text{strict}}$ | $\text{F1 Score}_{\text{strict}}$ |
|---|---|---|---|---|---|---|
| DISCOVER | 0.769 | 0.052 | 0.098 | 0.769 | 0.052 | 0.097 |
| Fisher's Exact Test | 1.000 | 0.010 | 0.020 | 1.000 | 0.010 | 0.020 |
| MEGSA | 1.000 | 0.021 | 0.041 | 1.000 | 0.021 | 0.041 |
| WExT | 0.757 | 0.151 | 0.252 | 0.794 | 0.146 | 0.247 |

(g) Metrics for STAD data. (438 samples | 72 CGC-CGC pairs)

| Method | Precision | Sensitivity | F1 Score | $\text{Precision}_{\text{strict}}$ | $\text{Sensitivity}_{\text{strict}}$ | $\text{F1 Score}_{\text{strict}}$ |
|---|---|---|---|---|---|---|
| DISCOVER | 0.556 | 0.072 | 0.127 | 0.571 | 0.058 | 0.105 |
| Fisher's Exact Test | NaN | 0.000 | NaN | NaN | 0.000 | NaN |
| MEGSA | 1.000 | 0.028 | 0.055 | 1.000 | 0.028 | 0.054 |
| WExT | 0.533 | 0.113 | 0.186 | 0.583 | 0.099 | 0.169 |

(h) Metrics for UCEC data. (531 samples | 606 CGC-CGC pairs)

| Method | Precision | Sensitivity | F1 Score | $\text{Precision}_{\text{strict}}$ | $\text{Sensitivity}_{\text{strict}}$ | $\text{F1 Score}_{\text{strict}}$ |
|---|---|---|---|---|---|---|
| DISCOVER | 0.666 | 0.218 | 0.329 | 0.729 | 0.171 | 0.277 |
| Fisher's Exact Test | 0.800 | 0.013 | 0.026 | 0.800 | 0.013 | 0.026 |
| MEGSA | 0.818 | 0.015 | 0.029 | 0.818 | 0.015 | 0.029 |
| WExT | 0.624 | 0.306 | 0.411 | 0.693 | 0.249 | 0.366 |

Table 10: **Results of network-centric ME evaluation framework with $\mathcal{G}$ = HINT, $\mathcal{S}$ = CGC, c= $X_1$, $p_t$ = 0.05, t=20, robustness_iterations = 100**

(a) Metrics for BLCA data. (411 samples | 76 CGC-CGC pairs)

| Method | Precision | Sensitivity | F1 Score | $\text{Precision}_{\text{strict}}$ | $\text{Sensitivity}_{\text{strict}}$ | $\text{F1 Score}_{\text{strict}}$ |
|---|---|---|---|---|---|---|
| DISCOVER | 0.750 | 0.080 | 0.145 | 0.750 | 0.080 | 0.145 |
| Fisher's Exact Test | 1.000 | 0.054 | 0.103 | 1.000 | 0.054 | 0.102 |
| MEGSA | 1.000 | 0.081 | 0.149 | 1.000 | 0.081 | 0.150 |
| MEMO | 0.667 | 0.082 | 0.146 | 0.667 | 0.082 | 0.146 |
| WExT | 0.615 | 0.110 | 0.186 | 0.667 | 0.110 | 0.189 |

(b) Metrics for BRCA data. (1026 samples | 42 CGC-CGC pairs)

| Method | Precision | Sensitivity | F1 Score | $\text{Precision}_{\text{strict}}$ | $\text{Sensitivity}_{\text{strict}}$ | $\text{F1 Score}_{\text{strict}}$ |
|---|---|---|---|---|---|---|
| DISCOVER | 0.643 | 0.450 | 0.529 | 0.652 | 0.375 | 0.476 |
| DISCOVER Strat | 0.750 | 0.405 | 0.526 | 0.789 | 0.405 | 0.535 |
| Fisher's Exact Test | 0.667 | 0.049 | 0.091 | 0.667 | 0.049 | 0.091 |
| MEGSA | 0.800 | 0.095 | 0.170 | 0.800 | 0.095 | 0.170 |
| MEMO | 0.655 | 0.480 | 0.554 | 0.682 | 0.400 | 0.504 |
| WExT | 0.606 | 0.556 | 0.580 | 0.654 | 0.472 | 0.548 |

(c) Metrics for COADREAD data. (498 samples | 198 CGC-CGC pairs)

| Method | Precision | Sensitivity | F1 Score | $\text{Precision}_{\text{strict}}$ | $\text{Sensitivity}_{\text{strict}}$ | $\text{F1 Score}_{\text{strict}}$ |
|---|---|---|---|---|---|---|
| DISCOVER | 0.623 | 0.204 | 0.308 | 0.702 | 0.177 | 0.283 |
| DISCOVER Strat | 0.800 | 0.064 | 0.118 | 0.800 | 0.064 | 0.119 |
| Fisher's Exact Test | 0.757 | 0.071 | 0.130 | 0.788 | 0.066 | 0.122 |
| MEGSA | 0.714 | 0.076 | 0.138 | 0.722 | 0.066 | 0.121 |
| MEMO | 0.620 | 0.302 | 0.406 | 0.597 | 0.212 | 0.313 |
| WExT | 0.636 | 0.367 | 0.466 | 0.716 | 0.304 | 0.427 |

(d) Metrics for LUAD data. (568 samples | 78 CGC-CGC pairs)

| Method | Precision | Sensitivity | F1 Score | $\text{Precision}_{\text{strict}}$ | $\text{Sensitivity}_{\text{strict}}$ | $\text{F1 Score}_{\text{strict}}$ |
|---|---|---|---|---|---|---|
| DISCOVER | 0.625 | 0.067 | 0.121 | 0.625 | 0.067 | 0.121 |
| Fisher's Exact Test | 0.667 | 0.026 | 0.050 | 0.667 | 0.026 | 0.050 |
| MEGSA | 0.667 | 0.026 | 0.050 | 0.667 | 0.026 | 0.050 |
| MEMO | 0.708 | 0.116 | 0.199 | 0.739 | 0.116 | 0.201 |
| WExT | 0.645 | 0.140 | 0.230 | 0.645 | 0.140 | 0.230 |

(e) Metrics for LUSC data. (485 samples | 60 CGC-CGC pairs)

| Method | Precision | Sensitivity | F1 Score | $\text{Precision}_{\text{strict}}$ | $\text{Sensitivity}_{\text{strict}}$ | $\text{F1 Score}_{\text{strict}}$ |
|---|---|---|---|---|---|---|
| DISCOVER | 1.000 | 0.033 | 0.065 | 1.000 | 0.033 | 0.064 |
| Fisher's Exact Test | 1.000 | 0.033 | 0.065 | 1.000 | 0.033 | 0.064 |
| MEGSA | 1.000 | 0.067 | 0.126 | 1.000 | 0.067 | 0.126 |
| MEMO | 0.667 | 0.069 | 0.125 | 0.667 | 0.069 | 0.125 |
| WExT | 0.667 | 0.068 | 0.123 | 0.667 | 0.068 | 0.123 |

(f) Metrics for SKCM data. (468 samples | 438 CGC-CGC pairs)

| Method | Precision | Sensitivity | F1 Score | $\text{Precision}_{\text{strict}}$ | $\text{Sensitivity}_{\text{strict}}$ | $\text{F1 Score}_{\text{strict}}$ |
|---|---|---|---|---|---|---|
| DISCOVER | 0.875 | 0.032 | 0.062 | 0.875 | 0.032 | 0.062 |
| Fisher's Exact Test | NaN | 0.000 | NaN | NaN | 0.000 | NaN |
| MEGSA | 1.000 | 0.014 | 0.027 | 1.000 | 0.014 | 0.028 |
| WExT | 0.644 | 0.077 | 0.138 | 0.640 | 0.076 | 0.136 |

(g) Metrics for STAD data. (438 samples | 150 CGC-CGC pairs)

| Method | Precision | Sensitivity | F1 Score | $\text{Precision}_{\text{strict}}$ | $\text{Sensitivity}_{\text{strict}}$ | $\text{F1 Score}_{\text{strict}}$ |
|---|---|---|---|---|---|---|
| DISCOVER | 0.581 | 0.124 | 0.204 | 0.625 | 0.103 | 0.177 |
| Fisher's Exact Test | 0.800 | 0.026 | 0.051 | 0.800 | 0.026 | 0.050 |
| MEGSA | 0.800 | 0.027 | 0.052 | 0.800 | 0.027 | 0.052 |
| WExT | 0.600 | 0.185 | 0.283 | 0.629 | 0.151 | 0.244 |

(h) Metrics for UCEC data. (531 samples | 1024 CGC-CGC pairs)

| Method | Precision | Sensitivity | F1 Score | $\text{Precision}_{\text{strict}}$ | $\text{Sensitivity}_{\text{strict}}$ | $\text{F1 Score}_{\text{strict}}$ |
|---|---|---|---|---|---|---|
| DISCOVER | 0.658 | 0.209 | 0.317 | 0.733 | 0.170 | 0.276 |
| Fisher's Exact Test | 0.846 | 0.011 | 0.021 | 0.846 | 0.011 | 0.022 |
| MEGSA | 0.778 | 0.014 | 0.027 | 0.824 | 0.014 | 0.028 |
| WExT | 0.628 | 0.304 | 0.410 | 0.704 | 0.255 | 0.374 |

Table 11: **Results of network-centric ME evaluation framework with $\mathcal{G}$ = STRING, $\mathcal{S}$ = CGC, c= $X_1$, $p_t$ = 0.05, t=20, robustness_iterations = 100**

(a) Metrics for BLCA data. (411 samples | 122 CGC-CGC pairs)

| Method | Precision | Sensitivity | F1 Score | $\text{Precision}_{\text{strict}}$ | $\text{Sensitivity}_{\text{strict}}$ | $\text{F1 Score}_{\text{strict}}$ |
|---|---|---|---|---|---|---|
| DISCOVER | 0.800 | 0.067 | 0.124 | 0.800 | 0.067 | 0.124 |
| Fisher's Exact Test | 1.000 | 0.034 | 0.065 | 1.000 | 0.034 | 0.066 |
| MEGSA | 1.000 | 0.033 | 0.065 | 1.000 | 0.033 | 0.064 |
| MEMO | 0.667 | 0.067 | 0.122 | 0.667 | 0.067 | 0.122 |
| WExT | 0.714 | 0.085 | 0.152 | 0.714 | 0.085 | 0.152 |

(b) Metrics for BRCA data. (1026 samples | 94 CGC-CGC pairs)

| Method | Precision | Sensitivity | F1 Score | $\text{Precision}_{\text{strict}}$ | $\text{Sensitivity}_{\text{strict}}$ | $\text{F1 Score}_{\text{strict}}$ |
|---|---|---|---|---|---|---|
| DISCOVER | 0.667 | 0.322 | 0.434 | 0.714 | 0.287 | 0.409 |
| DISCOVER Strat | 0.730 | 0.314 | 0.439 | 0.812 | 0.302 | 0.440 |
| Fisher's Exact Test | 0.667 | 0.022 | 0.042 | 0.667 | 0.022 | 0.043 |
| MEGSA | 0.800 | 0.043 | 0.081 | 0.800 | 0.043 | 0.082 |
| MEMO | 0.628 | 0.309 | 0.414 | 0.657 | 0.263 | 0.376 |
| WExT | 0.618 | 0.380 | 0.470 | 0.667 | 0.325 | 0.437 |

(c) Metrics for COADREAD data. (498 samples | 292 CGC-CGC pairs)

| Method | Precision | Sensitivity | F1 Score | $\text{Precision}_{\text{strict}}$ | $\text{Sensitivity}_{\text{strict}}$ | $\text{F1 Score}_{\text{strict}}$ |
|---|---|---|---|---|---|---|
| DISCOVER | 0.652 | 0.215 | 0.323 | 0.750 | 0.194 | 0.308 |
| DISCOVER Strat | 0.754 | 0.081 | 0.146 | 0.807 | 0.081 | 0.147 |
| Fisher's Exact Test | 0.882 | 0.052 | 0.097 | 0.933 | 0.048 | 0.091 |
| MEGSA | 0.842 | 0.055 | 0.103 | 0.882 | 0.052 | 0.098 |
| MEMO | 0.613 | 0.245 | 0.350 | 0.639 | 0.192 | 0.295 |
| WExT | 0.577 | 0.246 | 0.345 | 0.670 | 0.214 | 0.324 |

(d) Metrics for LUAD data. (568 samples | 110 CGC-CGC pairs)

| Method | Precision | Sensitivity | F1 Score | $\text{Precision}_{\text{strict}}$ | $\text{Sensitivity}_{\text{strict}}$ | $\text{F1 Score}_{\text{strict}}$ |
|---|---|---|---|---|---|---|
| DISCOVER | 0.750 | 0.113 | 0.196 | 0.923 | 0.113 | 0.201 |
| Fisher's Exact Test | 1.000 | 0.073 | 0.137 | 1.000 | 0.073 | 0.136 |
| MEGSA | 1.000 | 0.073 | 0.137 | 1.000 | 0.073 | 0.136 |
| MEMO | 0.600 | 0.115 | 0.193 | 0.706 | 0.115 | 0.198 |
| WExT | 0.621 | 0.176 | 0.274 | 0.750 | 0.176 | 0.285 |

(e) Metrics for LUSC data. (485 samples | 90 CGC-CGC pairs)

| Method | Precision | Sensitivity | F1 Score | $\text{Precision}_{\text{strict}}$ | $\text{Sensitivity}_{\text{strict}}$ | $\text{F1 Score}_{\text{strict}}$ |
|---|---|---|---|---|---|---|
| DISCOVER | 1.0 | 0.045 | 0.087 | 1.0 | 0.045 | 0.086 |
| Fisher's Exact Test | 1.0 | 0.022 | 0.044 | 1.0 | 0.022 | 0.043 |
| MEGSA | 1.0 | 0.047 | 0.089 | 1.0 | 0.047 | 0.090 |
| MEMO | 1.0 | 0.047 | 0.090 | 1.0 | 0.047 | 0.090 |
| WExT | 1.0 | 0.071 | 0.133 | 1.0 | 0.071 | 0.133 |

(f) Metrics for SKCM data. (468 samples | 560 CGC-CGC pairs)

| Method | Precision | Sensitivity | F1 Score | $\text{Precision}_{\text{strict}}$ | $\text{Sensitivity}_{\text{strict}}$ | $\text{F1 Score}_{\text{strict}}$ |
|---|---|---|---|---|---|---|
| DISCOVER | 0.806 | 0.046 | 0.087 | 0.828 | 0.044 | 0.084 |
| Fisher's Exact Test | 1.000 | 0.004 | 0.007 | 1.000 | 0.004 | 0.008 |
| MEGSA | 0.889 | 0.015 | 0.029 | 0.889 | 0.015 | 0.030 |
| WExT | 0.686 | 0.112 | 0.192 | 0.717 | 0.105 | 0.183 |

(g) Metrics for STAD data. (438 samples | 194 CGC-CGC pairs)

| Method | Precision | Sensitivity | F1 Score | $\text{Precision}_{\text{strict}}$ | $\text{Sensitivity}_{\text{strict}}$ | $\text{F1 Score}_{\text{strict}}$ |
|---|---|---|---|---|---|---|
| DISCOVER | 0.636 | 0.113 | 0.192 | 0.667 | 0.097 | 0.169 |
| Fisher's Exact Test | 1.000 | 0.010 | 0.021 | 1.000 | 0.010 | 0.020 |
| MEGSA | 0.667 | 0.010 | 0.020 | 0.667 | 0.010 | 0.020 |
| WExT | 0.605 | 0.139 | 0.227 | 0.639 | 0.123 | 0.206 |

(h) Metrics for UCEC data. (531 samples | 1566 CGC-CGC pairs)

| Method | Precision | Sensitivity | F1 Score | $\text{Precision}_{\text{strict}}$ | $\text{Sensitivity}_{\text{strict}}$ | $\text{F1 Score}_{\text{strict}}$ |
|---|---|---|---|---|---|---|
| DISCOVER | 0.672 | 0.170 | 0.271 | 0.745 | 0.140 | 0.236 |
| Fisher's Exact Test | 1.000 | 0.008 | 0.015 | 1.000 | 0.008 | 0.016 |
| MEGSA | 1.000 | 0.008 | 0.016 | 1.000 | 0.008 | 0.016 |
| WExT | 0.619 | 0.276 | 0.382 | 0.680 | 0.228 | 0.341 |

Table 12: **Degree-normalized network-centric evaluations $X_1$ and $t = 20$**

(a) Metrics for BLCA data.

| Method | Precision | Sensitivity | F1 Score | $\text{Precision}_{\text{strict}}$ | $\text{Sensitivity}_{\text{strict}}$ | $\text{F1 Score}_{\text{strict}}$ |
|---|---|---|---|---|---|---|
| DISCOVER | 0.875 | 0.432 | 0.531 | 0.875 | 0.432 | 0.531 |
| Fisher's Exact Test | 1.0 | 0.238 | 0.375 | 1.0 | 0.238 | 0.375 |
| MEGSA | 1.0 | 0.494 | 0.604 | 1.0 | 0.494 | 0.604 |
| MEMO | 0.667 | 0.343 | 0.523 | 0.667 | 0.343 | 0.523 |
| WExT | 0.667 | 0.345 | 0.525 | 0.667 | 0.345 | 0.525 |

(b) Metrics for BRCA data.

| Method | Precision | Sensitivity | F1 Score | $\text{Precision}_{\text{strict}}$ | $\text{Sensitivity}_{\text{strict}}$ | $\text{F1 Score}_{\text{strict}}$ |
|---|---|---|---|---|---|---|
| DISCOVER | 0.786 | 0.758 | 0.821 | 0.767 | 0.683 | 0.853 |
| DISCOVER Strat | 0.847 | 0.8 | 0.88 | 0.875 | 0.783 | 0.889 |
| Fisher's Exact Test | 1.0 | 0.6 | 0.667 | 1.0 | 0.6 | 0.667 |
| MEGSA | 1.0 | 0.6 | 0.667 | 1.0 | 0.6 | 0.667 |
| MEMO | 0.716 | 0.717 | 0.83 | 0.767 | 0.683 | 0.853 |
| WExT | 0.705 | 0.724 | 0.744 | 0.731 | 0.647 | 0.88 |

(c) Metrics for COADREAD data.

| Method | Precision | Sensitivity | F1 Score | $\text{Precision}_{\text{strict}}$ | $\text{Sensitivity}_{\text{strict}}$ | $\text{F1 Score}_{\text{strict}}$ |
|---|---|---|---|---|---|---|
| DISCOVER | 0.859 | 0.656 | 0.669 | 0.891 | 0.61 | 0.673 |
| DISCOVER Strat | 0.906 | 0.366 | 0.477 | 0.906 | 0.366 | 0.477 |
| Fisher's Exact Test | 0.819 | 0.3 | 0.377 | 0.819 | 0.3 | 0.377 |
| MEGSA | 0.865 | 0.484 | 0.55 | 0.852 | 0.472 | 0.537 |
| MEMO | 0.853 | 0.739 | 0.731 | 0.848 | 0.644 | 0.739 |
| WExT | 0.866 | 0.718 | 0.743 | 0.891 | 0.668 | 0.742 |

(d) Metrics for LUAD data.

| Method | Precision | Sensitivity | F1 Score | $\text{Precision}_{\text{strict}}$ | $\text{Sensitivity}_{\text{strict}}$ | $\text{F1 Score}_{\text{strict}}$ |
|---|---|---|---|---|---|---|
| DISCOVER | 0.923 | 0.727 | 0.756 | 0.889 | 0.653 | 0.706 |
| Fisher's Exact Test | 0.0 | 0.0 | NaN | 0.0 | 0.0 | NaN |
| MEGSA | 0.667 | 0.208 | 0.311 | 0.667 | 0.208 | 0.311 |
| MEMO | 0.85 | 0.699 | 0.703 | 0.89 | 0.659 | 0.715 |
| WExT | 0.769 | 0.633 | 0.762 | 0.803 | 0.607 | 0.778 |

(e) Metrics for LUSC data.

| Method | Precision | Sensitivity | F1 Score | $\text{Precision}_{\text{strict}}$ | $\text{Sensitivity}_{\text{strict}}$ | $\text{F1 Score}_{\text{strict}}$ |
|---|---|---|---|---|---|---|
| DISCOVER | 1.0 | 1.0 | 1.0 | 1.0 | 1.0 | 1.0 |
| Fisher's Exact Test | 1.0 | 0.75 | 0.833 | 1.0 | 0.75 | 0.833 |
| MEGSA | 1.0 | 0.75 | 0.833 | 1.0 | 0.75 | 0.833 |
| MEMO | 1.0 | 0.75 | 0.833 | 1.0 | 0.75 | 0.833 |
| WExT | 1.0 | 0.75 | 0.833 | 1.0 | 0.75 | 0.833 |

(f) Metrics for SKCM data.

| Method | Precision | Sensitivity | F1 Score | $\text{Precision}_{\text{strict}}$ | $\text{Sensitivity}_{\text{strict}}$ | $\text{F1 Score}_{\text{strict}}$ |
|---|---|---|---|---|---|---|
| DISCOVER | 0.892 | 0.353 | 0.477 | 0.892 | 0.353 | 0.477 |
| Fisher's Exact Test | 1.0 | 0.417 | 0.583 | 1.0 | 0.417 | 0.583 |
| MEGSA | 0.889 | 0.318 | 0.477 | 0.889 | 0.318 | 0.477 |
| WExT | 0.794 | 0.392 | 0.5 | 0.793 | 0.38 | 0.494 |

(g) Metrics for STAD data.

| Method | Precision | Sensitivity | F1 Score | $\text{Precision}_{\text{strict}}$ | $\text{Sensitivity}_{\text{strict}}$ | $\text{F1 Score}_{\text{strict}}$ |
|---|---|---|---|---|---|---|
| DISCOVER | 0.833 | 0.115 | 0.539 | 0.779 | 0.097 | 0.173 |
| Fisher's Exact Test | 0.000 | 0.000 | NaN | 0.000 | 0.000 | NaN |
| MEGSA | 0.750 | 0.010 | 0.375 | 0.769 | 0.010 | 0.020 |
| WExT | 0.829 | 0.190 | 0.585 | 0.799 | 0.173 | 0.284 |

(h) Metrics for UCEC data.

| Method | Precision | Sensitivity | F1 Score | $\text{Precision}_{\text{strict}}$ | $\text{Sensitivity}_{\text{strict}}$ | $\text{F1 Score}_{\text{strict}}$ |
|---|---|---|---|---|---|---|
| DISCOVER | 0.863 | 0.37 | 0.458 | 0.87 | 0.336 | 0.454 |
| Fisher's Exact Test | 0.939 | 0.126 | 0.208 | 0.939 | 0.126 | 0.208 |
| MEGSA | 0.946 | 0.174 | 0.268 | 0.946 | 0.174 | 0.268 |
| WExT | 0.78 | 0.464 | 0.55 | 0.781 | 0.411 | 0.541 |

Table 13: **Results of network-centric ME evaluation framework with $\mathcal{G}$ = Intact (w conf. threshold 0.35), $\mathcal{S}$ = CGC, c= $X_2$, $p_t$ = 0.05, t=20, robustness_iterations = 100**

(a) Metrics for BLCA data. (411 samples | 24 CGC-CGC pairs)

| Method | Precision | Sensitivity | F1 Score | $\text{Precision}_{\text{strict}}$ | $\text{Sensitivity}_{\text{strict}}$ | $\text{F1 Score}_{\text{strict}}$ |
|---|---|---|---|---|---|---|
| DISCOVER | 1.000 | 0.083 | 0.154 | 1.000 | 0.083 | 0.153 |
| Fisher's Exact Test | 1.000 | 0.042 | 0.080 | 1.000 | 0.042 | 0.081 |
| MEGSA | 1.000 | 0.083 | 0.154 | 1.000 | 0.083 | 0.153 |
| MEMO | 0.667 | 0.083 | 0.148 | 0.667 | 0.083 | 0.148 |
| WExT | 0.500 | 0.083 | 0.143 | 0.500 | 0.083 | 0.142 |

(b) Metrics for BRCA data. (1026 samples | 9 CGC-CGC pairs)

| Method | Precision | Sensitivity | F1 Score | $\text{Precision}_{\text{strict}}$ | $\text{Sensitivity}_{\text{strict}}$ | $\text{F1 Score}_{\text{strict}}$ |
|---|---|---|---|---|---|---|
| DISCOVER | 0.625 | 0.556 | 0.588 | 0.714 | 0.556 | 0.625 |
| DISCOVER Strat | 0.714 | 0.556 | 0.625 | 0.714 | 0.556 | 0.625 |
| Fisher's Exact Test | NaN | 0.000 | NaN | NaN | 0.000 | NaN |
| MEGSA | NaN | 0.000 | NaN | NaN | 0.000 | NaN |
| MEMO | 0.625 | 0.556 | 0.588 | 0.714 | 0.556 | 0.625 |
| WExT | 0.625 | 0.556 | 0.588 | 0.714 | 0.556 | 0.625 |

(c) Metrics for COADREAD data. (498 samples | 107 CGC-CGC pairs)

| Method | Precision | Sensitivity | F1 Score | $\text{Precision}_{\text{strict}}$ | $\text{Sensitivity}_{\text{strict}}$ | $\text{F1 Score}_{\text{strict}}$ |
|---|---|---|---|---|---|---|
| DISCOVER | 0.537 | 0.276 | 0.365 | 0.579 | 0.210 | 0.308 |
| DISCOVER Strat | 0.455 | 0.048 | 0.086 | 0.400 | 0.038 | 0.069 |
| Fisher's Exact Test | 0.444 | 0.038 | 0.069 | 0.375 | 0.028 | 0.052 |
| MEGSA | 0.571 | 0.075 | 0.133 | 0.538 | 0.066 | 0.118 |
| MEMO | 0.566 | 0.388 | 0.460 | 0.495 | 0.215 | 0.300 |
| WExT | 0.575 | 0.438 | 0.497 | 0.596 | 0.295 | 0.395 |

(d) Metrics for LUAD data. (568 samples | 54 CGC-CGC pairs)

| Method | Precision | Sensitivity | F1 Score | $\text{Precision}_{\text{strict}}$ | $\text{Sensitivity}_{\text{strict}}$ | $\text{F1 Score}_{\text{strict}}$ |
|---|---|---|---|---|---|---|
| DISCOVER | 0.583 | 0.123 | 0.203 | 0.600 | 0.105 | 0.179 |
| Fisher's Exact Test | NaN | 0.000 | NaN | NaN | 0.000 | NaN |
| MEGSA | 0.400 | 0.034 | 0.062 | 0.400 | 0.034 | 0.063 |
| MEMO | 0.688 | 0.193 | 0.301 | 0.714 | 0.175 | 0.281 |
| WExT | 0.684 | 0.228 | 0.342 | 0.688 | 0.193 | 0.301 |

(e) Metrics for LUSC data. (485 samples | 22 CGC-CGC pairs)

| Method | Precision | Sensitivity | F1 Score | $\text{Precision}_{\text{strict}}$ | $\text{Sensitivity}_{\text{strict}}$ | $\text{F1 Score}_{\text{strict}}$ |
|---|---|---|---|---|---|---|
| DISCOVER | NaN | 0.000 | NaN | NaN | 0.000 | NaN |
| Fisher's Exact Test | NaN | 0.000 | NaN | NaN | 0.000 | NaN |
| MEGSA | 1.000 | 0.136 | 0.240 | 1.000 | 0.136 | 0.239 |
| MEMO | 0.667 | 0.087 | 0.154 | 0.667 | 0.087 | 0.154 |
| WExT | 0.400 | 0.087 | 0.143 | 0.500 | 0.087 | 0.148 |

(f) Metrics for SKCM data. (468 samples | 313 CGC-CGC pairs)

| Method | Precision | Sensitivity | F1 Score | $\text{Precision}_{\text{strict}}$ | $\text{Sensitivity}_{\text{strict}}$ | $\text{F1 Score}_{\text{strict}}$ |
|---|---|---|---|---|---|---|
| DISCOVER | 0.757 | 0.045 | 0.085 | 0.757 | 0.045 | 0.085 |
| Fisher's Exact Test | 1.000 | 0.006 | 0.013 | 1.000 | 0.006 | 0.012 |
| MEGSA | 0.923 | 0.019 | 0.038 | 0.923 | 0.019 | 0.037 |
| WExT | 0.679 | 0.118 | 0.201 | 0.714 | 0.115 | 0.198 |

(g) Metrics for STAD data. (438 samples | 70 CGC-CGC pairs)

| Method | Precision | Sensitivity | F1 Score | $\text{Precision}_{\text{strict}}$ | $\text{Sensitivity}_{\text{strict}}$ | $\text{F1 Score}_{\text{strict}}$ |
|---|---|---|---|---|---|---|
| DISCOVER | 0.600 | 0.129 | 0.212 | 0.727 | 0.114 | 0.197 |
| Fisher's Exact Test | NaN | 0.000 | NaN | NaN | 0.000 | NaN |
| MEGSA | 1.000 | 0.014 | 0.028 | 1.000 | 0.014 | 0.028 |
| WExT | 0.696 | 0.229 | 0.344 | 0.824 | 0.200 | 0.322 |

(h) Metrics for UCEC data. (531 samples | 1179 CGC-CGC pairs)

| Method | Precision | Sensitivity | F1 Score | $\text{Precision}_{\text{strict}}$ | $\text{Sensitivity}_{\text{strict}}$ | $\text{F1 Score}_{\text{strict}}$ |
|---|---|---|---|---|---|---|
| DISCOVER | 0.649 | 0.184 | 0.287 | 0.744 | 0.154 | 0.255 |
| Fisher's Exact Test | 0.833 | 0.008 | 0.017 | 0.833 | 0.008 | 0.016 |
| MEGSA | 0.793 | 0.010 | 0.019 | 0.793 | 0.010 | 0.020 |
| WExT | 0.607 | 0.286 | 0.389 | 0.686 | 0.241 | 0.357 |

Table 14: **Results of network-centric ME evaluation framework with $\mathcal{G}$ = Intact (w conf. threshold 0.35), $\mathcal{S}$ = CGC, c= $X_2$, $p_t$ = 0.05, t=20, robustness_iterations = 300**

(a) Metrics for BLCA data. (411 samples | 24 CGC-CGC pairs)

| Method | Precision | Sensitivity | F1 Score | $\text{Precision}_{\text{strict}}$ | $\text{Sensitivity}_{\text{strict}}$ | $\text{F1 Score}_{\text{strict}}$ |
|---|---|---|---|---|---|---|
| DISCOVER | 1.000 | 0.083 | 0.154 | 1.000 | 0.083 | 0.153 |
| Fisher's Exact Test | 1.000 | 0.042 | 0.080 | 1.000 | 0.042 | 0.081 |
| MEGSA | 1.000 | 0.083 | 0.154 | 1.000 | 0.083 | 0.153 |
| MEMO | 0.667 | 0.083 | 0.148 | 0.667 | 0.083 | 0.148 |
| WExT | 0.500 | 0.083 | 0.143 | 0.500 | 0.083 | 0.142 |

(b) Metrics for BRCA data. (1026 samples | 9 CGC-CGC pairs)

| Method | Precision | Sensitivity | F1 Score | $\text{Precision}_{\text{strict}}$ | $\text{Sensitivity}_{\text{strict}}$ | $\text{F1 Score}_{\text{strict}}$ |
|---|---|---|---|---|---|---|
| DISCOVER | 0.625 | 0.556 | 0.588 | 0.714 | 0.556 | 0.625 |
| DISCOVER Strat | 0.714 | 0.556 | 0.625 | 0.714 | 0.556 | 0.625 |
| Fisher's Exact Test | NaN | 0.000 | NaN | NaN | 0.000 | NaN |
| MEGSA | NaN | 0.000 | NaN | NaN | 0.000 | NaN |
| MEMO | 0.625 | 0.556 | 0.588 | 0.714 | 0.556 | 0.625 |
| WExT | 0.625 | 0.556 | 0.588 | 0.714 | 0.556 | 0.625 |

(c) Metrics for COADREAD data. (498 samples | 107 CGC-CGC pairs)

| Method | Precision | Sensitivity | F1 Score | $\text{Precision}_{\text{strict}}$ | $\text{Sensitivity}_{\text{strict}}$ | $\text{F1 Score}_{\text{strict}}$ |
|---|---|---|---|---|---|---|
| DISCOVER | 0.536 | 0.278 | 0.366 | 0.575 | 0.213 | 0.311 |
| DISCOVER Strat | 0.500 | 0.056 | 0.101 | 0.500 | 0.056 | 0.101 |
| Fisher's Exact Test | 0.429 | 0.029 | 0.054 | 0.429 | 0.029 | 0.054 |
| MEGSA | 0.571 | 0.075 | 0.132 | 0.538 | 0.065 | 0.116 |
| MEMO | 0.554 | 0.383 | 0.453 | 0.500 | 0.224 | 0.309 |
| WExT | 0.578 | 0.441 | 0.500 | 0.619 | 0.308 | 0.411 |

(d) Metrics for LUAD data. (568 samples | 54 CGC-CGC pairs)

| Method | Precision | Sensitivity | F1 Score | $\text{Precision}_{\text{strict}}$ | $\text{Sensitivity}_{\text{strict}}$ | $\text{F1 Score}_{\text{strict}}$ |
|---|---|---|---|---|---|---|
| DISCOVER | 0.571 | 0.138 | 0.222 | 0.545 | 0.103 | 0.173 |
| Fisher's Exact Test | NaN | 0.000 | NaN | NaN | 0.000 | NaN |
| MEGSA | 0.400 | 0.034 | 0.062 | 0.400 | 0.034 | 0.063 |
| MEMO | 0.688 | 0.193 | 0.301 | 0.714 | 0.175 | 0.281 |
| WExT | 0.706 | 0.214 | 0.329 | 0.733 | 0.196 | 0.309 |

(e) Metrics for LUSC data. (485 samples | 22 CGC-CGC pairs)

| Method | Precision | Sensitivity | F1 Score | $\text{Precision}_{\text{strict}}$ | $\text{Sensitivity}_{\text{strict}}$ | $\text{F1 Score}_{\text{strict}}$ |
|---|---|---|---|---|---|---|
| DISCOVER | NaN | 0.000 | NaN | NaN | 0.000 | NaN |
| Fisher's Exact Test | NaN | 0.000 | NaN | NaN | 0.000 | NaN |
| MEGSA | 1.0 | 0.136 | 0.240 | 1.0 | 0.136 | 0.239 |
| MEMO | 1.0 | 0.095 | 0.174 | 1.0 | 0.095 | 0.174 |
| WExT | 0.4 | 0.095 | 0.154 | 0.5 | 0.095 | 0.160 |

(f) Metrics for SKCM data. (468 samples | 313 CGC-CGC pairs)

| Method | Precision | Sensitivity | F1 Score | $\text{Precision}_{\text{strict}}$ | $\text{Sensitivity}_{\text{strict}}$ | $\text{F1 Score}_{\text{strict}}$ |
|---|---|---|---|---|---|---|
| DISCOVER | 0.778 | 0.045 | 0.085 | 0.778 | 0.045 | 0.085 |
| Fisher's Exact Test | 1.000 | 0.006 | 0.013 | 1.000 | 0.006 | 0.012 |
| MEGSA | 0.857 | 0.019 | 0.037 | 0.857 | 0.019 | 0.037 |
| WExT | 0.667 | 0.119 | 0.202 | 0.700 | 0.116 | 0.199 |

(g) Metrics for STAD data. (438 samples | 70 CGC-CGC pairs)

| Method | Precision | Sensitivity | F1 Score | $\text{Precision}_{\text{strict}}$ | $\text{Sensitivity}_{\text{strict}}$ | $\text{F1 Score}_{\text{strict}}$ |
|---|---|---|---|---|---|---|
| DISCOVER | 0.600 | 0.129 | 0.212 | 0.636 | 0.100 | 0.173 |
| Fisher's Exact Test | NaN | 0.000 | NaN | NaN | 0.000 | NaN |
| MEGSA | 1.000 | 0.014 | 0.028 | 1.000 | 0.014 | 0.028 |
| WExT | 0.696 | 0.229 | 0.344 | 0.824 | 0.200 | 0.322 |

(h) Metrics for UCEC data. (531 samples | 1179 CGC-CGC pairs)

| Method | Precision | Sensitivity | F1 Score | $\text{Precision}_{\text{strict}}$ | $\text{Sensitivity}_{\text{strict}}$ | $\text{F1 Score}_{\text{strict}}$ |
|---|---|---|---|---|---|---|
| DISCOVER | 0.651 | 0.184 | 0.286 | 0.756 | 0.155 | 0.257 |
| Fisher's Exact Test | 0.833 | 0.008 | 0.017 | 0.833 | 0.008 | 0.016 |
| MEGSA | 0.846 | 0.009 | 0.018 | 0.846 | 0.009 | 0.018 |
| WExT | 0.601 | 0.286 | 0.388 | 0.668 | 0.238 | 0.351 |

Table 15: **Results of network-centric ME evaluation framework with $\mathcal{G}$ = Intact (w conf. threshold 0.35), $\mathcal{S}$ = CGC, c= $X_2$, $p_t$ = 0.05, t=20, robustness_iterations = 500**

(a) Metrics for BLCA data. (411 samples | 24 CGC-CGC pairs)

| Method | Precision | Sensitivity | F1 Score | $\text{Precision}_{\text{strict}}$ | $\text{Sensitivity}_{\text{strict}}$ | $\text{F1 Score}_{\text{strict}}$ |
|---|---|---|---|---|---|---|
| DISCOVER | 1.000 | 0.083 | 0.154 | 1.000 | 0.083 | 0.153 |
| Fisher's Exact Test | 1.000 | 0.042 | 0.080 | 1.000 | 0.042 | 0.081 |
| MEGSA | 1.000 | 0.083 | 0.154 | 1.000 | 0.083 | 0.153 |
| MEMO | 0.667 | 0.083 | 0.148 | 0.667 | 0.083 | 0.148 |
| WExT | 0.500 | 0.083 | 0.143 | 0.500 | 0.083 | 0.142 |

(b) Metrics for BRCA data. (1026 samples | 9 CGC-CGC pairs)

| Method | Precision | Sensitivity | F1 Score | $\text{Precision}_{\text{strict}}$ | $\text{Sensitivity}_{\text{strict}}$ | $\text{F1 Score}_{\text{strict}}$ |
|---|---|---|---|---|---|---|
| DISCOVER | 0.625 | 0.556 | 0.588 | 0.714 | 0.556 | 0.625 |
| DISCOVER Strat | 0.714 | 0.556 | 0.625 | 0.714 | 0.556 | 0.625 |
| Fisher's Exact Test | NaN | 0.000 | NaN | NaN | 0.000 | NaN |
| MEGSA | NaN | 0.000 | NaN | NaN | 0.000 | NaN |
| MEMO | 0.625 | 0.556 | 0.588 | 0.714 | 0.556 | 0.625 |
| WExT | 0.625 | 0.556 | 0.588 | 0.714 | 0.556 | 0.625 |

(c) Metrics for COADREAD data. (498 samples | 107 CGC-CGC pairs)

| Method | Precision | Sensitivity | F1 Score | $\text{Precision}_{\text{strict}}$ | $\text{Sensitivity}_{\text{strict}}$ | $\text{F1 Score}_{\text{strict}}$ |
|---|---|---|---|---|---|---|
| DISCOVER | 0.536 | 0.280 | 0.368 | 0.575 | 0.215 | 0.313 |
| DISCOVER Strat | 0.500 | 0.052 | 0.094 | 0.500 | 0.052 | 0.094 |
| Fisher's Exact Test | 0.500 | 0.038 | 0.070 | 0.500 | 0.038 | 0.071 |
| MEGSA | 0.615 | 0.075 | 0.134 | 0.583 | 0.066 | 0.119 |
| MEMO | 0.568 | 0.396 | 0.467 | 0.489 | 0.217 | 0.301 |
| WExT | 0.575 | 0.438 | 0.497 | 0.596 | 0.295 | 0.395 |

(d) Metrics for LUAD data. (568 samples | 54 CGC-CGC pairs)

| Method | Precision | Sensitivity | F1 Score | $\text{Precision}_{\text{strict}}$ | $\text{Sensitivity}_{\text{strict}}$ | $\text{F1 Score}_{\text{strict}}$ |
|---|---|---|---|---|---|---|
| DISCOVER | 0.571 | 0.138 | 0.222 | 0.545 | 0.103 | 0.173 |
| Fisher's Exact Test | NaN | 0.000 | NaN | NaN | 0.000 | NaN |
| MEGSA | 0.400 | 0.034 | 0.062 | 0.400 | 0.034 | 0.063 |
| MEMO | 0.688 | 0.193 | 0.301 | 0.714 | 0.175 | 0.281 |
| WExT | 0.684 | 0.228 | 0.342 | 0.688 | 0.193 | 0.301 |

(e) Metrics for LUSC data. (485 samples | 22 CGC-CGC pairs)

| Method | Precision | Sensitivity | F1 Score | $\text{Precision}_{\text{strict}}$ | $\text{Sensitivity}_{\text{strict}}$ | $\text{F1 Score}_{\text{strict}}$ |
|---|---|---|---|---|---|---|
| DISCOVER | NaN | 0.000 | NaN | NaN | 0.000 | NaN |
| Fisher's Exact Test | NaN | 0.000 | NaN | NaN | 0.000 | NaN |
| MEGSA | 1.0 | 0.136 | 0.240 | 1.000 | 0.136 | 0.239 |
| MEMO | 1.0 | 0.095 | 0.174 | 1.000 | 0.095 | 0.174 |
| WExT | 0.5 | 0.091 | 0.154 | 0.667 | 0.091 | 0.160 |

(f) Metrics for SKCM data. (468 samples | 313 CGC-CGC pairs)

| Method | Precision | Sensitivity | F1 Score | $\text{Precision}_{\text{strict}}$ | $\text{Sensitivity}_{\text{strict}}$ | $\text{F1 Score}_{\text{strict}}$ |
|---|---|---|---|---|---|---|
| DISCOVER | 0.737 | 0.045 | 0.084 | 0.737 | 0.045 | 0.085 |
| Fisher's Exact Test | 1.000 | 0.006 | 0.013 | 1.000 | 0.006 | 0.012 |
| MEGSA | 0.857 | 0.019 | 0.037 | 0.857 | 0.019 | 0.037 |
| WExT | 0.679 | 0.118 | 0.201 | 0.714 | 0.115 | 0.198 |

(g) Metrics for STAD data. (438 samples | 70 CGC-CGC pairs)

| Method | Precision | Sensitivity | F1 Score | $\text{Precision}_{\text{strict}}$ | $\text{Sensitivity}_{\text{strict}}$ | $\text{F1 Score}_{\text{strict}}$ |
|---|---|---|---|---|---|---|
| DISCOVER | 0.615 | 0.116 | 0.195 | 0.700 | 0.101 | 0.177 |
| Fisher's Exact Test | NaN | 0.000 | NaN | NaN | 0.000 | NaN |
| MEGSA | 1.000 | 0.014 | 0.028 | 1.000 | 0.014 | 0.028 |
| WExT | 0.696 | 0.229 | 0.344 | 0.824 | 0.200 | 0.322 |

(h) Metrics for UCEC data. (531 samples | 1179 CGC-CGC pairs)

| Method | Precision | Sensitivity | F1 Score | $\text{Precision}_{\text{strict}}$ | $\text{Sensitivity}_{\text{strict}}$ | $\text{F1 Score}_{\text{strict}}$ |
|---|---|---|---|---|---|---|
| DISCOVER | 0.653 | 0.184 | 0.287 | 0.755 | 0.155 | 0.257 |
| Fisher's Exact Test | 0.833 | 0.008 | 0.017 | 0.833 | 0.008 | 0.016 |
| MEGSA | 0.786 | 0.009 | 0.018 | 0.786 | 0.009 | 0.018 |
| WExT | 0.603 | 0.286 | 0.388 | 0.683 | 0.242 | 0.357 |

Table 16: **Results of network-centric ME evaluation framework with $\mathcal{G}$ = Intact (w conf. threshold 0.35), $\mathcal{S}$ = CGC, c= $X_2$, $p_t$ = 0.01, t=20, robustness_iterations = 100**

(a) Metrics for BLCA data. (411 samples | 24 CGC-CGC pairs)

| Method | Precision | Sensitivity | F1 Score | $\text{Precision}_{\text{strict}}$ | $\text{Sensitivity}_{\text{strict}}$ | $\text{F1 Score}_{\text{strict}}$ |
|---|---|---|---|---|---|---|
| DISCOVER | 1.0 | 0.042 | 0.08 | 1.0 | 0.042 | 0.081 |
| Fisher's Exact Test | NaN | 0.000 | NaN | NaN | 0.000 | NaN |
| MEGSA | NaN | 0.000 | NaN | NaN | 0.000 | NaN |
| MEMO | 1.0 | 0.042 | 0.08 | 1.0 | 0.042 | 0.081 |
| WExT | 1.0 | 0.042 | 0.08 | 1.0 | 0.042 | 0.081 |

(b) Metrics for BRCA data. (1026 samples | 9 CGC-CGC pairs)

| Method | Precision | Sensitivity | F1 Score | $\text{Precision}_{\text{strict}}$ | $\text{Sensitivity}_{\text{strict}}$ | $\text{F1 Score}_{\text{strict}}$ |
|---|---|---|---|---|---|---|
| DISCOVER | 0.75 | 0.333 | 0.462 | 0.75 | 0.333 | 0.461 |
| DISCOVER Strat | 0.50 | 0.111 | 0.182 | 0.50 | 0.111 | 0.182 |
| Fisher's Exact Test | NaN | 0.000 | NaN | NaN | 0.000 | NaN |
| MEGSA | NaN | 0.000 | NaN | NaN | 0.000 | NaN |
| MEMO | 0.60 | 0.333 | 0.429 | 0.60 | 0.333 | 0.428 |
| WExT | 0.60 | 0.333 | 0.429 | 0.60 | 0.333 | 0.428 |

(c) Metrics for COADREAD data. (498 samples | 107 CGC-CGC pairs)

| Method | Precision | Sensitivity | F1 Score | $\text{Precision}_{\text{strict}}$ | $\text{Sensitivity}_{\text{strict}}$ | $\text{F1 Score}_{\text{strict}}$ |
|---|---|---|---|---|---|---|
| DISCOVER | 0.613 | 0.178 | 0.275 | 0.625 | 0.140 | 0.229 |
| DISCOVER Strat | 0.000 | 0.000 | NaN | 0.000 | 0.000 | NaN |
| Fisher's Exact Test | 0.000 | 0.000 | NaN | 0.000 | 0.000 | NaN |
| MEGSA | 0.500 | 0.009 | 0.018 | 0.500 | 0.009 | 0.018 |
| MEMO | 0.596 | 0.321 | 0.417 | 0.474 | 0.170 | 0.250 |
| WExT | 0.603 | 0.330 | 0.427 | 0.590 | 0.217 | 0.317 |

(d) Metrics for LUAD data. (568 samples | 54 CGC-CGC pairs)

| Method | Precision | Sensitivity | F1 Score | $\text{Precision}_{\text{strict}}$ | $\text{Sensitivity}_{\text{strict}}$ | $\text{F1 Score}_{\text{strict}}$ |
|---|---|---|---|---|---|---|
| DISCOVER | 0.750 | 0.051 | 0.095 | 0.750 | 0.051 | 0.096 |
| Fisher's Exact Test | NaN | 0.000 | NaN | NaN | 0.000 | NaN |
| MEGSA | 1.000 | 0.034 | 0.066 | 1.000 | 0.034 | 0.066 |
| MEMO | 0.778 | 0.119 | 0.206 | 0.750 | 0.102 | 0.180 |
| WExT | 0.640 | 0.137 | 0.225 | 0.667 | 0.120 | 0.203 |

(e) Metrics for LUSC data. (485 samples | 22 CGC-CGC pairs)

| Method | Precision | Sensitivity | F1 Score | $\text{Precision}_{\text{strict}}$ | $\text{Sensitivity}_{\text{strict}}$ | $\text{F1 Score}_{\text{strict}}$ |
|---|---|---|---|---|---|---|
| DISCOVER | NaN | 0.0 | NaN | NaN | 0.0 | NaN |
| Fisher's Exact Test | NaN | 0.0 | NaN | NaN | 0.0 | NaN |
| MEGSA | NaN | 0.0 | NaN | NaN | 0.0 | NaN |
| MEMO | NaN | 0.0 | NaN | NaN | 0.0 | NaN |
| WExT | NaN | 0.0 | NaN | NaN | 0.0 | NaN |

(f) Metrics for SKCM data. (468 samples | 313 CGC-CGC pairs)

| Method | Precision | Sensitivity | F1 Score | $\text{Precision}_{\text{strict}}$ | $\text{Sensitivity}_{\text{strict}}$ | $\text{F1 Score}_{\text{strict}}$ |
|---|---|---|---|---|---|---|
| DISCOVER | 0.833 | 0.016 | 0.031 | 0.833 | 0.016 | 0.031 |
| Fisher's Exact Test | 1.000 | 0.006 | 0.013 | 1.000 | 0.006 | 0.012 |
| MEGSA | 1.000 | 0.006 | 0.013 | 1.000 | 0.006 | 0.012 |
| WExT | 0.710 | 0.035 | 0.067 | 0.710 | 0.035 | 0.067 |

(g) Metrics for STAD data. (438 samples | 70 CGC-CGC pairs)

| Method | Precision | Sensitivity | F1 Score | $\text{Precision}_{\text{strict}}$ | $\text{Sensitivity}_{\text{strict}}$ | $\text{F1 Score}_{\text{strict}}$ |
|---|---|---|---|---|---|---|
| DISCOVER | 0.625 | 0.071 | 0.128 | 0.667 | 0.057 | 0.105 |
| Fisher's Exact Test | NaN | 0.000 | NaN | NaN | 0.000 | NaN |
| MEGSA | NaN | 0.000 | NaN | NaN | 0.000 | NaN |
| WExT | 0.583 | 0.101 | 0.173 | 0.714 | 0.072 | 0.131 |

(h) Metrics for UCEC data. (531 samples | 1179 CGC-CGC pairs)

| Method | Precision | Sensitivity | F1 Score | $\text{Precision}_{\text{strict}}$ | $\text{Sensitivity}_{\text{strict}}$ | $\text{F1 Score}_{\text{strict}}$ |
|---|---|---|---|---|---|---|
| DISCOVER | 0.673 | 0.129 | 0.217 | 0.782 | 0.113 | 0.197 |
| Fisher's Exact Test | 1.000 | 0.005 | 0.010 | 1.000 | 0.005 | 0.010 |
| MEGSA | 1.000 | 0.005 | 0.010 | 1.000 | 0.005 | 0.010 |
| WExT | 0.640 | 0.205 | 0.311 | 0.744 | 0.170 | 0.277 |

Table 17: **Results of network-centric ME evaluation framework with $\mathcal{G}$ = Intact (w conf. threshold 0.35), $\mathcal{S}$ = CGC, c= $X_2$, $p_t$ = 0.1, t=20, robustness_iterations = 100**

(a) Metrics for BLCA data. (411 samples | 24 CGC-CGC pairs)

| Method | Precision | Sensitivity | F1 Score | $\text{Precision}_{\text{strict}}$ | $\text{Sensitivity}_{\text{strict}}$ | $\text{F1 Score}_{\text{strict}}$ |
|---|---|---|---|---|---|---|
| DISCOVER | 0.667 | 0.083 | 0.148 | 0.667 | 0.083 | 0.148 |
| Fisher's Exact Test | 1.000 | 0.042 | 0.080 | 1.000 | 0.042 | 0.081 |
| MEGSA | 1.000 | 0.083 | 0.154 | 1.000 | 0.083 | 0.153 |
| MEMO | 0.500 | 0.083 | 0.143 | 0.500 | 0.083 | 0.142 |
| WExT | 0.400 | 0.083 | 0.138 | 0.400 | 0.083 | 0.137 |

(b) Metrics for BRCA data. (1026 samples | 9 CGC-CGC pairs)

| Method | Precision | Sensitivity | F1 Score | $\text{Precision}_{\text{strict}}$ | $\text{Sensitivity}_{\text{strict}}$ | $\text{F1 Score}_{\text{strict}}$ |
|---|---|---|---|---|---|---|
| DISCOVER | 0.625 | 0.556 | 0.588 | 0.714 | 0.556 | 0.625 |
| DISCOVER Strat | 0.667 | 0.667 | 0.667 | 0.714 | 0.556 | 0.625 |
| Fisher's Exact Test | NaN | 0.000 | NaN | NaN | 0.000 | NaN |
| MEGSA | NaN | 0.000 | NaN | NaN | 0.000 | NaN |
| MEMO | 0.625 | 0.556 | 0.588 | 0.714 | 0.556 | 0.625 |
| WExT | 0.667 | 0.667 | 0.667 | 0.714 | 0.556 | 0.625 |

(c) Metrics for COADREAD data. (498 samples | 107 CGC-CGC pairs)

| Method | Precision | Sensitivity | F1 Score | $\text{Precision}_{\text{strict}}$ | $\text{Sensitivity}_{\text{strict}}$ | $\text{F1 Score}_{\text{strict}}$ |
|---|---|---|---|---|---|---|
| DISCOVER | 0.529 | 0.333 | 0.409 | 0.553 | 0.241 | 0.336 |
| DISCOVER Strat | 0.547 | 0.136 | 0.218 | 0.535 | 0.108 | 0.180 |
| Fisher's Exact Test | 0.650 | 0.124 | 0.208 | 0.588 | 0.095 | 0.164 |
| MEGSA | 0.577 | 0.142 | 0.227 | 0.550 | 0.104 | 0.175 |
| MEMO | 0.578 | 0.449 | 0.505 | 0.509 | 0.262 | 0.346 |
| WExT | 0.558 | 0.457 | 0.503 | 0.600 | 0.314 | 0.412 |

(d) Metrics for LUAD data. (568 samples | 54 CGC-CGC pairs)

| Method | Precision | Sensitivity | F1 Score | $\text{Precision}_{\text{strict}}$ | $\text{Sensitivity}_{\text{strict}}$ | $\text{F1 Score}_{\text{strict}}$ |
|---|---|---|---|---|---|---|
| DISCOVER | 0.686 | 0.212 | 0.324 | 0.710 | 0.195 | 0.306 |
| Fisher's Exact Test | 1.000 | 0.034 | 0.067 | 1.000 | 0.034 | 0.066 |
| MEGSA | 0.500 | 0.086 | 0.147 | 0.556 | 0.086 | 0.149 |
| MEMO | 0.636 | 0.237 | 0.346 | 0.684 | 0.220 | 0.333 |
| WExT | 0.667 | 0.339 | 0.449 | 0.692 | 0.305 | 0.423 |

(e) Metrics for LUSC data. (485 samples | 22 CGC-CGC pairs)

| Method | Precision | Sensitivity | F1 Score | $\text{Precision}_{\text{strict}}$ | $\text{Sensitivity}_{\text{strict}}$ | $\text{F1 Score}_{\text{strict}}$ |
|---|---|---|---|---|---|---|
| DISCOVER | 0.500 | 0.100 | 0.167 | 0.667 | 0.100 | 0.174 |
| Fisher's Exact Test | NaN | 0.000 | NaN | NaN | 0.000 | NaN |
| MEGSA | 0.500 | 0.130 | 0.207 | 0.600 | 0.130 | 0.214 |
| MEMO | 0.364 | 0.089 | 0.143 | 0.444 | 0.089 | 0.148 |
| WExT | 0.500 | 0.136 | 0.214 | 0.600 | 0.136 | 0.222 |

(f) Metrics for SKCM data. (468 samples | 313 CGC-CGC pairs)

| Method | Precision | Sensitivity | F1 Score | $\text{Precision}_{\text{strict}}$ | $\text{Sensitivity}_{\text{strict}}$ | $\text{F1 Score}_{\text{strict}}$ |
|---|---|---|---|---|---|---|
| DISCOVER | 0.697 | 0.101 | 0.176 | 0.713 | 0.101 | 0.177 |
| Fisher's Exact Test | 1.000 | 0.006 | 0.013 | 1.000 | 0.006 | 0.012 |
| MEGSA | 0.818 | 0.029 | 0.056 | 0.818 | 0.029 | 0.056 |
| WExT | 0.616 | 0.176 | 0.274 | 0.650 | 0.173 | 0.273 |

(g) Metrics for STAD data. (438 samples | 70 CGC-CGC pairs)

| Method | Precision | Sensitivity | F1 Score | $\text{Precision}_{\text{strict}}$ | $\text{Sensitivity}_{\text{strict}}$ | $\text{F1 Score}_{\text{strict}}$ |
|---|---|---|---|---|---|---|
| DISCOVER | 0.696 | 0.229 | 0.344 | 0.765 | 0.186 | 0.299 |
| Fisher's Exact Test | 0.500 | 0.014 | 0.028 | 0.500 | 0.014 | 0.027 |
| MEGSA | 0.500 | 0.014 | 0.027 | 0.500 | 0.014 | 0.027 |
| WExT | 0.704 | 0.271 | 0.392 | 0.800 | 0.229 | 0.356 |

(h) Metrics for UCEC data. (531 samples | 1179 CGC-CGC pairs)

| Method | Precision | Sensitivity | F1 Score | $\text{Precision}_{\text{strict}}$ | $\text{Sensitivity}_{\text{strict}}$ | $\text{F1 Score}_{\text{strict}}$ |
|---|---|---|---|---|---|---|
| DISCOVER | 0.622 | 0.226 | 0.331 | 0.708 | 0.186 | 0.295 |
| Fisher's Exact Test | 0.857 | 0.015 | 0.030 | 0.850 | 0.014 | 0.028 |
| MEGSA | 0.862 | 0.021 | 0.041 | 0.889 | 0.020 | 0.039 |
| WExT | 0.584 | 0.353 | 0.440 | 0.657 | 0.294 | 0.406 |

Table 18: **Results of network-centric ME evaluation framework with $\mathcal{G}$ = Intact (w conf. threshold 0.35), $\mathcal{S}$ = $CGC_{SNV}$ , c= $X_2$, $p_t$ = 0.05, t=20, robustness_iterations = 100**

(a) Metrics for BLCA data. (411 samples | 21 $CGC_{SNV}$-$CGC_{SNV}$ pairs)

| Method | Precision | Sensitivity | F1 Score | $\text{Precision}_{\text{strict}}$ | $\text{Sensitivity}_{\text{strict}}$ | $\text{F1 Score}_{\text{strict}}$ |
|---|---|---|---|---|---|---|
| DISCOVER | 1.00 | 0.143 | 0.250 | 1.00 | 0.143 | 0.250 |
| Fisher's Exact Test | 1.00 | 0.095 | 0.174 | 1.00 | 0.095 | 0.174 |
| MEGSA | 1.00 | 0.143 | 0.250 | 1.00 | 0.143 | 0.250 |
| MEMO | 1.00 | 0.146 | 0.255 | 1.00 | 0.146 | 0.255 |
| WExT | 0.75 | 0.143 | 0.240 | 0.75 | 0.143 | 0.240 |

(b) Metrics for BRCA data. (1026 samples | 10 $CGC_{SNV}$-$CGC_{SNV}$ pairs)

| Method | Precision | Sensitivity | F1 Score | $\text{Precision}_{\text{strict}}$ | $\text{Sensitivity}_{\text{strict}}$ | $\text{F1 Score}_{\text{strict}}$ |
|---|---|---|---|---|---|---|
| DISCOVER | 0.571 | 0.4 | 0.471 | 0.667 | 0.4 | 0.5 |
| DISCOVER Strat | 0.667 | 0.4 | 0.500 | 0.667 | 0.4 | 0.5 |
| Fisher's Exact Test | NaN | 0.0 | NaN | NaN | 0.0 | NaN |
| MEGSA | NaN | 0.0 | NaN | NaN | 0.0 | NaN |
| MEMO | 0.571 | 0.4 | 0.471 | 0.667 | 0.4 | 0.5 |
| WExT | 0.571 | 0.4 | 0.471 | 0.667 | 0.4 | 0.5 |

(c) Metrics for COADREAD data. (498 samples | 101 $CGC_{SNV}$-$CGC_{SNV}$ pairs)

| Method | Precision | Sensitivity | F1 Score | $\text{Precision}_{\text{strict}}$ | $\text{Sensitivity}_{\text{strict}}$ | $\text{F1 Score}_{\text{strict}}$ |
|---|---|---|---|---|---|---|
| DISCOVER | 0.581 | 0.248 | 0.347 | 0.645 | 0.198 | 0.303 |
| DISCOVER Strat | 0.600 | 0.060 | 0.109 | 0.600 | 0.060 | 0.109 |
| Fisher's Exact Test | 0.500 | 0.040 | 0.073 | 0.500 | 0.040 | 0.074 |
| MEGSA | 0.667 | 0.080 | 0.143 | 0.636 | 0.070 | 0.126 |
| MEMO | 0.593 | 0.327 | 0.421 | 0.583 | 0.214 | 0.313 |
| WExT | 0.612 | 0.408 | 0.490 | 0.696 | 0.318 | 0.437 |

(d) Metrics for LUAD data. (568 samples | 46 $CGC_{SNV}$-$CGC_{SNV}$ pairs)

| Method | Precision | Sensitivity | F1 Score | $\text{Precision}_{\text{strict}}$ | $\text{Sensitivity}_{\text{strict}}$ | $\text{F1 Score}_{\text{strict}}$ |
|---|---|---|---|---|---|---|
| DISCOVER | 0.545 | 0.136 | 0.218 | 0.556 | 0.114 | 0.189 |
| Fisher's Exact Test | NaN | 0.000 | NaN | NaN | 0.000 | NaN |
| MEGSA | 0.500 | 0.043 | 0.080 | 0.500 | 0.043 | 0.079 |
| MEMO | 0.690 | 0.227 | 0.342 | 0.720 | 0.205 | 0.319 |
| WExT | 0.719 | 0.258 | 0.380 | 0.741 | 0.225 | 0.345 |

(e) Metrics for LUSC data. (485 samples | 22 $CGC_{SNV}$-$CGC_{SNV}$ pairs)

| Method | Precision | Sensitivity | F1 Score | $\text{Precision}_{\text{strict}}$ | $\text{Sensitivity}_{\text{strict}}$ | $\text{F1 Score}_{\text{strict}}$ |
|---|---|---|---|---|---|---|
| DISCOVER | NaN | 0.000 | NaN | NaN | 0.000 | NaN |
| Fisher's Exact Test | NaN | 0.000 | NaN | NaN | 0.000 | NaN |
| MEGSA | 1.000 | 0.136 | 0.240 | 1.000 | 0.136 | 0.239 |
| MEMO | 0.667 | 0.091 | 0.160 | 0.667 | 0.091 | 0.160 |
| WExT | 0.400 | 0.087 | 0.143 | 0.500 | 0.087 | 0.148 |

(f) Metrics for SKCM data. (468 samples | 228 $CGC_{SNV}$-$CGC_{SNV}$ pairs)

| Method | Precision | Sensitivity | F1 Score | $\text{Precision}_{\text{strict}}$ | $\text{Sensitivity}_{\text{strict}}$ | $\text{F1 Score}_{\text{strict}}$ |
|---|---|---|---|---|---|---|
| DISCOVER | 0.842 | 0.035 | 0.068 | 0.842 | 0.035 | 0.067 |
| Fisher's Exact Test | 1.000 | 0.009 | 0.017 | 1.000 | 0.009 | 0.018 |
| MEGSA | 0.800 | 0.018 | 0.034 | 0.800 | 0.018 | 0.035 |
| WExT | 0.699 | 0.129 | 0.218 | 0.747 | 0.125 | 0.214 |

(g) Metrics for STAD data. (438 samples | 53 $CGC_{SNV}$-$CGC_{SNV}$ pairs)

| Method | Precision | Sensitivity | F1 Score | $\text{Precision}_{\text{strict}}$ | $\text{Sensitivity}_{\text{strict}}$ | $\text{F1 Score}_{\text{strict}}$ |
|---|---|---|---|---|---|---|
| DISCOVER | 0.579 | 0.107 | 0.180 | 0.692 | 0.087 | 0.155 |
| Fisher's Exact Test | NaN | 0.000 | NaN | NaN | 0.000 | NaN |
| MEGSA | 1.000 | 0.019 | 0.037 | 1.000 | 0.019 | 0.037 |
| WExT | 0.667 | 0.226 | 0.338 | 0.769 | 0.189 | 0.303 |

(h) Metrics for UCEC data. (531 samples | 739 $CGC_{SNV}$-$CGC_{SNV}$ pairs)

| Method | Precision | Sensitivity | F1 Score | $\text{Precision}_{\text{strict}}$ | $\text{Sensitivity}_{\text{strict}}$ | $\text{F1 Score}_{\text{strict}}$ |
|---|---|---|---|---|---|---|
| DISCOVER | 0.658 | 0.212 | 0.321 | 0.771 | 0.182 | 0.294 |
| Fisher's Exact Test | 0.833 | 0.014 | 0.027 | 0.833 | 0.014 | 0.028 |
| MEGSA | 0.846 | 0.015 | 0.029 | 0.846 | 0.015 | 0.029 |
| WExT | 0.637 | 0.329 | 0.434 | 0.731 | 0.282 | 0.407 |

Table 19: **Results of network-centric ME evaluation framework with $\mathcal{G}$ = Intact (w conf. threshold 0.35), $\mathcal{S}$ = Intogen , c= $X_2$, $p_t$ = 0.05, t=20, robustness_iterations = 100**

(a) Metrics for BLCA data. (411 samples | 23 Intogen-Intogen pairs)

| Method | Precision | Sensitivity | F1 Score | $\text{Precision}_{\text{strict}}$ | $\text{Sensitivity}_{\text{strict}}$ | $\text{F1 Score}_{\text{strict}}$ |
|---|---|---|---|---|---|---|
| DISCOVER | 1.00 | 0.130 | 0.231 | 1.00 | 0.130 | 0.230 |
| Fisher's Exact Test | 1.00 | 0.043 | 0.083 | 1.00 | 0.043 | 0.082 |
| MEGSA | 1.00 | 0.087 | 0.160 | 1.00 | 0.087 | 0.160 |
| MEMO | 0.75 | 0.130 | 0.222 | 0.75 | 0.130 | 0.222 |
| WExT | 0.60 | 0.130 | 0.214 | 0.60 | 0.130 | 0.214 |

(b) Metrics for BRCA data. (1026 samples | 7 Intogen-Intogen pairs)

| Method | Precision | Sensitivity | F1 Score | $\text{Precision}_{\text{strict}}$ | $\text{Sensitivity}_{\text{strict}}$ | $\text{F1 Score}_{\text{strict}}$ |
|---|---|---|---|---|---|---|
| DISCOVER | 0.6 | 0.429 | 0.5 | 0.6 | 0.429 | 0.5 |
| DISCOVER Strat | 0.6 | 0.429 | 0.5 | 0.6 | 0.429 | 0.5 |
| Fisher's Exact Test | NaN | 0.000 | NaN | NaN | 0.000 | NaN |
| MEGSA | NaN | 0.000 | NaN | NaN | 0.000 | NaN |
| MEMO | 0.6 | 0.429 | 0.5 | 0.6 | 0.429 | 0.5 |
| WExT | 0.6 | 0.429 | 0.5 | 0.6 | 0.429 | 0.5 |

(c) Metrics for COADREAD data. (498 samples | 105 Intogen-Intogen pairs)

| Method | Precision | Sensitivity | F1 Score | $\text{Precision}_{\text{strict}}$ | $\text{Sensitivity}_{\text{strict}}$ | $\text{F1 Score}_{\text{strict}}$ |
|---|---|---|---|---|---|---|
| DISCOVER | 0.564 | 0.301 | 0.392 | 0.590 | 0.223 | 0.324 |
| DISCOVER Strat | 0.545 | 0.058 | 0.104 | 0.545 | 0.058 | 0.105 |
| Fisher's Exact Test | 0.400 | 0.038 | 0.070 | 0.333 | 0.029 | 0.053 |
| MEGSA | 0.571 | 0.076 | 0.134 | 0.538 | 0.067 | 0.119 |
| MEMO | 0.581 | 0.417 | 0.486 | 0.562 | 0.262 | 0.357 |
| WExT | 0.590 | 0.467 | 0.521 | 0.630 | 0.324 | 0.428 |

(d) Metrics for LUAD data. (568 samples | 53 Intogen-Intogen pairs)

| Method | Precision | Sensitivity | F1 Score | $\text{Precision}_{\text{strict}}$ | $\text{Sensitivity}_{\text{strict}}$ | $\text{F1 Score}_{\text{strict}}$ |
|---|---|---|---|---|---|---|
| DISCOVER | 0.538 | 0.135 | 0.215 | 0.500 | 0.096 | 0.161 |
| Fisher's Exact Test | NaN | 0.000 | NaN | NaN | 0.000 | NaN |
| MEGSA | 0.250 | 0.019 | 0.035 | 0.250 | 0.019 | 0.035 |
| MEMO | 0.667 | 0.196 | 0.303 | 0.692 | 0.176 | 0.281 |
| WExT | 0.667 | 0.231 | 0.343 | 0.667 | 0.192 | 0.298 |

(e) Metrics for LUSC data. (485 samples | 24 Intogen-Intogen pairs)

| Method | Precision | Sensitivity | F1 Score | $\text{Precision}_{\text{strict}}$ | $\text{Sensitivity}_{\text{strict}}$ | $\text{F1 Score}_{\text{strict}}$ |
|---|---|---|---|---|---|---|
| DISCOVER | 1.0 | 0.042 | 0.080 | 1.0 | 0.042 | 0.081 |
| Fisher's Exact Test | NaN | 0.000 | NaN | NaN | 0.000 | NaN |
| MEGSA | 1.0 | 0.125 | 0.222 | 1.0 | 0.125 | 0.222 |
| MEMO | 1.0 | 0.125 | 0.222 | 1.0 | 0.125 | 0.222 |
| WExT | 0.5 | 0.120 | 0.194 | 0.6 | 0.120 | 0.200 |

(f) Metrics for SKCM data. (468 samples | 335 Intogen-Intogen pairs)

| Method | Precision | Sensitivity | F1 Score | $\text{Precision}_{\text{strict}}$ | $\text{Sensitivity}_{\text{strict}}$ | $\text{F1 Score}_{\text{strict}}$ |
|---|---|---|---|---|---|---|
| DISCOVER | 0.600 | 0.036 | 0.067 | 0.600 | 0.036 | 0.068 |
| Fisher's Exact Test | 1.000 | 0.006 | 0.012 | 1.000 | 0.006 | 0.012 |
| MEGSA | 0.750 | 0.018 | 0.035 | 0.750 | 0.018 | 0.035 |
| WExT | 0.615 | 0.110 | 0.186 | 0.642 | 0.107 | 0.183 |

(g) Metrics for STAD data. (438 samples | 65 Intogen-Intogen pairs)

| Method | Precision | Sensitivity | F1 Score | $\text{Precision}_{\text{strict}}$ | $\text{Sensitivity}_{\text{strict}}$ | $\text{F1 Score}_{\text{strict}}$ |
|---|---|---|---|---|---|---|
| DISCOVER | 0.600 | 0.138 | 0.225 | 0.636 | 0.108 | 0.185 |
| Fisher's Exact Test | NaN | 0.000 | NaN | NaN | 0.000 | NaN |
| MEGSA | 1.000 | 0.015 | 0.030 | 1.000 | 0.015 | 0.030 |
| WExT | 0.696 | 0.250 | 0.368 | 0.781 | 0.195 | 0.312 |

(h) Metrics for UCEC data. (531 samples | 1111 Intogen-Intogen pairs)

| Method | Precision | Sensitivity | F1 Score | $\text{Precision}_{\text{strict}}$ | $\text{Sensitivity}_{\text{strict}}$ | $\text{F1 Score}_{\text{strict}}$ |
|---|---|---|---|---|---|---|
| DISCOVER | 0.657 | 0.214 | 0.323 | 0.771 | 0.183 | 0.296 |
| Fisher's Exact Test | 0.875 | 0.013 | 0.025 | 0.875 | 0.013 | 0.026 |
| MEGSA | 0.882 | 0.014 | 0.027 | 0.882 | 0.014 | 0.028 |
| WExT | 0.616 | 0.323 | 0.424 | 0.707 | 0.271 | 0.392 |

Table 20: **Results of network-centric ME evaluation framework with $\mathcal{G}$ = Intact (w conf. threshold 0.25), $\mathcal{S}$ = CGC, c= $X_2$, $p_t$ = 0.05, t=20, robustness_iterations = 100**

(a) Metrics for BLCA data. (411 samples | 30 CGC-CGC pairs)

| Method | Precision | Sensitivity | F1 Score | $\text{Precision}_{\text{strict}}$ | $\text{Sensitivity}_{\text{strict}}$ | $\text{F1 Score}_{\text{strict}}$ |
|---|---|---|---|---|---|---|
| DISCOVER | 1.00 | 0.100 | 0.182 | 1.00 | 0.100 | 0.182 |
| Fisher's Exact Test | 1.00 | 0.067 | 0.125 | 1.00 | 0.067 | 0.126 |
| MEGSA | 1.00 | 0.133 | 0.235 | 1.00 | 0.133 | 0.235 |
| MEMO | 0.75 | 0.100 | 0.176 | 0.75 | 0.100 | 0.176 |
| WExT | 0.80 | 0.133 | 0.229 | 0.80 | 0.133 | 0.228 |

(b) Metrics for BRCA data. (1026 samples | 15 CGC-CGC pairs)

| Method | Precision | Sensitivity | F1 Score | $\text{Precision}_{\text{strict}}$ | $\text{Sensitivity}_{\text{strict}}$ | $\text{F1 Score}_{\text{strict}}$ |
|---|---|---|---|---|---|---|
| DISCOVER | 0.900 | 0.600 | 0.720 | 1.0 | 0.600 | 0.75 |
| DISCOVER Strat | 1.000 | 0.600 | 0.750 | 1.0 | 0.600 | 0.75 |
| Fisher's Exact Test | NaN | 0.000 | NaN | NaN | 0.000 | NaN |
| MEGSA | NaN | 0.000 | NaN | NaN | 0.000 | NaN |
| MEMO | 0.900 | 0.600 | 0.720 | 1.0 | 0.600 | 0.75 |
| WExT | 0.909 | 0.667 | 0.769 | 1.0 | 0.667 | 0.80 |

(c) Metrics for COADREAD data. (498 samples | 177 CGC-CGC pairs)

| Method | Precision | Sensitivity | F1 Score | $\text{Precision}_{\text{strict}}$ | $\text{Sensitivity}_{\text{strict}}$ | $\text{F1 Score}_{\text{strict}}$ |
|---|---|---|---|---|---|---|
| DISCOVER | 0.560 | 0.237 | 0.333 | 0.611 | 0.186 | 0.285 |
| DISCOVER Strat | 0.545 | 0.068 | 0.121 | 0.550 | 0.062 | 0.111 |
| Fisher's Exact Test | 0.600 | 0.068 | 0.122 | 0.611 | 0.062 | 0.113 |
| MEGSA | 0.655 | 0.102 | 0.177 | 0.667 | 0.091 | 0.160 |
| MEMO | 0.585 | 0.335 | 0.426 | 0.519 | 0.198 | 0.287 |
| WExT | 0.602 | 0.371 | 0.459 | 0.635 | 0.269 | 0.378 |

(d) Metrics for LUAD data. (568 samples | 80 CGC-CGC pairs)

| Method | Precision | Sensitivity | F1 Score | $\text{Precision}_{\text{strict}}$ | $\text{Sensitivity}_{\text{strict}}$ | $\text{F1 Score}_{\text{strict}}$ |
|---|---|---|---|---|---|---|
| DISCOVER | 0.632 | 0.150 | 0.242 | 0.688 | 0.138 | 0.230 |
| Fisher's Exact Test | NaN | 0.000 | NaN | NaN | 0.000 | NaN |
| MEGSA | 0.571 | 0.051 | 0.093 | 0.571 | 0.051 | 0.094 |
| MEMO | 0.680 | 0.215 | 0.327 | 0.714 | 0.190 | 0.300 |
| WExT | 0.679 | 0.232 | 0.345 | 0.708 | 0.207 | 0.320 |

(e) Metrics for LUSC data. (485 samples | 31 CGC-CGC pairs)

| Method | Precision | Sensitivity | F1 Score | $\text{Precision}_{\text{strict}}$ | $\text{Sensitivity}_{\text{strict}}$ | $\text{F1 Score}_{\text{strict}}$ |
|---|---|---|---|---|---|---|
| DISCOVER | 1.00 | 0.033 | 0.065 | 1.00 | 0.033 | 0.064 |
| Fisher's Exact Test | 1.00 | 0.032 | 0.062 | 1.00 | 0.032 | 0.062 |
| MEGSA | 1.00 | 0.097 | 0.176 | 1.00 | 0.097 | 0.177 |
| MEMO | 1.00 | 0.103 | 0.188 | 1.00 | 0.103 | 0.187 |
| WExT | 0.75 | 0.100 | 0.176 | 0.75 | 0.100 | 0.176 |

(f) Metrics for SKCM data. (468 samples | 522 CGC-CGC pairs)

| Method | Precision | Sensitivity | F1 Score | $\text{Precision}_{\text{strict}}$ | $\text{Sensitivity}_{\text{strict}}$ | $\text{F1 Score}_{\text{strict}}$ |
|---|---|---|---|---|---|---|
| DISCOVER | 0.814 | 0.046 | 0.088 | 0.814 | 0.046 | 0.087 |
| Fisher's Exact Test | 1.000 | 0.004 | 0.008 | 1.000 | 0.004 | 0.008 |
| MEGSA | 0.778 | 0.013 | 0.026 | 0.778 | 0.013 | 0.026 |
| WExT | 0.691 | 0.101 | 0.176 | 0.716 | 0.099 | 0.174 |

(g) Metrics for STAD data. (438 samples | 107 CGC-CGC pairs)

| Method | Precision | Sensitivity | F1 Score | $\text{Precision}_{\text{strict}}$ | $\text{Sensitivity}_{\text{strict}}$ | $\text{F1 Score}_{\text{strict}}$ |
|---|---|---|---|---|---|---|
| DISCOVER | 0.621 | 0.171 | 0.269 | 0.750 | 0.143 | 0.240 |
| Fisher's Exact Test | 1.000 | 0.019 | 0.037 | 1.000 | 0.019 | 0.037 |
| MEGSA | 0.800 | 0.037 | 0.071 | 0.800 | 0.037 | 0.071 |
| WExT | 0.619 | 0.243 | 0.349 | 0.724 | 0.196 | 0.308 |

(h) Metrics for UCEC data. (531 samples | 1794 CGC-CGC pairs)

| Method | Precision | Sensitivity | F1 Score | $\text{Precision}_{\text{strict}}$ | $\text{Sensitivity}_{\text{strict}}$ | $\text{F1 Score}_{\text{strict}}$ |
|---|---|---|---|---|---|---|
| DISCOVER | 0.664 | 0.164 | 0.264 | 0.768 | 0.140 | 0.237 |
| Fisher's Exact Test | 0.824 | 0.008 | 0.015 | 0.824 | 0.008 | 0.016 |
| MEGSA | 0.783 | 0.010 | 0.020 | 0.818 | 0.010 | 0.020 |
| WExT | 0.619 | 0.267 | 0.373 | 0.695 | 0.229 | 0.344 |

Table 21: **Results of network-centric ME evaluation framework with $\mathcal{G}$ = Intact (w conf. threshold 0.45), $\mathcal{S}$ = CGC, c= $X_2$, $p_t$ = 0.05, t=20, robustness_iterations = 100**

(a) Metrics for BLCA data. (411 samples | 12 CGC-CGC pairs)

| Method | Precision | Sensitivity | F1 Score | $\text{Precision}_{\text{strict}}$ | $\text{Sensitivity}_{\text{strict}}$ | $\text{F1 Score}_{\text{strict}}$ |
|---|---|---|---|---|---|---|
| DISCOVER | 1.0 | 0.083 | 0.154 | 1.0 | 0.083 | 0.153 |
| Fisher's Exact Test | 1.0 | 0.083 | 0.154 | 1.0 | 0.083 | 0.153 |
| MEGSA | 1.0 | 0.083 | 0.154 | 1.0 | 0.083 | 0.153 |
| MEMO | 0.5 | 0.077 | 0.133 | 0.5 | 0.077 | 0.133 |
| WExT | 0.5 | 0.083 | 0.143 | 0.5 | 0.083 | 0.142 |

(b) Metrics for BRCA data. (1026 samples | 1 CGC-CGC pairs)

| Method | Precision | Sensitivity | F1 Score | $\text{Precision}_{\text{strict}}$ | $\text{Sensitivity}_{\text{strict}}$ | $\text{F1 Score}_{\text{strict}}$ |
|---|---|---|---|---|---|---|
| DISCOVER | 1.0 | 1.0 | 1.0 | 1.0 | 1.0 | 1.0 |
| DISCOVER Strat | 1.0 | 1.0 | 1.0 | 1.0 | 1.0 | 1.0 |
| Fisher's Exact Test | NaN | 0.0 | NaN | NaN | 0.0 | NaN |
| MEGSA | NaN | 0.0 | NaN | NaN | 0.0 | NaN |
| MEMO | 1.0 | 1.0 | 1.0 | 1.0 | 1.0 | 1.0 |
| WExT | 1.0 | 1.0 | 1.0 | 1.0 | 1.0 | 1.0 |

(c) Metrics for COADREAD data. (498 samples | 15 CGC-CGC pairs)

| Method | Precision | Sensitivity | F1 Score | $\text{Precision}_{\text{strict}}$ | $\text{Sensitivity}_{\text{strict}}$ | $\text{F1 Score}_{\text{strict}}$ |
|---|---|---|---|---|---|---|
| DISCOVER | 0.467 | 0.467 | 0.467 | 0.556 | 0.333 | 0.417 |
| DISCOVER Strat | 0.333 | 0.133 | 0.190 | 0.200 | 0.067 | 0.100 |
| Fisher's Exact Test | 0.333 | 0.062 | 0.105 | 0.333 | 0.062 | 0.105 |
| MEGSA | 0.400 | 0.133 | 0.200 | 0.250 | 0.067 | 0.106 |
| MEMO | 0.485 | 0.516 | 0.500 | 0.286 | 0.194 | 0.231 |
| WExT | 0.529 | 0.643 | 0.581 | 0.600 | 0.429 | 0.500 |

(d) Metrics for LUAD data. (568 samples | 12 CGC-CGC pairs)

| Method | Precision | Sensitivity | F1 Score | $\text{Precision}_{\text{strict}}$ | $\text{Sensitivity}_{\text{strict}}$ | $\text{F1 Score}_{\text{strict}}$ |
|---|---|---|---|---|---|---|
| DISCOVER | 0.571 | 0.308 | 0.400 | 0.600 | 0.231 | 0.334 |
| Fisher's Exact Test | NaN | 0.000 | NaN | NaN | 0.000 | NaN |
| MEGSA | 0.667 | 0.167 | 0.267 | 0.667 | 0.167 | 0.267 |
| MEMO | 0.625 | 0.385 | 0.476 | 0.667 | 0.308 | 0.421 |
| WExT | 0.667 | 0.333 | 0.444 | 0.600 | 0.250 | 0.353 |

(e) Metrics for LUSC data. (485 samples | 5 CGC-CGC pairs)

| Method | Precision | Sensitivity | F1 Score | $\text{Precision}_{\text{strict}}$ | $\text{Sensitivity}_{\text{strict}}$ | $\text{F1 Score}_{\text{strict}}$ |
|---|---|---|---|---|---|---|
| DISCOVER | NaN | 0.0 | NaN | NaN | 0.0 | NaN |
| Fisher's Exact Test | NaN | 0.0 | NaN | NaN | 0.0 | NaN |
| MEGSA | 0.667 | 0.4 | 0.500 | 0.5 | 0.2 | 0.286 |
| MEMO | 1.000 | 0.2 | 0.333 | 1.0 | 0.2 | 0.333 |
| WExT | 0.333 | 0.2 | 0.250 | 0.5 | 0.2 | 0.286 |

(f) Metrics for SKCM data. (468 samples | 104 CGC-CGC pairs)

| Method | Precision | Sensitivity | F1 Score | $\text{Precision}_{\text{strict}}$ | $\text{Sensitivity}_{\text{strict}}$ | $\text{F1 Score}_{\text{strict}}$ |
|---|---|---|---|---|---|---|
| DISCOVER | 0.800 | 0.039 | 0.074 | 0.8 | 0.039 | 0.074 |
| Fisher's Exact Test | NaN | 0.000 | NaN | NaN | 0.000 | NaN |
| MEGSA | NaN | 0.000 | NaN | NaN | 0.000 | NaN |
| WExT | 0.714 | 0.144 | 0.240 | 0.7 | 0.135 | 0.226 |

(g) Metrics for STAD data. (438 samples | 16 CGC-CGC pairs)

| Method | Precision | Sensitivity | F1 Score | $\text{Precision}_{\text{strict}}$ | $\text{Sensitivity}_{\text{strict}}$ | $\text{F1 Score}_{\text{strict}}$ |
|---|---|---|---|---|---|---|
| DISCOVER | 0.667 | 0.250 | 0.364 | 1.0 | 0.250 | 0.400 |
| Fisher's Exact Test | NaN | 0.000 | NaN | NaN | 0.000 | NaN |
| MEGSA | 1.000 | 0.062 | 0.118 | 1.0 | 0.062 | 0.117 |
| WExT | 0.571 | 0.250 | 0.348 | 1.0 | 0.250 | 0.400 |

(h) Metrics for UCEC data. (531 samples | 425 CGC-CGC pairs)

| Method | Precision | Sensitivity | F1 Score | $\text{Precision}_{\text{strict}}$ | $\text{Sensitivity}_{\text{strict}}$ | $\text{F1 Score}_{\text{strict}}$ |
|---|---|---|---|---|---|---|
| DISCOVER | 0.614 | 0.230 | 0.334 | 0.713 | 0.182 | 0.290 |
| Fisher's Exact Test | 0.778 | 0.016 | 0.032 | 0.778 | 0.016 | 0.031 |
| MEGSA | 0.818 | 0.021 | 0.041 | 0.818 | 0.021 | 0.041 |
| WExT | 0.565 | 0.319 | 0.408 | 0.633 | 0.250 | 0.358 |

Table 22: **Results of network-centric ME evaluation framework with $\mathcal{G}$ = HINT, $\mathcal{S}$ = CGC, c= $X_2$, $p_t$ = 0.05, t=20, robustness_iterations = 100**

(a) Metrics for BLCA data. (411 samples | 6 CGC-CGC pairs)

| Method | Precision | Sensitivity | F1 Score | $\text{Precision}_{\text{strict}}$ | $\text{Sensitivity}_{\text{strict}}$ | $\text{F1 Score}_{\text{strict}}$ |
|---|---|---|---|---|---|---|
| DISCOVER | NaN | 0.0 | NaN | NaN | 0.0 | NaN |
| Fisher's Exact Test | NaN | 0.0 | NaN | NaN | 0.0 | NaN |
| MEGSA | NaN | 0.0 | NaN | NaN | 0.0 | NaN |
| MEMO | NaN | 0.0 | NaN | NaN | 0.0 | NaN |
| WExT | NaN | 0.0 | NaN | NaN | 0.0 | NaN |

(b) Metrics for BRCA data. (1026 samples | 3 CGC-CGC pairs)

| Method | Precision | Sensitivity | F1 Score | $\text{Precision}_{\text{strict}}$ | $\text{Sensitivity}_{\text{strict}}$ | $\text{F1 Score}_{\text{strict}}$ |
|---|---|---|---|---|---|---|
| DISCOVER | NaN | 0.0 | NaN | NaN | 0.0 | NaN |
| DISCOVER Strat | NaN | 0.0 | NaN | NaN | 0.0 | NaN |
| Fisher's Exact Test | NaN | 0.0 | NaN | NaN | 0.0 | NaN |
| MEGSA | NaN | 0.0 | NaN | NaN | 0.0 | NaN |
| MEMO | NaN | 0.0 | NaN | NaN | 0.0 | NaN |
| WExT | NaN | 0.0 | NaN | NaN | 0.0 | NaN |

(c) Metrics for COADREAD data. (498 samples | 58 CGC-CGC pairs)

| Method | Precision | Sensitivity | F1 Score | $\text{Precision}_{\text{strict}}$ | $\text{Sensitivity}_{\text{strict}}$ | $\text{F1 Score}_{\text{strict}}$ |
|---|---|---|---|---|---|---|
| DISCOVER | 0.455 | 0.273 | 0.341 | 0.478 | 0.200 | 0.282 |
| DISCOVER Strat | 0.364 | 0.069 | 0.116 | 0.333 | 0.052 | 0.090 |
| Fisher's Exact Test | 0.600 | 0.158 | 0.250 | 0.583 | 0.123 | 0.203 |
| MEGSA | 0.600 | 0.155 | 0.247 | 0.583 | 0.121 | 0.200 |
| MEMO | 0.529 | 0.474 | 0.500 | 0.344 | 0.193 | 0.247 |
| WExT | 0.537 | 0.518 | 0.527 | 0.559 | 0.339 | 0.422 |

(d) Metrics for LUAD data. (568 samples | 24 CGC-CGC pairs)

| Method | Precision | Sensitivity | F1 Score | $\text{Precision}_{\text{strict}}$ | $\text{Sensitivity}_{\text{strict}}$ | $\text{F1 Score}_{\text{strict}}$ |
|---|---|---|---|---|---|---|
| DISCOVER | 0.667 | 0.083 | 0.148 | 0.667 | 0.083 | 0.148 |
| Fisher's Exact Test | NaN | 0.000 | NaN | NaN | 0.000 | NaN |
| MEGSA | 0.000 | 0.000 | NaN | 0.000 | 0.000 | NaN |
| MEMO | 0.800 | 0.167 | 0.276 | 0.800 | 0.167 | 0.276 |
| WExT | 0.800 | 0.167 | 0.276 | 0.800 | 0.167 | 0.276 |

(e) Metrics for LUSC data. (485 samples | 9 CGC-CGC pairs)

| Method | Precision | Sensitivity | F1 Score | $\text{Precision}_{\text{strict}}$ | $\text{Sensitivity}_{\text{strict}}$ | $\text{F1 Score}_{\text{strict}}$ |
|---|---|---|---|---|---|---|
| DISCOVER | NaN | 0.000 | NaN | NaN | 0.000 | NaN |
| Fisher's Exact Test | NaN | 0.000 | NaN | NaN | 0.000 | NaN |
| MEGSA | 1.000 | 0.222 | 0.364 | 1.0 | 0.222 | 0.363 |
| MEMO | 1.000 | 0.250 | 0.400 | 1.0 | 0.250 | 0.400 |
| WExT | 0.667 | 0.250 | 0.364 | 1.0 | 0.250 | 0.400 |

(f) Metrics for SKCM data. (468 samples | 237 CGC-CGC pairs)

| Method | Precision | Sensitivity | F1 Score | $\text{Precision}_{\text{strict}}$ | $\text{Sensitivity}_{\text{strict}}$ | $\text{F1 Score}_{\text{strict}}$ |
|---|---|---|---|---|---|---|
| DISCOVER | 0.900 | 0.038 | 0.073 | 0.900 | 0.038 | 0.073 |
| Fisher's Exact Test | NaN | 0.000 | NaN | NaN | 0.000 | NaN |
| MEGSA | 0.800 | 0.017 | 0.033 | 0.800 | 0.017 | 0.033 |
| WExT | 0.667 | 0.085 | 0.152 | 0.655 | 0.081 | 0.144 |

(g) Metrics for STAD data. (438 samples | 33 CGC-CGC pairs)

| Method | Precision | Sensitivity | F1 Score | $\text{Precision}_{\text{strict}}$ | $\text{Sensitivity}_{\text{strict}}$ | $\text{F1 Score}_{\text{strict}}$ |
|---|---|---|---|---|---|---|
| DISCOVER | 0.588 | 0.303 | 0.400 | 0.818 | 0.273 | 0.409 |
| Fisher's Exact Test | 0.750 | 0.091 | 0.162 | 0.750 | 0.091 | 0.162 |
| MEGSA | 0.750 | 0.091 | 0.162 | 0.750 | 0.091 | 0.162 |
| WExT | 0.609 | 0.424 | 0.500 | 0.786 | 0.333 | 0.468 |

(h) Metrics for UCEC data. (531 samples | 721 CGC-CGC pairs)

| Method | Precision | Sensitivity | F1 Score | $\text{Precision}_{\text{strict}}$ | $\text{Sensitivity}_{\text{strict}}$ | $\text{F1 Score}_{\text{strict}}$ |
|---|---|---|---|---|---|---|
| DISCOVER | 0.629 | 0.233 | 0.340 | 0.748 | 0.193 | 0.307 |
| Fisher's Exact Test | 0.714 | 0.014 | 0.027 | 0.769 | 0.014 | 0.027 |
| MEGSA | 0.733 | 0.015 | 0.030 | 0.786 | 0.015 | 0.029 |
| WExT | 0.605 | 0.343 | 0.438 | 0.713 | 0.285 | 0.407 |

Table 23: **Results of network-centric ME evaluation framework with $\mathcal{G}$ = STRING, $\mathcal{S}$ = CGC, c= $X_2$, $p_t$ = 0.05, t=20, robustness_iterations = 100**

(a) Metrics for BLCA data. (411 samples | 13 CGC-CGC pairs)

| Method | Precision | Sensitivity | F1 Score | $\text{Precision}_{\text{strict}}$ | $\text{Sensitivity}_{\text{strict}}$ | $\text{F1 Score}_{\text{strict}}$ |
|---|---|---|---|---|---|---|
| DISCOVER | 0.500 | 0.071 | 0.125 | 0.500 | 0.071 | 0.124 |
| Fisher's Exact Test | NaN | 0.000 | NaN | NaN | 0.000 | NaN |
| MEGSA | NaN | 0.000 | NaN | NaN | 0.000 | NaN |
| MEMO | 0.500 | 0.077 | 0.133 | 0.500 | 0.077 | 0.133 |
| WExT | 0.667 | 0.080 | 0.143 | 0.667 | 0.080 | 0.143 |

(b) Metrics for BRCA data. (1026 samples | 7 CGC-CGC pairs)

| Method | Precision | Sensitivity | F1 Score | $\text{Precision}_{\text{strict}}$ | $\text{Sensitivity}_{\text{strict}}$ | $\text{F1 Score}_{\text{strict}}$ |
|---|---|---|---|---|---|---|
| DISCOVER | 0.333 | 0.143 | 0.200 | 0.5 | 0.143 | 0.222 |
| DISCOVER Strat | 0.500 | 0.143 | 0.222 | 1.0 | 0.143 | 0.250 |
| Fisher's Exact Test | NaN | 0.000 | NaN | NaN | 0.000 | NaN |
| MEGSA | NaN | 0.000 | NaN | NaN | 0.000 | NaN |
| MEMO | 0.333 | 0.143 | 0.200 | 0.5 | 0.143 | 0.222 |
| WExT | 0.333 | 0.143 | 0.200 | 0.5 | 0.143 | 0.222 |

(c) Metrics for COADREAD data. (498 samples | 116 CGC-CGC pairs)

| Method | Precision | Sensitivity | F1 Score | $\text{Precision}_{\text{strict}}$ | $\text{Sensitivity}_{\text{strict}}$ | $\text{F1 Score}_{\text{strict}}$ |
|---|---|---|---|---|---|---|
| DISCOVER | 0.581 | 0.218 | 0.317 | 0.767 | 0.201 | 0.319 |
| DISCOVER Strat | 0.650 | 0.113 | 0.192 | 0.722 | 0.113 | 0.195 |
| Fisher's Exact Test | 1.000 | 0.096 | 0.175 | 1.000 | 0.096 | 0.175 |
| MEGSA | 1.000 | 0.113 | 0.203 | 1.000 | 0.113 | 0.203 |
| MEMO | 0.551 | 0.237 | 0.331 | 0.469 | 0.132 | 0.206 |
| WExT | 0.547 | 0.252 | 0.345 | 0.714 | 0.217 | 0.333 |

(d) Metrics for LUAD data. (568 samples | 49 CGC-CGC pairs)

| Method | Precision | Sensitivity | F1 Score | $\text{Precision}_{\text{strict}}$ | $\text{Sensitivity}_{\text{strict}}$ | $\text{F1 Score}_{\text{strict}}$ |
|---|---|---|---|---|---|---|
| DISCOVER | 0.750 | 0.184 | 0.295 | 0.818 | 0.184 | 0.300 |
| Fisher's Exact Test | 1.000 | 0.102 | 0.185 | 1.000 | 0.102 | 0.185 |
| MEGSA | 0.833 | 0.102 | 0.182 | 0.833 | 0.102 | 0.182 |
| MEMO | 0.588 | 0.208 | 0.308 | 0.769 | 0.208 | 0.327 |
| WExT | 0.650 | 0.265 | 0.377 | 0.812 | 0.265 | 0.400 |

(e) Metrics for LUSC data. (485 samples | 16 CGC-CGC pairs)

| Method | Precision | Sensitivity | F1 Score | $\text{Precision}_{\text{strict}}$ | $\text{Sensitivity}_{\text{strict}}$ | $\text{F1 Score}_{\text{strict}}$ |
|---|---|---|---|---|---|---|
| DISCOVER | NaN | 0.000 | NaN | NaN | 0.000 | NaN |
| Fisher's Exact Test | NaN | 0.000 | NaN | NaN | 0.000 | NaN |
| MEGSA | NaN | 0.000 | NaN | NaN | 0.000 | NaN |
| MEMO | NaN | 0.000 | NaN | NaN | 0.000 | NaN |
| WExT | 1.0 | 0.062 | 0.118 | 1.0 | 0.062 | 0.117 |

(f) Metrics for SKCM data. (468 samples | 373 CGC-CGC pairs)

| Method | Precision | Sensitivity | F1 Score | $\text{Precision}_{\text{strict}}$ | $\text{Sensitivity}_{\text{strict}}$ | $\text{F1 Score}_{\text{strict}}$ |
|---|---|---|---|---|---|---|
| DISCOVER | 0.717 | 0.052 | 0.096 | 0.735 | 0.049 | 0.092 |
| Fisher's Exact Test | 1.000 | 0.005 | 0.011 | 1.000 | 0.005 | 0.010 |
| MEGSA | 0.727 | 0.021 | 0.042 | 0.727 | 0.021 | 0.041 |
| WExT | 0.708 | 0.125 | 0.213 | 0.741 | 0.117 | 0.202 |

(g) Metrics for STAD data. (438 samples | 48 CGC-CGC pairs)

| Method | Precision | Sensitivity | F1 Score | $\text{Precision}_{\text{strict}}$ | $\text{Sensitivity}_{\text{strict}}$ | $\text{F1 Score}_{\text{strict}}$ |
|---|---|---|---|---|---|---|
| DISCOVER | 1.0 | 0.062 | 0.118 | 1.0 | 0.062 | 0.117 |
| Fisher's Exact Test | NaN | 0.000 | NaN | NaN | 0.000 | NaN |
| MEGSA | NaN | 0.000 | NaN | NaN | 0.000 | NaN |
| WExT | 0.8 | 0.083 | 0.151 | 0.8 | 0.083 | 0.150 |

(h) Metrics for UCEC data. (531 samples | 1440 CGC-CGC pairs)

| Method | Precision | Sensitivity | F1 Score | $\text{Precision}_{\text{strict}}$ | $\text{Sensitivity}_{\text{strict}}$ | $\text{F1 Score}_{\text{strict}}$ |
|---|---|---|---|---|---|---|
| DISCOVER | 0.677 | 0.172 | 0.274 | 0.776 | 0.148 | 0.249 |
| Fisher's Exact Test | 0.909 | 0.007 | 0.014 | 1.000 | 0.007 | 0.014 |
| MEGSA | 0.917 | 0.008 | 0.015 | 1.000 | 0.008 | 0.016 |
| WExT | 0.629 | 0.282 | 0.389 | 0.715 | 0.235 | 0.354 |

Table 24: **Degree-normalized network-centric evaluations $X_2$ and $t = 20$**

(a) Metrics for BLCA data.

| Method | Precision | Sensitivity | F1 Score | $\text{Precision}_{\text{strict}}$ | $\text{Sensitivity}_{\text{strict}}$ | $\text{F1 Score}_{\text{strict}}$ |
|---|---|---|---|---|---|---|
| DISCOVER | 1.0 | 1.0 | 0.625 | 1.0 | 0.571 | 0.625 |
| Fisher's Exact Test | 1.0 | 1.0 | 0.25 | 1.0 | 0.143 | 0.25 |
| MEGSA | 1.0 | 1.0 | 0.625 | 1.0 | 0.571 | 0.625 |
| MEMO | 0.75 | 0.929 | 0.611 | 0.75 | 0.571 | 0.611 |
| WExT | 0.667 | 0.857 | 0.6 | 0.667 | 0.571 | 0.6 |

(b) Metrics for BRCA data.

| Method | Precision | Sensitivity | F1 Score | $\text{Precision}_{\text{strict}}$ | $\text{Sensitivity}_{\text{strict}}$ | $\text{F1 Score}_{\text{strict}}$ |
|---|---|---|---|---|---|---|
| DISCOVER | 0.733 | 0.7 | 0.867 | 0.8 | 0.7 | 0.917 |
| DISCOVER Strat | 0.8 | 0.8 | 0.917 | 0.8 | 0.7 | 0.917 |
| Fisher's Exact Test | NaN | NaN | NaN | NaN | NaN | NaN |
| MEGSA | NaN | NaN | NaN | NaN | NaN | NaN |
| MEMO | 0.733 | 0.7 | 0.867 | 0.8 | 0.7 | 0.917 |
| WExT | 0.733 | 0.7 | 0.867 | 0.8 | 0.7 | 0.917 |

(c) Metrics for COADREAD data.

| Method | Precision | Sensitivity | F1 Score | $\text{Precision}_{\text{strict}}$ | $\text{Sensitivity}_{\text{strict}}$ | $\text{F1 Score}_{\text{strict}}$ |
|---|---|---|---|---|---|---|
| DISCOVER | 0.637 | 0.53 | 0.652 | 0.663 | 0.478 | 0.664 |
| DISCOVER Strat | 0.679 | 0.653 | 0.516 | 0.607 | 0.285 | 0.486 |
| Fisher's Exact Test | 0.482 | 0.76 | 0.276 | 0.482 | 0.194 | 0.276 |
| MEGSA | 0.817 | 0.88 | 0.676 | 0.79 | 0.585 | 0.65 |
| MEMO | 0.685 | 0.581 | 0.728 | 0.7 | 0.564 | 0.727 |
| WExT | 0.729 | 0.607 | 0.721 | 0.766 | 0.614 | 0.739 |

(d) Metrics for LUAD data.

| Method | Precision | Sensitivity | F1 Score | $\text{Precision}_{\text{strict}}$ | $\text{Sensitivity}_{\text{strict}}$ | $\text{F1 Score}_{\text{strict}}$ |
|---|---|---|---|---|---|---|
| DISCOVER | 0.643 | 0.702 | 0.68 | 0.643 | 0.429 | 0.66 |
| Fisher's Exact Test | NaN | NaN | NaN | NaN | NaN | NaN |
| MEGSA | 0.375 | 0.625 | 0.325 | 0.375 | 0.104 | 0.325 |
| MEMO | 0.75 | 0.792 | 0.674 | 0.75 | 0.471 | 0.661 |
| WExT | 0.75 | 0.787 | 0.698 | 0.74 | 0.473 | 0.671 |

(e) Metrics for LUSC data.

| Method | Precision | Sensitivity | F1 Score | $\text{Precision}_{\text{strict}}$ | $\text{Sensitivity}_{\text{strict}}$ | $\text{F1 Score}_{\text{strict}}$ |
|---|---|---|---|---|---|---|
| DISCOVER | NaN | NaN | NaN | NaN | NaN | NaN |
| Fisher's Exact Test | NaN | NaN | NaN | NaN | NaN | NaN |
| MEGSA | 1.0 | 1.0 | 0.778 | 1.0 | 0.667 | 0.778 |
| MEMO | 1.0 | 1.0 | 0.833 | 1.0 | 0.75 | 0.833 |
| WExT | 0.667 | 0.5 | 0.7 | 0.75 | 0.75 | 0.75 |

(f) Metrics for SKCM data.

| Method | Precision | Sensitivity | F1 Score | $\text{Precision}_{\text{strict}}$ | $\text{Sensitivity}_{\text{strict}}$ | $\text{F1 Score}_{\text{strict}}$ |
|---|---|---|---|---|---|---|
| DISCOVER | 0.812 | 0.992 | 0.575 | 0.913 | 0.074 | 0.137 |
| Fisher's Exact Test | 1.000 | 1.000 | 0.583 | 1.000 | 0.010 | 0.020 |
| MEGSA | 0.857 | 0.999 | 0.578 | 0.971 | 0.033 | 0.064 |
| WExT | 0.765 | 0.957 | 0.547 | 0.846 | 0.155 | 0.262 |

(g) Metrics for STAD data.

| Method | Precision | Sensitivity | F1 Score | $\text{Precision}_{\text{strict}}$ | $\text{Sensitivity}_{\text{strict}}$ | $\text{F1 Score}_{\text{strict}}$ |
|---|---|---|---|---|---|---|
| DISCOVER | 0.9 | 0.833 | 0.532 | 0.9 | 0.44 | 0.507 |
| Fisher's Exact Test | NaN | NaN | NaN | NaN | NaN | NaN |
| MEGSA | 1.0 | 1.0 | 0.286 | 1.0 | 0.167 | 0.286 |
| WExT | 0.9 | 0.864 | 0.602 | 0.907 | 0.529 | 0.588 |

(h) Metrics for UCEC data.

| Method | Precision | Sensitivity | F1 Score | $\text{Precision}_{\text{strict}}$ | $\text{Sensitivity}_{\text{strict}}$ | $\text{F1 Score}_{\text{strict}}$ |
|---|---|---|---|---|---|---|
| DISCOVER | 0.882 | 0.958 | 0.465 | 0.868 | 0.133 | 0.231 |
| Fisher's Exact Test | 0.952 | 1.000 | 0.212 | 1.000 | 0.004 | 0.008 |
| MEGSA | 0.946 | 0.999 | 0.268 | 0.857 | 0.006 | 0.012 |
| WExT | 0.773 | 0.890 | 0.563 | 0.757 | 0.234 | 0.357 |

Table 25: **Results of network-centric ME evaluation framework with $\mathcal{G}$ = Intact (w conf. threshold 0.35), $\mathcal{S}$ = CGC, c= $X_1$, $p_t$ = 0.05, t=5, robustness_iterations = 100**

(a) Metrics for BLCA data. (411 samples | 1048 CGC-CGC pairs)

| Method | Precision | Sensitivity | F1 Score | $\text{Precision}_{\text{strict}}$ | $\text{Sensitivity}_{\text{strict}}$ | $\text{F1 Score}_{\text{strict}}$ |
|---|---|---|---|---|---|---|
| DISCOVER | 0.923 | 0.012 | 0.023 | 0.923 | 0.012 | 0.024 |
| Fisher's Exact Test | 1.000 | 0.002 | 0.004 | 1.000 | 0.002 | 0.004 |
| WExT | 0.690 | 0.019 | 0.037 | 0.690 | 0.019 | 0.037 |

(b) Metrics for BRCA data. (1026 samples | 958 CGC-CGC pairs)

| Method | Precision | Sensitivity | F1 Score | $\text{Precision}_{\text{strict}}$ | $\text{Sensitivity}_{\text{strict}}$ | $\text{F1 Score}_{\text{strict}}$ |
|---|---|---|---|---|---|---|
| DISCOVER | 0.727 | 0.025 | 0.049 | 0.727 | 0.025 | 0.048 |
| DISCOVER Strat | 0.789 | 0.031 | 0.060 | 0.789 | 0.031 | 0.060 |
| Fisher's Exact Test | 0.667 | 0.002 | 0.004 | 0.667 | 0.002 | 0.004 |
| WExT | 0.691 | 0.059 | 0.109 | 0.707 | 0.056 | 0.104 |

(c) Metrics for COADREAD data. (498 samples | 1748 CGC-CGC pairs)

| Method | Precision | Sensitivity | F1 Score | $\text{Precision}_{\text{strict}}$ | $\text{Sensitivity}_{\text{strict}}$ | $\text{F1 Score}_{\text{strict}}$ |
|---|---|---|---|---|---|---|
| DISCOVER | 0.647 | 0.052 | 0.096 | 0.658 | 0.046 | 0.086 |
| DISCOVER Strat | 0.618 | 0.012 | 0.024 | 0.618 | 0.012 | 0.024 |
| Fisher's Exact Test | 0.583 | 0.008 | 0.016 | 0.565 | 0.007 | 0.014 |
| WExT | 0.645 | 0.121 | 0.203 | 0.668 | 0.102 | 0.177 |

(d) Metrics for LUAD data. (568 samples | 1344 CGC-CGC pairs)

| Method | Precision | Sensitivity | F1 Score | $\text{Precision}_{\text{strict}}$ | $\text{Sensitivity}_{\text{strict}}$ | $\text{F1 Score}_{\text{strict}}$ |
|---|---|---|---|---|---|---|
| DISCOVER | 0.769 | 0.015 | 0.029 | 0.760 | 0.014 | 0.027 |
| Fisher's Exact Test | 0.000 | 0.000 | NaN | 0.000 | 0.000 | NaN |
| WExT | 0.788 | 0.031 | 0.060 | 0.812 | 0.029 | 0.056 |

(e) Metrics for LUSC data. (485 samples | 1080 CGC-CGC pairs)

| Method | Precision | Sensitivity | F1 Score | $\text{Precision}_{\text{strict}}$ | $\text{Sensitivity}_{\text{strict}}$ | $\text{F1 Score}_{\text{strict}}$ |
|---|---|---|---|---|---|---|
| DISCOVER | 1.000 | 0.002 | 0.004 | 1.000 | 0.002 | 0.004 |
| Fisher's Exact Test | 1.000 | 0.002 | 0.004 | 1.000 | 0.002 | 0.004 |
| WExT | 0.857 | 0.006 | 0.011 | 0.857 | 0.006 | 0.012 |

(f) Metrics for SKCM data. (468 samples | 2254 CGC-CGC pairs)

| Method | Precision | Sensitivity | F1 Score | $\text{Precision}_{\text{strict}}$ | $\text{Sensitivity}_{\text{strict}}$ | $\text{F1 Score}_{\text{strict}}$ |
|---|---|---|---|---|---|---|
| DISCOVER | 0.838 | 0.014 | 0.027 | 0.838 | 0.014 | 0.028 |
| Fisher's Exact Test | 1.000 | 0.001 | 0.002 | 1.000 | 0.001 | 0.002 |
| WExT | 0.725 | 0.042 | 0.080 | 0.732 | 0.041 | 0.078 |

(g) Metrics for STAD data. (438 samples | 1460 CGC-CGC pairs)

| Method | Precision | Sensitivity | F1 Score | $\text{Precision}_{\text{strict}}$ | $\text{Sensitivity}_{\text{strict}}$ | $\text{F1 Score}_{\text{strict}}$ |
|---|---|---|---|---|---|---|
| DISCOVER | 0.667 | 0.028 | 0.053 | 0.673 | 0.025 | 0.048 |
| Fisher's Exact Test | 0.667 | 0.003 | 0.005 | 0.667 | 0.003 | 0.006 |
| WExT | 0.688 | 0.069 | 0.125 | 0.705 | 0.060 | 0.111 |

(h) Metrics for UCEC data. (531 samples | 2274 CGC-CGC pairs)

| Method | Precision | Sensitivity | F1 Score | $\text{Precision}_{\text{strict}}$ | $\text{Sensitivity}_{\text{strict}}$ | $\text{F1 Score}_{\text{strict}}$ |
|---|---|---|---|---|---|---|
| DISCOVER | 0.666 | 0.129 | 0.216 | 0.711 | 0.111 | 0.192 |
| Fisher's Exact Test | 0.833 | 0.004 | 0.007 | 0.833 | 0.004 | 0.008 |
| WExT | 0.626 | 0.210 | 0.314 | 0.674 | 0.178 | 0.282 |

Table 26: **Results of network-centric ME evaluation framework with $\mathcal{G}$ = Intact (w conf. threshold 0.35), $\mathcal{S}$ = CGC, c= $X_1$, $p_t$ = 0.05, t=5, robustness_iterations = 300**

(a) Metrics for BLCA data. (411 samples | 1048 CGC-CGC pairs)

| Method | Precision | Sensitivity | F1 Score | $\text{Precision}_{\text{strict}}$ | $\text{Sensitivity}_{\text{strict}}$ | $\text{F1 Score}_{\text{strict}}$ |
|---|---|---|---|---|---|---|
| DISCOVER | 0.923 | 0.011 | 0.023 | 0.923 | 0.011 | 0.022 |
| Fisher's Exact Test | 1.000 | 0.002 | 0.004 | 1.000 | 0.002 | 0.004 |
| WExT | 0.690 | 0.019 | 0.037 | 0.690 | 0.019 | 0.037 |

(b) Metrics for BRCA data. (1026 samples | 958 CGC-CGC pairs)

| Method | Precision | Sensitivity | F1 Score | $\text{Precision}_{\text{strict}}$ | $\text{Sensitivity}_{\text{strict}}$ | $\text{F1 Score}_{\text{strict}}$ |
|---|---|---|---|---|---|---|
| DISCOVER | 0.742 | 0.024 | 0.047 | 0.742 | 0.024 | 0.046 |
| DISCOVER Strat | 0.789 | 0.032 | 0.061 | 0.789 | 0.032 | 0.062 |
| Fisher's Exact Test | 0.667 | 0.002 | 0.004 | 0.667 | 0.002 | 0.004 |
| WExT | 0.705 | 0.058 | 0.107 | 0.726 | 0.056 | 0.104 |

(c) Metrics for COADREAD data. (498 samples | 1748 CGC-CGC pairs)

| Method | Precision | Sensitivity | F1 Score | $\text{Precision}_{\text{strict}}$ | $\text{Sensitivity}_{\text{strict}}$ | $\text{F1 Score}_{\text{strict}}$ |
|---|---|---|---|---|---|---|
| DISCOVER | 0.645 | 0.052 | 0.096 | 0.658 | 0.046 | 0.086 |
| DISCOVER Strat | 0.636 | 0.012 | 0.024 | 0.636 | 0.012 | 0.024 |
| Fisher's Exact Test | 0.583 | 0.008 | 0.016 | 0.565 | 0.007 | 0.014 |
| WExT | 0.644 | 0.120 | 0.202 | 0.669 | 0.102 | 0.177 |

(d) Metrics for LUAD data. (568 samples | 1344 CGC-CGC pairs)

| Method | Precision | Sensitivity | F1 Score | $\text{Precision}_{\text{strict}}$ | $\text{Sensitivity}_{\text{strict}}$ | $\text{F1 Score}_{\text{strict}}$ |
|---|---|---|---|---|---|---|
| DISCOVER | 0.800 | 0.015 | 0.029 | 0.792 | 0.014 | 0.028 |
| Fisher's Exact Test | 0.000 | 0.000 | NaN | 0.000 | 0.000 | NaN |
| WExT | 0.769 | 0.030 | 0.058 | 0.776 | 0.029 | 0.056 |

(e) Metrics for LUSC data. (485 samples | 1080 CGC-CGC pairs)

| Method | Precision | Sensitivity | F1 Score | $\text{Precision}_{\text{strict}}$ | $\text{Sensitivity}_{\text{strict}}$ | $\text{F1 Score}_{\text{strict}}$ |
|---|---|---|---|---|---|---|
| DISCOVER | 1.000 | 0.002 | 0.004 | 1.000 | 0.002 | 0.004 |
| Fisher's Exact Test | 1.000 | 0.002 | 0.004 | 1.000 | 0.002 | 0.004 |
| WExT | 0.857 | 0.006 | 0.011 | 0.857 | 0.006 | 0.012 |

(f) Metrics for SKCM data. (468 samples | 2254 CGC-CGC pairs)

| Method | Precision | Sensitivity | F1 Score | $\text{Precision}_{\text{strict}}$ | $\text{Sensitivity}_{\text{strict}}$ | $\text{F1 Score}_{\text{strict}}$ |
|---|---|---|---|---|---|---|
| DISCOVER | 0.800 | 0.014 | 0.028 | 0.800 | 0.014 | 0.028 |
| Fisher's Exact Test | 1.000 | 0.001 | 0.002 | 1.000 | 0.001 | 0.002 |
| WExT | 0.728 | 0.042 | 0.080 | 0.735 | 0.041 | 0.078 |

(g) Metrics for STAD data. (438 samples | 1460 CGC-CGC pairs)

| Method | Precision | Sensitivity | F1 Score | $\text{Precision}_{\text{strict}}$ | $\text{Sensitivity}_{\text{strict}}$ | $\text{F1 Score}_{\text{strict}}$ |
|---|---|---|---|---|---|---|
| DISCOVER | 0.667 | 0.028 | 0.053 | 0.673 | 0.026 | 0.050 |
| Fisher's Exact Test | 0.667 | 0.003 | 0.005 | 0.667 | 0.003 | 0.006 |
| WExT | 0.688 | 0.069 | 0.125 | 0.705 | 0.060 | 0.111 |

(h) Metrics for UCEC data. (531 samples | 2274 CGC-CGC pairs)

| Method | Precision | Sensitivity | F1 Score | $\text{Precision}_{\text{strict}}$ | $\text{Sensitivity}_{\text{strict}}$ | $\text{F1 Score}_{\text{strict}}$ |
|---|---|---|---|---|---|---|
| DISCOVER | 0.662 | 0.129 | 0.216 | 0.706 | 0.111 | 0.192 |
| Fisher's Exact Test | 0.769 | 0.004 | 0.007 | 0.769 | 0.004 | 0.008 |
| WExT | 0.623 | 0.211 | 0.315 | 0.670 | 0.178 | 0.281 |

Table 27: **Results of network-centric ME evaluation framework with $\mathcal{G}$ = Intact (w conf. threshold 0.35), $\mathcal{S}$ = CGC, c= $X_1$, $p_t$ = 0.05, t=5, robustness_iterations = 500**

(a) Metrics for BLCA data. (411 samples | 1048 CGC-CGC pairs)

| Method | Precision | Sensitivity | F1 Score | $\text{Precision}_{\text{strict}}$ | $\text{Sensitivity}_{\text{strict}}$ | $\text{F1 Score}_{\text{strict}}$ |
|---|---|---|---|---|---|---|
| DISCOVER | 0.923 | 0.012 | 0.023 | 0.923 | 0.012 | 0.024 |
| Fisher's Exact Test | 1.000 | 0.002 | 0.004 | 1.000 | 0.002 | 0.004 |
| WExT | 0.690 | 0.019 | 0.037 | 0.690 | 0.019 | 0.037 |

(b) Metrics for BRCA data. (1026 samples | 958 CGC-CGC pairs)

| Method | Precision | Sensitivity | F1 Score | $\text{Precision}_{\text{strict}}$ | $\text{Sensitivity}_{\text{strict}}$ | $\text{F1 Score}_{\text{strict}}$ |
|---|---|---|---|---|---|---|
| DISCOVER | 0.719 | 0.024 | 0.047 | 0.719 | 0.024 | 0.046 |
| DISCOVER Strat | 0.789 | 0.031 | 0.060 | 0.789 | 0.031 | 0.060 |
| Fisher's Exact Test | 0.667 | 0.002 | 0.004 | 0.667 | 0.002 | 0.004 |
| WExT | 0.705 | 0.058 | 0.107 | 0.726 | 0.056 | 0.104 |

(c) Metrics for COADREAD data. (498 samples | 1748 CGC-CGC pairs)

| Method | Precision | Sensitivity | F1 Score | $\text{Precision}_{\text{strict}}$ | $\text{Sensitivity}_{\text{strict}}$ | $\text{F1 Score}_{\text{strict}}$ |
|---|---|---|---|---|---|---|
| DISCOVER | 0.643 | 0.052 | 0.097 | 0.653 | 0.046 | 0.086 |
| DISCOVER Strat | 0.600 | 0.012 | 0.024 | 0.600 | 0.012 | 0.024 |
| Fisher's Exact Test | 0.591 | 0.007 | 0.015 | 0.591 | 0.007 | 0.014 |
| WExT | 0.652 | 0.120 | 0.202 | 0.677 | 0.102 | 0.177 |

(d) Metrics for LUAD data. (568 samples | 1344 CGC-CGC pairs)

| Method | Precision | Sensitivity | F1 Score | $\text{Precision}_{\text{strict}}$ | $\text{Sensitivity}_{\text{strict}}$ | $\text{F1 Score}_{\text{strict}}$ |
|---|---|---|---|---|---|---|
| DISCOVER | 0.800 | 0.015 | 0.029 | 0.792 | 0.014 | 0.028 |
| Fisher's Exact Test | 0.000 | 0.000 | NaN | 0.000 | 0.000 | NaN |
| WExT | 0.784 | 0.030 | 0.058 | 0.792 | 0.029 | 0.056 |

(e) Metrics for LUSC data. (485 samples | 1080 CGC-CGC pairs)

| Method | Precision | Sensitivity | F1 Score | $\text{Precision}_{\text{strict}}$ | $\text{Sensitivity}_{\text{strict}}$ | $\text{F1 Score}_{\text{strict}}$ |
|---|---|---|---|---|---|---|
| DISCOVER | 1.000 | 0.002 | 0.004 | 1.000 | 0.002 | 0.004 |
| Fisher's Exact Test | 1.000 | 0.002 | 0.004 | 1.000 | 0.002 | 0.004 |
| WExT | 0.857 | 0.006 | 0.011 | 0.857 | 0.006 | 0.012 |

(f) Metrics for SKCM data. (468 samples | 2254 CGC-CGC pairs)

| Method | Precision | Sensitivity | F1 Score | $\text{Precision}_{\text{strict}}$ | $\text{Sensitivity}_{\text{strict}}$ | $\text{F1 Score}_{\text{strict}}$ |
|---|---|---|---|---|---|---|
| DISCOVER | 0.821 | 0.014 | 0.028 | 0.821 | 0.014 | 0.028 |
| Fisher's Exact Test | 1.000 | 0.001 | 0.002 | 1.000 | 0.001 | 0.002 |
| WExT | 0.723 | 0.043 | 0.080 | 0.730 | 0.042 | 0.079 |

(g) Metrics for STAD data. (438 samples | 1460 CGC-CGC pairs)

| Method | Precision | Sensitivity | F1 Score | $\text{Precision}_{\text{strict}}$ | $\text{Sensitivity}_{\text{strict}}$ | $\text{F1 Score}_{\text{strict}}$ |
|---|---|---|---|---|---|---|
| DISCOVER | 0.667 | 0.028 | 0.053 | 0.673 | 0.026 | 0.050 |
| Fisher's Exact Test | 0.667 | 0.003 | 0.005 | 0.667 | 0.003 | 0.006 |
| WExT | 0.692 | 0.069 | 0.125 | 0.711 | 0.060 | 0.111 |

(h) Metrics for UCEC data. (531 samples | 2274 CGC-CGC pairs)

| Method | Precision | Sensitivity | F1 Score | $\text{Precision}_{\text{strict}}$ | $\text{Sensitivity}_{\text{strict}}$ | $\text{F1 Score}_{\text{strict}}$ |
|---|---|---|---|---|---|---|
| DISCOVER | 0.661 | 0.128 | 0.215 | 0.704 | 0.110 | 0.190 |
| Fisher's Exact Test | 0.769 | 0.004 | 0.007 | 0.769 | 0.004 | 0.008 |
| WExT | 0.628 | 0.213 | 0.318 | 0.675 | 0.180 | 0.284 |

Table 28: **Results of network-centric ME evaluation framework with $\mathcal{G}$ = Intact (w conf. threshold 0.35), $\mathcal{S}$ = CGC, c= $X_1$, $p_t$ = 0.01, t=5, robustness_iterations = 100**

(a) Metrics for BLCA data. (411 samples | 1048 CGC-CGC pairs)

| Method | Precision | Sensitivity | F1 Score | $\text{Precision}_{\text{strict}}$ | $\text{Sensitivity}_{\text{strict}}$ | $\text{F1 Score}_{\text{strict}}$ |
|---|---|---|---|---|---|---|
| DISCOVER | 1.0 | 0.002 | 0.004 | 1.0 | 0.002 | 0.004 |
| Fisher's Exact Test | NaN | 0.000 | NaN | NaN | 0.000 | NaN |
| WExT | 0.8 | 0.004 | 0.008 | 0.8 | 0.004 | 0.008 |

(b) Metrics for BRCA data. (1026 samples | 958 CGC-CGC pairs)

| Method | Precision | Sensitivity | F1 Score | $\text{Precision}_{\text{strict}}$ | $\text{Sensitivity}_{\text{strict}}$ | $\text{F1 Score}_{\text{strict}}$ |
|---|---|---|---|---|---|---|
| DISCOVER | 0.667 | 0.006 | 0.012 | 0.667 | 0.006 | 0.012 |
| DISCOVER Strat | 0.750 | 0.006 | 0.012 | 0.750 | 0.006 | 0.012 |
| Fisher's Exact Test | NaN | 0.000 | NaN | NaN | 0.000 | NaN |
| WExT | 0.556 | 0.011 | 0.021 | 0.556 | 0.011 | 0.022 |

(c) Metrics for COADREAD data. (498 samples | 1748 CGC-CGC pairs)

| Method | Precision | Sensitivity | F1 Score | $\text{Precision}_{\text{strict}}$ | $\text{Sensitivity}_{\text{strict}}$ | $\text{F1 Score}_{\text{strict}}$ |
|---|---|---|---|---|---|---|
| DISCOVER | 0.673 | 0.020 | 0.038 | 0.674 | 0.018 | 0.035 |
| DISCOVER Strat | 0.000 | 0.000 | NaN | 0.000 | 0.000 | NaN |
| Fisher's Exact Test | 0.400 | 0.001 | 0.002 | 0.400 | 0.001 | 0.002 |
| WExT | 0.678 | 0.069 | 0.125 | 0.703 | 0.055 | 0.102 |

(d) Metrics for LUAD data. (568 samples | 1344 CGC-CGC pairs)

| Method | Precision | Sensitivity | F1 Score | $\text{Precision}_{\text{strict}}$ | $\text{Sensitivity}_{\text{strict}}$ | $\text{F1 Score}_{\text{strict}}$ |
|---|---|---|---|---|---|---|
| DISCOVER | 0.667 | 0.003 | 0.006 | 0.667 | 0.003 | 0.006 |
| Fisher's Exact Test | NaN | 0.000 | NaN | NaN | 0.000 | NaN |
| WExT | 0.829 | 0.013 | 0.025 | 0.829 | 0.013 | 0.026 |

(e) Metrics for LUSC data. (485 samples | 1080 CGC-CGC pairs)

| Method | Precision | Sensitivity | F1 Score | $\text{Precision}_{\text{strict}}$ | $\text{Sensitivity}_{\text{strict}}$ | $\text{F1 Score}_{\text{strict}}$ |
|---|---|---|---|---|---|---|
| DISCOVER | 1.0 | 0.002 | 0.004 | 1.0 | 0.002 | 0.004 |
| Fisher's Exact Test | NaN | 0.000 | NaN | NaN | 0.000 | NaN |
| WExT | 1.0 | 0.002 | 0.004 | 1.0 | 0.002 | 0.004 |

(f) Metrics for SKCM data. (468 samples | 2254 CGC-CGC pairs)

| Method | Precision | Sensitivity | F1 Score | $\text{Precision}_{\text{strict}}$ | $\text{Sensitivity}_{\text{strict}}$ | $\text{F1 Score}_{\text{strict}}$ |
|---|---|---|---|---|---|---|
| DISCOVER | 0.750 | 0.003 | 0.005 | 0.750 | 0.003 | 0.006 |
| Fisher's Exact Test | 1.000 | 0.001 | 0.002 | 1.000 | 0.001 | 0.002 |
| WExT | 0.792 | 0.008 | 0.017 | 0.792 | 0.008 | 0.016 |

(g) Metrics for STAD data. (438 samples | 1460 CGC-CGC pairs)

| Method | Precision | Sensitivity | F1 Score | $\text{Precision}_{\text{strict}}$ | $\text{Sensitivity}_{\text{strict}}$ | $\text{F1 Score}_{\text{strict}}$ |
|---|---|---|---|---|---|---|
| DISCOVER | 0.615 | 0.008 | 0.016 | 0.595 | 0.008 | 0.016 |
| Fisher's Exact Test | 0.800 | 0.001 | 0.003 | 0.800 | 0.001 | 0.002 |
| WExT | 0.631 | 0.028 | 0.054 | 0.654 | 0.023 | 0.044 |

(h) Metrics for UCEC data. (531 samples | 2274 CGC-CGC pairs)

| Method | Precision | Sensitivity | F1 Score | $\text{Precision}_{\text{strict}}$ | $\text{Sensitivity}_{\text{strict}}$ | $\text{F1 Score}_{\text{strict}}$ |
|---|---|---|---|---|---|---|
| DISCOVER | 0.676 | 0.072 | 0.129 | 0.721 | 0.063 | 0.116 |
| Fisher's Exact Test | 0.857 | 0.002 | 0.004 | 0.857 | 0.002 | 0.004 |
| WExT | 0.650 | 0.136 | 0.224 | 0.710 | 0.116 | 0.199 |

Table 29: **Results of network-centric ME evaluation framework with $\mathcal{G}$ = Intact (w conf. threshold 0.35), $\mathcal{S}$ = CGC, c= $X_1$, $p_t = 0.1$, t=5, robustness_iterations = 100**

(a) Metrics for BLCA data. (411 samples | 1048 CGC-CGC pairs)

| Method | Precision | Sensitivity | F1 Score | $\text{Precision}_{\text{strict}}$ | $\text{Sensitivity}_{\text{strict}}$ | $\text{F1 Score}_{\text{strict}}$ |
|---|---|---|---|---|---|---|
| DISCOVER | 0.769 | 0.019 | 0.038 | 0.769 | 0.019 | 0.037 |
| Fisher's Exact Test | 1.000 | 0.002 | 0.004 | 1.000 | 0.002 | 0.004 |
| WExT | 0.746 | 0.047 | 0.089 | 0.742 | 0.046 | 0.087 |

(b) Metrics for BRCA data. (1026 samples | 958 CGC-CGC pairs)

| Method | Precision | Sensitivity | F1 Score | $\text{Precision}_{\text{strict}}$ | $\text{Sensitivity}_{\text{strict}}$ | $\text{F1 Score}_{\text{strict}}$ |
|---|---|---|---|---|---|---|
| DISCOVER | 0.696 | 0.051 | 0.094 | 0.697 | 0.049 | 0.092 |
| DISCOVER Strat | 0.728 | 0.056 | 0.105 | 0.741 | 0.054 | 0.101 |
| Fisher's Exact Test | 0.667 | 0.002 | 0.004 | 0.667 | 0.002 | 0.004 |
| WExT | 0.670 | 0.109 | 0.187 | 0.695 | 0.099 | 0.173 |

(c) Metrics for COADREAD data. (498 samples | 1748 CGC-CGC pairs)

| Method | Precision | Sensitivity | F1 Score | $\text{Precision}_{\text{strict}}$ | $\text{Sensitivity}_{\text{strict}}$ | $\text{F1 Score}_{\text{strict}}$ |
|---|---|---|---|---|---|---|
| DISCOVER | 0.647 | 0.089 | 0.156 | 0.662 | 0.079 | 0.141 |
| DISCOVER Strat | 0.663 | 0.039 | 0.074 | 0.660 | 0.038 | 0.072 |
| Fisher's Exact Test | 0.656 | 0.017 | 0.034 | 0.646 | 0.015 | 0.029 |
| WExT | 0.648 | 0.191 | 0.295 | 0.672 | 0.166 | 0.266 |

(d) Metrics for LUAD data. (568 samples | 1344 CGC-CGC pairs)

| Method | Precision | Sensitivity | F1 Score | $\text{Precision}_{\text{strict}}$ | $\text{Sensitivity}_{\text{strict}}$ | $\text{F1 Score}_{\text{strict}}$ |
|---|---|---|---|---|---|---|
| DISCOVER | 0.764 | 0.031 | 0.060 | 0.784 | 0.030 | 0.058 |
| Fisher's Exact Test | 0.750 | 0.004 | 0.009 | 0.750 | 0.004 | 0.008 |
| WExT | 0.737 | 0.057 | 0.106 | 0.757 | 0.052 | 0.097 |

(e) Metrics for LUSC data. (485 samples | 1080 CGC-CGC pairs)

| Method | Precision | Sensitivity | F1 Score | $\text{Precision}_{\text{strict}}$ | $\text{Sensitivity}_{\text{strict}}$ | $\text{F1 Score}_{\text{strict}}$ |
|---|---|---|---|---|---|---|
| DISCOVER | 1.0 | 0.004 | 0.007 | 1.0 | 0.004 | 0.008 |
| Fisher's Exact Test | 1.0 | 0.002 | 0.004 | 1.0 | 0.002 | 0.004 |
| WExT | 0.7 | 0.013 | 0.026 | 0.7 | 0.013 | 0.026 |

(f) Metrics for SKCM data. (468 samples | 2254 CGC-CGC pairs)

| Method | Precision | Sensitivity | F1 Score | $\text{Precision}_{\text{strict}}$ | $\text{Sensitivity}_{\text{strict}}$ | $\text{F1 Score}_{\text{strict}}$ |
|---|---|---|---|---|---|---|
| DISCOVER | 0.763 | 0.039 | 0.075 | 0.768 | 0.039 | 0.074 |
| Fisher's Exact Test | 1.000 | 0.002 | 0.004 | 1.000 | 0.002 | 0.004 |
| WExT | 0.662 | 0.080 | 0.144 | 0.666 | 0.078 | 0.140 |

(g) Metrics for STAD data. (438 samples | 1460 CGC-CGC pairs)

| Method | Precision | Sensitivity | F1 Score | $\text{Precision}_{\text{strict}}$ | $\text{Sensitivity}_{\text{strict}}$ | $\text{F1 Score}_{\text{strict}}$ |
|---|---|---|---|---|---|---|
| DISCOVER | 0.698 | 0.052 | 0.097 | 0.715 | 0.048 | 0.090 |
| Fisher's Exact Test | 0.667 | 0.005 | 0.011 | 0.667 | 0.005 | 0.010 |
| WExT | 0.639 | 0.093 | 0.163 | 0.652 | 0.081 | 0.144 |

(h) Metrics for UCEC data. (531 samples | 2274 CGC-CGC pairs)

| Method | Precision | Sensitivity | F1 Score | $\text{Precision}_{\text{strict}}$ | $\text{Sensitivity}_{\text{strict}}$ | $\text{F1 Score}_{\text{strict}}$ |
|---|---|---|---|---|---|---|
| DISCOVER | 0.640 | 0.174 | 0.274 | 0.684 | 0.148 | 0.243 |
| Fisher's Exact Test | 0.704 | 0.007 | 0.014 | 0.720 | 0.006 | 0.012 |
| WExT | 0.599 | 0.273 | 0.375 | 0.640 | 0.230 | 0.338 |

Table 30: **Results of network-centric ME evaluation framework with $\mathcal{G}$ = Intact (w conf. threshold 0.35), $\mathcal{S}$ = $CGC_{SNV}$ , c= $X_1$, $p_t$ = 0.05, t=5, robustness_iterations = 100**

(a) Metrics for BLCA data. (411 samples | 363 $CGC_{SNV}$-$CGC_{SNV}$ pairs)

| Method | Precision | Sensitivity | F1 Score | $\text{Precision}_{\text{strict}}$ | $\text{Sensitivity}_{\text{strict}}$ | $\text{F1 Score}_{\text{strict}}$ |
|---|---|---|---|---|---|---|
| DISCOVER | 0.889 | 0.013 | 0.025 | 0.889 | 0.013 | 0.026 |
| Fisher's Exact Test | 1.000 | 0.003 | 0.006 | 1.000 | 0.003 | 0.006 |
| WExT | 0.667 | 0.022 | 0.043 | 0.667 | 0.022 | 0.043 |

(b) Metrics for BRCA data. (1026 samples | 568 $CGC_{SNV}$-$CGC_{SNV}$ pairs)

| Method | Precision | Sensitivity | F1 Score | $\text{Precision}_{\text{strict}}$ | $\text{Sensitivity}_{\text{strict}}$ | $\text{F1 Score}_{\text{strict}}$ |
|---|---|---|---|---|---|---|
| DISCOVER | 0.653 | 0.029 | 0.055 | 0.653 | 0.029 | 0.056 |
| DISCOVER Strat | 0.759 | 0.039 | 0.074 | 0.759 | 0.039 | 0.074 |
| Fisher's Exact Test | 0.667 | 0.004 | 0.007 | 0.667 | 0.004 | 0.008 |
| WExT | 0.672 | 0.080 | 0.144 | 0.694 | 0.077 | 0.139 |

(c) Metrics for COADREAD data. (498 samples | 636 $CGC_{SNV}$-$CGC_{SNV}$ pairs)

| Method | Precision | Sensitivity | F1 Score | $\text{Precision}_{\text{strict}}$ | $\text{Sensitivity}_{\text{strict}}$ | $\text{F1 Score}_{\text{strict}}$ |
|---|---|---|---|---|---|---|
| DISCOVER | 0.621 | 0.065 | 0.117 | 0.637 | 0.057 | 0.105 |
| DISCOVER Strat | 0.704 | 0.020 | 0.039 | 0.704 | 0.020 | 0.039 |
| Fisher's Exact Test | 0.632 | 0.012 | 0.025 | 0.611 | 0.011 | 0.022 |
| WExT | 0.608 | 0.147 | 0.237 | 0.622 | 0.123 | 0.205 |

(d) Metrics for LUAD data. (568 samples | 694 $CGC_{SNV}$-$CGC_{SNV}$ pairs)

| Method | Precision | Sensitivity | F1 Score | $\text{Precision}_{\text{strict}}$ | $\text{Sensitivity}_{\text{strict}}$ | $\text{F1 Score}_{\text{strict}}$ |
|---|---|---|---|---|---|---|
| DISCOVER | 0.737 | 0.020 | 0.040 | 0.722 | 0.019 | 0.037 |
| Fisher's Exact Test | 0.000 | 0.000 | NaN | 0.000 | 0.000 | NaN |
| WExT | 0.780 | 0.047 | 0.089 | 0.789 | 0.044 | 0.083 |

(e) Metrics for LUSC data. (485 samples | 646 $CGC_{SNV}$-$CGC_{SNV}$ pairs)

| Method | Precision | Sensitivity | F1 Score | $\text{Precision}_{\text{strict}}$ | $\text{Sensitivity}_{\text{strict}}$ | $\text{F1 Score}_{\text{strict}}$ |
|---|---|---|---|---|---|---|
| DISCOVER | 1.000 | 0.003 | 0.006 | 1.000 | 0.003 | 0.006 |
| Fisher's Exact Test | 1.000 | 0.003 | 0.006 | 1.000 | 0.003 | 0.006 |
| WExT | 0.857 | 0.009 | 0.019 | 0.857 | 0.009 | 0.018 |

(f) Metrics for SKCM data. (468 samples | 1144 $CGC_{SNV}$-$CGC_{SNV}$ pairs)

| Method | Precision | Sensitivity | F1 Score | $\text{Precision}_{\text{strict}}$ | $\text{Sensitivity}_{\text{strict}}$ | $\text{F1 Score}_{\text{strict}}$ |
|---|---|---|---|---|---|---|
| DISCOVER | 0.640 | 0.014 | 0.028 | 0.640 | 0.014 | 0.027 |
| Fisher's Exact Test | 1.000 | 0.002 | 0.003 | 1.000 | 0.002 | 0.004 |
| WExT | 0.677 | 0.057 | 0.105 | 0.682 | 0.054 | 0.100 |

(g) Metrics for STAD data. (438 samples | 800 $CGC_{SNV}$-$CGC_{SNV}$ pairs)

| Method | Precision | Sensitivity | F1 Score | $\text{Precision}_{\text{strict}}$ | $\text{Sensitivity}_{\text{strict}}$ | $\text{F1 Score}_{\text{strict}}$ |
|---|---|---|---|---|---|---|
| DISCOVER | 0.615 | 0.030 | 0.058 | 0.629 | 0.028 | 0.054 |
| Fisher's Exact Test | 0.667 | 0.003 | 0.005 | 0.667 | 0.003 | 0.006 |
| WExT | 0.667 | 0.080 | 0.143 | 0.692 | 0.069 | 0.125 |

(h) Metrics for UCEC data. (531 samples | 1294 $CGC_{SNV}$-$CGC_{SNV}$ pairs)

| Method | Precision | Sensitivity | F1 Score | $\text{Precision}_{\text{strict}}$ | $\text{Sensitivity}_{\text{strict}}$ | $\text{F1 Score}_{\text{strict}}$ |
|---|---|---|---|---|---|---|
| DISCOVER | 0.657 | 0.180 | 0.283 | 0.696 | 0.152 | 0.250 |
| Fisher's Exact Test | 0.833 | 0.008 | 0.015 | 0.833 | 0.008 | 0.016 |
| WExT | 0.627 | 0.291 | 0.397 | 0.675 | 0.242 | 0.356 |

Table 31: **Results of network-centric ME evaluation framework with $\mathcal{G}$ = Intact (w conf. threshold 0.35), $\mathcal{S}$ = Intogen , c= $X_1$, $p_t$ = 0.05, t=5, robustness_iterations = 100**

(a) Metrics for BLCA data. (411 samples | 968 Intogen-Intogen pairs)

| Method | Precision | Sensitivity | F1 Score | $\text{Precision}_{\text{strict}}$ | $\text{Sensitivity}_{\text{strict}}$ | $\text{F1 Score}_{\text{strict}}$ |
|---|---|---|---|---|---|---|
| DISCOVER | 0.941 | 0.017 | 0.033 | 0.941 | 0.017 | 0.033 |
| Fisher's Exact Test | 1.000 | 0.002 | 0.004 | 1.000 | 0.002 | 0.004 |
| WExT | 0.733 | 0.023 | 0.045 | 0.733 | 0.023 | 0.045 |

(b) Metrics for BRCA data. (1026 samples | 908 Intogen-Intogen pairs)

| Method | Precision | Sensitivity | F1 Score | $\text{Precision}_{\text{strict}}$ | $\text{Sensitivity}_{\text{strict}}$ | $\text{F1 Score}_{\text{strict}}$ |
|---|---|---|---|---|---|---|
| DISCOVER | 0.711 | 0.030 | 0.057 | 0.711 | 0.030 | 0.058 |
| DISCOVER Strat | 0.780 | 0.035 | 0.068 | 0.780 | 0.035 | 0.067 |
| Fisher's Exact Test | 0.667 | 0.002 | 0.004 | 0.667 | 0.002 | 0.004 |
| WExT | 0.693 | 0.068 | 0.124 | 0.704 | 0.064 | 0.117 |

(c) Metrics for COADREAD data. (498 samples | 1618 Intogen-Intogen pairs)

| Method | Precision | Sensitivity | F1 Score | $\text{Precision}_{\text{strict}}$ | $\text{Sensitivity}_{\text{strict}}$ | $\text{F1 Score}_{\text{strict}}$ |
|---|---|---|---|---|---|---|
| DISCOVER | 0.655 | 0.068 | 0.124 | 0.669 | 0.060 | 0.110 |
| DISCOVER Strat | 0.680 | 0.021 | 0.041 | 0.673 | 0.021 | 0.041 |
| Fisher's Exact Test | 0.558 | 0.009 | 0.018 | 0.551 | 0.008 | 0.016 |
| WExT | 0.644 | 0.145 | 0.237 | 0.661 | 0.122 | 0.206 |

(d) Metrics for LUAD data. (568 samples | 1164 Intogen-Intogen pairs)

| Method | Precision | Sensitivity | F1 Score | $\text{Precision}_{\text{strict}}$ | $\text{Sensitivity}_{\text{strict}}$ | $\text{F1 Score}_{\text{strict}}$ |
|---|---|---|---|---|---|---|
| DISCOVER | 0.720 | 0.016 | 0.030 | 0.708 | 0.015 | 0.029 |
| Fisher's Exact Test | 0.000 | 0.000 | NaN | 0.000 | 0.000 | NaN |
| WExT | 0.754 | 0.038 | 0.071 | 0.755 | 0.035 | 0.067 |

(e) Metrics for LUSC data. (485 samples | 996 Intogen-Intogen pairs)

| Method | Precision | Sensitivity | F1 Score | $\text{Precision}_{\text{strict}}$ | $\text{Sensitivity}_{\text{strict}}$ | $\text{F1 Score}_{\text{strict}}$ |
|---|---|---|---|---|---|---|
| DISCOVER | 1.000 | 0.004 | 0.008 | 1.000 | 0.004 | 0.008 |
| Fisher's Exact Test | 1.000 | 0.002 | 0.004 | 1.000 | 0.002 | 0.004 |
| WExT | 0.909 | 0.010 | 0.020 | 0.909 | 0.010 | 0.020 |

(f) Metrics for SKCM data. (468 samples | 1844 Intogen-Intogen pairs)

| Method | Precision | Sensitivity | F1 Score | $\text{Precision}_{\text{strict}}$ | $\text{Sensitivity}_{\text{strict}}$ | $\text{F1 Score}_{\text{strict}}$ |
|---|---|---|---|---|---|---|
| DISCOVER | 0.691 | 0.015 | 0.030 | 0.691 | 0.015 | 0.029 |
| Fisher's Exact Test | 1.000 | 0.001 | 0.002 | 1.000 | 0.001 | 0.002 |
| WExT | 0.702 | 0.048 | 0.090 | 0.708 | 0.047 | 0.088 |

(g) Metrics for STAD data. (438 samples | 1398 Intogen-Intogen pairs)

| Method | Precision | Sensitivity | F1 Score | $\text{Precision}_{\text{strict}}$ | $\text{Sensitivity}_{\text{strict}}$ | $\text{F1 Score}_{\text{strict}}$ |
|---|---|---|---|---|---|---|
| DISCOVER | 0.648 | 0.033 | 0.063 | 0.656 | 0.030 | 0.057 |
| Fisher's Exact Test | 0.667 | 0.004 | 0.009 | 0.667 | 0.004 | 0.008 |
| WExT | 0.693 | 0.084 | 0.150 | 0.712 | 0.073 | 0.132 |

(h) Metrics for UCEC data. (531 samples | 2082 Intogen-Intogen pairs)

| Method | Precision | Sensitivity | F1 Score | $\text{Precision}_{\text{strict}}$ | $\text{Sensitivity}_{\text{strict}}$ | $\text{F1 Score}_{\text{strict}}$ |
|---|---|---|---|---|---|---|
| DISCOVER | 0.643 | 0.170 | 0.268 | 0.692 | 0.142 | 0.236 |
| Fisher's Exact Test | 0.765 | 0.006 | 0.012 | 0.765 | 0.006 | 0.012 |
| WExT | 0.602 | 0.258 | 0.361 | 0.645 | 0.212 | 0.319 |

Table 32: **Results of network-centric ME evaluation framework with $\mathcal{G}$ = Intact (w conf. threshold 0.25), $\mathcal{S}$ = CGC, c= $X_1$, $p_t$ = 0.05, t=5, robustness_iterations = 100**

(a) Metrics for BLCA data. (411 samples | 1530 CGC-CGC pairs)

| Method | Precision | Sensitivity | F1 Score | $\text{Precision}_{\text{strict}}$ | $\text{Sensitivity}_{\text{strict}}$ | $\text{F1 Score}_{\text{strict}}$ |
|---|---|---|---|---|---|---|
| DISCOVER | 0.947 | 0.012 | 0.023 | 0.947 | 0.012 | 0.024 |
| Fisher's Exact Test | 1.000 | 0.004 | 0.008 | 1.000 | 0.004 | 0.008 |
| WExT | 0.732 | 0.020 | 0.039 | 0.732 | 0.020 | 0.039 |

(b) Metrics for BRCA data. (1026 samples | 1376 CGC-CGC pairs)

| Method | Precision | Sensitivity | F1 Score | $\text{Precision}_{\text{strict}}$ | $\text{Sensitivity}_{\text{strict}}$ | $\text{F1 Score}_{\text{strict}}$ |
|---|---|---|---|---|---|---|
| DISCOVER | 0.720 | 0.026 | 0.051 | 0.720 | 0.026 | 0.050 |
| DISCOVER Strat | 0.820 | 0.030 | 0.058 | 0.820 | 0.030 | 0.058 |
| Fisher's Exact Test | 0.800 | 0.003 | 0.006 | 0.800 | 0.003 | 0.006 |
| WExT | 0.707 | 0.058 | 0.106 | 0.727 | 0.055 | 0.102 |

(c) Metrics for COADREAD data. (498 samples | 2588 CGC-CGC pairs)

| Method | Precision | Sensitivity | F1 Score | $\text{Precision}_{\text{strict}}$ | $\text{Sensitivity}_{\text{strict}}$ | $\text{F1 Score}_{\text{strict}}$ |
|---|---|---|---|---|---|---|
| DISCOVER | 0.646 | 0.044 | 0.083 | 0.669 | 0.039 | 0.074 |
| DISCOVER Strat | 0.645 | 0.012 | 0.023 | 0.659 | 0.012 | 0.024 |
| Fisher's Exact Test | 0.611 | 0.009 | 0.017 | 0.618 | 0.008 | 0.016 |
| WExT | 0.652 | 0.107 | 0.183 | 0.676 | 0.092 | 0.162 |

(d) Metrics for LUAD data. (568 samples | 1948 CGC-CGC pairs)

| Method | Precision | Sensitivity | F1 Score | $\text{Precision}_{\text{strict}}$ | $\text{Sensitivity}_{\text{strict}}$ | $\text{F1 Score}_{\text{strict}}$ |
|---|---|---|---|---|---|---|
| DISCOVER | 0.775 | 0.016 | 0.031 | 0.789 | 0.015 | 0.029 |
| Fisher's Exact Test | 0.667 | 0.002 | 0.004 | 0.667 | 0.002 | 0.004 |
| WExT | 0.767 | 0.029 | 0.056 | 0.783 | 0.028 | 0.054 |

(e) Metrics for LUSC data. (485 samples | 1536 CGC-CGC pairs)

| Method | Precision | Sensitivity | F1 Score | $\text{Precision}_{\text{strict}}$ | $\text{Sensitivity}_{\text{strict}}$ | $\text{F1 Score}_{\text{strict}}$ |
|---|---|---|---|---|---|---|
| DISCOVER | 1.000 | 0.001 | 0.003 | 1.000 | 0.001 | 0.002 |
| Fisher's Exact Test | 1.000 | 0.001 | 0.003 | 1.000 | 0.001 | 0.002 |
| WExT | 0.727 | 0.005 | 0.010 | 0.727 | 0.005 | 0.010 |

(f) Metrics for SKCM data. (468 samples | 3208 CGC-CGC pairs)

| Method | Precision | Sensitivity | F1 Score | $\text{Precision}_{\text{strict}}$ | $\text{Sensitivity}_{\text{strict}}$ | $\text{F1 Score}_{\text{strict}}$ |
|---|---|---|---|---|---|---|
| DISCOVER | 0.774 | 0.013 | 0.025 | 0.774 | 0.013 | 0.026 |
| Fisher's Exact Test | 1.000 | 0.001 | 0.001 | 1.000 | 0.001 | 0.002 |
| WExT | 0.704 | 0.036 | 0.068 | 0.709 | 0.035 | 0.067 |

(g) Metrics for STAD data. (438 samples | 2286 CGC-CGC pairs)

| Method | Precision | Sensitivity | F1 Score | $\text{Precision}_{\text{strict}}$ | $\text{Sensitivity}_{\text{strict}}$ | $\text{F1 Score}_{\text{strict}}$ |
|---|---|---|---|---|---|---|
| DISCOVER | 0.685 | 0.028 | 0.053 | 0.707 | 0.026 | 0.050 |
| Fisher's Exact Test | 0.727 | 0.004 | 0.007 | 0.727 | 0.004 | 0.008 |
| WExT | 0.687 | 0.061 | 0.111 | 0.722 | 0.054 | 0.100 |

(h) Metrics for UCEC data. (531 samples | 4120 CGC-CGC pairs)

| Method | Precision | Sensitivity | F1 Score | $\text{Precision}_{\text{strict}}$ | $\text{Sensitivity}_{\text{strict}}$ | $\text{F1 Score}_{\text{strict}}$ |
|---|---|---|---|---|---|---|
| DISCOVER | 0.652 | 0.116 | 0.196 | 0.695 | 0.100 | 0.175 |
| Fisher's Exact Test | 0.737 | 0.003 | 0.007 | 0.737 | 0.003 | 0.006 |
| WExT | 0.618 | 0.202 | 0.305 | 0.654 | 0.172 | 0.272 |

Table 33: **Results of network-centric ME evaluation framework with $\mathcal{G}$ = Intact (w conf. threshold 0.45), $\mathcal{S}$ = CGC, c= $X_1$, $p_t$ = 0.05, t=5, robustness_iterations = 100**

(a) Metrics for BLCA data. (411 samples | 468 CGC-CGC pairs)

| Method | Precision | Sensitivity | F1 Score | $\text{Precision}_{\text{strict}}$ | $\text{Sensitivity}_{\text{strict}}$ | $\text{F1 Score}_{\text{strict}}$ |
|---|---|---|---|---|---|---|
| DISCOVER | 1.000 | 0.013 | 0.026 | 1.000 | 0.013 | 0.026 |
| Fisher's Exact Test | 1.000 | 0.004 | 0.009 | 1.000 | 0.004 | 0.008 |
| WExT | 0.727 | 0.017 | 0.034 | 0.727 | 0.017 | 0.033 |

(b) Metrics for BRCA data. (1026 samples | 420 CGC-CGC pairs)

| Method | Precision | Sensitivity | F1 Score | $\text{Precision}_{\text{strict}}$ | $\text{Sensitivity}_{\text{strict}}$ | $\text{F1 Score}_{\text{strict}}$ |
|---|---|---|---|---|---|---|
| DISCOVER | 0.667 | 0.024 | 0.046 | 0.667 | 0.024 | 0.046 |
| DISCOVER Strat | 0.824 | 0.034 | 0.065 | 0.824 | 0.034 | 0.065 |
| Fisher's Exact Test | 1.000 | 0.005 | 0.010 | 1.000 | 0.005 | 0.010 |
| WExT | 0.750 | 0.066 | 0.121 | 0.743 | 0.063 | 0.116 |

(c) Metrics for COADREAD data. (498 samples | 784 CGC-CGC pairs)

| Method | Precision | Sensitivity | F1 Score | $\text{Precision}_{\text{strict}}$ | $\text{Sensitivity}_{\text{strict}}$ | $\text{F1 Score}_{\text{strict}}$ |
|---|---|---|---|---|---|---|
| DISCOVER | 0.687 | 0.059 | 0.109 | 0.702 | 0.052 | 0.097 |
| DISCOVER Strat | 0.706 | 0.015 | 0.030 | 0.706 | 0.015 | 0.029 |
| Fisher's Exact Test | 0.583 | 0.009 | 0.018 | 0.583 | 0.009 | 0.018 |
| WExT | 0.668 | 0.133 | 0.221 | 0.705 | 0.110 | 0.190 |

(d) Metrics for LUAD data. (568 samples | 636 CGC-CGC pairs)

| Method | Precision | Sensitivity | F1 Score | $\text{Precision}_{\text{strict}}$ | $\text{Sensitivity}_{\text{strict}}$ | $\text{F1 Score}_{\text{strict}}$ |
|---|---|---|---|---|---|---|
| DISCOVER | 0.800 | 0.013 | 0.025 | 0.80 | 0.013 | 0.026 |
| Fisher's Exact Test | NaN | 0.000 | NaN | NaN | 0.000 | NaN |
| WExT | 0.769 | 0.032 | 0.061 | 0.76 | 0.030 | 0.058 |

(e) Metrics for LUSC data. (485 samples | 478 CGC-CGC pairs)

| Method | Precision | Sensitivity | F1 Score | $\text{Precision}_{\text{strict}}$ | $\text{Sensitivity}_{\text{strict}}$ | $\text{F1 Score}_{\text{strict}}$ |
|---|---|---|---|---|---|---|
| DISCOVER | 1.000 | 0.004 | 0.008 | 1.000 | 0.004 | 0.008 |
| Fisher's Exact Test | 1.000 | 0.004 | 0.008 | 1.000 | 0.004 | 0.008 |
| WExT | 0.857 | 0.013 | 0.025 | 0.857 | 0.013 | 0.026 |

(f) Metrics for SKCM data. (468 samples | 1036 CGC-CGC pairs)

| Method | Precision | Sensitivity | F1 Score | $\text{Precision}_{\text{strict}}$ | $\text{Sensitivity}_{\text{strict}}$ | $\text{F1 Score}_{\text{strict}}$ |
|---|---|---|---|---|---|---|
| DISCOVER | 0.842 | 0.016 | 0.031 | 0.842 | 0.016 | 0.031 |
| Fisher's Exact Test | 1.000 | 0.002 | 0.004 | 1.000 | 0.002 | 0.004 |
| WExT | 0.807 | 0.048 | 0.090 | 0.832 | 0.047 | 0.089 |

(g) Metrics for STAD data. (438 samples | 714 CGC-CGC pairs)

| Method | Precision | Sensitivity | F1 Score | $\text{Precision}_{\text{strict}}$ | $\text{Sensitivity}_{\text{strict}}$ | $\text{F1 Score}_{\text{strict}}$ |
|---|---|---|---|---|---|---|
| DISCOVER | 0.684 | 0.037 | 0.069 | 0.706 | 0.034 | 0.065 |
| Fisher's Exact Test | 0.800 | 0.006 | 0.011 | 0.800 | 0.006 | 0.012 |
| WExT | 0.667 | 0.077 | 0.138 | 0.692 | 0.064 | 0.117 |

(h) Metrics for UCEC data. (531 samples | 1256 CGC-CGC pairs)

| Method | Precision | Sensitivity | F1 Score | $\text{Precision}_{\text{strict}}$ | $\text{Sensitivity}_{\text{strict}}$ | $\text{F1 Score}_{\text{strict}}$ |
|---|---|---|---|---|---|---|
| DISCOVER | 0.673 | 0.154 | 0.250 | 0.721 | 0.130 | 0.220 |
| Fisher's Exact Test | 0.800 | 0.006 | 0.013 | 0.800 | 0.006 | 0.012 |
| WExT | 0.637 | 0.231 | 0.339 | 0.683 | 0.190 | 0.297 |

Table 34: **Results of network-centric ME evaluation framework with $\mathcal{G}$ = HINT, $\mathcal{S}$ = CGC, c= $X_1$, $p_t$ = 0.05, t=5, robustness_iterations = 100**

(a) Metrics for BLCA data. (411 samples | 800 CGC-CGC pairs)

| Method | Precision | Sensitivity | F1 Score | $\text{Precision}_{\text{strict}}$ | $\text{Sensitivity}_{\text{strict}}$ | $\text{F1 Score}_{\text{strict}}$ |
|---|---|---|---|---|---|---|
| DISCOVER | 0.903 | 0.018 | 0.035 | 0.903 | 0.018 | 0.035 |
| Fisher's Exact Test | 1.000 | 0.005 | 0.010 | 1.000 | 0.005 | 0.010 |
| WExT | 0.750 | 0.030 | 0.058 | 0.750 | 0.030 | 0.058 |

(b) Metrics for BRCA data. (1026 samples | 736 CGC-CGC pairs)

| Method | Precision | Sensitivity | F1 Score | $\text{Precision}_{\text{strict}}$ | $\text{Sensitivity}_{\text{strict}}$ | $\text{F1 Score}_{\text{strict}}$ |
|---|---|---|---|---|---|---|
| DISCOVER | 0.741 | 0.027 | 0.053 | 0.741 | 0.027 | 0.052 |
| DISCOVER Strat | 0.800 | 0.033 | 0.063 | 0.800 | 0.033 | 0.063 |
| Fisher's Exact Test | 1.000 | 0.003 | 0.005 | 1.000 | 0.003 | 0.006 |
| WExT | 0.770 | 0.075 | 0.136 | 0.785 | 0.071 | 0.130 |

(c) Metrics for COADREAD data. (498 samples | 1278 CGC-CGC pairs)

| Method | Precision | Sensitivity | F1 Score | $\text{Precision}_{\text{strict}}$ | $\text{Sensitivity}_{\text{strict}}$ | $\text{F1 Score}_{\text{strict}}$ |
|---|---|---|---|---|---|---|
| DISCOVER | 0.695 | 0.071 | 0.128 | 0.736 | 0.064 | 0.118 |
| DISCOVER Strat | 0.771 | 0.029 | 0.056 | 0.783 | 0.028 | 0.054 |
| Fisher's Exact Test | 0.738 | 0.018 | 0.034 | 0.754 | 0.017 | 0.033 |
| WExT | 0.657 | 0.137 | 0.226 | 0.707 | 0.118 | 0.202 |

(d) Metrics for LUAD data. (568 samples | 1018 CGC-CGC pairs)

| Method | Precision | Sensitivity | F1 Score | $\text{Precision}_{\text{strict}}$ | $\text{Sensitivity}_{\text{strict}}$ | $\text{F1 Score}_{\text{strict}}$ |
|---|---|---|---|---|---|---|
| DISCOVER | 0.769 | 0.010 | 0.019 | 0.769 | 0.010 | 0.020 |
| Fisher's Exact Test | 0.667 | 0.002 | 0.004 | 0.667 | 0.002 | 0.004 |
| WExT | 0.722 | 0.026 | 0.050 | 0.714 | 0.025 | 0.048 |

(e) Metrics for LUSC data. (485 samples | 794 CGC-CGC pairs)

| Method | Precision | Sensitivity | F1 Score | $\text{Precision}_{\text{strict}}$ | $\text{Sensitivity}_{\text{strict}}$ | $\text{F1 Score}_{\text{strict}}$ |
|---|---|---|---|---|---|---|
| DISCOVER | 1.000 | 0.003 | 0.005 | 1.000 | 0.003 | 0.006 |
| Fisher's Exact Test | 1.000 | 0.003 | 0.005 | 1.000 | 0.003 | 0.006 |
| WExT | 0.667 | 0.005 | 0.010 | 0.667 | 0.005 | 0.010 |

(f) Metrics for SKCM data. (468 samples | 1560 CGC-CGC pairs)

| Method | Precision | Sensitivity | F1 Score | $\text{Precision}_{\text{strict}}$ | $\text{Sensitivity}_{\text{strict}}$ | $\text{F1 Score}_{\text{strict}}$ |
|---|---|---|---|---|---|---|
| DISCOVER | 0.800 | 0.010 | 0.020 | 0.800 | 0.010 | 0.020 |
| Fisher's Exact Test | NaN | 0.000 | NaN | NaN | 0.000 | NaN |
| WExT | 0.721 | 0.039 | 0.074 | 0.718 | 0.038 | 0.072 |

(g) Metrics for STAD data. (438 samples | 1100 CGC-CGC pairs)

| Method | Precision | Sensitivity | F1 Score | $\text{Precision}_{\text{strict}}$ | $\text{Sensitivity}_{\text{strict}}$ | $\text{F1 Score}_{\text{strict}}$ |
|---|---|---|---|---|---|---|
| DISCOVER | 0.630 | 0.029 | 0.055 | 0.656 | 0.027 | 0.052 |
| Fisher's Exact Test | 0.500 | 0.002 | 0.004 | 0.500 | 0.002 | 0.004 |
| WExT | 0.713 | 0.081 | 0.145 | 0.750 | 0.070 | 0.128 |

(h) Metrics for UCEC data. (531 samples | 1884 CGC-CGC pairs)

| Method | Precision | Sensitivity | F1 Score | $\text{Precision}_{\text{strict}}$ | $\text{Sensitivity}_{\text{strict}}$ | $\text{F1 Score}_{\text{strict}}$ |
|---|---|---|---|---|---|---|
| DISCOVER | 0.681 | 0.153 | 0.249 | 0.741 | 0.132 | 0.224 |
| Fisher's Exact Test | 0.800 | 0.006 | 0.013 | 0.800 | 0.006 | 0.012 |
| WExT | 0.657 | 0.252 | 0.365 | 0.717 | 0.217 | 0.333 |

Table 35: **Results of network-centric ME evaluation framework with $\mathcal{G}$ = STRING, $\mathcal{S}$ = CGC, c= $X_1$, $p_t$ = 0.05, t=5, robustness_iterations = 100**

(a) Metrics for BLCA data. (411 samples | 1368 CGC-CGC pairs)

| Method | Precision | Sensitivity | F1 Score | $\text{Precision}_{\text{strict}}$ | $\text{Sensitivity}_{\text{strict}}$ | $\text{F1 Score}_{\text{strict}}$ |
|---|---|---|---|---|---|---|
| DISCOVER | 0.933 | 0.010 | 0.020 | 0.933 | 0.010 | 0.020 |
| Fisher's Exact Test | 1.000 | 0.003 | 0.006 | 1.000 | 0.003 | 0.006 |
| WExT | 0.706 | 0.018 | 0.035 | 0.706 | 0.018 | 0.035 |

(b) Metrics for BRCA data. (1026 samples | 1110 CGC-CGC pairs)

| Method | Precision | Sensitivity | F1 Score | $\text{Precision}_{\text{strict}}$ | $\text{Sensitivity}_{\text{strict}}$ | $\text{F1 Score}_{\text{strict}}$ |
|---|---|---|---|---|---|---|
| DISCOVER | 0.824 | 0.038 | 0.073 | 0.824 | 0.038 | 0.073 |
| DISCOVER Strat | 0.900 | 0.041 | 0.078 | 0.900 | 0.041 | 0.078 |
| Fisher's Exact Test | 1.000 | 0.002 | 0.004 | 1.000 | 0.002 | 0.004 |
| WExT | 0.775 | 0.077 | 0.141 | 0.788 | 0.073 | 0.134 |

(c) Metrics for COADREAD data. (498 samples | 1994 CGC-CGC pairs)

| Method | Precision | Sensitivity | F1 Score | $\text{Precision}_{\text{strict}}$ | $\text{Sensitivity}_{\text{strict}}$ | $\text{F1 Score}_{\text{strict}}$ |
|---|---|---|---|---|---|---|
| DISCOVER | 0.701 | 0.048 | 0.090 | 0.746 | 0.045 | 0.085 |
| DISCOVER Strat | 0.862 | 0.017 | 0.034 | 0.873 | 0.017 | 0.033 |
| Fisher's Exact Test | 0.850 | 0.009 | 0.017 | 0.850 | 0.009 | 0.018 |
| WExT | 0.664 | 0.096 | 0.168 | 0.711 | 0.085 | 0.152 |

(d) Metrics for LUAD data. (568 samples | 1464 CGC-CGC pairs)

| Method | Precision | Sensitivity | F1 Score | $\text{Precision}_{\text{strict}}$ | $\text{Sensitivity}_{\text{strict}}$ | $\text{F1 Score}_{\text{strict}}$ |
|---|---|---|---|---|---|---|
| DISCOVER | 0.800 | 0.014 | 0.027 | 0.833 | 0.014 | 0.028 |
| Fisher's Exact Test | 0.909 | 0.007 | 0.014 | 0.909 | 0.007 | 0.014 |
| WExT | 0.745 | 0.024 | 0.047 | 0.786 | 0.023 | 0.045 |

(e) Metrics for LUSC data. (485 samples | 1188 CGC-CGC pairs)

| Method | Precision | Sensitivity | F1 Score | $\text{Precision}_{\text{strict}}$ | $\text{Sensitivity}_{\text{strict}}$ | $\text{F1 Score}_{\text{strict}}$ |
|---|---|---|---|---|---|---|
| DISCOVER | 0.800 | 0.003 | 0.007 | 0.800 | 0.003 | 0.006 |
| Fisher's Exact Test | 1.000 | 0.002 | 0.003 | 1.000 | 0.002 | 0.004 |
| WExT | 0.857 | 0.005 | 0.010 | 0.857 | 0.005 | 0.010 |

(f) Metrics for SKCM data. (468 samples | 2400 CGC-CGC pairs)

| Method | Precision | Sensitivity | F1 Score | $\text{Precision}_{\text{strict}}$ | $\text{Sensitivity}_{\text{strict}}$ | $\text{F1 Score}_{\text{strict}}$ |
|---|---|---|---|---|---|---|
| DISCOVER | 0.830 | 0.016 | 0.032 | 0.844 | 0.016 | 0.031 |
| Fisher's Exact Test | 1.000 | 0.002 | 0.003 | 1.000 | 0.002 | 0.004 |
| WExT | 0.714 | 0.037 | 0.070 | 0.740 | 0.036 | 0.069 |

(g) Metrics for STAD data. (438 samples | 1628 CGC-CGC pairs)

| Method | Precision | Sensitivity | F1 Score | $\text{Precision}_{\text{strict}}$ | $\text{Sensitivity}_{\text{strict}}$ | $\text{F1 Score}_{\text{strict}}$ |
|---|---|---|---|---|---|---|
| DISCOVER | 0.718 | 0.017 | 0.034 | 0.750 | 0.017 | 0.033 |
| Fisher's Exact Test | 0.667 | 0.001 | 0.002 | 0.667 | 0.001 | 0.002 |
| WExT | 0.714 | 0.039 | 0.074 | 0.765 | 0.036 | 0.069 |

(h) Metrics for UCEC data. (531 samples | 3044 CGC-CGC pairs)

| Method | Precision | Sensitivity | F1 Score | $\text{Precision}_{\text{strict}}$ | $\text{Sensitivity}_{\text{strict}}$ | $\text{F1 Score}_{\text{strict}}$ |
|---|---|---|---|---|---|---|
| DISCOVER | 0.679 | 0.110 | 0.190 | 0.734 | 0.096 | 0.170 |
| Fisher's Exact Test | 0.923 | 0.004 | 0.008 | 0.923 | 0.004 | 0.008 |
| WExT | 0.620 | 0.194 | 0.295 | 0.666 | 0.164 | 0.263 |

Table 36: **Degree-normalized network-centric evaluations $X_1$ and $t = 5$**

(a) Metrics for BLCA data.

| Method | Precision | Sensitivity | F1 Score | $\text{Precision}_{\text{strict}}$ | $\text{Sensitivity}_{\text{strict}}$ | $\text{F1 Score}_{\text{strict}}$ |
|---|---|---|---|---|---|---|
| DISCOVER | 0.958 | 0.268 | 0.367 | 0.958 | 0.268 | 0.367 |
| Fisher's Exact Test | 1.0 | 0.072 | 0.133 | 1.0 | 0.072 | 0.133 |
| WExT | 0.772 | 0.275 | 0.404 | 0.772 | 0.275 | 0.404 |

(b) Metrics for BRCA data.

| Method | Precision | Sensitivity | F1 Score | $\text{Precision}_{\text{strict}}$ | $\text{Sensitivity}_{\text{strict}}$ | $\text{F1 Score}_{\text{strict}}$ |
|---|---|---|---|---|---|---|
| DISCOVER | 0.866 | 0.407 | 0.496 | 0.866 | 0.407 | 0.496 |
| DISCOVER Strat | 0.927 | 0.391 | 0.478 | 0.927 | 0.391 | 0.478 |
| Fisher's Exact Test | 0.667 | 0.125 | 0.29 | 0.667 | 0.125 | 0.29 |
| WExT | 0.88 | 0.347 | 0.457 | 0.882 | 0.343 | 0.454 |

(c) Metrics for COADREAD data.

| Method | Precision | Sensitivity | F1 Score | $\text{Precision}_{\text{strict}}$ | $\text{Sensitivity}_{\text{strict}}$ | $\text{F1 Score}_{\text{strict}}$ |
|---|---|---|---|---|---|---|
| DISCOVER | 0.857 | 0.303 | 0.394 | 0.852 | 0.286 | 0.383 |
| DISCOVER Strat | 0.773 | 0.111 | 0.203 | 0.773 | 0.111 | 0.203 |
| Fisher's Exact Test | 0.881 | 0.199 | 0.294 | 0.875 | 0.196 | 0.289 |
| WExT | 0.824 | 0.343 | 0.443 | 0.823 | 0.318 | 0.432 |

(d) Metrics for LUAD data.

| Method | Precision | Sensitivity | F1 Score | $\text{Precision}_{\text{strict}}$ | $\text{Sensitivity}_{\text{strict}}$ | $\text{F1 Score}_{\text{strict}}$ |
|---|---|---|---|---|---|---|
| DISCOVER | 0.938 | 0.342 | 0.445 | 0.936 | 0.34 | 0.442 |
| Fisher's Exact Test | 0.0 | 0.0 | NaN | 0.0 | 0.0 | NaN |
| WExT | 0.936 | 0.297 | 0.393 | 0.935 | 0.295 | 0.391 |

(e) Metrics for LUSC data.

| Method | Precision | Sensitivity | F1 Score | $\text{Precision}_{\text{strict}}$ | $\text{Sensitivity}_{\text{strict}}$ | $\text{F1 Score}_{\text{strict}}$ |
|---|---|---|---|---|---|---|
| DISCOVER | 1.0 | 0.611 | 0.682 | 1.0 | 0.611 | 0.682 |
| Fisher's Exact Test | 1.0 | 0.6 | 0.667 | 1.0 | 0.6 | 0.667 |
| WExT | 0.917 | 0.334 | 0.436 | 0.917 | 0.334 | 0.436 |

(f) Metrics for SKCM data.

| Method | Precision | Sensitivity | F1 Score | $\text{Precision}_{\text{strict}}$ | $\text{Sensitivity}_{\text{strict}}$ | $\text{F1 Score}_{\text{strict}}$ |
|---|---|---|---|---|---|---|
| DISCOVER | 0.813 | 0.015 | 0.303 | 0.875 | 0.014 | 0.028 |
| Fisher's Exact Test | 1.000 | 0.001 | 0.267 | 1.000 | 0.001 | 0.002 |
| WExT | 0.801 | 0.035 | 0.307 | 0.872 | 0.035 | 0.067 |

(g) Metrics for STAD data.

| Method | Precision | Sensitivity | F1 Score | $\text{Precision}_{\text{strict}}$ | $\text{Sensitivity}_{\text{strict}}$ | $\text{F1 Score}_{\text{strict}}$ |
|---|---|---|---|---|---|---|
| DISCOVER | 0.891 | 0.019 | 0.316 | 0.857 | 0.018 | 0.035 |
| Fisher's Exact Test | 0.833 | 0.002 | 0.331 | 1.000 | 0.002 | 0.004 |
| WExT | 0.891 | 0.056 | 0.401 | 0.881 | 0.053 | 0.100 |

(h) Metrics for UCEC data.

| Method | Precision | Sensitivity | F1 Score | $\text{Precision}_{\text{strict}}$ | $\text{Sensitivity}_{\text{strict}}$ | $\text{F1 Score}_{\text{strict}}$ |
|---|---|---|---|---|---|---|
| DISCOVER | 0.875 | 0.098 | 0.396 | 0.824 | 0.092 | 0.166 |
| Fisher's Exact Test | 0.952 | 0.001 | 0.152 | 1.000 | 0.001 | 0.002 |
| WExT | 0.809 | 0.184 | 0.476 | 0.809 | 0.167 | 0.277 |

Table 37: **Results of network-centric ME evaluation framework with $\mathcal{G}$ = Intact (w conf. threshold 0.35), $\mathcal{S}$ = CGC, c= $X_2$, $p_t$ = 0.05, t=5, robustness_iterations = 100**

(a) Metrics for BLCA data. (411 samples | 895 CGC-CGC pairs)

| Method | Precision | Sensitivity | F1 Score | $\text{Precision}_{\text{strict}}$ | $\text{Sensitivity}_{\text{strict}}$ | $\text{F1 Score}_{\text{strict}}$ |
|---|---|---|---|---|---|---|
| DISCOVER | 1.0 | 0.013 | 0.026 | 1.0 | 0.013 | 0.026 |
| Fisher's Exact Test | 1.0 | 0.002 | 0.004 | 1.0 | 0.002 | 0.004 |
| WExT | 0.8 | 0.022 | 0.043 | 0.8 | 0.022 | 0.043 |

(b) Metrics for BRCA data. (1026 samples | 737 CGC-CGC pairs)

| Method | Precision | Sensitivity | F1 Score | $\text{Precision}_{\text{strict}}$ | $\text{Sensitivity}_{\text{strict}}$ | $\text{F1 Score}_{\text{strict}}$ |
|---|---|---|---|---|---|---|
| DISCOVER | 0.773 | 0.023 | 0.045 | 0.773 | 0.023 | 0.045 |
| DISCOVER Strat | 0.852 | 0.031 | 0.060 | 0.852 | 0.031 | 0.060 |
| Fisher's Exact Test | 1.000 | 0.003 | 0.005 | 1.000 | 0.003 | 0.006 |
| WExT | 0.698 | 0.060 | 0.110 | 0.724 | 0.057 | 0.106 |

(c) Metrics for COADREAD data. (498 samples | 1625 CGC-CGC pairs)

| Method | Precision | Sensitivity | F1 Score | $\text{Precision}_{\text{strict}}$ | $\text{Sensitivity}_{\text{strict}}$ | $\text{F1 Score}_{\text{strict}}$ |
|---|---|---|---|---|---|---|
| DISCOVER | 0.721 | 0.052 | 0.097 | 0.746 | 0.048 | 0.090 |
| DISCOVER Strat | 0.641 | 0.013 | 0.025 | 0.641 | 0.013 | 0.025 |
| Fisher's Exact Test | 0.619 | 0.008 | 0.016 | 0.619 | 0.008 | 0.016 |
| WExT | 0.670 | 0.118 | 0.200 | 0.712 | 0.103 | 0.180 |

(d) Metrics for LUAD data. (568 samples | 1197 CGC-CGC pairs)

| Method | Precision | Sensitivity | F1 Score | $\text{Precision}_{\text{strict}}$ | $\text{Sensitivity}_{\text{strict}}$ | $\text{F1 Score}_{\text{strict}}$ |
|---|---|---|---|---|---|---|
| DISCOVER | 0.889 | 0.013 | 0.026 | 0.889 | 0.013 | 0.026 |
| Fisher's Exact Test | NaN | 0.000 | NaN | NaN | 0.000 | NaN |
| WExT | 0.804 | 0.031 | 0.060 | 0.818 | 0.030 | 0.058 |

(e) Metrics for LUSC data. (485 samples | 906 CGC-CGC pairs)

| Method | Precision | Sensitivity | F1 Score | $\text{Precision}_{\text{strict}}$ | $\text{Sensitivity}_{\text{strict}}$ | $\text{F1 Score}_{\text{strict}}$ |
|---|---|---|---|---|---|---|
| DISCOVER | 1.000 | 0.002 | 0.004 | 1.000 | 0.002 | 0.004 |
| Fisher's Exact Test | 0.667 | 0.002 | 0.004 | 0.667 | 0.002 | 0.004 |
| WExT | 0.600 | 0.007 | 0.013 | 0.600 | 0.007 | 0.014 |

(f) Metrics for SKCM data. (468 samples | 2142 CGC-CGC pairs)

| Method | Precision | Sensitivity | F1 Score | $\text{Precision}_{\text{strict}}$ | $\text{Sensitivity}_{\text{strict}}$ | $\text{F1 Score}_{\text{strict}}$ |
|---|---|---|---|---|---|---|
| DISCOVER | 0.784 | 0.014 | 0.027 | 0.806 | 0.014 | 0.028 |
| Fisher's Exact Test | 1.000 | 0.001 | 0.002 | 1.000 | 0.001 | 0.002 |
| WExT | 0.743 | 0.042 | 0.080 | 0.753 | 0.042 | 0.080 |

(g) Metrics for STAD data. (438 samples | 1308 CGC-CGC pairs)

| Method | Precision | Sensitivity | F1 Score | $\text{Precision}_{\text{strict}}$ | $\text{Sensitivity}_{\text{strict}}$ | $\text{F1 Score}_{\text{strict}}$ |
|---|---|---|---|---|---|---|
| DISCOVER | 0.743 | 0.029 | 0.055 | 0.763 | 0.027 | 0.052 |
| Fisher's Exact Test | 0.800 | 0.003 | 0.006 | 0.800 | 0.003 | 0.006 |
| WExT | 0.718 | 0.072 | 0.132 | 0.757 | 0.065 | 0.120 |

(h) Metrics for UCEC data. (531 samples | 2668 CGC-CGC pairs)

| Method | Precision | Sensitivity | F1 Score | $\text{Precision}_{\text{strict}}$ | $\text{Sensitivity}_{\text{strict}}$ | $\text{F1 Score}_{\text{strict}}$ |
|---|---|---|---|---|---|---|
| DISCOVER | 0.703 | 0.130 | 0.219 | 0.777 | 0.117 | 0.203 |
| Fisher's Exact Test | 0.833 | 0.004 | 0.007 | 0.833 | 0.004 | 0.008 |
| WExT | 0.651 | 0.213 | 0.321 | 0.728 | 0.189 | 0.300 |

Table 38: **Results of network-centric ME evaluation framework with $\mathcal{G}$ = Intact (w conf. threshold 0.35), $\mathcal{S}$ = CGC, c= $X_2$, $p_t$ = 0.05, t=5, robustness_iterations = 300**

(a) Metrics for BLCA data. (411 samples | 895 CGC-CGC pairs)

| Method | Precision | Sensitivity | F1 Score | $\text{Precision}_{\text{strict}}$ | $\text{Sensitivity}_{\text{strict}}$ | $\text{F1 Score}_{\text{strict}}$ |
|---|---|---|---|---|---|---|
| DISCOVER | 1.0 | 0.013 | 0.026 | 1.0 | 0.013 | 0.026 |
| Fisher's Exact Test | 1.0 | 0.002 | 0.004 | 1.0 | 0.002 | 0.004 |
| WExT | 0.8 | 0.022 | 0.043 | 0.8 | 0.022 | 0.043 |

(b) Metrics for BRCA data. (1026 samples | 737 CGC-CGC pairs)

| Method | Precision | Sensitivity | F1 Score | $\text{Precision}_{\text{strict}}$ | $\text{Sensitivity}_{\text{strict}}$ | $\text{F1 Score}_{\text{strict}}$ |
|---|---|---|---|---|---|---|
| DISCOVER | 0.773 | 0.023 | 0.045 | 0.773 | 0.023 | 0.045 |
| DISCOVER Strat | 0.852 | 0.031 | 0.060 | 0.852 | 0.031 | 0.060 |
| Fisher's Exact Test | 1.000 | 0.003 | 0.005 | 1.000 | 0.003 | 0.006 |
| WExT | 0.698 | 0.060 | 0.110 | 0.724 | 0.057 | 0.106 |

(c) Metrics for COADREAD data. (498 samples | 1625 CGC-CGC pairs)

| Method | Precision | Sensitivity | F1 Score | $\text{Precision}_{\text{strict}}$ | $\text{Sensitivity}_{\text{strict}}$ | $\text{F1 Score}_{\text{strict}}$ |
|---|---|---|---|---|---|---|
| DISCOVER | 0.714 | 0.053 | 0.098 | 0.736 | 0.048 | 0.090 |
| DISCOVER Strat | 0.645 | 0.012 | 0.024 | 0.645 | 0.012 | 0.024 |
| Fisher's Exact Test | 0.619 | 0.008 | 0.016 | 0.619 | 0.008 | 0.016 |
| WExT | 0.676 | 0.116 | 0.198 | 0.720 | 0.101 | 0.177 |

(d) Metrics for LUAD data. (568 samples | 1197 CGC-CGC pairs)

| Method | Precision | Sensitivity | F1 Score | $\text{Precision}_{\text{strict}}$ | $\text{Sensitivity}_{\text{strict}}$ | $\text{F1 Score}_{\text{strict}}$ |
|---|---|---|---|---|---|---|
| DISCOVER | 0.842 | 0.013 | 0.026 | 0.842 | 0.013 | 0.026 |
| Fisher's Exact Test | NaN | 0.000 | NaN | NaN | 0.000 | NaN |
| WExT | 0.800 | 0.030 | 0.058 | 0.814 | 0.029 | 0.056 |

(e) Metrics for LUSC data. (485 samples | 906 CGC-CGC pairs)

| Method | Precision | Sensitivity | F1 Score | $\text{Precision}_{\text{strict}}$ | $\text{Sensitivity}_{\text{strict}}$ | $\text{F1 Score}_{\text{strict}}$ |
|---|---|---|---|---|---|---|
| DISCOVER | 1.0 | 0.002 | 0.004 | 1.0 | 0.002 | 0.004 |
| Fisher's Exact Test | 1.0 | 0.002 | 0.004 | 1.0 | 0.002 | 0.004 |
| WExT | 0.6 | 0.007 | 0.013 | 0.6 | 0.007 | 0.014 |

(f) Metrics for SKCM data. (468 samples | 2142 CGC-CGC pairs)

| Method | Precision | Sensitivity | F1 Score | $\text{Precision}_{\text{strict}}$ | $\text{Sensitivity}_{\text{strict}}$ | $\text{F1 Score}_{\text{strict}}$ |
|---|---|---|---|---|---|---|
| DISCOVER | 0.800 | 0.013 | 0.026 | 0.800 | 0.013 | 0.026 |
| Fisher's Exact Test | 1.000 | 0.001 | 0.002 | 1.000 | 0.001 | 0.002 |
| WExT | 0.754 | 0.042 | 0.079 | 0.765 | 0.041 | 0.078 |

(g) Metrics for STAD data. (438 samples | 1308 CGC-CGC pairs)

| Method | Precision | Sensitivity | F1 Score | $\text{Precision}_{\text{strict}}$ | $\text{Sensitivity}_{\text{strict}}$ | $\text{F1 Score}_{\text{strict}}$ |
|---|---|---|---|---|---|---|
| DISCOVER | 0.738 | 0.029 | 0.056 | 0.758 | 0.028 | 0.054 |
| Fisher's Exact Test | 0.800 | 0.003 | 0.006 | 0.800 | 0.003 | 0.006 |
| WExT | 0.725 | 0.074 | 0.134 | 0.771 | 0.066 | 0.122 |

(h) Metrics for UCEC data. (531 samples | 2668 CGC-CGC pairs)

| Method | Precision | Sensitivity | F1 Score | $\text{Precision}_{\text{strict}}$ | $\text{Sensitivity}_{\text{strict}}$ | $\text{F1 Score}_{\text{strict}}$ |
|---|---|---|---|---|---|---|
| DISCOVER | 0.700 | 0.131 | 0.220 | 0.772 | 0.117 | 0.203 |
| Fisher's Exact Test | 0.833 | 0.004 | 0.007 | 0.833 | 0.004 | 0.008 |
| WExT | 0.649 | 0.214 | 0.322 | 0.725 | 0.188 | 0.299 |

Table 39: **Results of network-centric ME evaluation framework with $\mathcal{G}$ = Intact (w conf. threshold 0.35), $\mathcal{S}$ = CGC, c= $X_2$, $p_t$ = 0.05, t=5, robustness_iterations = 500**

(a) Metrics for BLCA data. (411 samples | 895 CGC-CGC pairs)

| Method | Precision | Sensitivity | F1 Score | $\text{Precision}_{\text{strict}}$ | $\text{Sensitivity}_{\text{strict}}$ | $\text{F1 Score}_{\text{strict}}$ |
|---|---|---|---|---|---|---|
| DISCOVER | 0.923 | 0.012 | 0.023 | 0.923 | 0.012 | 0.024 |
| Fisher's Exact Test | 1.000 | 0.002 | 0.004 | 1.000 | 0.002 | 0.004 |
| WExT | 0.690 | 0.019 | 0.037 | 0.690 | 0.019 | 0.037 |

(b) Metrics for BRCA data. (1026 samples | 737 CGC-CGC pairs)

| Method | Precision | Sensitivity | F1 Score | $\text{Precision}_{\text{strict}}$ | $\text{Sensitivity}_{\text{strict}}$ | $\text{F1 Score}_{\text{strict}}$ |
|---|---|---|---|---|---|---|
| DISCOVER | 0.719 | 0.024 | 0.047 | 0.719 | 0.024 | 0.046 |
| DISCOVER Strat | 0.789 | 0.031 | 0.060 | 0.789 | 0.031 | 0.060 |
| Fisher's Exact Test | 0.667 | 0.002 | 0.004 | 0.667 | 0.002 | 0.004 |
| WExT | 0.705 | 0.058 | 0.107 | 0.726 | 0.056 | 0.104 |

(c) Metrics for COADREAD data. (498 samples | 1625 CGC-CGC pairs)

| Method | Precision | Sensitivity | F1 Score | $\text{Precision}_{\text{strict}}$ | $\text{Sensitivity}_{\text{strict}}$ | $\text{F1 Score}_{\text{strict}}$ |
|---|---|---|---|---|---|---|
| DISCOVER | 0.643 | 0.052 | 0.097 | 0.653 | 0.046 | 0.086 |
| DISCOVER Strat | 0.600 | 0.012 | 0.024 | 0.600 | 0.012 | 0.024 |
| Fisher's Exact Test | 0.591 | 0.007 | 0.015 | 0.591 | 0.007 | 0.014 |
| WExT | 0.652 | 0.120 | 0.202 | 0.677 | 0.102 | 0.177 |

(d) Metrics for LUAD data. (568 samples | 1197 CGC-CGC pairs)

| Method | Precision | Sensitivity | F1 Score | $\text{Precision}_{\text{strict}}$ | $\text{Sensitivity}_{\text{strict}}$ | $\text{F1 Score}_{\text{strict}}$ |
|---|---|---|---|---|---|---|
| DISCOVER | 0.800 | 0.015 | 0.029 | 0.792 | 0.014 | 0.028 |
| Fisher's Exact Test | 0.000 | 0.000 | NaN | 0.000 | 0.000 | NaN |
| WExT | 0.784 | 0.030 | 0.058 | 0.792 | 0.029 | 0.056 |

(e) Metrics for LUSC data. (485 samples | 906 CGC-CGC pairs)

| Method | Precision | Sensitivity | F1 Score | $\text{Precision}_{\text{strict}}$ | $\text{Sensitivity}_{\text{strict}}$ | $\text{F1 Score}_{\text{strict}}$ |
|---|---|---|---|---|---|---|
| DISCOVER | 1.000 | 0.002 | 0.004 | 1.000 | 0.002 | 0.004 |
| Fisher's Exact Test | 1.000 | 0.002 | 0.004 | 1.000 | 0.002 | 0.004 |
| WExT | 0.857 | 0.006 | 0.011 | 0.857 | 0.006 | 0.012 |

(f) Metrics for SKCM data. (468 samples | 2142 CGC-CGC pairs)

| Method | Precision | Sensitivity | F1 Score | $\text{Precision}_{\text{strict}}$ | $\text{Sensitivity}_{\text{strict}}$ | $\text{F1 Score}_{\text{strict}}$ |
|---|---|---|---|---|---|---|
| DISCOVER | 0.821 | 0.014 | 0.028 | 0.821 | 0.014 | 0.028 |
| Fisher's Exact Test | 1.000 | 0.001 | 0.002 | 1.000 | 0.001 | 0.002 |
| WExT | 0.723 | 0.043 | 0.080 | 0.730 | 0.042 | 0.079 |

(g) Metrics for STAD data. (438 samples | 1308 CGC-CGC pairs)

| Method | Precision | Sensitivity | F1 Score | $\text{Precision}_{\text{strict}}$ | $\text{Sensitivity}_{\text{strict}}$ | $\text{F1 Score}_{\text{strict}}$ |
|---|---|---|---|---|---|---|
| DISCOVER | 0.667 | 0.028 | 0.053 | 0.673 | 0.026 | 0.050 |
| Fisher's Exact Test | 0.667 | 0.003 | 0.005 | 0.667 | 0.003 | 0.006 |
| WExT | 0.692 | 0.069 | 0.125 | 0.711 | 0.060 | 0.111 |

(h) Metrics for UCEC data. (531 samples | 2668 CGC-CGC pairs)

| Method | Precision | Sensitivity | F1 Score | $\text{Precision}_{\text{strict}}$ | $\text{Sensitivity}_{\text{strict}}$ | $\text{F1 Score}_{\text{strict}}$ |
|---|---|---|---|---|---|---|
| DISCOVER | 0.661 | 0.128 | 0.215 | 0.704 | 0.110 | 0.190 |
| Fisher's Exact Test | 0.769 | 0.004 | 0.007 | 0.769 | 0.004 | 0.008 |
| WExT | 0.628 | 0.213 | 0.318 | 0.675 | 0.180 | 0.284 |

Table 40: **Results of network-centric ME evaluation framework with $\mathcal{G}$ = Intact (w conf. threshold 0.35), $\mathcal{S}$ = CGC, c= $X_2$, $p_t$ = 0.01, t=5, robustness_iterations = 100**

(a) Metrics for BLCA data. (411 samples | 895 CGC-CGC pairs)

| Method | Precision | Sensitivity | F1 Score | $\text{Precision}_{\text{strict}}$ | $\text{Sensitivity}_{\text{strict}}$ | $\text{F1 Score}_{\text{strict}}$ |
|---|---|---|---|---|---|---|
| DISCOVER | 1.0 | 0.002 | 0.004 | 1.0 | 0.002 | 0.004 |
| Fisher's Exact Test | NaN | 0.000 | NaN | NaN | 0.000 | NaN |
| WExT | 0.8 | 0.004 | 0.009 | 0.8 | 0.004 | 0.008 |

(b) Metrics for BRCA data. (1026 samples | 737 CGC-CGC pairs)

| Method | Precision | Sensitivity | F1 Score | $\text{Precision}_{\text{strict}}$ | $\text{Sensitivity}_{\text{strict}}$ | $\text{F1 Score}_{\text{strict}}$ |
|---|---|---|---|---|---|---|
| DISCOVER | 1.000 | 0.008 | 0.016 | 1.000 | 0.008 | 0.016 |
| DISCOVER Strat | 0.857 | 0.008 | 0.016 | 0.857 | 0.008 | 0.016 |
| Fisher's Exact Test | NaN | 0.000 | NaN | NaN | 0.000 | NaN |
| WExT | 0.692 | 0.012 | 0.024 | 0.692 | 0.012 | 0.024 |

(c) Metrics for COADREAD data. (498 samples | 1625 CGC-CGC pairs)

| Method | Precision | Sensitivity | F1 Score | $\text{Precision}_{\text{strict}}$ | $\text{Sensitivity}_{\text{strict}}$ | $\text{F1 Score}_{\text{strict}}$ |
|---|---|---|---|---|---|---|
| DISCOVER | 0.756 | 0.021 | 0.041 | 0.767 | 0.020 | 0.039 |
| DISCOVER Strat | 0.000 | 0.000 | NaN | 0.000 | 0.000 | NaN |
| Fisher's Exact Test | 0.286 | 0.001 | 0.002 | 0.286 | 0.001 | 0.002 |
| WExT | 0.716 | 0.068 | 0.125 | 0.775 | 0.057 | 0.106 |

(d) Metrics for LUAD data. (568 samples | 1197 CGC-CGC pairs)

| Method | Precision | Sensitivity | F1 Score | $\text{Precision}_{\text{strict}}$ | $\text{Sensitivity}_{\text{strict}}$ | $\text{F1 Score}_{\text{strict}}$ |
|---|---|---|---|---|---|---|
| DISCOVER | 0.800 | 0.003 | 0.007 | 0.800 | 0.003 | 0.006 |
| Fisher's Exact Test | NaN | 0.000 | NaN | NaN | 0.000 | NaN |
| WExT | 0.889 | 0.013 | 0.027 | 0.889 | 0.013 | 0.026 |

(e) Metrics for LUSC data. (485 samples | 906 CGC-CGC pairs)

| Method | Precision | Sensitivity | F1 Score | $\text{Precision}_{\text{strict}}$ | $\text{Sensitivity}_{\text{strict}}$ | $\text{F1 Score}_{\text{strict}}$ |
|---|---|---|---|---|---|---|
| DISCOVER | 1.000 | 0.002 | 0.004 | 1.000 | 0.002 | 0.004 |
| Fisher's Exact Test | NaN | 0.000 | NaN | NaN | 0.000 | NaN |
| WExT | 0.667 | 0.002 | 0.004 | 0.667 | 0.002 | 0.004 |

(f) Metrics for SKCM data. (468 samples | 2142 CGC-CGC pairs)

| Method | Precision | Sensitivity | F1 Score | $\text{Precision}_{\text{strict}}$ | $\text{Sensitivity}_{\text{strict}}$ | $\text{F1 Score}_{\text{strict}}$ |
|---|---|---|---|---|---|---|
| DISCOVER | 1.00 | 0.003 | 0.006 | 1.00 | 0.003 | 0.006 |
| Fisher's Exact Test | 1.00 | 0.001 | 0.002 | 1.00 | 0.001 | 0.002 |
| WExT | 0.87 | 0.009 | 0.019 | 0.87 | 0.009 | 0.018 |

(g) Metrics for STAD data. (438 samples | 1308 CGC-CGC pairs)

| Method | Precision | Sensitivity | F1 Score | $\text{Precision}_{\text{strict}}$ | $\text{Sensitivity}_{\text{strict}}$ | $\text{F1 Score}_{\text{strict}}$ |
|---|---|---|---|---|---|---|
| DISCOVER | 0.667 | 0.008 | 0.015 | 0.667 | 0.008 | 0.016 |
| Fisher's Exact Test | 1.000 | 0.002 | 0.003 | 1.000 | 0.002 | 0.004 |
| WExT | 0.679 | 0.028 | 0.053 | 0.744 | 0.025 | 0.048 |

(h) Metrics for UCEC data. (531 samples | 2668 CGC-CGC pairs)

| Method | Precision | Sensitivity | F1 Score | $\text{Precision}_{\text{strict}}$ | $\text{Sensitivity}_{\text{strict}}$ | $\text{F1 Score}_{\text{strict}}$ |
|---|---|---|---|---|---|---|
| DISCOVER | 0.740 | 0.074 | 0.134 | 0.813 | 0.069 | 0.127 |
| Fisher's Exact Test | 1.000 | 0.002 | 0.004 | 1.000 | 0.002 | 0.004 |
| WExT | 0.686 | 0.137 | 0.229 | 0.776 | 0.122 | 0.211 |

Table 41: **Results of network-centric ME evaluation framework with $\mathcal{G}$ = Intact (w conf. threshold 0.35), $\mathcal{S}$ = CGC, c= $X_2$, $p_t$ = 0.1, t=5, robustness_iterations = 100**

(a) Metrics for BLCA data. (411 samples | 895 CGC-CGC pairs)

| Method | Precision | Sensitivity | F1 Score | $\text{Precision}_{\text{strict}}$ | $\text{Sensitivity}_{\text{strict}}$ | $\text{F1 Score}_{\text{strict}}$ |
|---|---|---|---|---|---|---|
| DISCOVER | 0.909 | 0.022 | 0.044 | 0.909 | 0.022 | 0.043 |
| Fisher's Exact Test | 1.000 | 0.002 | 0.004 | 1.000 | 0.002 | 0.004 |
| WExT | 0.746 | 0.053 | 0.098 | 0.738 | 0.050 | 0.094 |

(b) Metrics for BRCA data. (1026 samples | 737 CGC-CGC pairs)

| Method | Precision | Sensitivity | F1 Score | $\text{Precision}_{\text{strict}}$ | $\text{Sensitivity}_{\text{strict}}$ | $\text{F1 Score}_{\text{strict}}$ |
|---|---|---|---|---|---|---|
| DISCOVER | 0.740 | 0.050 | 0.094 | 0.755 | 0.050 | 0.094 |
| DISCOVER Strat | 0.741 | 0.058 | 0.108 | 0.750 | 0.057 | 0.106 |
| Fisher's Exact Test | 1.000 | 0.003 | 0.005 | 1.000 | 0.003 | 0.006 |
| WExT | 0.700 | 0.104 | 0.181 | 0.720 | 0.093 | 0.165 |

(c) Metrics for COADREAD data. (498 samples | 1625 CGC-CGC pairs)

| Method | Precision | Sensitivity | F1 Score | $\text{Precision}_{\text{strict}}$ | $\text{Sensitivity}_{\text{strict}}$ | $\text{F1 Score}_{\text{strict}}$ |
|---|---|---|---|---|---|---|
| DISCOVER | 0.689 | 0.089 | 0.158 | 0.718 | 0.081 | 0.146 |
| DISCOVER Strat | 0.663 | 0.041 | 0.077 | 0.660 | 0.039 | 0.074 |
| Fisher's Exact Test | 0.698 | 0.018 | 0.036 | 0.692 | 0.017 | 0.033 |
| WExT | 0.659 | 0.183 | 0.286 | 0.703 | 0.163 | 0.265 |

(d) Metrics for LUAD data. (568 samples | 1197 CGC-CGC pairs)

| Method | Precision | Sensitivity | F1 Score | $\text{Precision}_{\text{strict}}$ | $\text{Sensitivity}_{\text{strict}}$ | $\text{F1 Score}_{\text{strict}}$ |
|---|---|---|---|---|---|---|
| DISCOVER | 0.813 | 0.031 | 0.060 | 0.847 | 0.030 | 0.058 |
| Fisher's Exact Test | 1.000 | 0.004 | 0.008 | 1.000 | 0.004 | 0.008 |
| WExT | 0.795 | 0.056 | 0.105 | 0.838 | 0.053 | 0.100 |

(e) Metrics for LUSC data. (485 samples | 906 CGC-CGC pairs)

| Method | Precision | Sensitivity | F1 Score | $\text{Precision}_{\text{strict}}$ | $\text{Sensitivity}_{\text{strict}}$ | $\text{F1 Score}_{\text{strict}}$ |
|---|---|---|---|---|---|---|
| DISCOVER | 0.800 | 0.004 | 0.009 | 0.800 | 0.004 | 0.008 |
| Fisher's Exact Test | 0.500 | 0.002 | 0.004 | 0.500 | 0.002 | 0.004 |
| WExT | 0.619 | 0.015 | 0.028 | 0.619 | 0.015 | 0.029 |

(f) Metrics for SKCM data. (468 samples | 2142 CGC-CGC pairs)

| Method | Precision | Sensitivity | F1 Score | $\text{Precision}_{\text{strict}}$ | $\text{Sensitivity}_{\text{strict}}$ | $\text{F1 Score}_{\text{strict}}$ |
|---|---|---|---|---|---|---|
| DISCOVER | 0.777 | 0.039 | 0.075 | 0.784 | 0.039 | 0.074 |
| Fisher's Exact Test | 1.000 | 0.002 | 0.004 | 1.000 | 0.002 | 0.004 |
| WExT | 0.716 | 0.080 | 0.144 | 0.726 | 0.079 | 0.142 |

(g) Metrics for STAD data. (438 samples | 1308 CGC-CGC pairs)

| Method | Precision | Sensitivity | F1 Score | $\text{Precision}_{\text{strict}}$ | $\text{Sensitivity}_{\text{strict}}$ | $\text{F1 Score}_{\text{strict}}$ |
|---|---|---|---|---|---|---|
| DISCOVER | 0.741 | 0.056 | 0.104 | 0.771 | 0.053 | 0.099 |
| Fisher's Exact Test | 0.727 | 0.006 | 0.012 | 0.727 | 0.006 | 0.012 |
| WExT | 0.658 | 0.098 | 0.171 | 0.706 | 0.089 | 0.158 |

(h) Metrics for UCEC data. (531 samples | 2668 CGC-CGC pairs)

| Method | Precision | Sensitivity | F1 Score | $\text{Precision}_{\text{strict}}$ | $\text{Sensitivity}_{\text{strict}}$ | $\text{F1 Score}_{\text{strict}}$ |
|---|---|---|---|---|---|---|
| DISCOVER | 0.658 | 0.173 | 0.274 | 0.725 | 0.154 | 0.254 |
| Fisher's Exact Test | 0.810 | 0.006 | 0.013 | 0.810 | 0.006 | 0.012 |
| WExT | 0.620 | 0.271 | 0.377 | 0.681 | 0.237 | 0.352 |

Table 42: **Results of network-centric ME evaluation framework with $\mathcal{G}$ = Intact (w conf. threshold 0.35), $\mathcal{S}$ = $CGC_{SNV}$ , c= $X_2$, $p_t$ = 0.05, t=5, robustness_iterations = 100**

(a) Metrics for BLCA data. (411 samples | 584 $CGC_{SNV}$-$CGC_{SNV}$ pairs)

| Method | Precision | Sensitivity | F1 Score | $\text{Precision}_{\text{strict}}$ | $\text{Sensitivity}_{\text{strict}}$ | $\text{F1 Score}_{\text{strict}}$ |
|---|---|---|---|---|---|---|
| DISCOVER | 1.000 | 0.014 | 0.027 | 1.000 | 0.014 | 0.028 |
| Fisher's Exact Test | 1.000 | 0.003 | 0.007 | 1.000 | 0.003 | 0.006 |
| WExT | 0.824 | 0.024 | 0.047 | 0.824 | 0.024 | 0.047 |

(b) Metrics for BRCA data. (1026 samples | 505 $CGC_{SNV}$-$CGC_{SNV}$ pairs)

| Method | Precision | Sensitivity | F1 Score | $\text{Precision}_{\text{strict}}$ | $\text{Sensitivity}_{\text{strict}}$ | $\text{F1 Score}_{\text{strict}}$ |
|---|---|---|---|---|---|---|
| DISCOVER | 0.800 | 0.032 | 0.061 | 0.800 | 0.032 | 0.062 |
| DISCOVER Strat | 0.846 | 0.044 | 0.083 | 0.846 | 0.044 | 0.084 |
| Fisher's Exact Test | 1.000 | 0.004 | 0.008 | 1.000 | 0.004 | 0.008 |
| WExT | 0.757 | 0.084 | 0.151 | 0.781 | 0.082 | 0.148 |

(c) Metrics for COADREAD data. (498 samples | 919 $CGC_{SNV}$-$CGC_{SNV}$ pairs)

| Method | Precision | Sensitivity | F1 Score | $\text{Precision}_{\text{strict}}$ | $\text{Sensitivity}_{\text{strict}}$ | $\text{F1 Score}_{\text{strict}}$ |
|---|---|---|---|---|---|---|
| DISCOVER | 0.725 | 0.068 | 0.124 | 0.771 | 0.064 | 0.118 |
| DISCOVER Strat | 0.731 | 0.021 | 0.040 | 0.731 | 0.021 | 0.041 |
| Fisher's Exact Test | 0.688 | 0.012 | 0.024 | 0.688 | 0.012 | 0.024 |
| WExT | 0.697 | 0.144 | 0.239 | 0.752 | 0.128 | 0.219 |

(d) Metrics for LUAD data. (568 samples | 645 $CGC_{SNV}$-$CGC_{SNV}$ pairs)

| Method | Precision | Sensitivity | F1 Score | $\text{Precision}_{\text{strict}}$ | $\text{Sensitivity}_{\text{strict}}$ | $\text{F1 Score}_{\text{strict}}$ |
|---|---|---|---|---|---|---|
| DISCOVER | 0.867 | 0.020 | 0.039 | 0.867 | 0.020 | 0.039 |
| Fisher's Exact Test | NaN | 0.000 | NaN | NaN | 0.000 | NaN |
| WExT | 0.816 | 0.049 | 0.092 | 0.833 | 0.047 | 0.089 |

(e) Metrics for LUSC data. (485 samples | 586 $CGC_{SNV}$-$CGC_{SNV}$ pairs)

| Method | Precision | Sensitivity | F1 Score | $\text{Precision}_{\text{strict}}$ | $\text{Sensitivity}_{\text{strict}}$ | $\text{F1 Score}_{\text{strict}}$ |
|---|---|---|---|---|---|---|
| DISCOVER | 1.0 | 0.003 | 0.007 | 1.0 | 0.003 | 0.006 |
| Fisher's Exact Test | 1.0 | 0.003 | 0.007 | 1.0 | 0.003 | 0.006 |
| WExT | 0.6 | 0.010 | 0.020 | 0.6 | 0.010 | 0.020 |

(f) Metrics for SKCM data. (468 samples | 1104 $CGC_{SNV}$-$CGC_{SNV}$ pairs)

| Method | Precision | Sensitivity | F1 Score | $\text{Precision}_{\text{strict}}$ | $\text{Sensitivity}_{\text{strict}}$ | $\text{F1 Score}_{\text{strict}}$ |
|---|---|---|---|---|---|---|
| DISCOVER | 0.789 | 0.014 | 0.027 | 0.833 | 0.014 | 0.028 |
| Fisher's Exact Test | 1.000 | 0.002 | 0.004 | 1.000 | 0.002 | 0.004 |
| WExT | 0.849 | 0.057 | 0.107 | 0.871 | 0.056 | 0.105 |

(g) Metrics for STAD data. (438 samples | 722 $CGC_{SNV}$-$CGC_{SNV}$ pairs)

| Method | Precision | Sensitivity | F1 Score | $\text{Precision}_{\text{strict}}$ | $\text{Sensitivity}_{\text{strict}}$ | $\text{F1 Score}_{\text{strict}}$ |
|---|---|---|---|---|---|---|
| DISCOVER | 0.734 | 0.033 | 0.062 | 0.750 | 0.031 | 0.060 |
| Fisher's Exact Test | 0.667 | 0.003 | 0.006 | 0.667 | 0.003 | 0.006 |
| WExT | 0.733 | 0.089 | 0.158 | 0.781 | 0.080 | 0.145 |

(h) Metrics for UCEC data. (531 samples | 1255 $CGC_{SNV}$-$CGC_{SNV}$ pairs)

| Method | Precision | Sensitivity | F1 Score | $\text{Precision}_{\text{strict}}$ | $\text{Sensitivity}_{\text{strict}}$ | $\text{F1 Score}_{\text{strict}}$ |
|---|---|---|---|---|---|---|
| DISCOVER | 0.721 | 0.179 | 0.287 | 0.812 | 0.163 | 0.271 |
| Fisher's Exact Test | 0.909 | 0.008 | 0.016 | 0.909 | 0.008 | 0.016 |
| WExT | 0.680 | 0.291 | 0.408 | 0.773 | 0.259 | 0.388 |

Table 43: **Results of network-centric ME evaluation framework with $\mathcal{G}$ = Intact (w conf. threshold 0.35), $\mathcal{S}$ = Intogen , c= $X_2$, $p_t$ = 0.05, t=5, robustness_iterations = 100**

(a) Metrics for BLCA data. (411 samples | 844 Intogen-Intogen pairs)

| Method | Precision | Sensitivity | F1 Score | $\text{Precision}_{\text{strict}}$ | $\text{Sensitivity}_{\text{strict}}$ | $\text{F1 Score}_{\text{strict}}$ |
|---|---|---|---|---|---|---|
| DISCOVER | 1.00 | 0.019 | 0.037 | 1.00 | 0.019 | 0.037 |
| Fisher's Exact Test | 1.00 | 0.002 | 0.005 | 1.00 | 0.002 | 0.004 |
| WExT | 0.83 | 0.026 | 0.051 | 0.83 | 0.026 | 0.050 |

(b) Metrics for BRCA data. (1026 samples | 754 Intogen-Intogen pairs)

| Method | Precision | Sensitivity | F1 Score | $\text{Precision}_{\text{strict}}$ | $\text{Sensitivity}_{\text{strict}}$ | $\text{F1 Score}_{\text{strict}}$ |
|---|---|---|---|---|---|---|
| DISCOVER | 0.800 | 0.027 | 0.051 | 0.800 | 0.027 | 0.052 |
| DISCOVER Strat | 0.862 | 0.033 | 0.064 | 0.862 | 0.033 | 0.064 |
| Fisher's Exact Test | 1.000 | 0.003 | 0.005 | 1.000 | 0.003 | 0.006 |
| WExT | 0.708 | 0.068 | 0.124 | 0.727 | 0.064 | 0.118 |

(c) Metrics for COADREAD data. (498 samples | 1495 Intogen-Intogen pairs)

| Method | Precision | Sensitivity | F1 Score | $\text{Precision}_{\text{strict}}$ | $\text{Sensitivity}_{\text{strict}}$ | $\text{F1 Score}_{\text{strict}}$ |
|---|---|---|---|---|---|---|
| DISCOVER | 0.754 | 0.068 | 0.124 | 0.790 | 0.063 | 0.117 |
| DISCOVER Strat | 0.775 | 0.021 | 0.041 | 0.775 | 0.021 | 0.041 |
| Fisher's Exact Test | 0.591 | 0.009 | 0.017 | 0.591 | 0.009 | 0.018 |
| WExT | 0.704 | 0.141 | 0.236 | 0.765 | 0.126 | 0.216 |

(d) Metrics for LUAD data. (568 samples | 1056 Intogen-Intogen pairs)

| Method | Precision | Sensitivity | F1 Score | $\text{Precision}_{\text{strict}}$ | $\text{Sensitivity}_{\text{strict}}$ | $\text{F1 Score}_{\text{strict}}$ |
|---|---|---|---|---|---|---|
| DISCOVER | 0.842 | 0.015 | 0.030 | 0.842 | 0.015 | 0.029 |
| Fisher's Exact Test | NaN | 0.000 | NaN | NaN | 0.000 | NaN |
| WExT | 0.812 | 0.037 | 0.071 | 0.826 | 0.036 | 0.069 |

(e) Metrics for LUSC data. (485 samples | 867 Intogen-Intogen pairs)

| Method | Precision | Sensitivity | F1 Score | $\text{Precision}_{\text{strict}}$ | $\text{Sensitivity}_{\text{strict}}$ | $\text{F1 Score}_{\text{strict}}$ |
|---|---|---|---|---|---|---|
| DISCOVER | 1.000 | 0.003 | 0.007 | 1.000 | 0.003 | 0.006 |
| Fisher's Exact Test | 0.667 | 0.002 | 0.005 | 0.667 | 0.002 | 0.004 |
| WExT | 0.692 | 0.010 | 0.021 | 0.692 | 0.010 | 0.020 |

(f) Metrics for SKCM data. (468 samples | 1774 Intogen-Intogen pairs)

| Method | Precision | Sensitivity | F1 Score | $\text{Precision}_{\text{strict}}$ | $\text{Sensitivity}_{\text{strict}}$ | $\text{F1 Score}_{\text{strict}}$ |
|---|---|---|---|---|---|---|
| DISCOVER | 0.833 | 0.014 | 0.028 | 0.833 | 0.014 | 0.028 |
| Fisher's Exact Test | 1.000 | 0.001 | 0.002 | 1.000 | 0.001 | 0.002 |
| WExT | 0.774 | 0.048 | 0.090 | 0.787 | 0.047 | 0.089 |

(g) Metrics for STAD data. (438 samples | 1258 Intogen-Intogen pairs)

| Method | Precision | Sensitivity | F1 Score | $\text{Precision}_{\text{strict}}$ | $\text{Sensitivity}_{\text{strict}}$ | $\text{F1 Score}_{\text{strict}}$ |
|---|---|---|---|---|---|---|
| DISCOVER | 0.778 | 0.033 | 0.064 | 0.800 | 0.032 | 0.062 |
| Fisher's Exact Test | 0.833 | 0.004 | 0.008 | 0.833 | 0.004 | 0.008 |
| WExT | 0.751 | 0.084 | 0.152 | 0.808 | 0.077 | 0.141 |

(h) Metrics for UCEC data. (531 samples | 2029 Intogen-Intogen pairs)

| Method | Precision | Sensitivity | F1 Score | $\text{Precision}_{\text{strict}}$ | $\text{Sensitivity}_{\text{strict}}$ | $\text{F1 Score}_{\text{strict}}$ |
|---|---|---|---|---|---|---|
| DISCOVER | 0.719 | 0.171 | 0.276 | 0.811 | 0.155 | 0.260 |
| Fisher's Exact Test | 0.875 | 0.007 | 0.014 | 0.875 | 0.007 | 0.014 |
| WExT | 0.664 | 0.257 | 0.371 | 0.762 | 0.228 | 0.351 |

Table 44: **Results of network-centric ME evaluation framework with $\mathcal{G}$ = Intact (w conf. threshold 0.25), $\mathcal{S}$ = CGC, c= $X_2$, $p_t$ = 0.05, t=5, robustness_iterations = 100**

(a) Metrics for BLCA data. (411 samples | 1439 CGC-CGC pairs)

| Method | Precision | Sensitivity | F1 Score | $\text{Precision}_{\text{strict}}$ | $\text{Sensitivity}_{\text{strict}}$ | $\text{F1 Score}_{\text{strict}}$ |
|---|---|---|---|---|---|---|
| DISCOVER | 1.000 | 0.013 | 0.025 | 1.000 | 0.013 | 0.026 |
| Fisher's Exact Test | 1.000 | 0.004 | 0.008 | 1.000 | 0.004 | 0.008 |
| WExT | 0.833 | 0.021 | 0.041 | 0.833 | 0.021 | 0.041 |

(b) Metrics for BRCA data. (1026 samples | 1238 CGC-CGC pairs)

| Method | Precision | Sensitivity | F1 Score | $\text{Precision}_{\text{strict}}$ | $\text{Sensitivity}_{\text{strict}}$ | $\text{F1 Score}_{\text{strict}}$ |
|---|---|---|---|---|---|---|
| DISCOVER | 0.827 | 0.027 | 0.053 | 0.827 | 0.027 | 0.052 |
| DISCOVER Strat | 0.927 | 0.031 | 0.059 | 0.927 | 0.031 | 0.060 |
| Fisher's Exact Test | 1.000 | 0.003 | 0.006 | 1.000 | 0.003 | 0.006 |
| WExT | 0.746 | 0.058 | 0.108 | 0.771 | 0.056 | 0.104 |

(c) Metrics for COADREAD data. (498 samples | 2494 CGC-CGC pairs)

| Method | Precision | Sensitivity | F1 Score | $\text{Precision}_{\text{strict}}$ | $\text{Sensitivity}_{\text{strict}}$ | $\text{F1 Score}_{\text{strict}}$ |
|---|---|---|---|---|---|---|
| DISCOVER | 0.702 | 0.043 | 0.081 | 0.736 | 0.040 | 0.076 |
| DISCOVER Strat | 0.659 | 0.012 | 0.023 | 0.659 | 0.012 | 0.024 |
| Fisher's Exact Test | 0.667 | 0.009 | 0.017 | 0.677 | 0.008 | 0.016 |
| WExT | 0.668 | 0.104 | 0.180 | 0.709 | 0.092 | 0.163 |

(d) Metrics for LUAD data. (568 samples | 1862 CGC-CGC pairs)

| Method | Precision | Sensitivity | F1 Score | $\text{Precision}_{\text{strict}}$ | $\text{Sensitivity}_{\text{strict}}$ | $\text{F1 Score}_{\text{strict}}$ |
|---|---|---|---|---|---|---|
| DISCOVER | 0.848 | 0.015 | 0.030 | 0.875 | 0.015 | 0.029 |
| Fisher's Exact Test | 1.000 | 0.002 | 0.004 | 1.000 | 0.002 | 0.004 |
| WExT | 0.818 | 0.030 | 0.058 | 0.840 | 0.030 | 0.058 |

(e) Metrics for LUSC data. (485 samples | 1417 CGC-CGC pairs)

| Method | Precision | Sensitivity | F1 Score | $\text{Precision}_{\text{strict}}$ | $\text{Sensitivity}_{\text{strict}}$ | $\text{F1 Score}_{\text{strict}}$ |
|---|---|---|---|---|---|---|
| DISCOVER | 1.000 | 0.001 | 0.003 | 1.000 | 0.001 | 0.002 |
| Fisher's Exact Test | 1.000 | 0.001 | 0.003 | 1.000 | 0.001 | 0.002 |
| WExT | 0.696 | 0.006 | 0.011 | 0.696 | 0.006 | 0.012 |

(f) Metrics for SKCM data. (468 samples | 3113 CGC-CGC pairs)

| Method | Precision | Sensitivity | F1 Score | $\text{Precision}_{\text{strict}}$ | $\text{Sensitivity}_{\text{strict}}$ | $\text{F1 Score}_{\text{strict}}$ |
|---|---|---|---|---|---|---|
| DISCOVER | 0.839 | 0.013 | 0.025 | 0.839 | 0.013 | 0.026 |
| Fisher's Exact Test | 1.000 | 0.001 | 0.001 | 1.000 | 0.001 | 0.002 |
| WExT | 0.766 | 0.036 | 0.069 | 0.775 | 0.036 | 0.069 |

(g) Metrics for STAD data. (438 samples | 2194 CGC-CGC pairs)

| Method | Precision | Sensitivity | F1 Score | $\text{Precision}_{\text{strict}}$ | $\text{Sensitivity}_{\text{strict}}$ | $\text{F1 Score}_{\text{strict}}$ |
|---|---|---|---|---|---|---|
| DISCOVER | 0.768 | 0.029 | 0.055 | 0.803 | 0.028 | 0.054 |
| Fisher's Exact Test | 0.889 | 0.004 | 0.007 | 0.889 | 0.004 | 0.008 |
| WExT | 0.729 | 0.062 | 0.115 | 0.794 | 0.056 | 0.105 |

(h) Metrics for UCEC data. (531 samples | 4046 CGC-CGC pairs)

| Method | Precision | Sensitivity | F1 Score | $\text{Precision}_{\text{strict}}$ | $\text{Sensitivity}_{\text{strict}}$ | $\text{F1 Score}_{\text{strict}}$ |
|---|---|---|---|---|---|---|
| DISCOVER | 0.702 | 0.116 | 0.200 | 0.776 | 0.105 | 0.185 |
| Fisher's Exact Test | 0.875 | 0.003 | 0.007 | 0.875 | 0.003 | 0.006 |
| WExT | 0.657 | 0.201 | 0.308 | 0.727 | 0.179 | 0.287 |

Table 45: **Results of network-centric ME evaluation framework with $\mathcal{G}$ = Intact (w conf. threshold 0.45), $\mathcal{S}$ = CGC, c= $X_2$, $p_t$ = 0.05, t=5, robustness_iterations = 100**

(a) Metrics for BLCA data. (411 samples | 359 CGC-CGC pairs)

| Method | Precision | Sensitivity | F1 Score | $\text{Precision}_{\text{strict}}$ | $\text{Sensitivity}_{\text{strict}}$ | $\text{F1 Score}_{\text{strict}}$ |
|---|---|---|---|---|---|---|
| DISCOVER | 1.000 | 0.011 | 0.022 | 1.000 | 0.011 | 0.022 |
| Fisher's Exact Test | 1.000 | 0.006 | 0.011 | 1.000 | 0.006 | 0.012 |
| WExT | 0.667 | 0.017 | 0.033 | 0.667 | 0.017 | 0.033 |

(b) Metrics for BRCA data. (1026 samples | 286 CGC-CGC pairs)

| Method | Precision | Sensitivity | F1 Score | $\text{Precision}_{\text{strict}}$ | $\text{Sensitivity}_{\text{strict}}$ | $\text{F1 Score}_{\text{strict}}$ |
|---|---|---|---|---|---|---|
| DISCOVER | 0.571 | 0.014 | 0.027 | 0.571 | 0.014 | 0.027 |
| DISCOVER Strat | 0.727 | 0.028 | 0.054 | 0.727 | 0.028 | 0.054 |
| Fisher's Exact Test | 1.000 | 0.003 | 0.007 | 1.000 | 0.003 | 0.006 |
| WExT | 0.649 | 0.065 | 0.119 | 0.636 | 0.062 | 0.113 |

(c) Metrics for COADREAD data. (498 samples | 605 CGC-CGC pairs)

| Method | Precision | Sensitivity | F1 Score | $\text{Precision}_{\text{strict}}$ | $\text{Sensitivity}_{\text{strict}}$ | $\text{F1 Score}_{\text{strict}}$ |
|---|---|---|---|---|---|---|
| DISCOVER | 0.712 | 0.061 | 0.113 | 0.739 | 0.056 | 0.104 |
| DISCOVER Strat | 0.643 | 0.015 | 0.029 | 0.643 | 0.015 | 0.029 |
| Fisher's Exact Test | 0.545 | 0.010 | 0.020 | 0.545 | 0.010 | 0.020 |
| WExT | 0.704 | 0.135 | 0.227 | 0.767 | 0.115 | 0.200 |

(d) Metrics for LUAD data. (568 samples | 448 CGC-CGC pairs)

| Method | Precision | Sensitivity | F1 Score | $\text{Precision}_{\text{strict}}$ | $\text{Sensitivity}_{\text{strict}}$ | $\text{F1 Score}_{\text{strict}}$ |
|---|---|---|---|---|---|---|
| DISCOVER | 0.857 | 0.013 | 0.026 | 0.857 | 0.013 | 0.026 |
| Fisher's Exact Test | NaN | 0.000 | NaN | NaN | 0.000 | NaN |
| WExT | 0.778 | 0.032 | 0.061 | 0.778 | 0.032 | 0.061 |

(e) Metrics for LUSC data. (485 samples | 359 CGC-CGC pairs)

| Method | Precision | Sensitivity | F1 Score | $\text{Precision}_{\text{strict}}$ | $\text{Sensitivity}_{\text{strict}}$ | $\text{F1 Score}_{\text{strict}}$ |
|---|---|---|---|---|---|---|
| DISCOVER | 1.000 | 0.006 | 0.011 | 1.000 | 0.006 | 0.012 |
| Fisher's Exact Test | 0.667 | 0.006 | 0.011 | 0.667 | 0.006 | 0.012 |
| WExT | 0.714 | 0.014 | 0.027 | 0.714 | 0.014 | 0.027 |

(f) Metrics for SKCM data. (468 samples | 860 CGC-CGC pairs)

| Method | Precision | Sensitivity | F1 Score | $\text{Precision}_{\text{strict}}$ | $\text{Sensitivity}_{\text{strict}}$ | $\text{F1 Score}_{\text{strict}}$ |
|---|---|---|---|---|---|---|
| DISCOVER | 0.857 | 0.014 | 0.028 | 0.857 | 0.014 | 0.028 |
| Fisher's Exact Test | NaN | 0.000 | NaN | NaN | 0.000 | NaN |
| WExT | 0.844 | 0.045 | 0.085 | 0.844 | 0.045 | 0.085 |

(g) Metrics for STAD data. (438 samples | 545 CGC-CGC pairs)

| Method | Precision | Sensitivity | F1 Score | $\text{Precision}_{\text{strict}}$ | $\text{Sensitivity}_{\text{strict}}$ | $\text{F1 Score}_{\text{strict}}$ |
|---|---|---|---|---|---|---|
| DISCOVER | 0.690 | 0.037 | 0.070 | 0.731 | 0.035 | 0.067 |
| Fisher's Exact Test | 0.667 | 0.004 | 0.007 | 0.667 | 0.004 | 0.008 |
| WExT | 0.652 | 0.083 | 0.147 | 0.696 | 0.072 | 0.130 |

(h) Metrics for UCEC data. (531 samples | 1096 CGC-CGC pairs)

| Method | Precision | Sensitivity | F1 Score | $\text{Precision}_{\text{strict}}$ | $\text{Sensitivity}_{\text{strict}}$ | $\text{F1 Score}_{\text{strict}}$ |
|---|---|---|---|---|---|---|
| DISCOVER | 0.683 | 0.152 | 0.249 | 0.762 | 0.135 | 0.229 |
| Fisher's Exact Test | 0.800 | 0.007 | 0.014 | 0.800 | 0.007 | 0.014 |
| WExT | 0.632 | 0.226 | 0.333 | 0.708 | 0.192 | 0.302 |

Table 46: **Results of network-centric ME evaluation framework with $\mathcal{G}$ = HINT, $\mathcal{S}$ = CGC, c= $X_2$, $p_t$ = 0.05, t=5, robustness_iterations = 100**

(a) Metrics for BLCA data. (411 samples | 577 CGC-CGC pairs)

| Method | Precision | Sensitivity | F1 Score | $\text{Precision}_{\text{strict}}$ | $\text{Sensitivity}_{\text{strict}}$ | $\text{F1 Score}_{\text{strict}}$ |
|---|---|---|---|---|---|---|
| DISCOVER | 0.867 | 0.023 | 0.044 | 0.867 | 0.023 | 0.045 |
| Fisher's Exact Test | 1.000 | 0.005 | 0.010 | 1.000 | 0.005 | 0.010 |
| WExT | 0.710 | 0.038 | 0.073 | 0.710 | 0.038 | 0.072 |

(b) Metrics for BRCA data. (1026 samples | 427 CGC-CGC pairs)

| Method | Precision | Sensitivity | F1 Score | $\text{Precision}_{\text{strict}}$ | $\text{Sensitivity}_{\text{strict}}$ | $\text{F1 Score}_{\text{strict}}$ |
|---|---|---|---|---|---|---|
| DISCOVER | 0.800 | 0.028 | 0.054 | 0.800 | 0.028 | 0.054 |
| DISCOVER Strat | 0.800 | 0.037 | 0.072 | 0.842 | 0.037 | 0.071 |
| Fisher's Exact Test | 1.000 | 0.005 | 0.009 | 1.000 | 0.005 | 0.010 |
| WExT | 0.813 | 0.087 | 0.157 | 0.833 | 0.082 | 0.149 |

(c) Metrics for COADREAD data. (498 samples | 1059 CGC-CGC pairs)

| Method | Precision | Sensitivity | F1 Score | $\text{Precision}_{\text{strict}}$ | $\text{Sensitivity}_{\text{strict}}$ | $\text{F1 Score}_{\text{strict}}$ |
|---|---|---|---|---|---|---|
| DISCOVER | 0.736 | 0.074 | 0.135 | 0.777 | 0.069 | 0.127 |
| DISCOVER Strat | 0.719 | 0.030 | 0.058 | 0.736 | 0.030 | 0.058 |
| Fisher's Exact Test | 0.733 | 0.021 | 0.040 | 0.750 | 0.020 | 0.039 |
| WExT | 0.644 | 0.143 | 0.234 | 0.721 | 0.126 | 0.215 |

(d) Metrics for LUAD data. (568 samples | 748 CGC-CGC pairs)

| Method | Precision | Sensitivity | F1 Score | $\text{Precision}_{\text{strict}}$ | $\text{Sensitivity}_{\text{strict}}$ | $\text{F1 Score}_{\text{strict}}$ |
|---|---|---|---|---|---|---|
| DISCOVER | 0.583 | 0.009 | 0.018 | 0.583 | 0.009 | 0.018 |
| Fisher's Exact Test | 1.000 | 0.001 | 0.003 | 1.000 | 0.001 | 0.002 |
| WExT | 0.733 | 0.030 | 0.057 | 0.724 | 0.028 | 0.054 |

(e) Metrics for LUSC data. (485 samples | 524 CGC-CGC pairs)

| Method | Precision | Sensitivity | F1 Score | $\text{Precision}_{\text{strict}}$ | $\text{Sensitivity}_{\text{strict}}$ | $\text{F1 Score}_{\text{strict}}$ |
|---|---|---|---|---|---|---|
| DISCOVER | 1.000 | 0.004 | 0.008 | 1.000 | 0.004 | 0.008 |
| Fisher's Exact Test | 0.667 | 0.004 | 0.008 | 0.667 | 0.004 | 0.008 |
| WExT | 0.500 | 0.008 | 0.015 | 0.500 | 0.008 | 0.016 |

(f) Metrics for SKCM data. (468 samples | 1324 CGC-CGC pairs)

| Method | Precision | Sensitivity | F1 Score | $\text{Precision}_{\text{strict}}$ | $\text{Sensitivity}_{\text{strict}}$ | $\text{F1 Score}_{\text{strict}}$ |
|---|---|---|---|---|---|---|
| DISCOVER | 0.875 | 0.011 | 0.021 | 0.875 | 0.011 | 0.022 |
| Fisher's Exact Test | NaN | 0.000 | NaN | NaN | 0.000 | NaN |
| WExT | 0.816 | 0.039 | 0.074 | 0.816 | 0.039 | 0.074 |

(g) Metrics for STAD data. (438 samples | 854 CGC-CGC pairs)

| Method | Precision | Sensitivity | F1 Score | $\text{Precision}_{\text{strict}}$ | $\text{Sensitivity}_{\text{strict}}$ | $\text{F1 Score}_{\text{strict}}$ |
|---|---|---|---|---|---|---|
| DISCOVER | 0.619 | 0.030 | 0.058 | 0.658 | 0.029 | 0.056 |
| Fisher's Exact Test | 0.667 | 0.002 | 0.005 | 0.667 | 0.002 | 0.004 |
| WExT | 0.692 | 0.090 | 0.160 | 0.735 | 0.079 | 0.143 |

(h) Metrics for UCEC data. (531 samples | 1707 CGC-CGC pairs)

| Method | Precision | Sensitivity | F1 Score | $\text{Precision}_{\text{strict}}$ | $\text{Sensitivity}_{\text{strict}}$ | $\text{F1 Score}_{\text{strict}}$ |
|---|---|---|---|---|---|---|
| DISCOVER | 0.701 | 0.160 | 0.261 | 0.800 | 0.145 | 0.246 |
| Fisher's Exact Test | 0.846 | 0.006 | 0.013 | 0.846 | 0.006 | 0.012 |
| WExT | 0.660 | 0.254 | 0.367 | 0.755 | 0.225 | 0.347 |

Table 47: **Results of network-centric ME evaluation framework with $\mathcal{G}$ = STRING, $\mathcal{S}$ = CGC, c= $X_2$, $p_t$ = 0.05, t=5, robustness_iterations = 100**

(a) Metrics for BLCA data. (411 samples | 1210 CGC-CGC pairs)

| Method | Precision | Sensitivity | F1 Score | $\text{Precision}_{\text{strict}}$ | $\text{Sensitivity}_{\text{strict}}$ | $\text{F1 Score}_{\text{strict}}$ |
|---|---|---|---|---|---|---|
| DISCOVER | 1.000 | 0.011 | 0.021 | 1.000 | 0.011 | 0.022 |
| Fisher's Exact Test | 1.000 | 0.003 | 0.007 | 1.000 | 0.003 | 0.006 |
| WExT | 0.724 | 0.017 | 0.034 | 0.724 | 0.017 | 0.033 |

(b) Metrics for BRCA data. (1026 samples | 818 CGC-CGC pairs)

| Method | Precision | Sensitivity | F1 Score | $\text{Precision}_{\text{strict}}$ | $\text{Sensitivity}_{\text{strict}}$ | $\text{F1 Score}_{\text{strict}}$ |
|---|---|---|---|---|---|---|
| DISCOVER | 0.903 | 0.034 | 0.066 | 0.933 | 0.034 | 0.066 |
| DISCOVER Strat | 0.968 | 0.037 | 0.071 | 0.968 | 0.037 | 0.071 |
| Fisher's Exact Test | NaN | 0.000 | NaN | NaN | 0.000 | NaN |
| WExT | 0.794 | 0.068 | 0.126 | 0.857 | 0.066 | 0.123 |

(c) Metrics for COADREAD data. (498 samples | 1906 CGC-CGC pairs)

| Method | Precision | Sensitivity | F1 Score | $\text{Precision}_{\text{strict}}$ | $\text{Sensitivity}_{\text{strict}}$ | $\text{F1 Score}_{\text{strict}}$ |
|---|---|---|---|---|---|---|
| DISCOVER | 0.742 | 0.048 | 0.091 | 0.796 | 0.045 | 0.085 |
| DISCOVER Strat | 0.687 | 0.018 | 0.035 | 0.716 | 0.018 | 0.035 |
| Fisher's Exact Test | 0.857 | 0.009 | 0.019 | 0.857 | 0.009 | 0.018 |
| WExT | 0.707 | 0.096 | 0.168 | 0.768 | 0.087 | 0.156 |

(d) Metrics for LUAD data. (568 samples | 1330 CGC-CGC pairs)

| Method | Precision | Sensitivity | F1 Score | $\text{Precision}_{\text{strict}}$ | $\text{Sensitivity}_{\text{strict}}$ | $\text{F1 Score}_{\text{strict}}$ |
|---|---|---|---|---|---|---|
| DISCOVER | 0.792 | 0.014 | 0.028 | 0.826 | 0.014 | 0.028 |
| Fisher's Exact Test | 1.000 | 0.008 | 0.015 | 1.000 | 0.008 | 0.016 |
| WExT | 0.721 | 0.023 | 0.045 | 0.763 | 0.022 | 0.043 |

(e) Metrics for LUSC data. (485 samples | 1057 CGC-CGC pairs)

| Method | Precision | Sensitivity | F1 Score | $\text{Precision}_{\text{strict}}$ | $\text{Sensitivity}_{\text{strict}}$ | $\text{F1 Score}_{\text{strict}}$ |
|---|---|---|---|---|---|---|
| DISCOVER | 1.000 | 0.003 | 0.006 | 1.000 | 0.003 | 0.006 |
| Fisher's Exact Test | 1.000 | 0.001 | 0.002 | 1.000 | 0.001 | 0.002 |
| WExT | 0.667 | 0.004 | 0.008 | 0.667 | 0.004 | 0.008 |

(f) Metrics for SKCM data. (468 samples | 2309 CGC-CGC pairs)

| Method | Precision | Sensitivity | F1 Score | $\text{Precision}_{\text{strict}}$ | $\text{Sensitivity}_{\text{strict}}$ | $\text{F1 Score}_{\text{strict}}$ |
|---|---|---|---|---|---|---|
| DISCOVER | 0.740 | 0.016 | 0.032 | 0.750 | 0.016 | 0.031 |
| Fisher's Exact Test | 1.000 | 0.002 | 0.003 | 1.000 | 0.002 | 0.004 |
| WExT | 0.722 | 0.036 | 0.069 | 0.743 | 0.035 | 0.067 |

(g) Metrics for STAD data. (438 samples | 1517 CGC-CGC pairs)

| Method | Precision | Sensitivity | F1 Score | $\text{Precision}_{\text{strict}}$ | $\text{Sensitivity}_{\text{strict}}$ | $\text{F1 Score}_{\text{strict}}$ |
|---|---|---|---|---|---|---|
| DISCOVER | 0.818 | 0.018 | 0.035 | 0.839 | 0.017 | 0.033 |
| Fisher's Exact Test | 1.000 | 0.001 | 0.003 | 1.000 | 0.001 | 0.002 |
| WExT | 0.742 | 0.039 | 0.074 | 0.806 | 0.037 | 0.071 |

(h) Metrics for UCEC data. (531 samples | 2890 CGC-CGC pairs)

| Method | Precision | Sensitivity | F1 Score | $\text{Precision}_{\text{strict}}$ | $\text{Sensitivity}_{\text{strict}}$ | $\text{F1 Score}_{\text{strict}}$ |
|---|---|---|---|---|---|---|
| DISCOVER | 0.728 | 0.114 | 0.197 | 0.803 | 0.104 | 0.184 |
| Fisher's Exact Test | 0.923 | 0.004 | 0.008 | 0.923 | 0.004 | 0.008 |
| WExT | 0.663 | 0.197 | 0.304 | 0.737 | 0.174 | 0.282 |

Table 48: **Degree-normalized network-centric evaluations $X_2$ and $t = 5$**

(a) Metrics for BLCA data.

| Method | Precision | Sensitivity | F1 Score | $\text{Precision}_{\text{strict}}$ | $\text{Sensitivity}_{\text{strict}}$ | $\text{F1 Score}_{\text{strict}}$ |
|---|---|---|---|---|---|---|
| DISCOVER | 1.0 | 1.0 | 0.366 | 1.0 | 0.267 | 0.366 |
| Fisher's Exact Test | 1.0 | 1.0 | 0.133 | 1.0 | 0.072 | 0.133 |
| WExT | 0.89 | 0.977 | 0.411 | 0.89 | 0.292 | 0.411 |

(b) Metrics for BRCA data.

| Method | Precision | Sensitivity | F1 Score | $\text{Precision}_{\text{strict}}$ | $\text{Sensitivity}_{\text{strict}}$ | $\text{F1 Score}_{\text{strict}}$ |
|---|---|---|---|---|---|---|
| DISCOVER | 0.808 | 0.899 | 0.509 | 0.808 | 0.411 | 0.509 |
| DISCOVER Strat | 0.951 | 0.992 | 0.476 | 0.951 | 0.406 | 0.476 |
| Fisher's Exact Test | 1.0 | 1.0 | 0.373 | 1.0 | 0.271 | 0.373 |
| WExT | 0.846 | 0.911 | 0.46 | 0.85 | 0.344 | 0.458 |

(c) Metrics for COADREAD data.

| Method | Precision | Sensitivity | F1 Score | $\text{Precision}_{\text{strict}}$ | $\text{Sensitivity}_{\text{strict}}$ | $\text{F1 Score}_{\text{strict}}$ |
|---|---|---|---|---|---|---|
| DISCOVER | 0.87 | 0.952 | 0.385 | 0.874 | 0.267 | 0.383 |
| DISCOVER Strat | 0.707 | 0.961 | 0.206 | 0.707 | 0.097 | 0.206 |
| Fisher's Exact Test | 0.881 | 0.961 | 0.289 | 0.881 | 0.195 | 0.289 |
| WExT | 0.818 | 0.906 | 0.441 | 0.822 | 0.307 | 0.435 |

(d) Metrics for LUAD data.

| Method | Precision | Sensitivity | F1 Score | $\text{Precision}_{\text{strict}}$ | $\text{Sensitivity}_{\text{strict}}$ | $\text{F1 Score}_{\text{strict}}$ |
|---|---|---|---|---|---|---|
| DISCOVER | 0.924 | 0.977 | 0.468 | 0.924 | 0.369 | 0.468 |
| Fisher's Exact Test | NaN | NaN | NaN | NaN | NaN | NaN |
| WExT | 0.942 | 0.981 | 0.399 | 0.943 | 0.299 | 0.398 |

(e) Metrics for LUSC data.

| Method | Precision | Sensitivity | F1 Score | $\text{Precision}_{\text{strict}}$ | $\text{Sensitivity}_{\text{strict}}$ | $\text{F1 Score}_{\text{strict}}$ |
|---|---|---|---|---|---|---|
| DISCOVER | 1.0 | 1.0 | 0.667 | 1.0 | 0.6 | 0.667 |
| Fisher's Exact Test | 0.667 | 0.986 | 0.667 | 0.667 | 0.4 | 0.667 |
| WExT | 0.792 | 0.948 | 0.413 | 0.792 | 0.322 | 0.413 |

(f) Metrics for SKCM data.

| Method | Precision | Sensitivity | F1 Score | $\text{Precision}_{\text{strict}}$ | $\text{Sensitivity}_{\text{strict}}$ | $\text{F1 Score}_{\text{strict}}$ |
|---|---|---|---|---|---|---|
| DISCOVER | 0.812 | 0.961 | 0.575 | 0.812 | 0.375 | 0.575 |
| Fisher's Exact Test | 1.0 | 1.0 | 0.583 | 1.0 | 0.417 | 0.583 |
| WExT | 0.857 | 0.99 | 0.578 | 0.857 | 0.381 | 0.578 |

(g) Metrics for STAD data.

| Method | Precision | Sensitivity | F1 Score | $\text{Precision}_{\text{strict}}$ | $\text{Sensitivity}_{\text{strict}}$ | $\text{F1 Score}_{\text{strict}}$ |
|---|---|---|---|---|---|---|
| DISCOVER | 0.863 | 0.997 | 0.317 | 0.857 | 0.018 | 0.035 |
| Fisher's Exact Test | 0.889 | 1.000 | 0.331 | 1.000 | 0.002 | 0.004 |
| WExT | 0.887 | 0.991 | 0.419 | 0.892 | 0.058 | 0.109 |

(h) Metrics for UCEC data.

| Method | Precision | Sensitivity | F1 Score | $\text{Precision}_{\text{strict}}$ | $\text{Sensitivity}_{\text{strict}}$ | $\text{F1 Score}_{\text{strict}}$ |
|---|---|---|---|---|---|---|
| DISCOVER | 0.889 | 0.888 | 0.466 | 0.902 | 0.359 | 0.465 |
| Fisher's Exact Test | 0.952 | 0.988 | 0.211 | 0.952 | 0.127 | 0.211 |
| WExT | 0.953 | 0.984 | 0.274 | 0.953 | 0.179 | 0.274 |

### Scatterplots of percentage significance of mutual exclusivity runs vs mutation load association (MLA) when $t = 20$

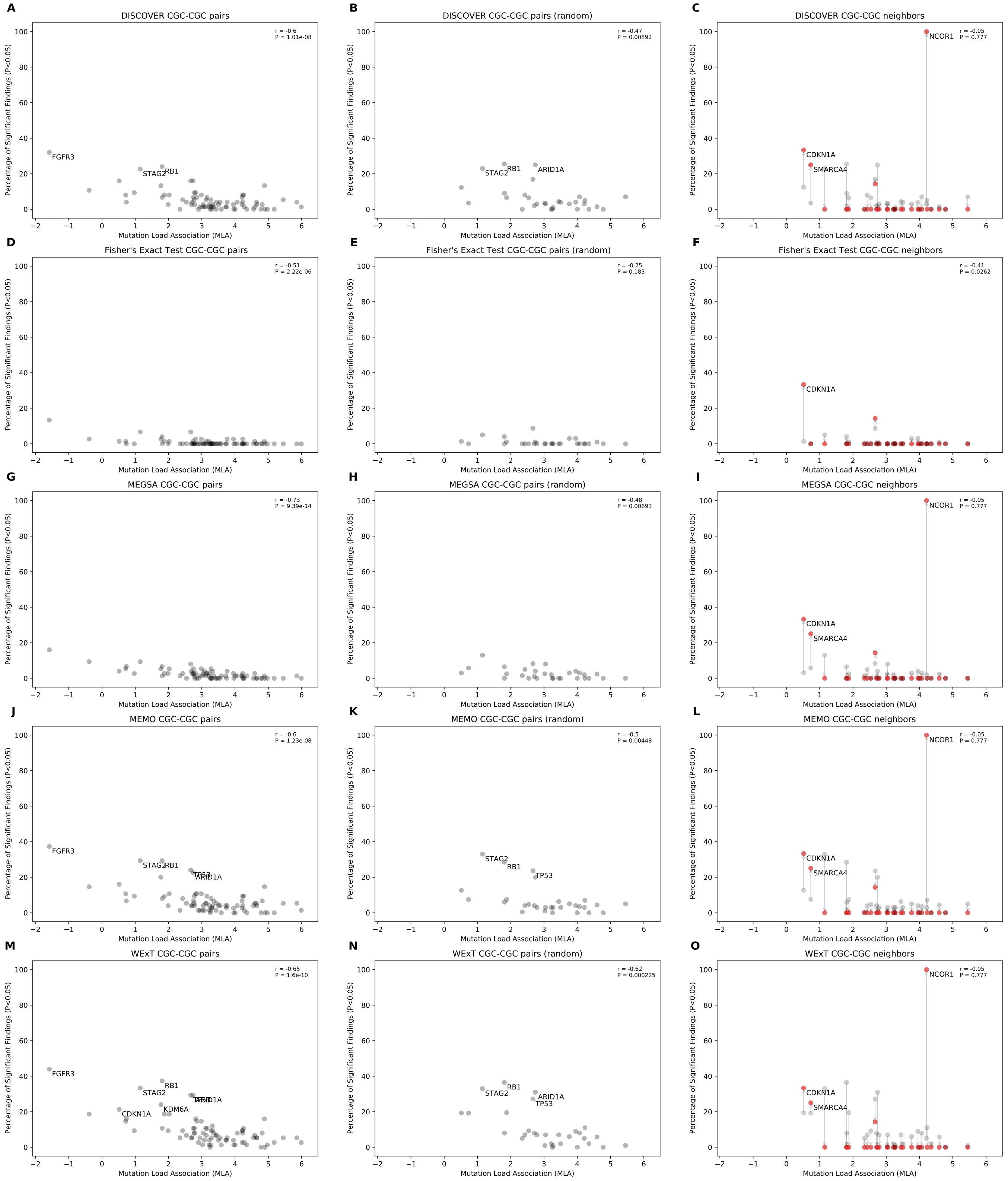


Fig 1: Comparison of mutual exclusivity results of DISCOVER, Fisher's Exact Test, MEGSA, MEMo and WExT on BLCA cohort with $t = 20$ (411 samples)

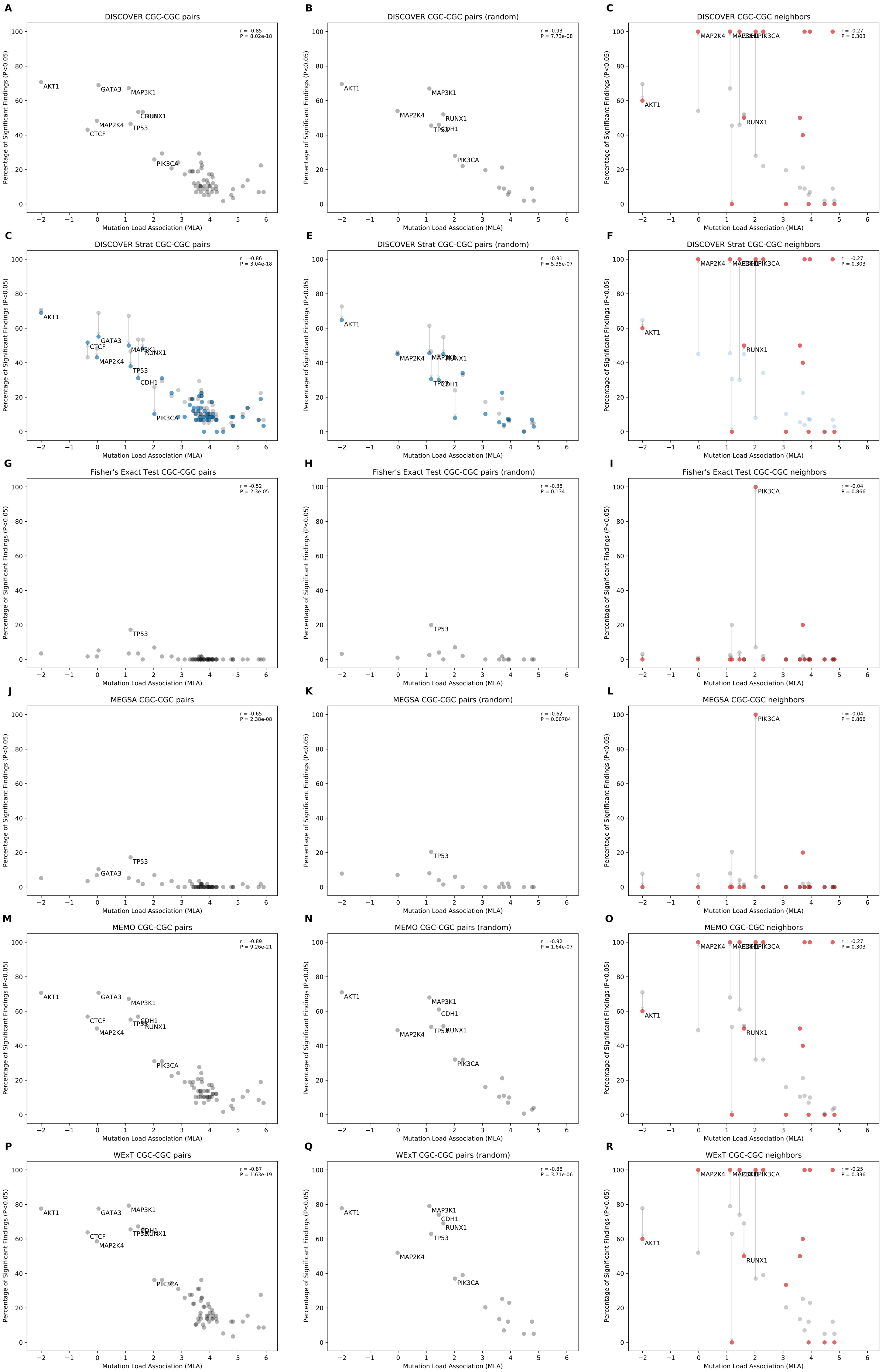


Fig 2: Comparison of mutual exclusivity results of DISCOVER, DISCOVER Strat, Fisher's Exact Test, MEGSA, MEMo and WExT on BRCA cohort with $t = 20$ (1026 samples)

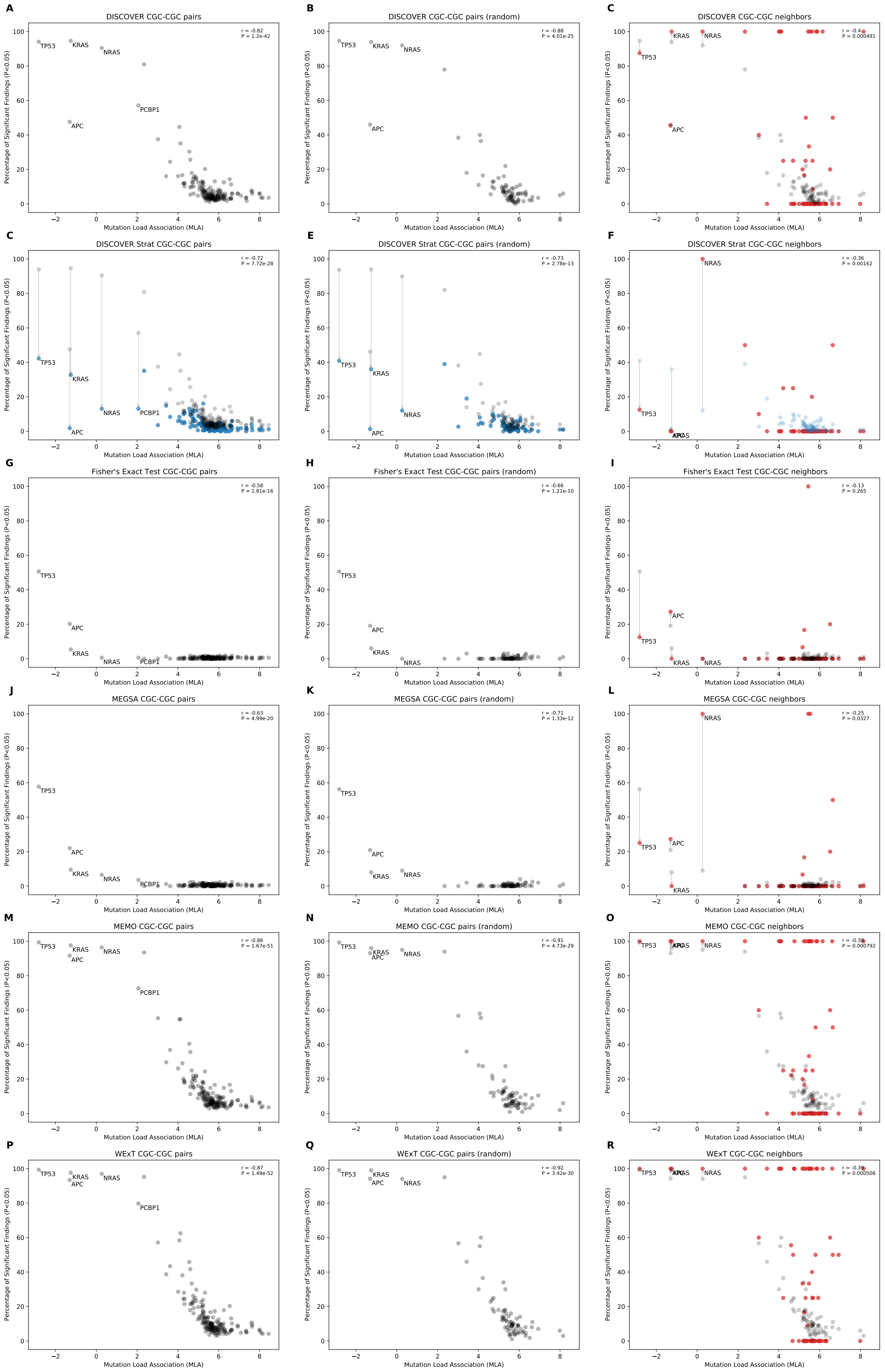


Fig 3: Comparison of mutual exclusivity results of DISCOVER, DISCOVER Strat, Fisher's Exact Test, MEGSA, MEMo and WExT on COADREAD cohort with $t = 20$ (498 samples)

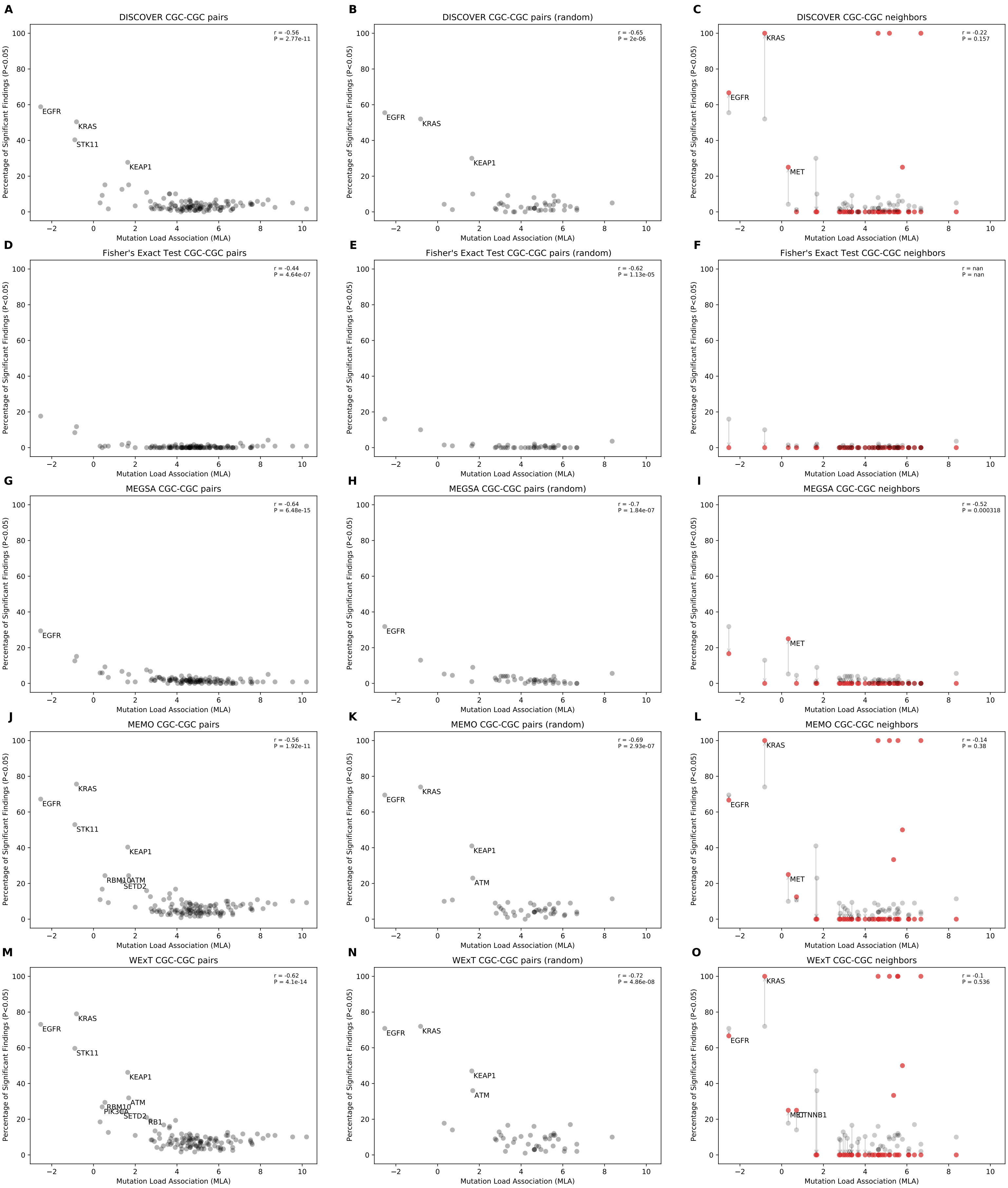


Fig 4: Comparison of mutual exclusivity results of DISCOVER, Fisher's Exact Test, MEGSA, MEMo and WExT on LUAD cohort with $t = 20$ (568 samples)

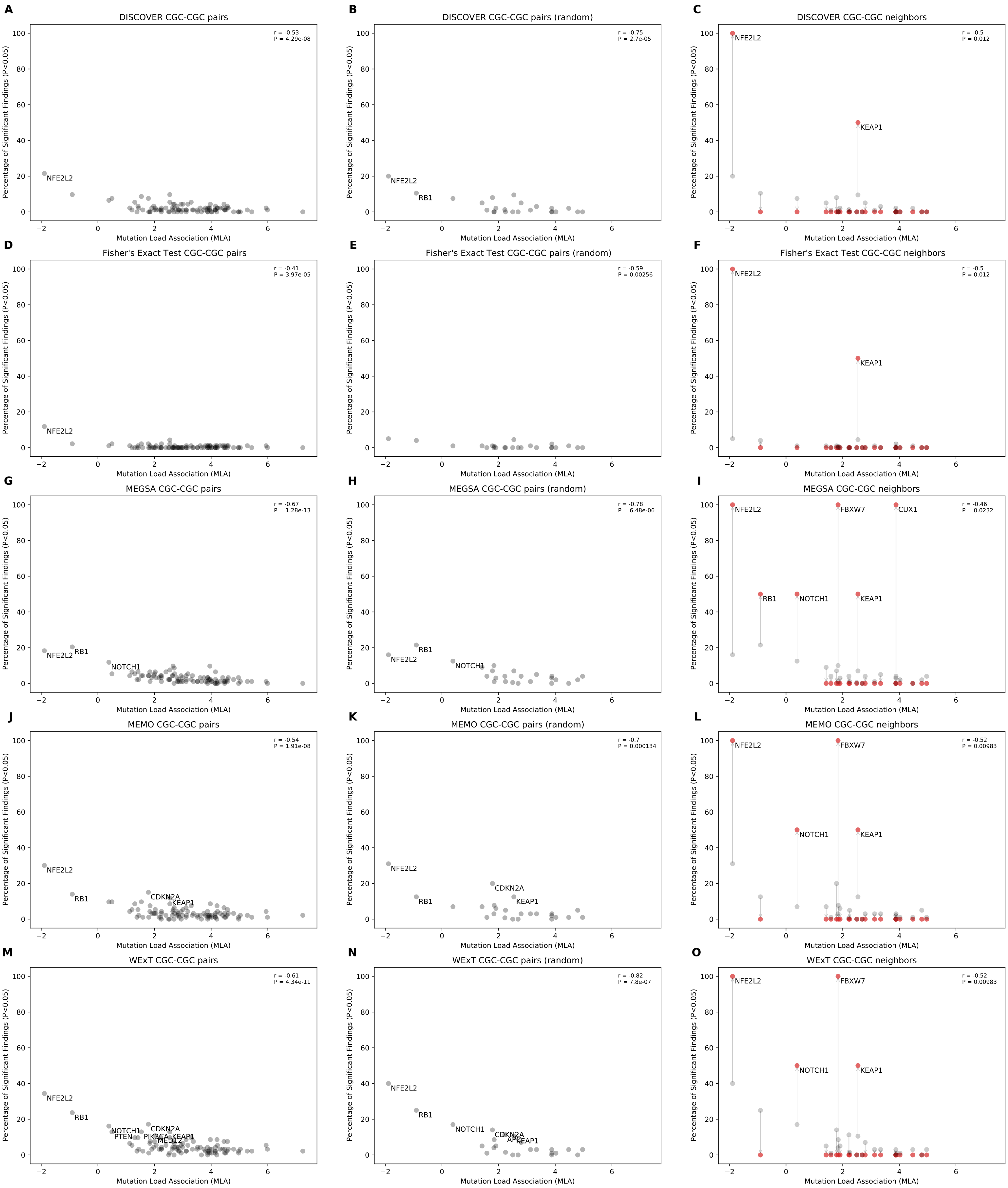


Fig 5: Comparison of mutual exclusivity results of DISCOVER, Fisher's Exact Test, MEGSA, MEMo and WExT on LUSC cohort with $t = 20$ (485 samples)

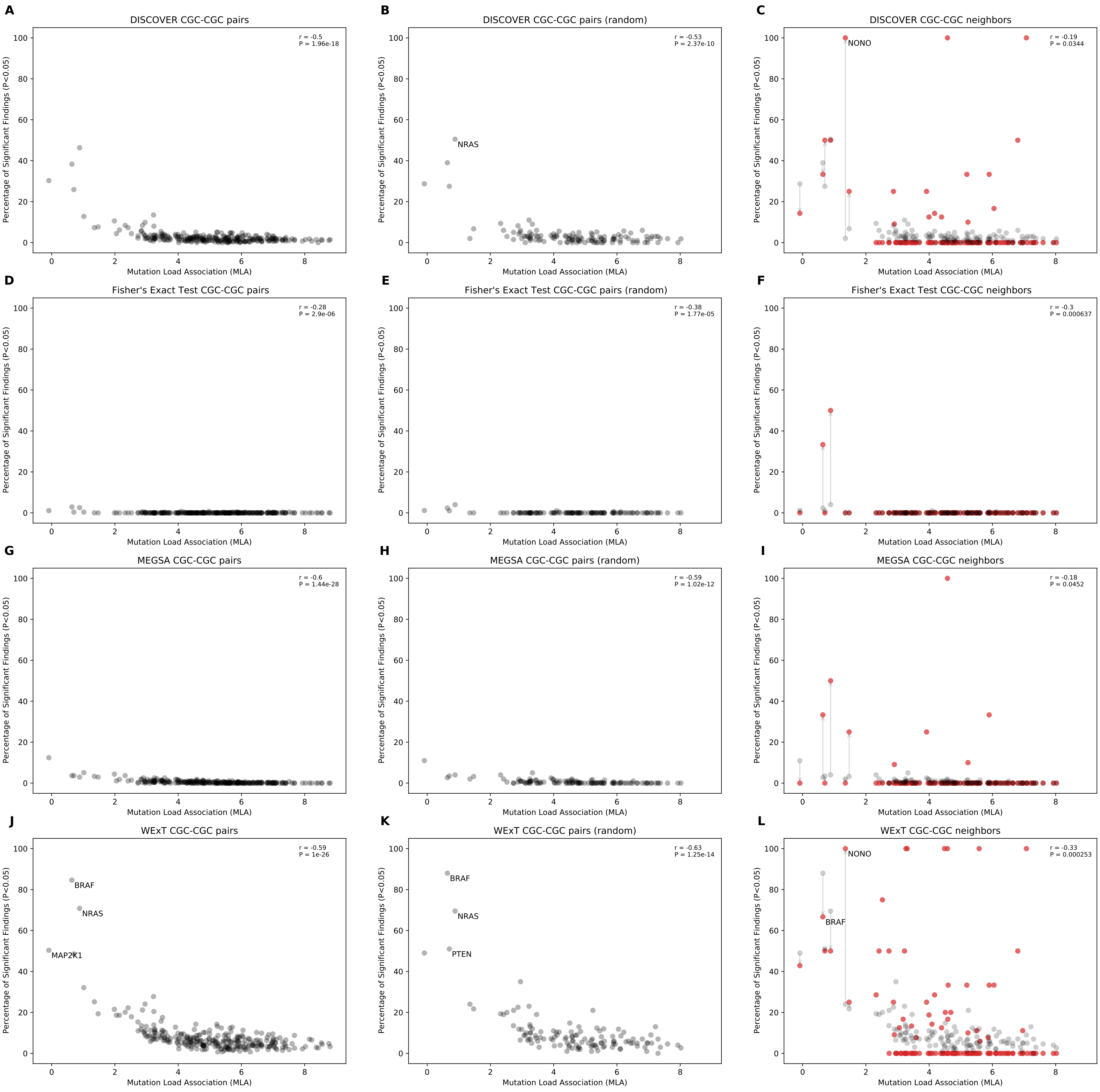


Fig 6: Comparison of mutual exclusivity results of DISCOVER, Fisher's Exact Test, MEGSA and WExT on SKCM cohort with $t = 20$ (468 samples)

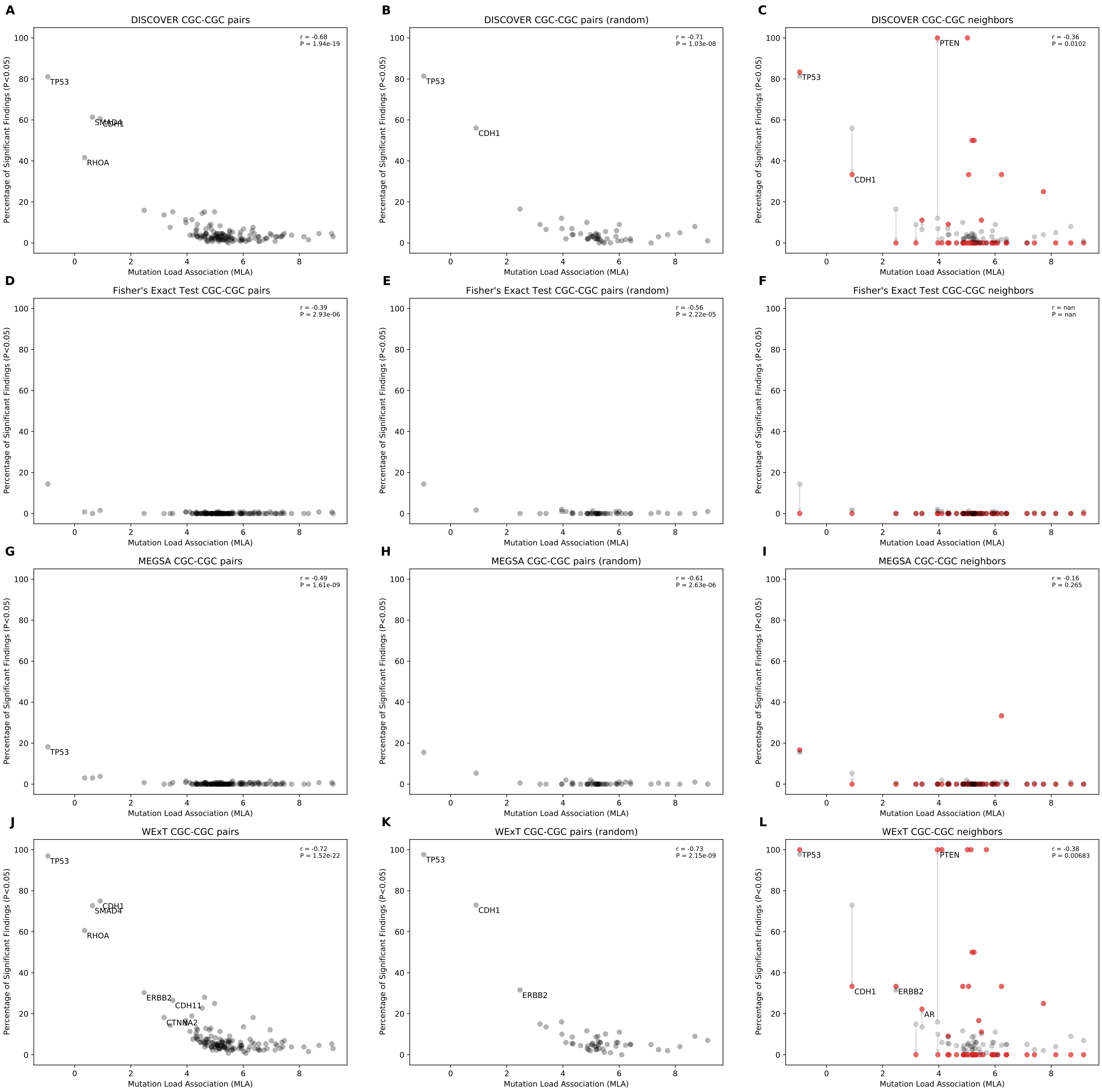


Fig 7: Comparison of mutual exclusivity results of DISCOVER, Fisher's Exact Test, MEGSA and WExT on STAD cohort with $t = 20$ (438 samples)

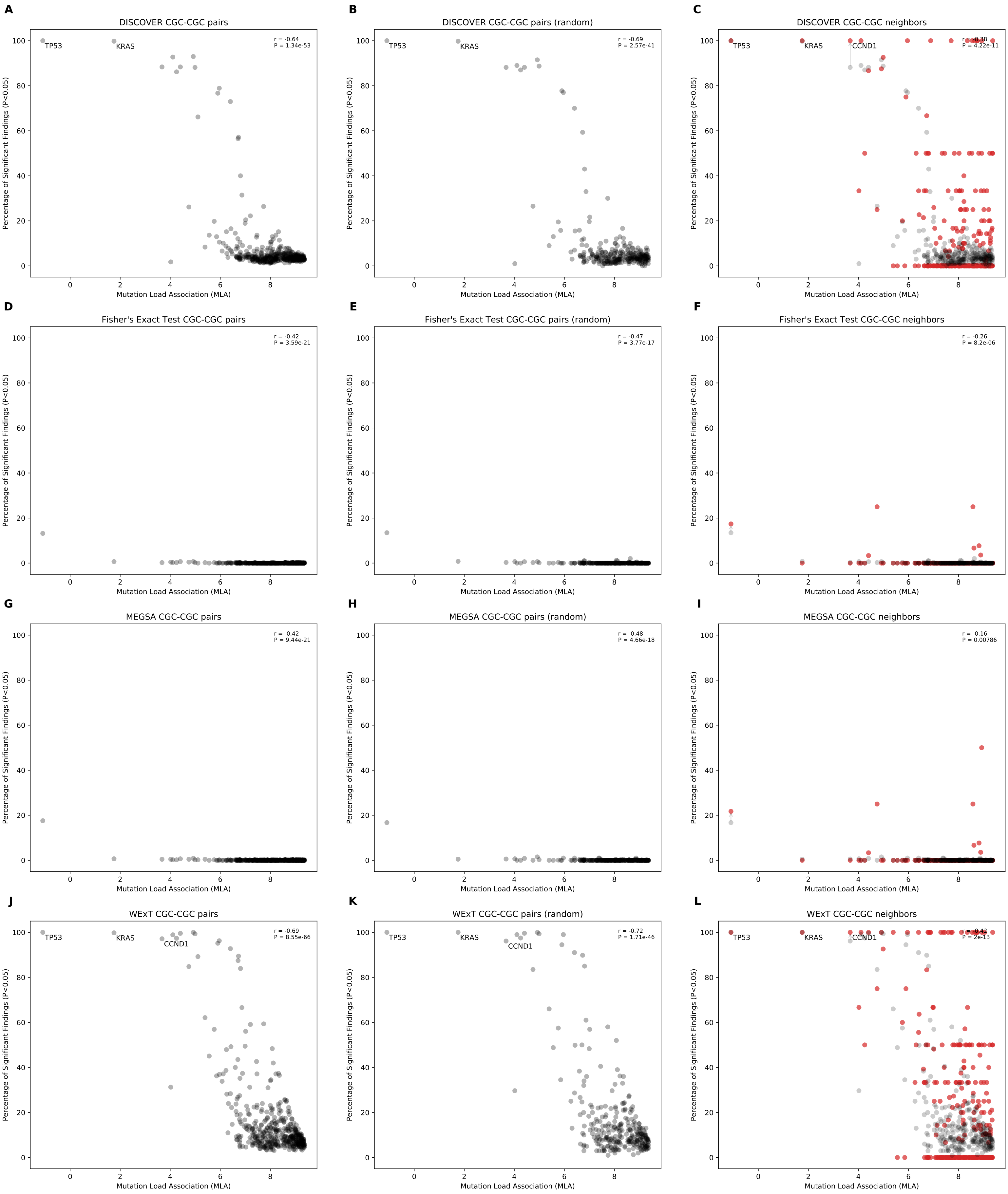


Fig 8: Comparison of mutual exclusivity results of DISCOVER, Fisher's Exact Test, MEGSA and WExT on UCEC cohort with $t = 20$ (531 samples)

### Scatterplots of percentage significance of mutual exclusivity runs vs mutation load association (MLA) when $t = 5$

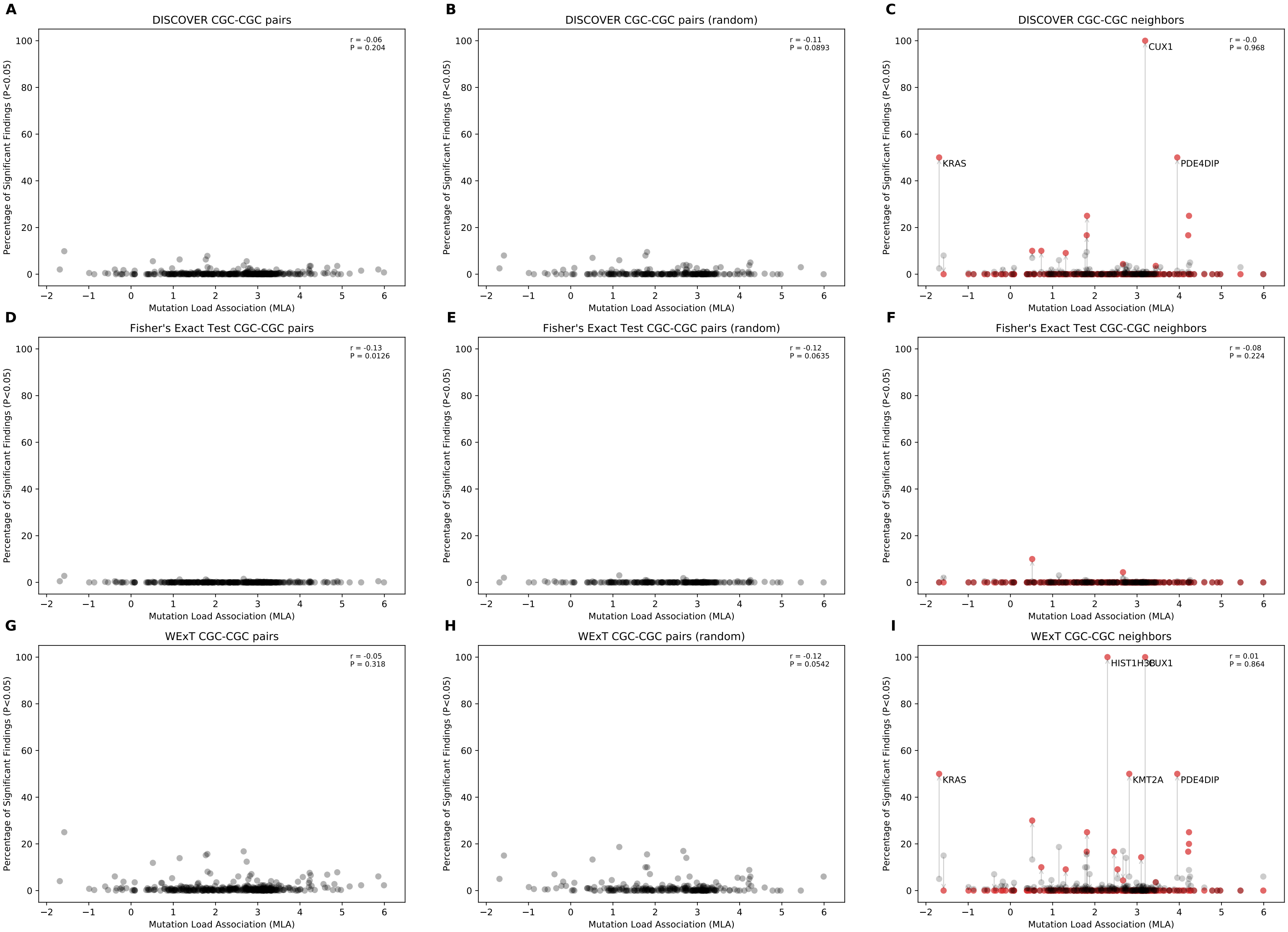


Fig 9: Comparison of mutual exclusivity results of DISCOVER, Fisher's Exact Test and WExT on BLCA cohort with $t = 5$ (411 samples)

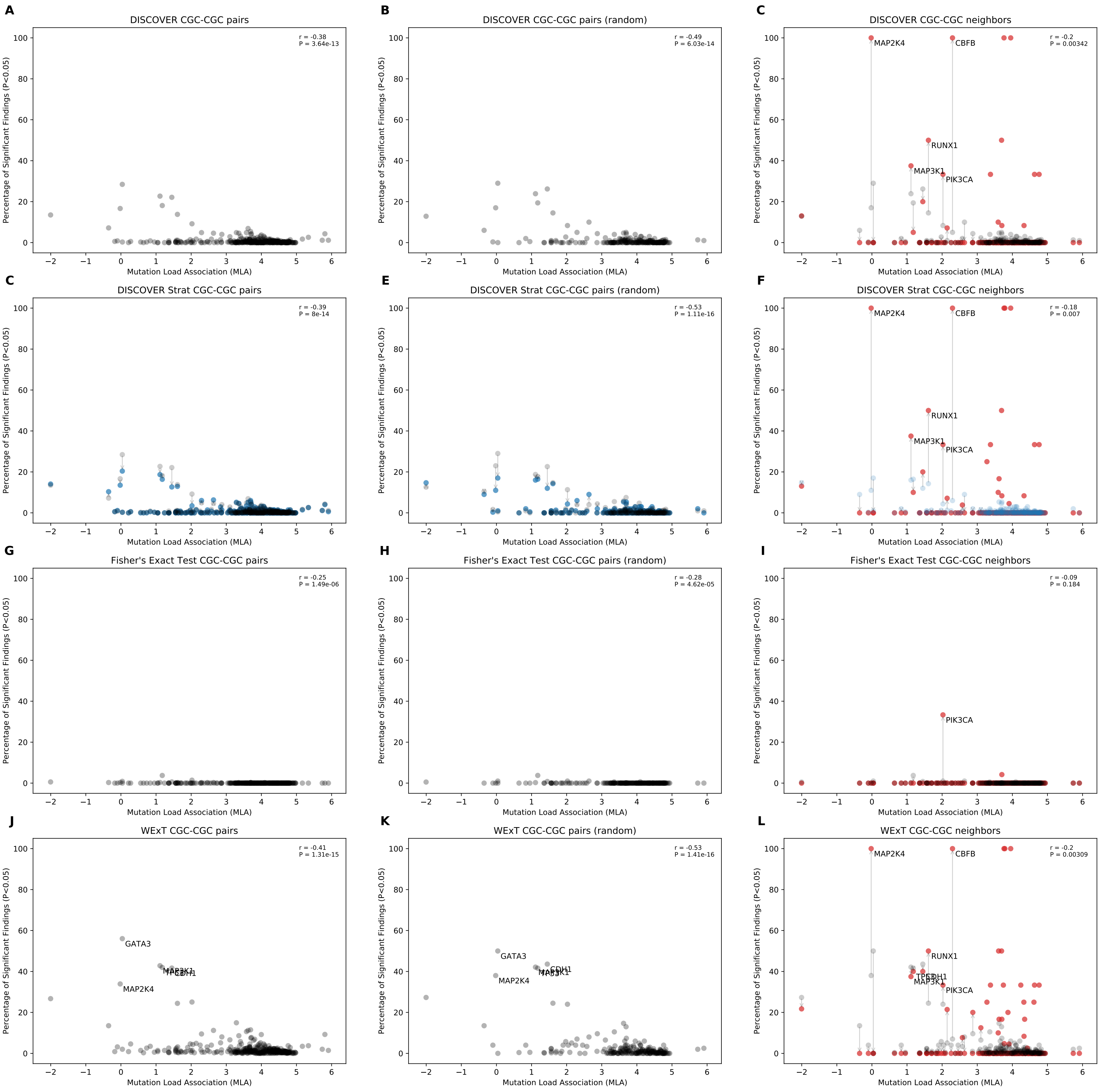


Fig 10: Comparison of mutual exclusivity results of DISCOVER, DISCOVER Strat, Fisher's Exact Test and WExT on BRCA cohort with $t = 5$ (1026 samples)

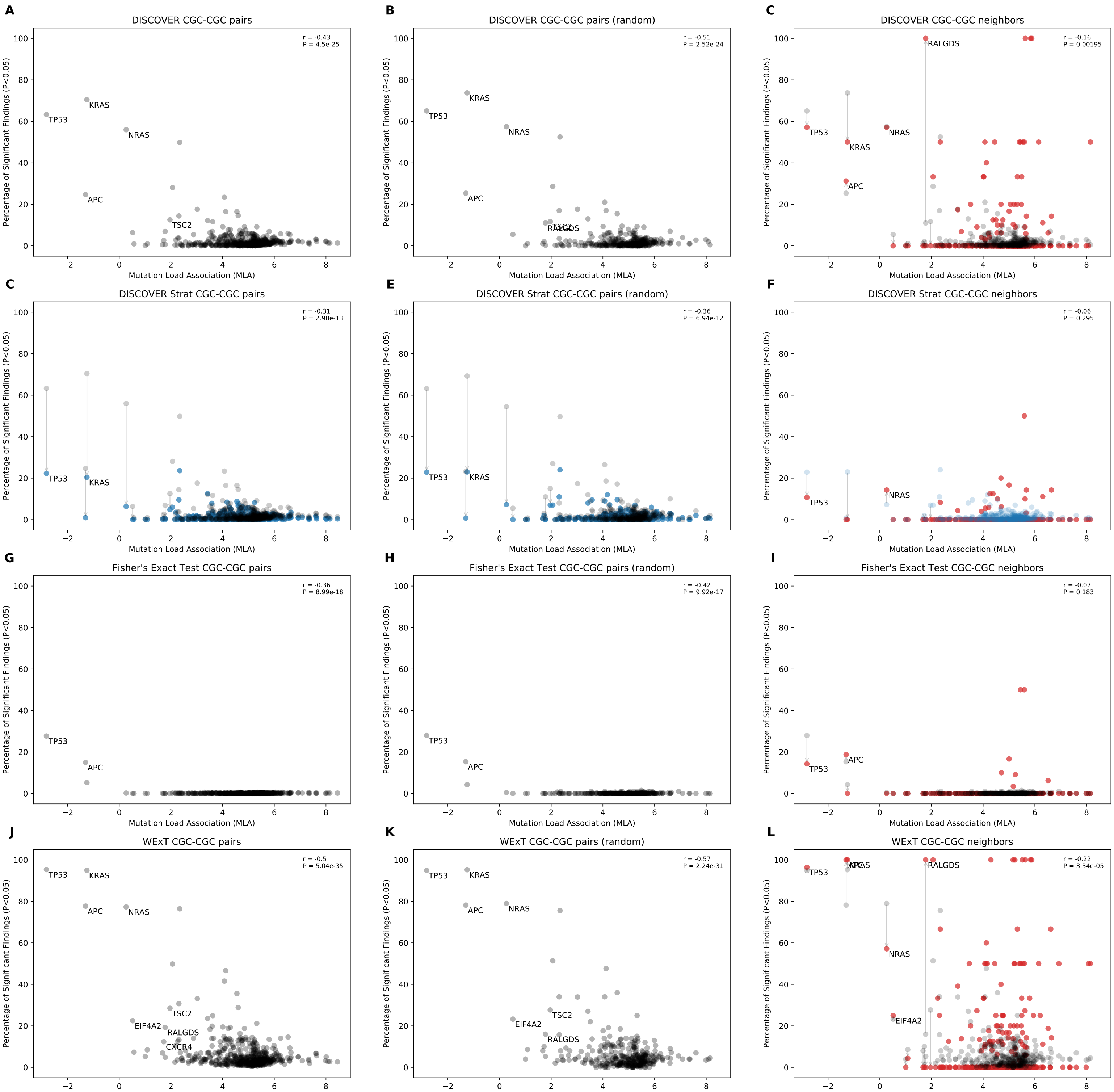


Fig 11: Comparison of mutual exclusivity results of DISCOVER, DISCOVER Strat, Fisher's Exact Test and WExT on COADREAD cohort with $t = 5$ (498 samples)

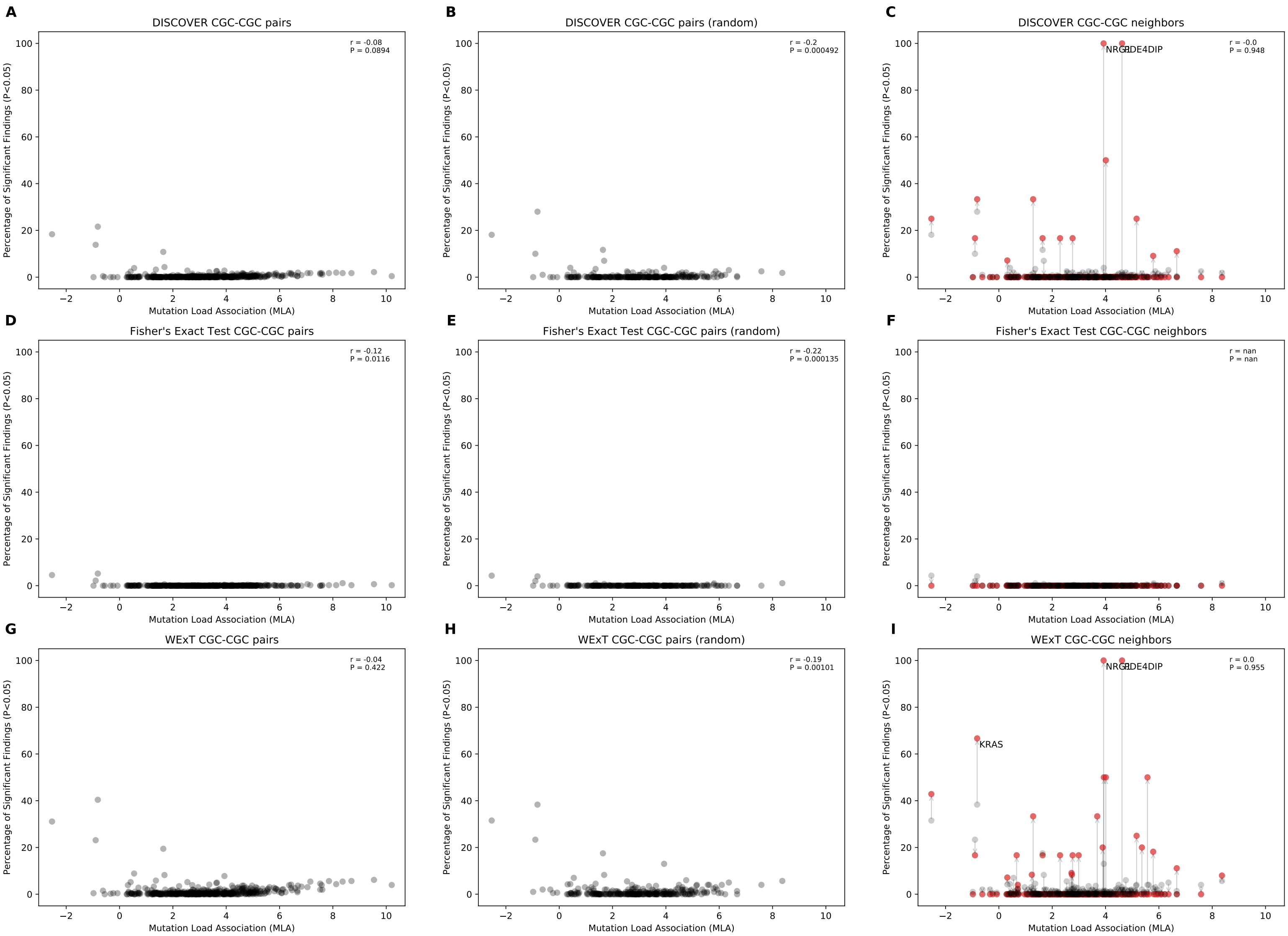


Fig 12: Comparison of mutual exclusivity results of DISCOVER, Fisher's Exact Test and WExT on LUAD cohort with $t = 5$ (568 samples)

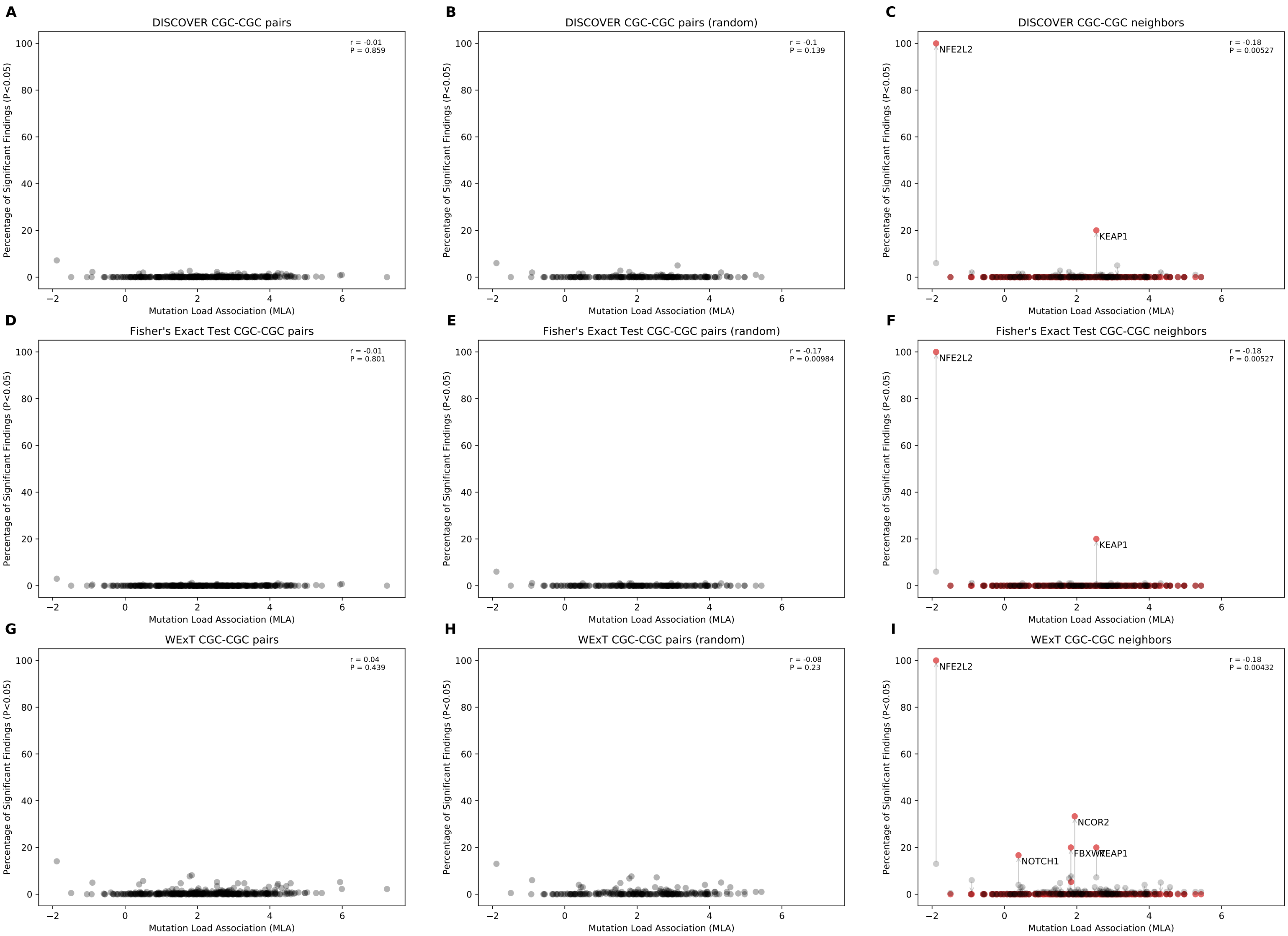


Fig 13: Comparison of mutual exclusivity results of DISCOVER, Fisher's Exact Test and WExT on LUSC cohort with $t = 5$ (485 samples)

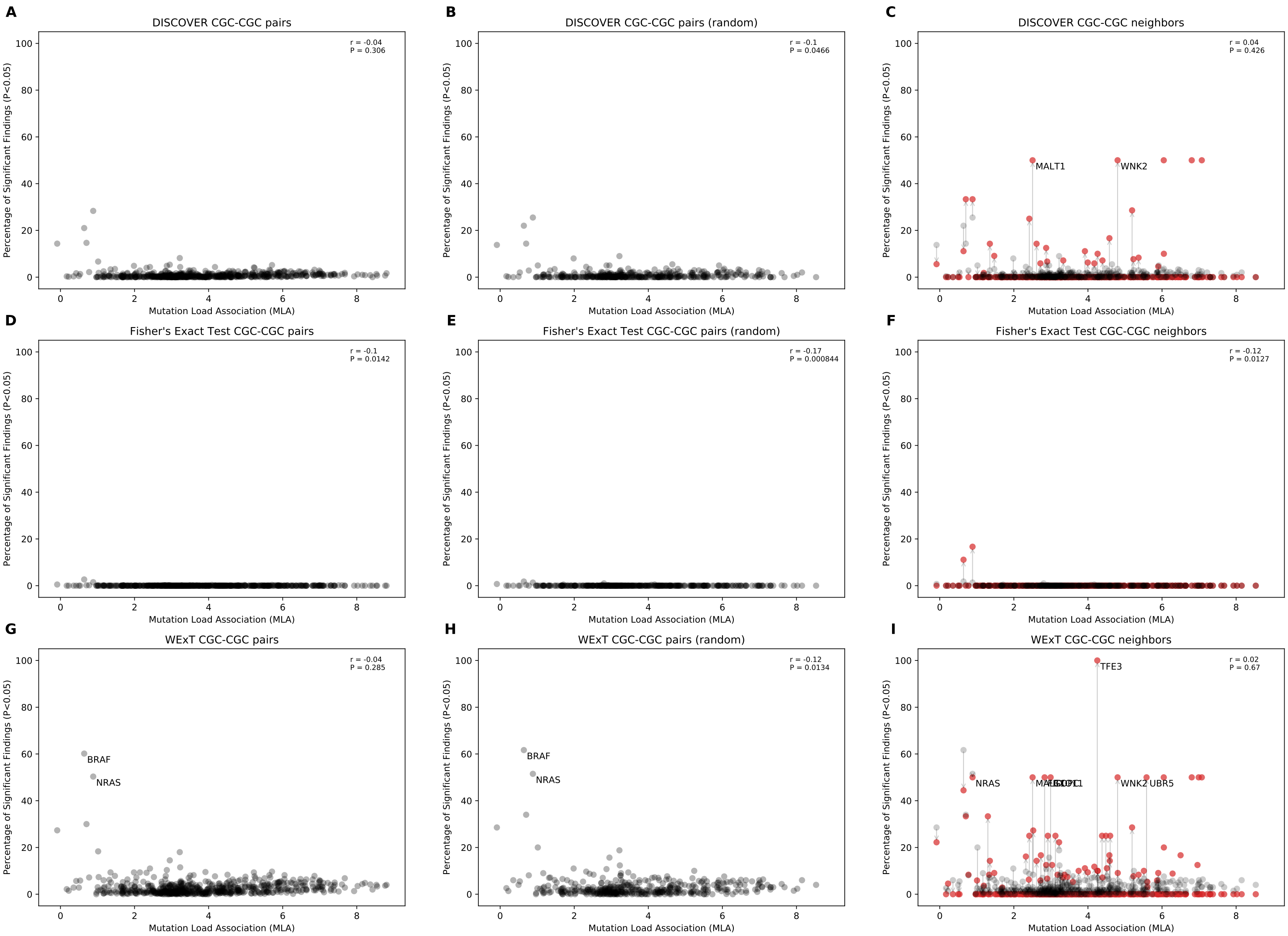


Fig 14: Comparison of mutual exclusivity results of DISCOVER, Fisher's Exact Test and WExT on SKCM cohort with $t = 5$ (468 samples)

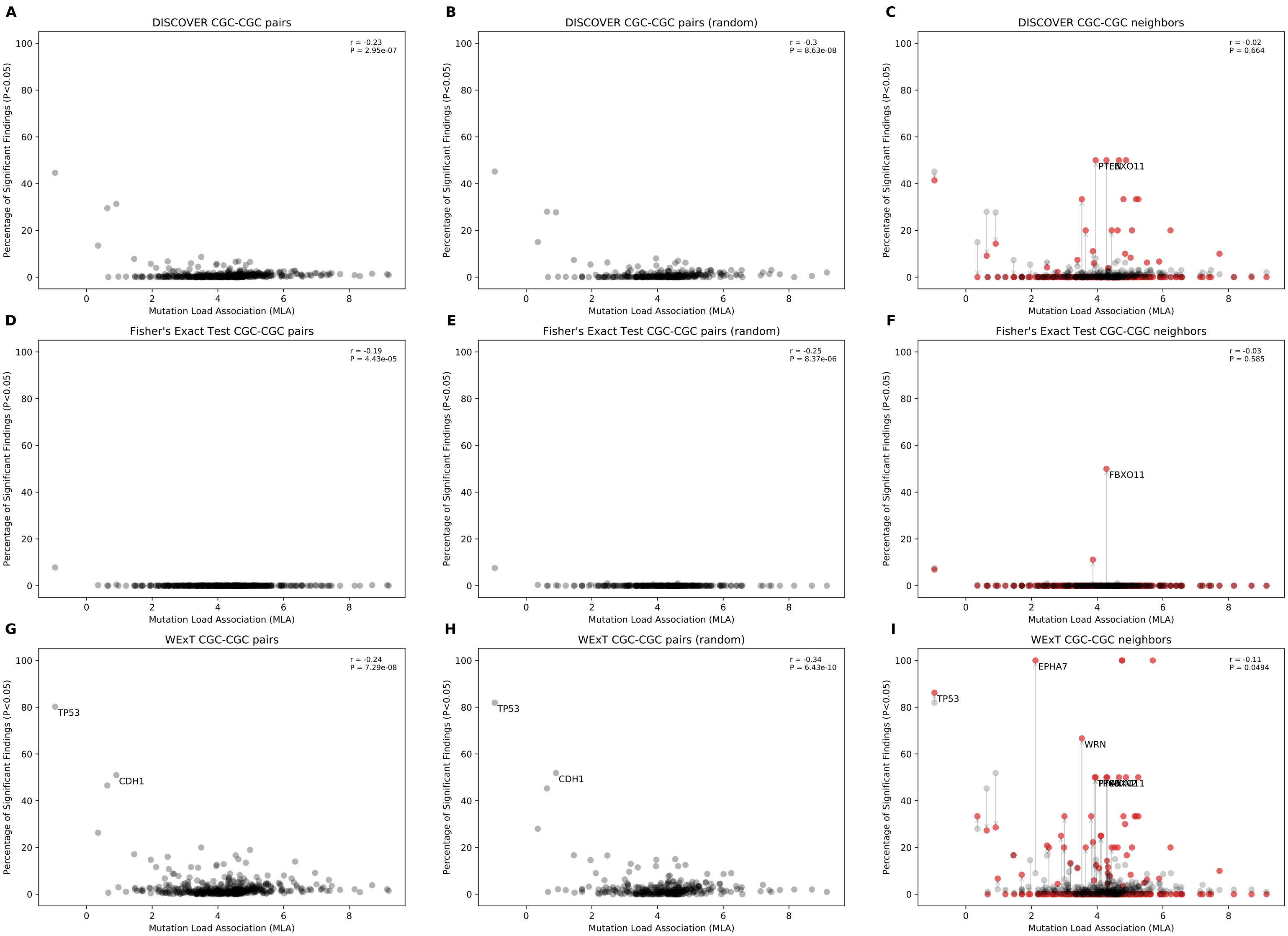


Fig 15: Comparison of mutual exclusivity results of DISCOVER, Fisher's Exact Test and WExT on STAD cohort with $t = 5$ (438 samples)

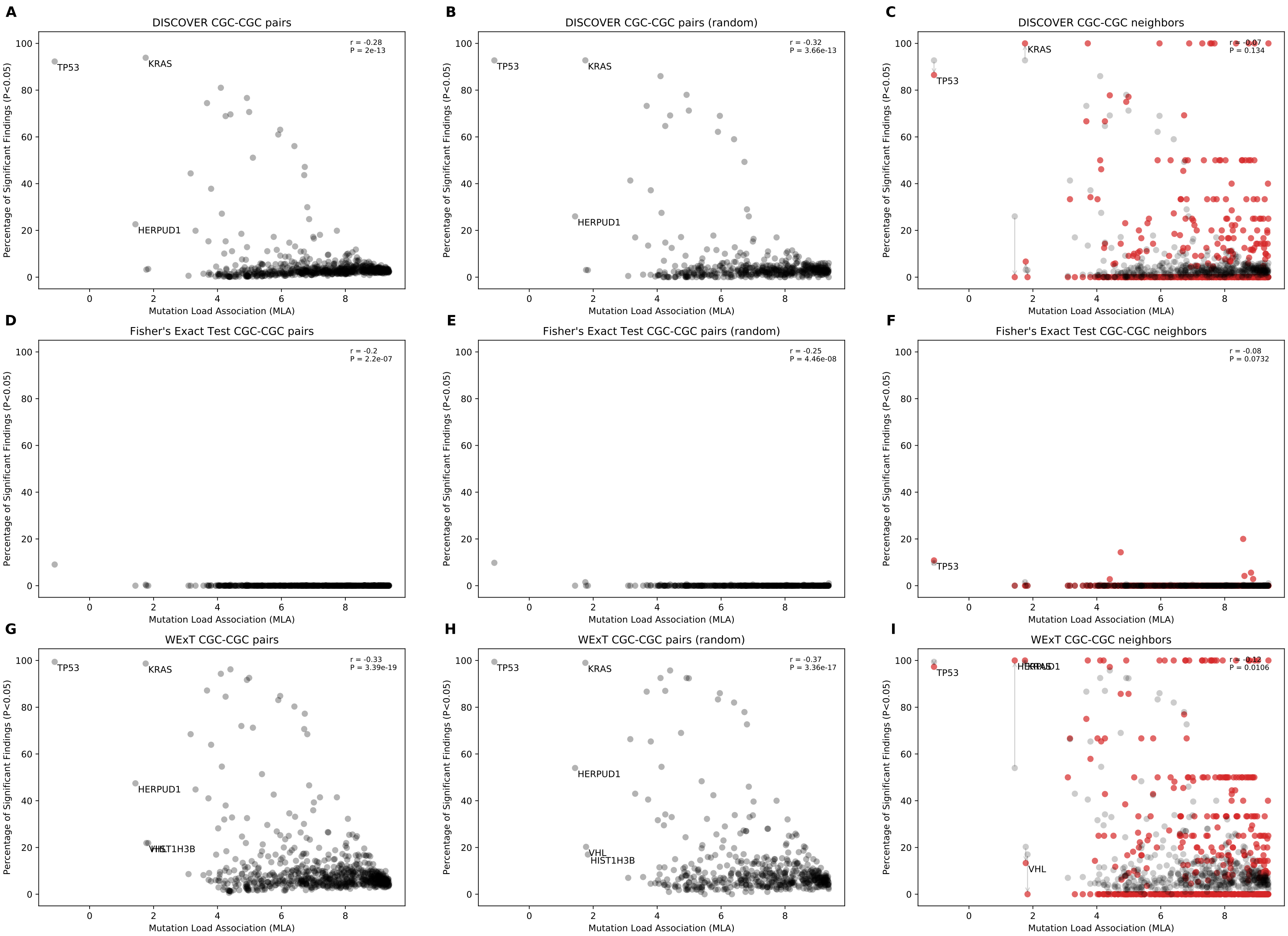


Fig 16: Comparison of mutual exclusivity results of DISCOVER, Fisher's Exact Test and WExT on UCEC cohort with $t = 5$ (531 samples)

Scatterplots of percentage significance of mutual exclusivity runs vs mutation load association (MLA) when only CGC genes that have $> 1$ neighbors are included ($t = 20$)

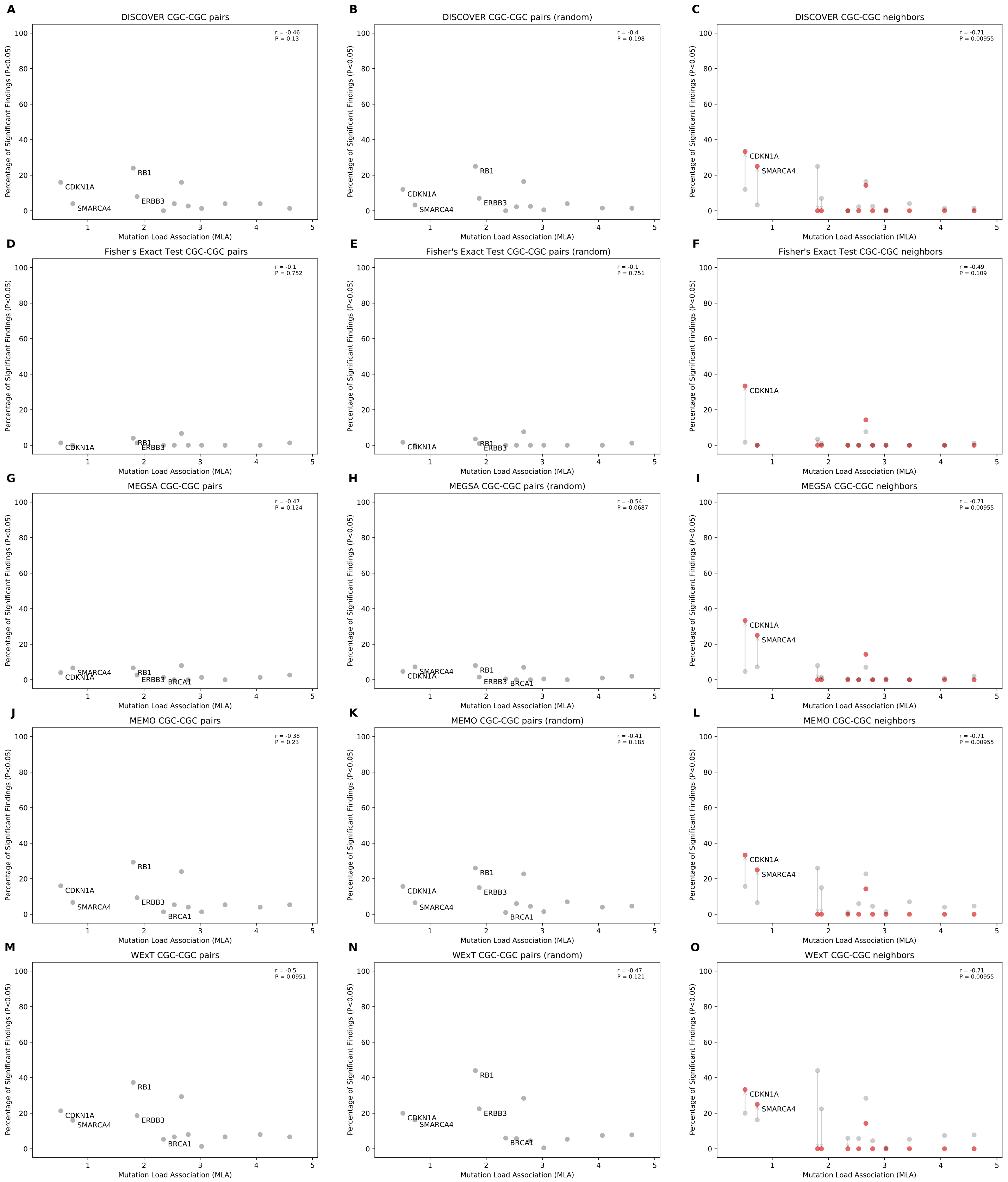


Fig 17: Comparison of mutual exclusivity results of DISCOVER, Fisher's Exact Test, MEGSA, MEMo and WExT on BLCA cohort with $t = 20$ for CGC genes that have $> 1$ neighbors (411 samples)

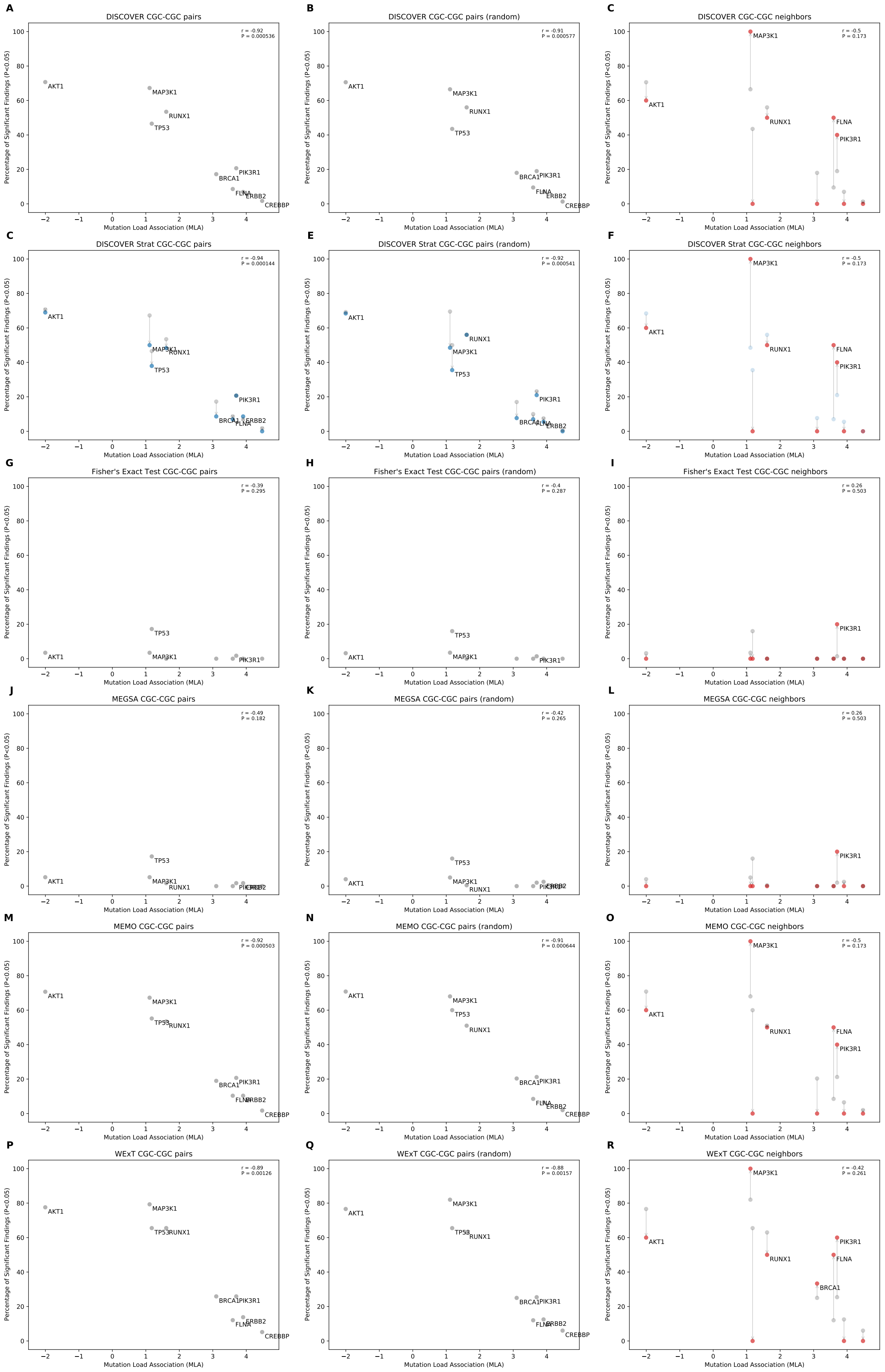


Fig 18: Comparison of mutual exclusivity results of DISCOVER, DISCOVER Strat, Fisher's Exact Test, MEGSA, MEMo and WExT on BRCA cohort with $t = 20$ for CGC genes that have $> 1$ neighbors (1026 samples)

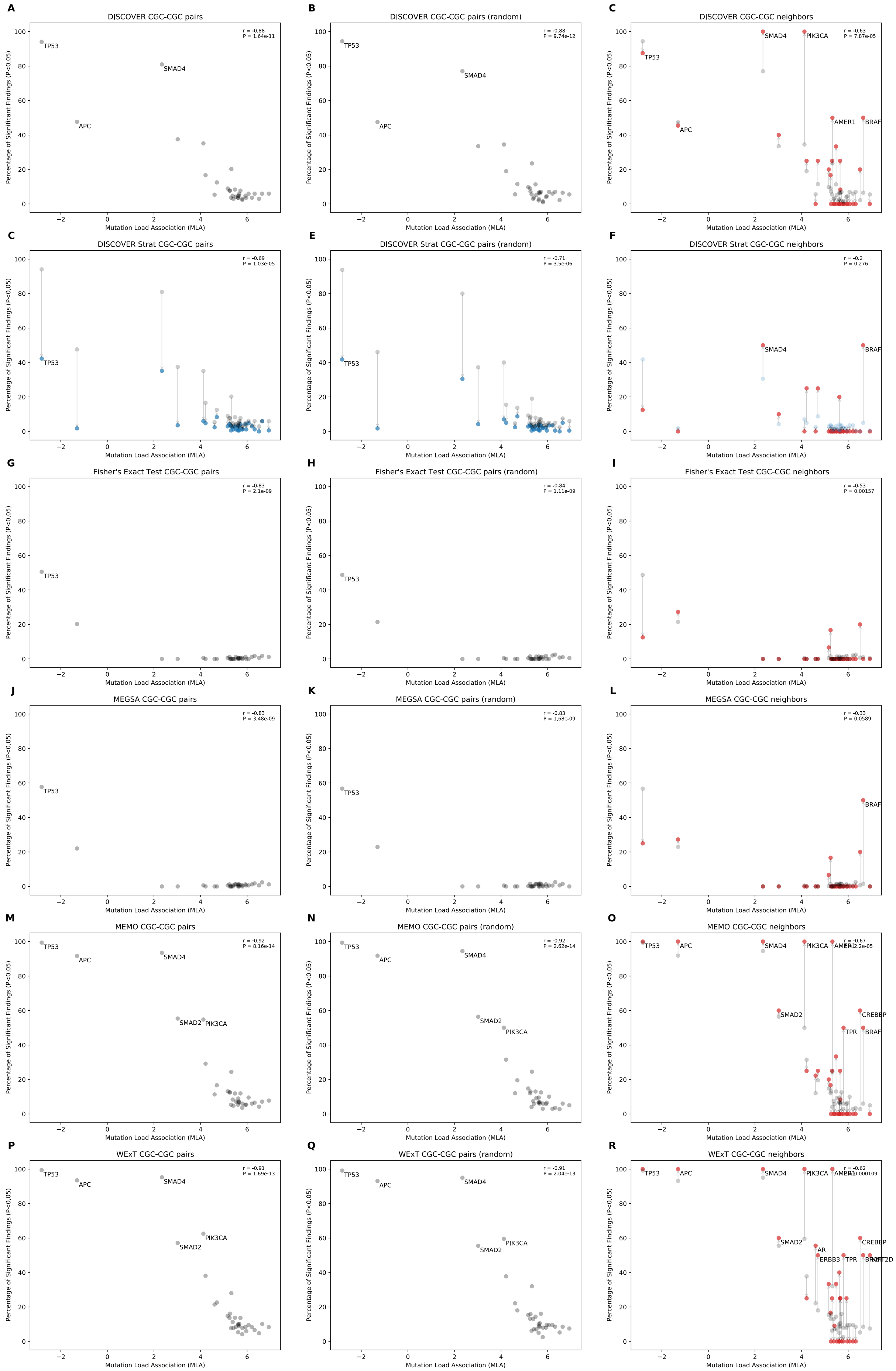

Fig 19: Comparison of mutual exclusivity results of DISCOVER, DISCOVER Strat, Fisher's Exact Test, MEGSA, MEMo and WExT on COADREAD cohort with $t = 20$ for CGC genes that have $> 1$ neighbors (498 samples)

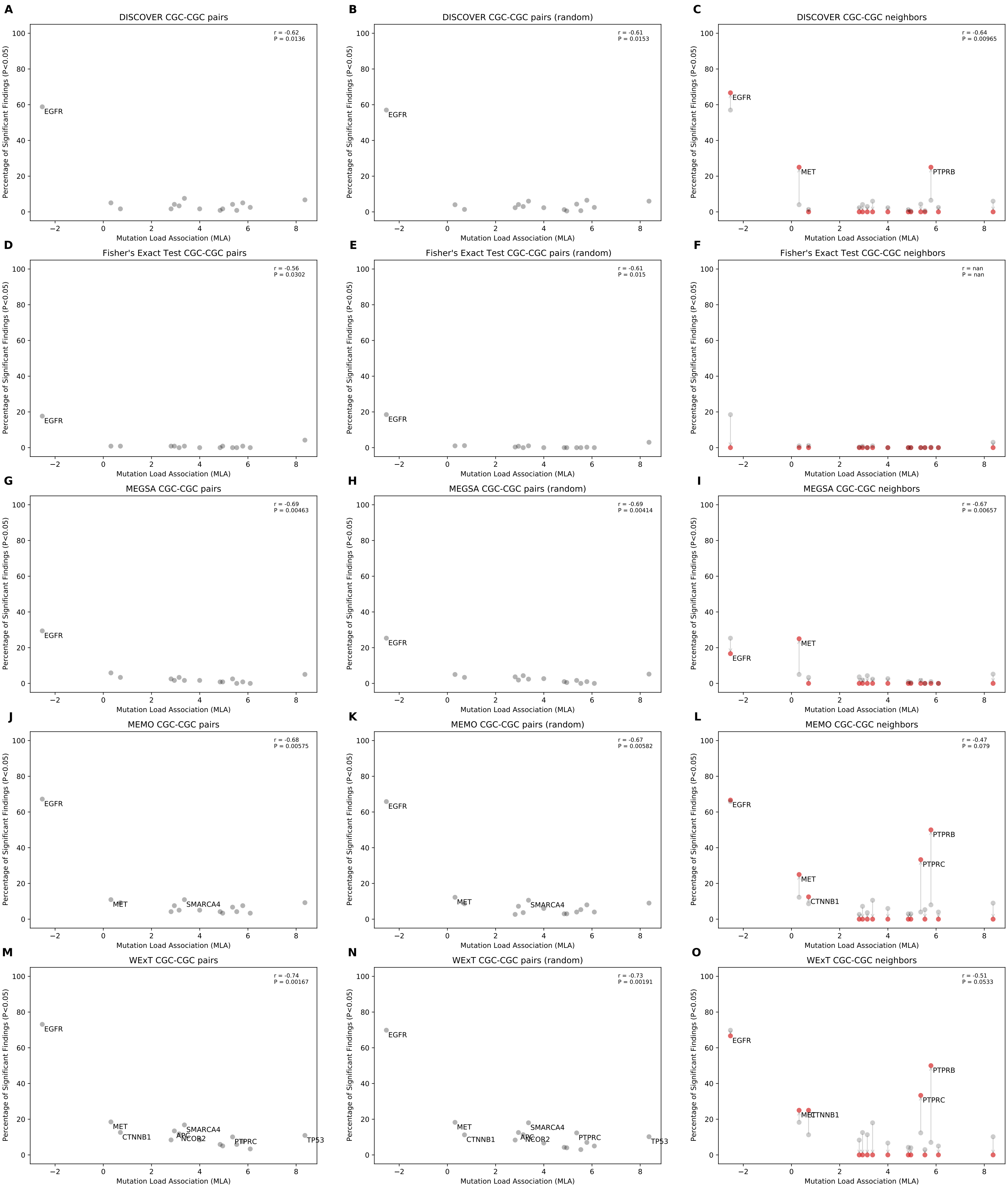


Fig 20: Comparison of mutual exclusivity results of DISCOVER, Fisher's Exact Test, MEGSA, MEMo and WExT on LUAD cohort with $t = 20$ for CGC genes that have $> 1$ neighbors (568 samples)

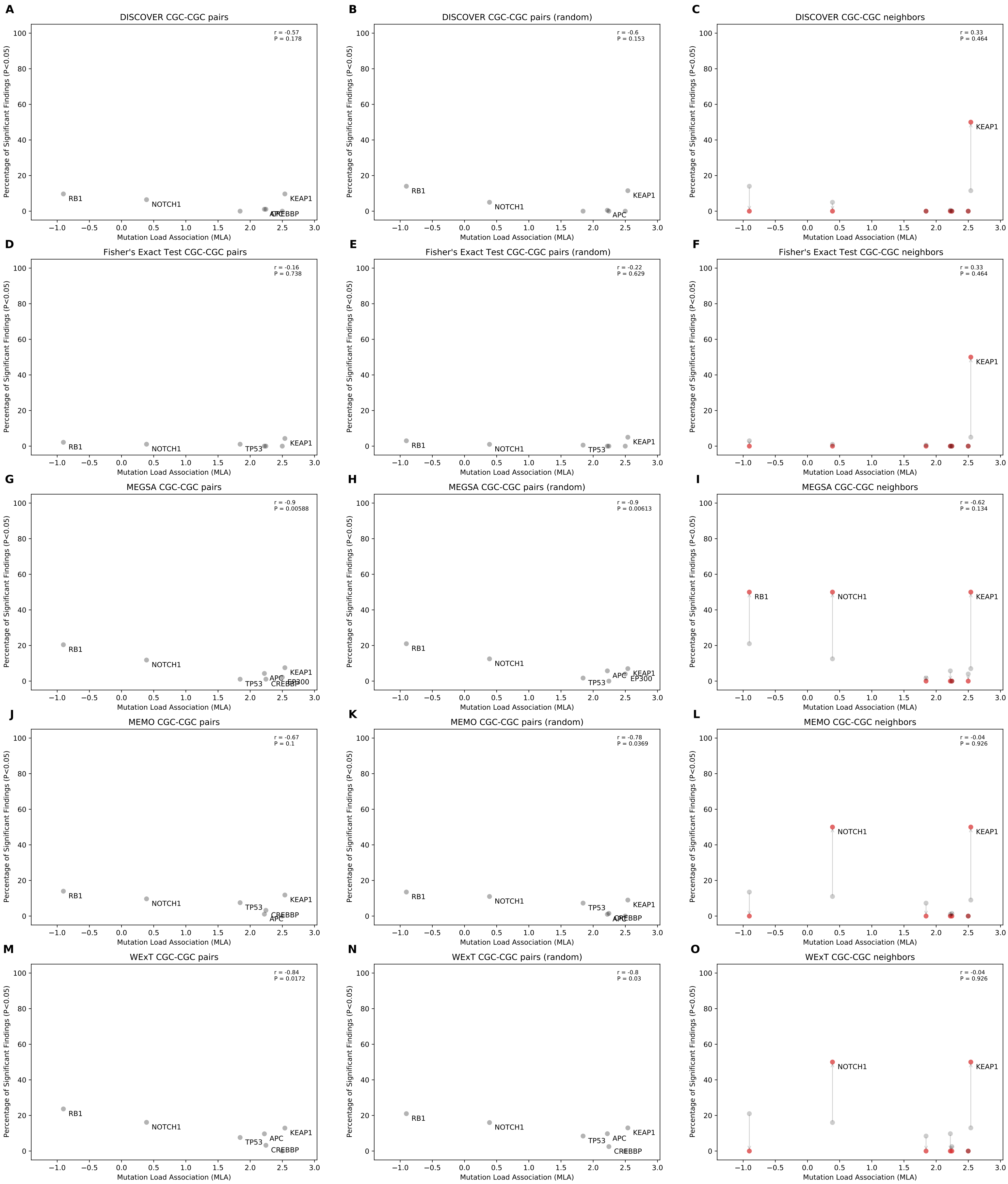


Fig 21: Comparison of mutual exclusivity results of DISCOVER, Fisher's Exact Test, MEGSA, MEMo and WExT on LUSC cohort with $t = 20$ for CGC genes that have $> 1$ neighbors (485 samples)

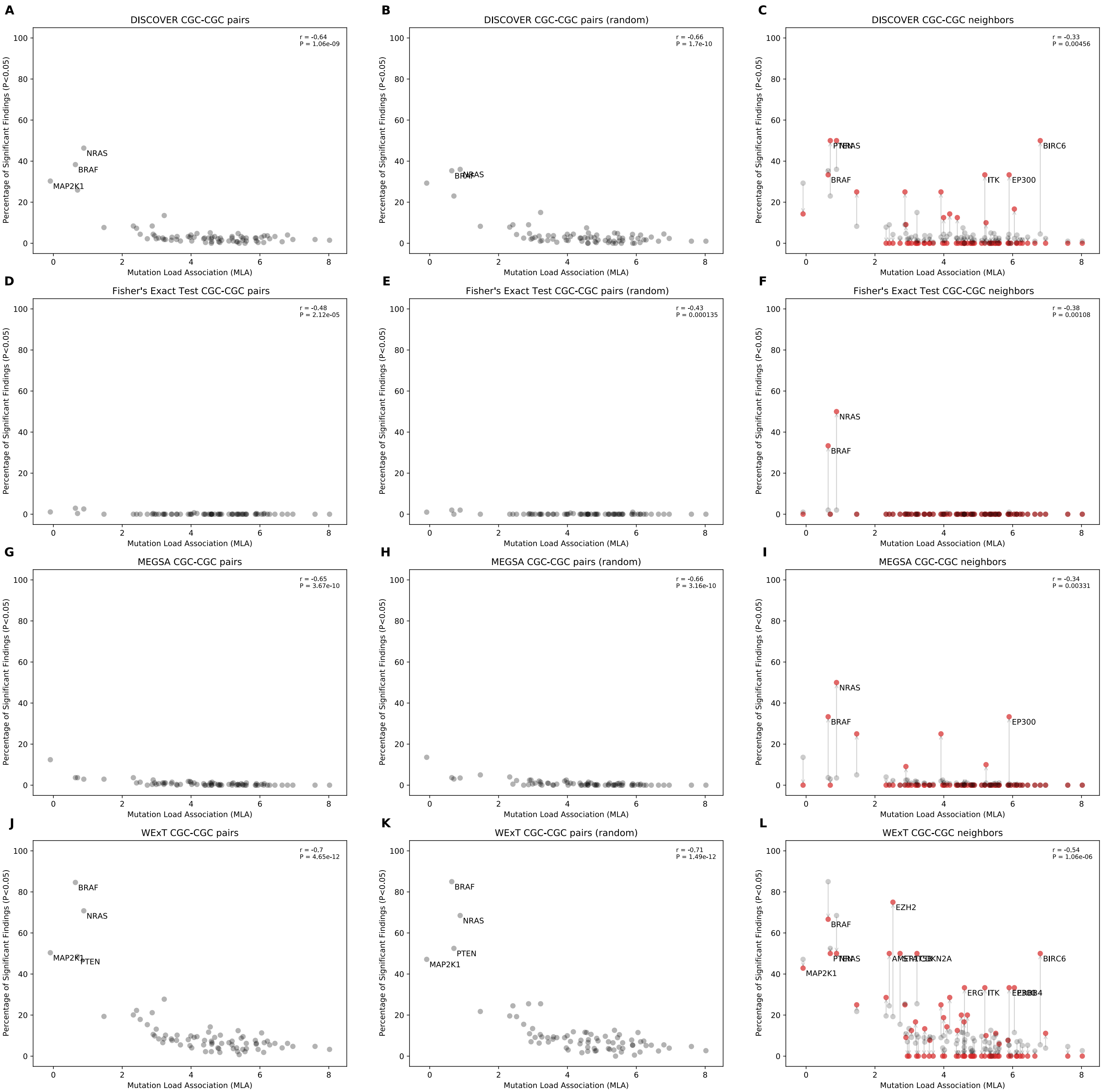


Fig 22: Comparison of mutual exclusivity results of DISCOVER, Fisher's Exact Test, MEGSA and WExT on SKCM cohort with $t = 20$ for CGC genes that have $> 1$ neighbors (468 samples)

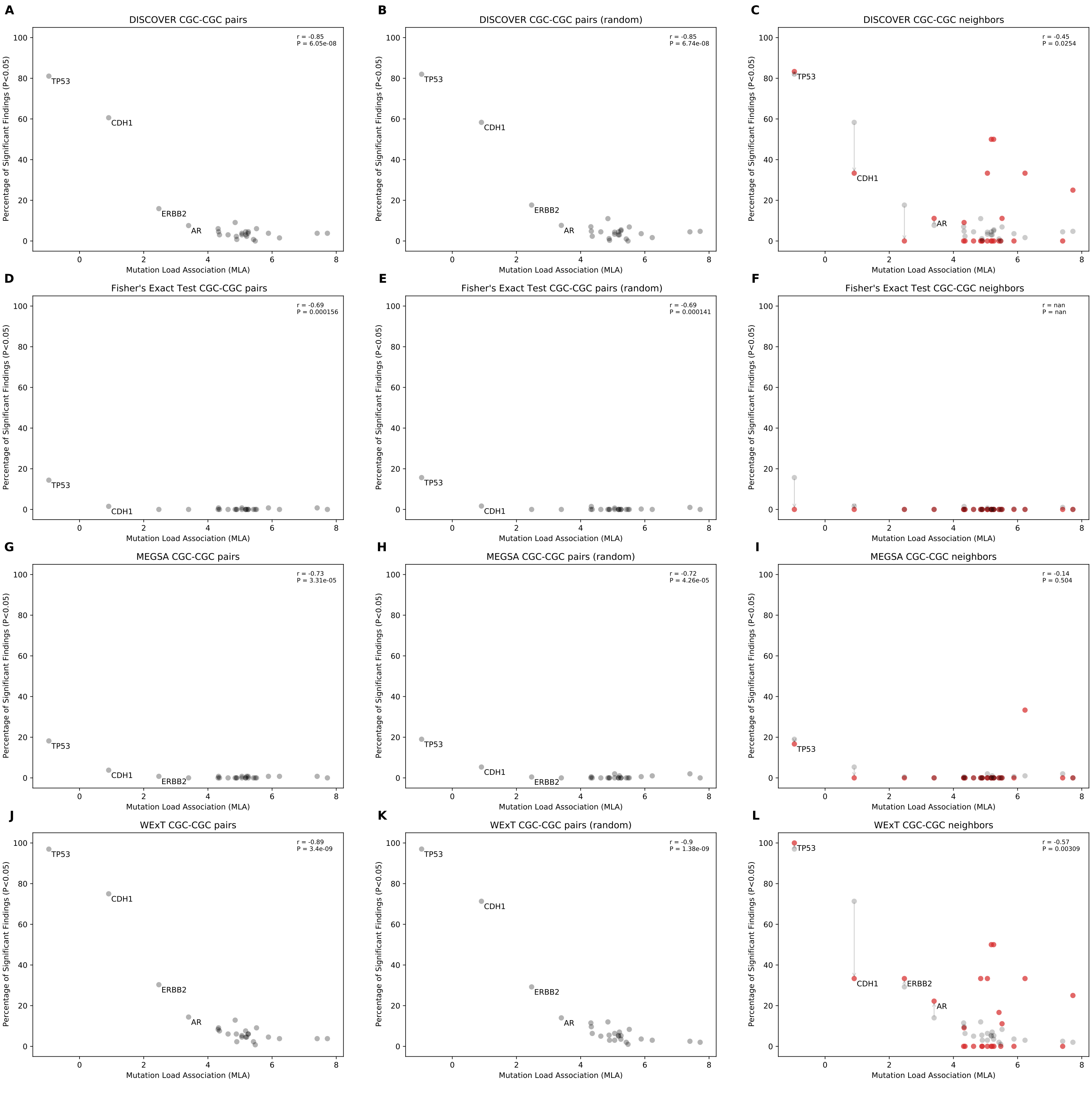


Fig 23: Comparison of mutual exclusivity results of DISCOVER, Fisher's Exact Test, MEGSA and WExT on STAD cohort with $t = 20$ for CGC genes that have $> 1$ neighbors (438 samples)

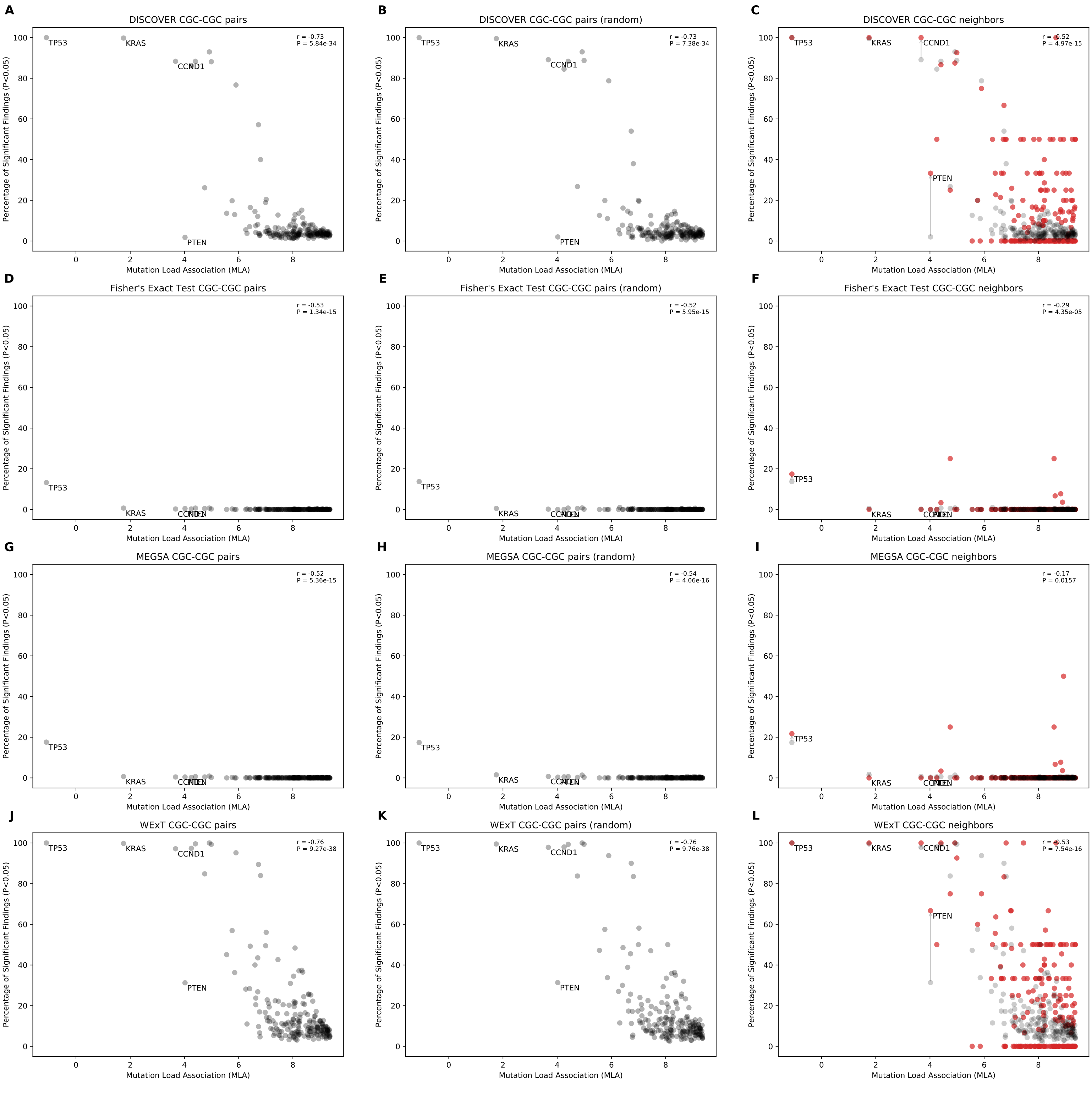


Fig 24: Comparison of mutual exclusivity results of DISCOVER, Fisher's Exact Test, MEGSA and WExT on UCEC cohort with $t = 20$ for CGC genes that have $> 1$ neighbors (531 samples)

### Scatterplots of percentage significance of mutual exclusivity runs vs mutation load association (MLA) when only CGC genes that have > 1 neighbors are included ($t = 5$)

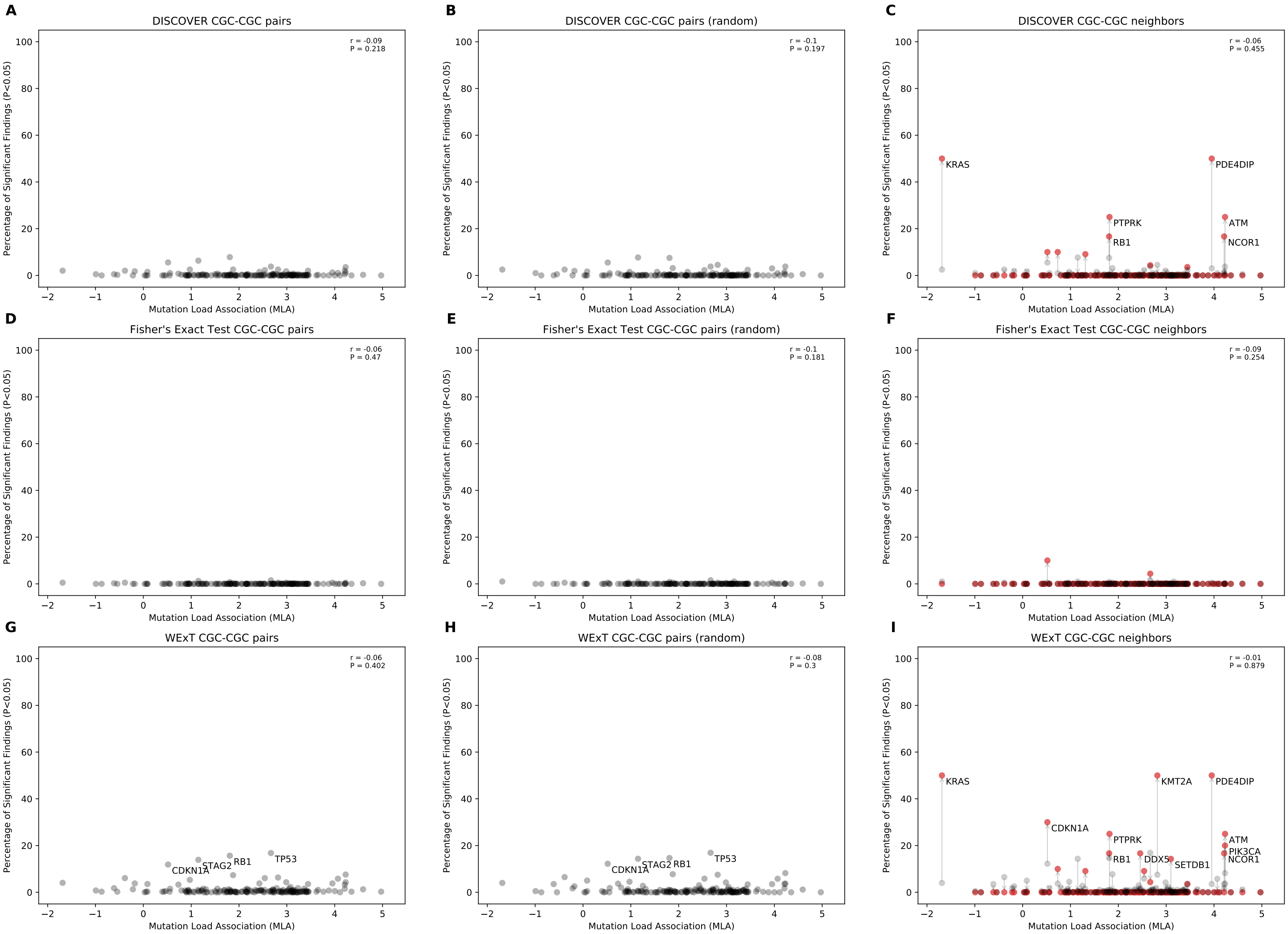


Fig 25: Comparison of mutual exclusivity results of DISCOVER, Fisher's Exact Test and WExT on BLCA cohort with $t = 5$ for CGC genes that have > 1 neighbors (411 samples)

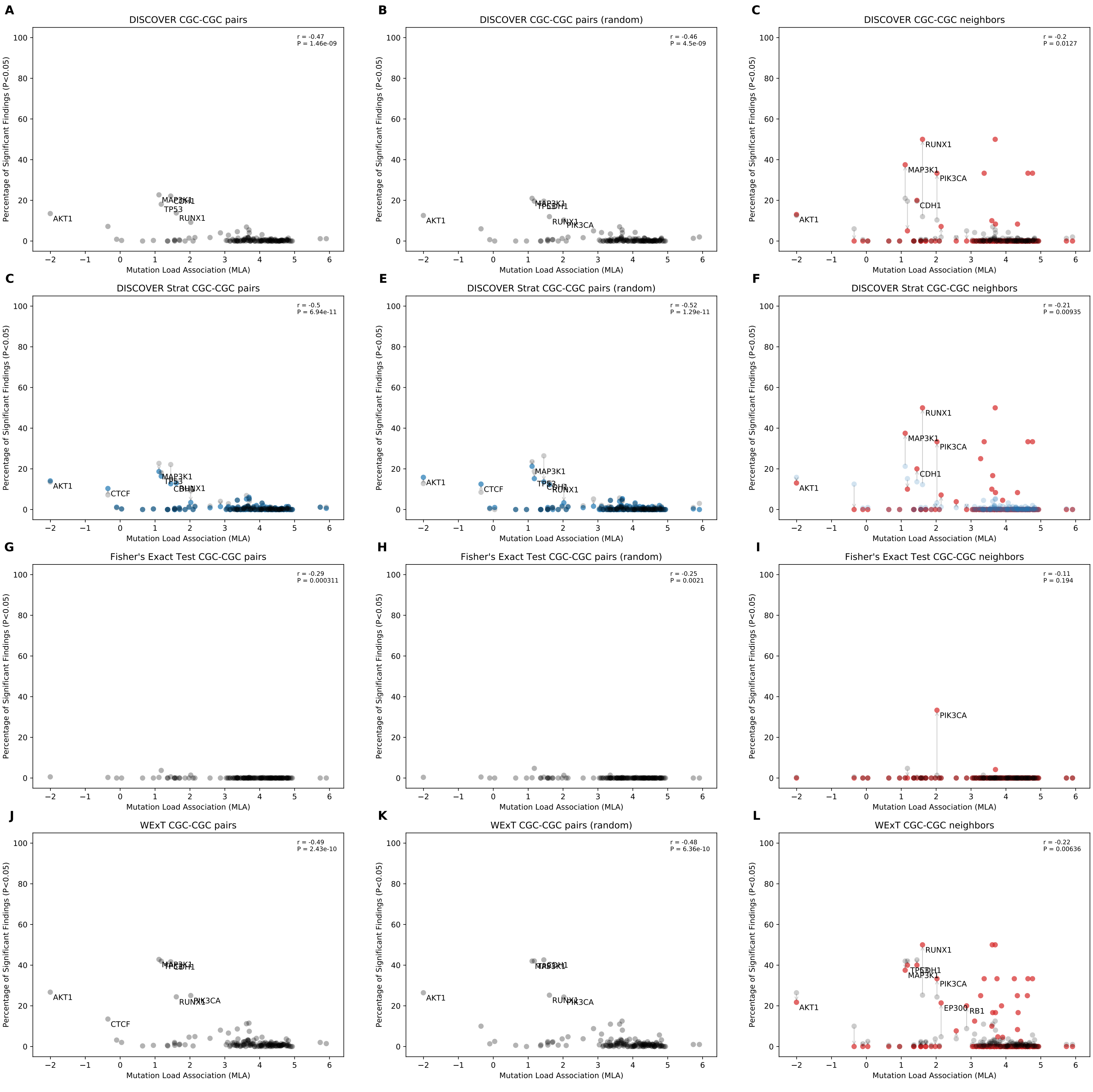


Fig 26: Comparison of mutual exclusivity results of DISCOVER, DISCOVER Strat, Fisher's Exact Test and WExT on BRCA cohort with $t = 5$ for CGC genes that have $> 1$ neighbors (1026 samples)

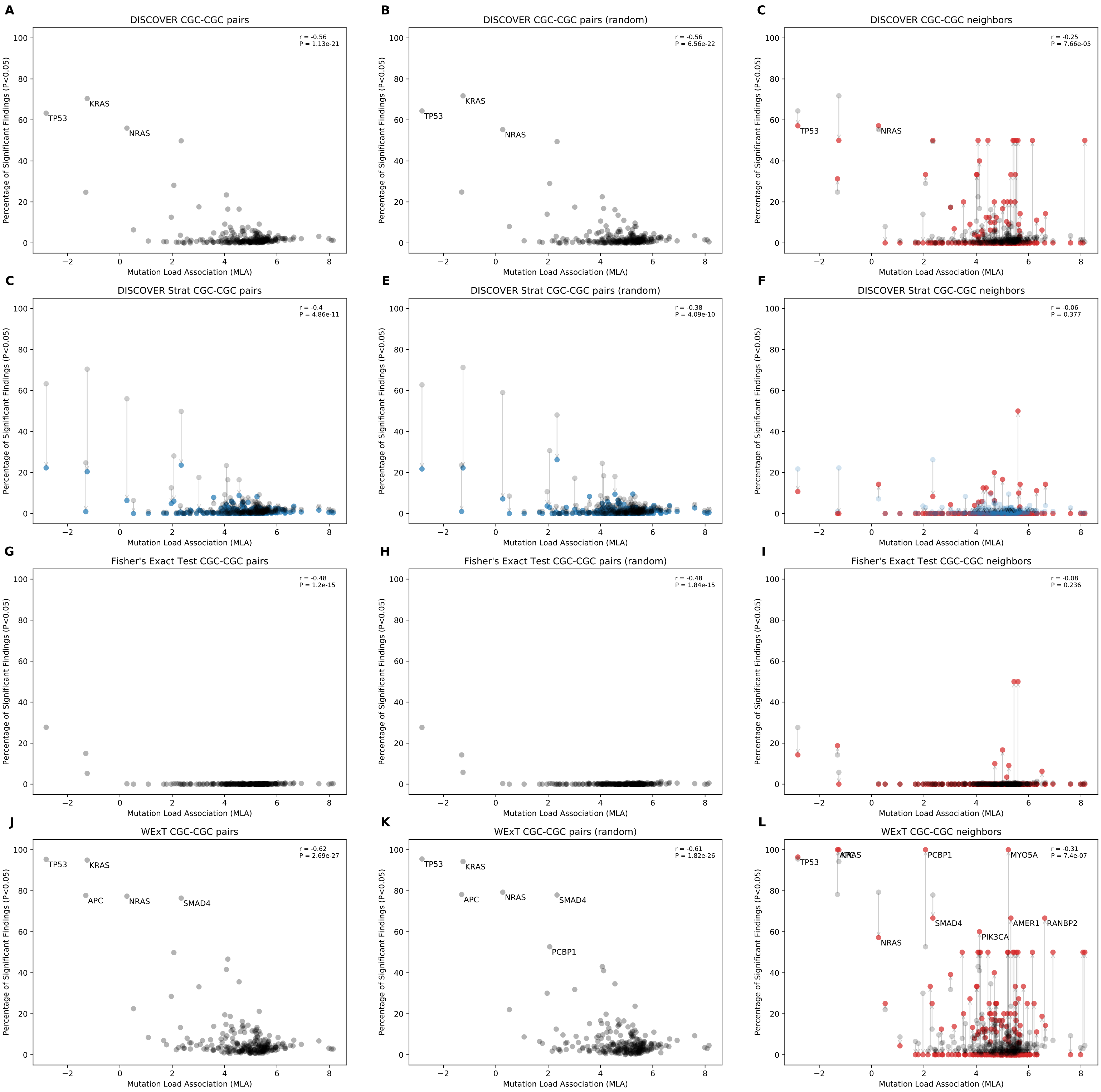


Fig 27: Comparison of mutual exclusivity results of DISCOVER, DISCOVER Strat, Fisher's Exact Test and WExT on COADREAD cohort with $t = 5$ for CGC genes that have $> 1$ neighbors (498 samples)

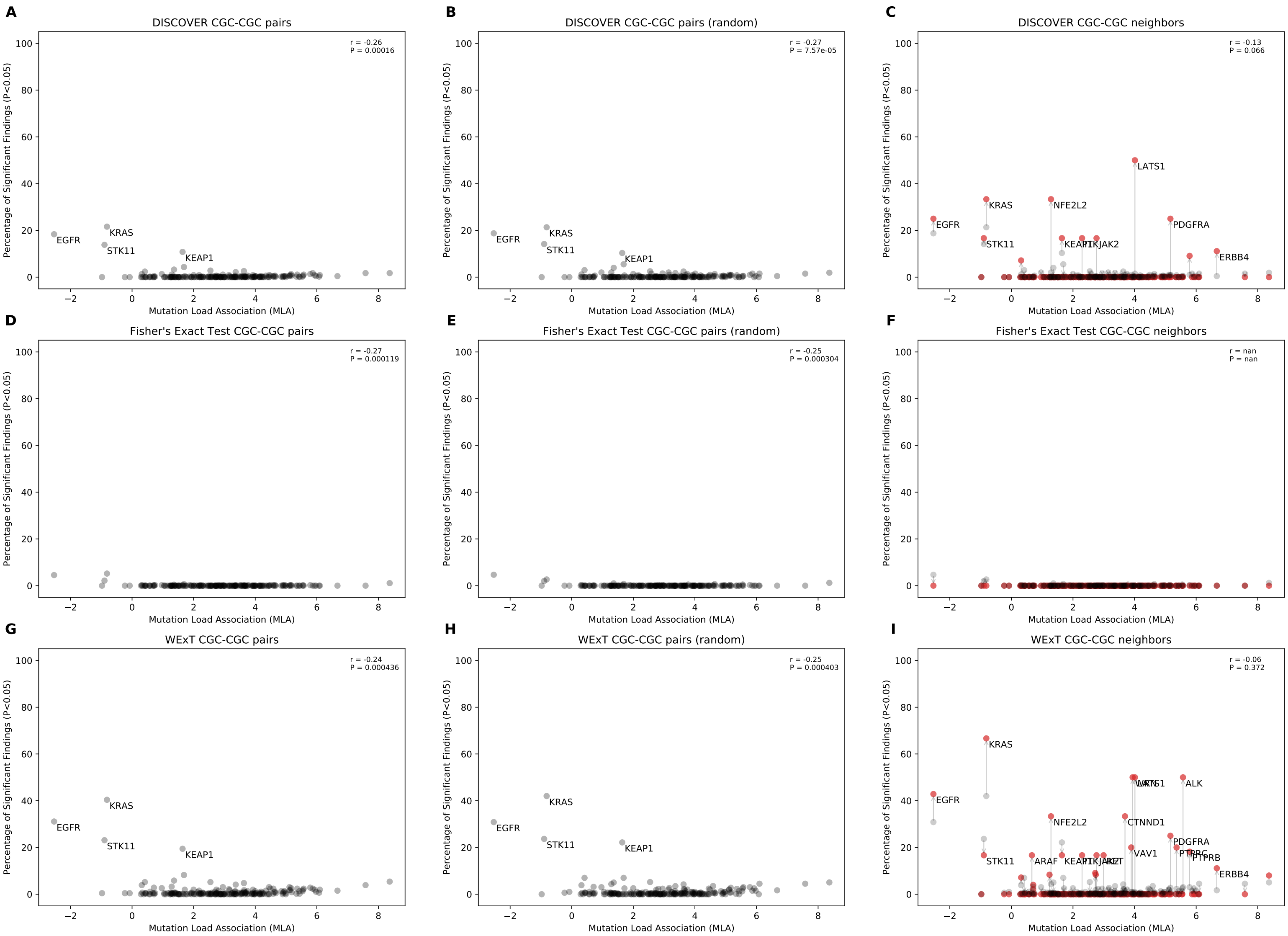


Fig 28: Comparison of mutual exclusivity results of DISCOVER, Fisher's Exact Test and WExT on LUAD cohort with $t = 5$ for CGC genes that have $> 1$ neighbors (568 samples)

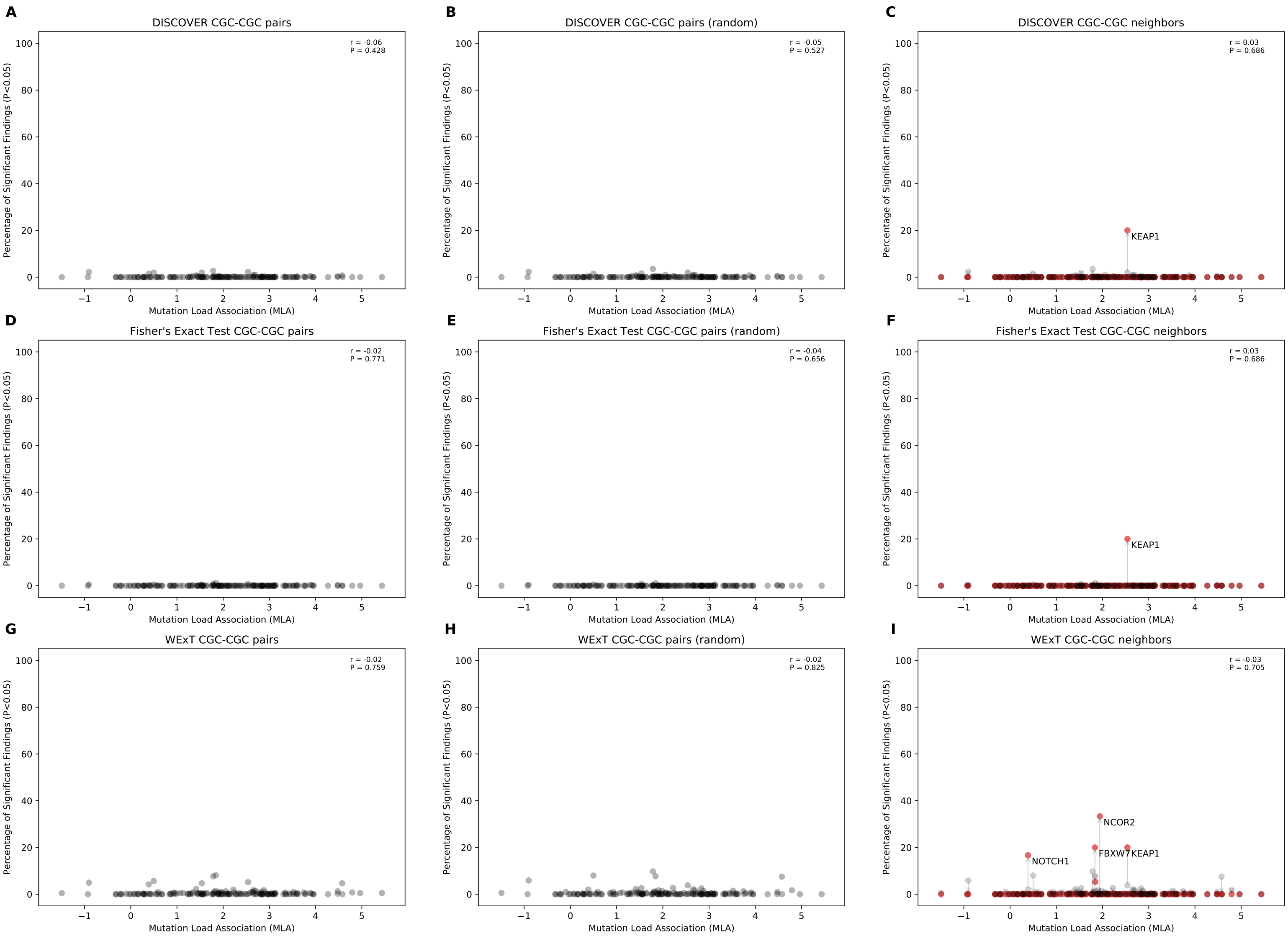


Fig 29: Comparison of mutual exclusivity results of DISCOVER, Fisher's Exact Test and WExT on LUSC cohort with $t = 5$ for CGC genes that have $> 1$ neighbors (485 samples)

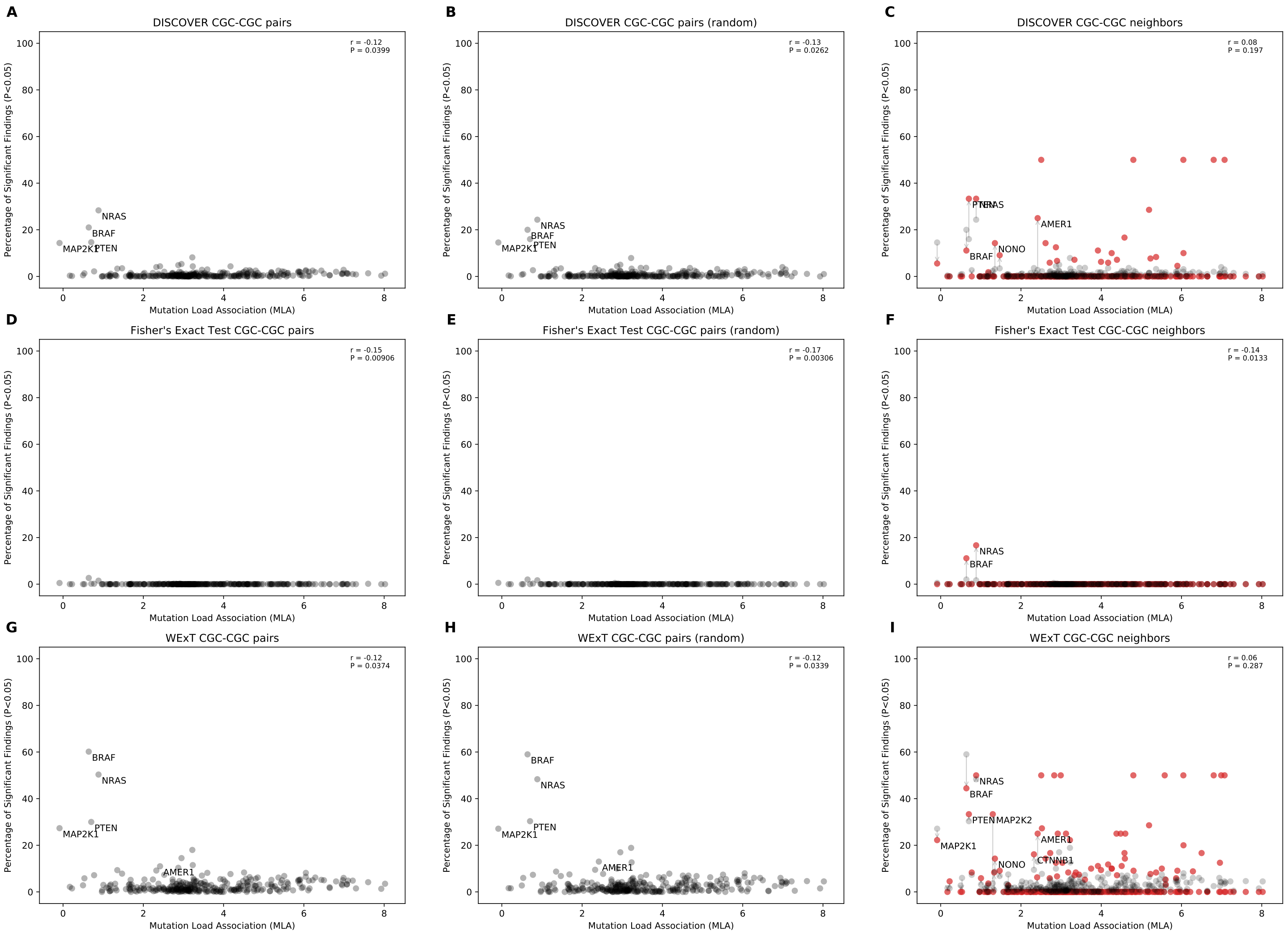


Fig 30: Comparison of mutual exclusivity results of DISCOVER, Fisher's Exact Test and WExT on SKCM cohort with $t = 5$ for CGC genes that have $> 1$ neighbors (468 samples)

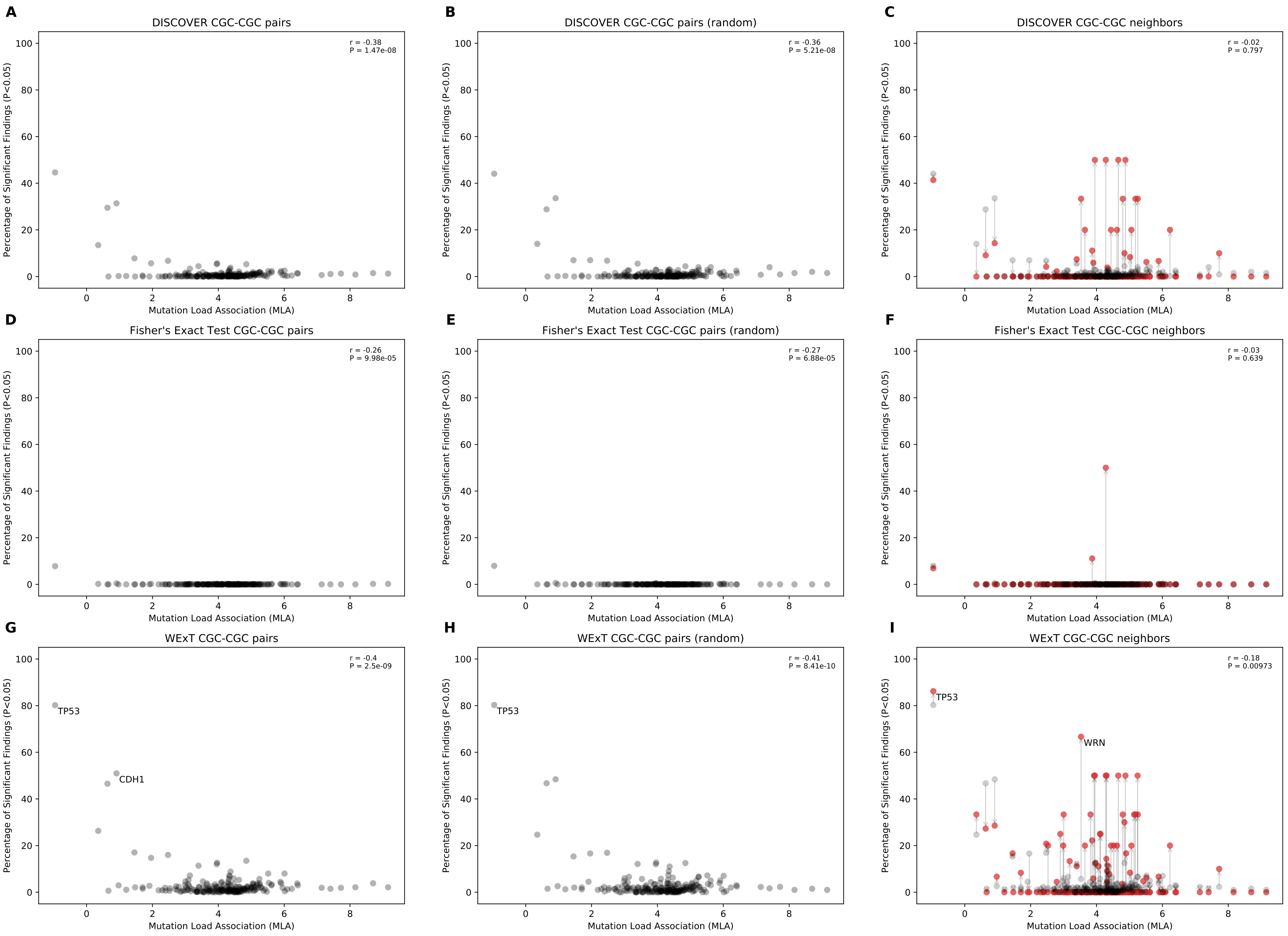


Fig 31: Comparison of mutual exclusivity results of DISCOVER, Fisher's Exact Test and WExT on STAD cohort with $t = 5$ for CGC genes that have $> 1$ neighbors (438 samples)

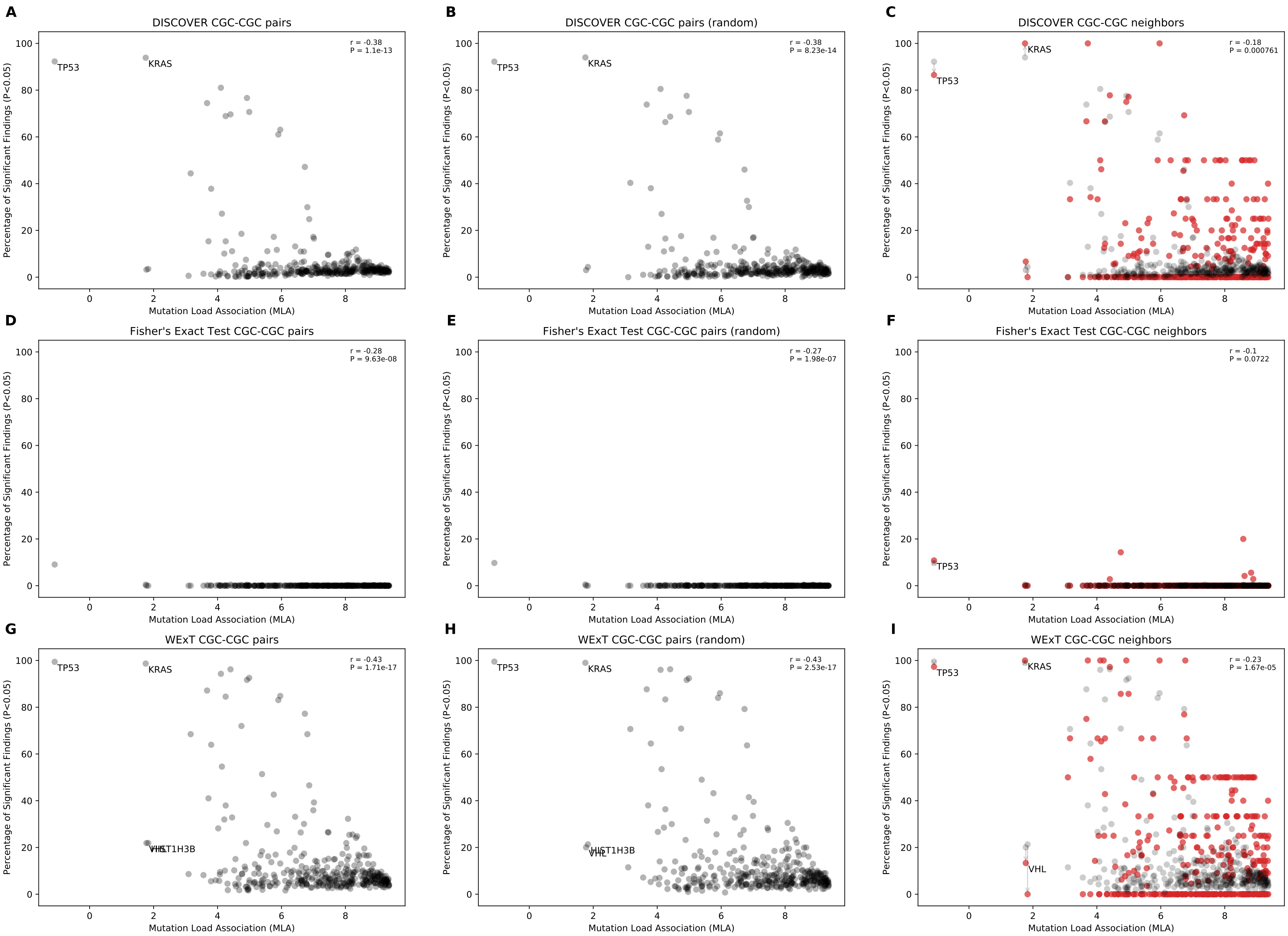


Fig 32: Comparison of mutual exclusivity results of DISCOVER, Fisher's Exact Test and WExT on UCEC cohort with $t = 5$ for CGC genes that have $> 1$ neighbors (531 samples)

Table 49: **Number of available samples for each tissue within the RNA-seq datasets downloaded from GTEx**

| Cancer Type | Tissue | Number of Samples |
|---|---|---|
| BLCA | Bladder | 21 |
| BRCA | Breast Mammary Tissue | 459 |
| COADREAD | Colon Sigmoid | 779 |
| | Colon Transverse | |
| LUAD | Lung | 578 |
| LUSC | Lung | 578 |
| SKCM | Skin Not Sun Exposed (Suprapubic) | 1809 |
| | Skin Sun Exposed (Lower leg) | |
| | Cells Cultured fibroblasts | |
| STAD | Stomach | 359 |
| UCEC | Uterus | 142 |

### Results of network-centric ME evaluation framework run on the tissue-specific network (TSN) with control group $X_1$ and $t = 20$

Table 50: **Metrics for BLCA data on the TSN network constructed with co-expression ratio threshold 0.0, $X_1$ as the control group and $t = 20$ (Bladder tissue | 21 samples | 56 CGC-CGC pairs)**

| Method | Precision | Sensitivity | F1 Score | $\text{Precision}_{\text{strict}}$ | $\text{Sensitivity}_{\text{strict}}$ | $\text{F1 Score}_{\text{strict}}$ |
|---|---|---|---|---|---|---|
| DISCOVER | 0.800 | 0.077 | 0.140 | 0.800 | 0.077 | 0.140 |
| Fisher's Exact Test | 1.000 | 0.037 | 0.071 | 1.000 | 0.037 | 0.071 |
| MEGSA | 1.000 | 0.072 | 0.134 | 1.000 | 0.072 | 0.134 |
| MEMO | 0.800 | 0.076 | 0.139 | 0.800 | 0.076 | 0.139 |
| WExT | 0.571 | 0.078 | 0.137 | 0.571 | 0.078 | 0.137 |

Table 51: **Metrics for BLCA data on the TSN network constructed with co-expression ratio threshold 0.5, $X_1$ as the control group and $t = 20$ (Bladder tissue | 21 samples | 52 CGC-CGC pairs)**

| Method | Precision | Sensitivity | F1 Score | $\text{Precision}_{\text{strict}}$ | $\text{Sensitivity}_{\text{strict}}$ | $\text{F1 Score}_{\text{strict}}$ |
|---|---|---|---|---|---|---|
| DISCOVER | 0.800 | 0.082 | 0.148 | 0.800 | 0.082 | 0.149 |
| Fisher's Exact Test | 1.000 | 0.040 | 0.077 | 1.000 | 0.040 | 0.077 |
| MEGSA | 1.000 | 0.077 | 0.143 | 1.000 | 0.077 | 0.143 |
| MEMO | 0.667 | 0.085 | 0.151 | 0.667 | 0.085 | 0.151 |
| WExT | 0.667 | 0.086 | 0.152 | 0.667 | 0.086 | 0.152 |

Table 52: **Metrics for BRCA data on the TSN network constructed with co-expression ratio threshold 0.0, $X_1$ as the control group and $t = 20$ (Breast tissue | 459 samples | 28 CGC-CGC pairs)**

| Method | Precision | Sensitivity | F1 Score | $\text{Precision}_{\text{strict}}$ | $\text{Sensitivity}_{\text{strict}}$ | $\text{F1 Score}_{\text{strict}}$ |
|---|---|---|---|---|---|---|
| DISCOVER | 0.632 | 0.444 | 0.522 | 0.625 | 0.370 | 0.465 |
| DISCOVER Strat | 0.833 | 0.417 | 0.556 | 0.833 | 0.417 | 0.556 |
| Fisher's Exact Test | 1.000 | 0.074 | 0.138 | 1.000 | 0.074 | 0.138 |
| MEGSA | 1.000 | 0.071 | 0.133 | 1.000 | 0.071 | 0.133 |
| MEMO | 0.636 | 0.467 | 0.538 | 0.630 | 0.378 | 0.472 |
| WExT | 0.550 | 0.440 | 0.489 | 0.562 | 0.360 | 0.439 |

Table 53: **Metrics for BLCA data on the TSN network constructed with co-expression ratio threshold 0.5, $X_1$ as the control group and $t = 20$ (Breast tissue | 459 samples | 28 CGC-CGC pairs)**

| Method | Precision | Sensitivity | F1 Score | $\text{Precision}_{\text{strict}}$ | $\text{Sensitivity}_{\text{strict}}$ | $\text{F1 Score}_{\text{strict}}$ |
|---|---|---|---|---|---|---|
| DISCOVER | 0.632 | 0.444 | 0.522 | 0.625 | 0.370 | 0.465 |
| DISCOVER Strat | 0.750 | 0.409 | 0.529 | 0.750 | 0.409 | 0.529 |
| Fisher's Exact Test | 1.000 | 0.071 | 0.133 | 1.000 | 0.071 | 0.133 |
| MEGSA | 1.000 | 0.071 | 0.133 | 1.000 | 0.071 | 0.133 |
| MEMO | 0.677 | 0.447 | 0.538 | 0.704 | 0.404 | 0.513 |
| WExT | 0.556 | 0.417 | 0.476 | 0.600 | 0.375 | 0.462 |

Table 54: **Metrics for COADREAD data on the TSN network constructed with co-expression ratio threshold 0.0, $X_1$ as the control group and $t = 20$ (Colon tissue | 779 samples | 194 CGC-CGC pairs)**

| Method | Precision | Sensitivity | F1 Score | $\text{Precision}_{\text{strict}}$ | $\text{Sensitivity}_{\text{strict}}$ | $\text{F1 Score}_{\text{strict}}$ |
|---|---|---|---|---|---|---|
| DISCOVER | 0.672 | 0.226 | 0.339 | 0.720 | 0.189 | 0.299 |
| DISCOVER Strat | 0.667 | 0.042 | 0.079 | 0.636 | 0.037 | 0.070 |
| Fisher's Exact Test | 0.583 | 0.037 | 0.069 | 0.583 | 0.037 | 0.070 |
| MEGSA | 0.632 | 0.062 | 0.113 | 0.611 | 0.057 | 0.104 |
| MEMO | 0.653 | 0.342 | 0.449 | 0.639 | 0.246 | 0.355 |
| WExT | 0.681 | 0.414 | 0.515 | 0.736 | 0.340 | 0.465 |

Table 55: **Metrics for COADREAD data on the TSN network constructed with co-expression ratio threshold 0.5, $X_1$ as the control group and $t = 20$ (Colon tissue | 779 samples | 194 CGC-CGC pairs)**

| Method | Precision | Sensitivity | F1 Score | $\text{Precision}_{\text{strict}}$ | $\text{Sensitivity}_{\text{strict}}$ | $\text{F1 Score}_{\text{strict}}$ |
|---|---|---|---|---|---|---|
| DISCOVER | 0.662 | 0.234 | 0.346 | 0.699 | 0.191 | 0.300 |
| DISCOVER Strat | 0.727 | 0.042 | 0.079 | 0.727 | 0.042 | 0.079 |
| Fisher's Exact Test | 0.565 | 0.034 | 0.064 | 0.565 | 0.034 | 0.064 |
| MEGSA | 0.611 | 0.057 | 0.104 | 0.588 | 0.052 | 0.096 |
| MEMO | 0.663 | 0.340 | 0.449 | 0.667 | 0.248 | 0.362 |
| WExT | 0.692 | 0.410 | 0.515 | 0.743 | 0.336 | 0.463 |

Table 56: **Metrics for LUAD data on the TSN network constructed with co-expression ratio threshold 0.0, $X_1$ as the control group and $t = 20$ (Lung tissue | 578 samples | 88 CGC-CGC pairs)**

| Method | Precision | Sensitivity | F1 Score | $\text{Precision}_{\text{strict}}$ | $\text{Sensitivity}_{\text{strict}}$ | $\text{F1 Score}_{\text{strict}}$ |
|---|---|---|---|---|---|---|
| DISCOVER | 0.750 | 0.106 | 0.186 | 0.700 | 0.082 | 0.147 |
| Fisher's Exact Test | 0.000 | 0.000 | NaN | 0.000 | 0.000 | NaN |
| MEGSA | 0.667 | 0.023 | 0.044 | 0.667 | 0.023 | 0.044 |
| MEMO | 0.722 | 0.160 | 0.263 | 0.733 | 0.136 | 0.229 |
| WExT | 0.667 | 0.187 | 0.292 | 0.700 | 0.164 | 0.266 |

Table 57: **Metrics for LUAD data on the TSN network constructed with co-expression ratio threshold 0.5, $X_1$ as the control group and $t = 20$ (Lung tissue | 578 samples | 82 CGC-CGC pairs)**

| Method | Precision | Sensitivity | F1 Score | $\text{Precision}_{\text{strict}}$ | $\text{Sensitivity}_{\text{strict}}$ | $\text{F1 Score}_{\text{strict}}$ |
|---|---|---|---|---|---|---|
| DISCOVER | 0.778 | 0.093 | 0.167 | 0.750 | 0.080 | 0.145 |
| Fisher's Exact Test | 0.000 | 0.000 | NaN | 0.000 | 0.000 | NaN |
| MEGSA | 0.667 | 0.025 | 0.048 | 0.667 | 0.025 | 0.048 |
| MEMO | 0.692 | 0.118 | 0.201 | 0.727 | 0.105 | 0.183 |
| WExT | 0.632 | 0.152 | 0.245 | 0.688 | 0.139 | 0.231 |

Table 58: **Metrics for LUSC data on the TSN network constructed with co-expression ratio threshold 0.0, $X_1$ as the control group and $t = 20$ (Lung tissue | 578 samples | 38 CGC-CGC pairs)**

| Method | Precision | Sensitivity | F1 Score | $\text{Precision}_{\text{strict}}$ | $\text{Sensitivity}_{\text{strict}}$ | $\text{F1 Score}_{\text{strict}}$ |
|---|---|---|---|---|---|---|
| DISCOVER | 1.0 | 0.053 | 0.100 | 1.0 | 0.053 | 0.101 |
| Fisher's Exact Test | 1.0 | 0.053 | 0.100 | 1.0 | 0.053 | 0.101 |
| MEGSA | 1.0 | 0.158 | 0.273 | 1.0 | 0.158 | 0.273 |
| MEMO | 1.0 | 0.113 | 0.203 | 1.0 | 0.113 | 0.203 |
| WExT | 1.0 | 0.110 | 0.198 | 1.0 | 0.110 | 0.198 |

Table 59: **Metrics for LUSC data on the TSN network constructed with co-expression ratio threshold 0.5, $X_1$ as the control group and $t = 20$ (Lung tissue | 578 samples | 36 CGC-CGC pairs)**

| Method | Precision | Sensitivity | F1 Score | $\text{Precision}_{\text{strict}}$ | $\text{Sensitivity}_{\text{strict}}$ | $\text{F1 Score}_{\text{strict}}$ |
|---|---|---|---|---|---|---|
| DISCOVER | 1.0 | 0.056 | 0.107 | 1.0 | 0.056 | 0.106 |
| Fisher's Exact Test | 1.0 | 0.056 | 0.105 | 1.0 | 0.056 | 0.106 |
| MEGSA | 1.0 | 0.167 | 0.286 | 1.0 | 0.167 | 0.286 |
| MEMO | 1.0 | 0.119 | 0.213 | 1.0 | 0.119 | 0.213 |
| WExT | 1.0 | 0.119 | 0.213 | 1.0 | 0.119 | 0.213 |

Table 60: **Metrics for SKCM data on the TSN network constructed with co-expression ratio threshold 0.0, $X_1$ as the control group and $t = 20$ (Skin tissue | 1809 samples | 456 CGC-CGC pairs)**

| Method | Precision | Sensitivity | F1 Score | $\text{Precision}_{\text{strict}}$ | $\text{Sensitivity}_{\text{strict}}$ | $\text{F1 Score}_{\text{strict}}$ |
|---|---|---|---|---|---|---|
| DISCOVER | 0.833 | 0.045 | 0.086 | 0.833 | 0.045 | 0.085 |
| Fisher's Exact Test | 1.000 | 0.004 | 0.009 | 1.000 | 0.004 | 0.008 |
| MEGSA | 0.889 | 0.018 | 0.034 | 0.889 | 0.018 | 0.035 |
| WExT | 0.687 | 0.118 | 0.202 | 0.699 | 0.115 | 0.198 |

Table 61: **Metrics for SKCM data on the TSN network constructed with co-expression ratio threshold 0.5, $X_1$ as the control group and $t = 20$ (Skin tissue | 1809 samples | 390 CGC-CGC pairs)**

| Method | Precision | Sensitivity | F1 Score | $\text{Precision}_{\text{strict}}$ | $\text{Sensitivity}_{\text{strict}}$ | $\text{F1 Score}_{\text{strict}}$ |
|---|---|---|---|---|---|---|
| DISCOVER | 0.842 | 0.042 | 0.08 | 0.842 | 0.042 | 0.080 |
| Fisher's Exact Test | 1.000 | 0.005 | 0.01 | 1.000 | 0.005 | 0.010 |
| MEGSA | 0.889 | 0.021 | 0.04 | 0.889 | 0.021 | 0.041 |
| WExT | 0.667 | 0.117 | 0.20 | 0.677 | 0.112 | 0.192 |

Table 62: **Metrics for STAD data on the TSN network constructed with co-expression ratio threshold 0.0, $X_1$ as the control group and $t = 20$ (Stomach tissue | 359 samples | 140 CGC-CGC pairs)**

| Method | Precision | Sensitivity | F1 Score | $\text{Precision}_{\text{strict}}$ | $\text{Sensitivity}_{\text{strict}}$ | $\text{F1 Score}_{\text{strict}}$ |
|---|---|---|---|---|---|---|
| DISCOVER | 0.653 | 0.119 | 0.201 | 0.667 | 0.096 | 0.168 |
| Fisher's Exact Test | 0.000 | 0.000 | NaN | 0.000 | 0.000 | NaN |
| MEGSA | 0.667 | 0.014 | 0.028 | 0.667 | 0.014 | 0.027 |
| WExT | 0.634 | 0.193 | 0.295 | 0.636 | 0.156 | 0.251 |

Table 63: **Metrics for STAD data on the TSN network constructed with co-expression ratio threshold 0.5, $X_1$ as the control group and $t = 20$ (Stomach tissue | 359 samples | 126 CGC-CGC pairs)**

| Method | Precision | Sensitivity | F1 Score | $\text{Precision}_{\text{strict}}$ | $\text{Sensitivity}_{\text{strict}}$ | $\text{F1 Score}_{\text{strict}}$ |
|---|---|---|---|---|---|---|
| DISCOVER | 0.667 | 0.130 | 0.217 | 0.684 | 0.105 | 0.182 |
| Fisher's Exact Test | 0.000 | 0.000 | NaN | 0.000 | 0.000 | NaN |
| MEGSA | 0.667 | 0.016 | 0.031 | 0.667 | 0.016 | 0.031 |
| WExT | 0.649 | 0.197 | 0.302 | 0.667 | 0.164 | 0.263 |

Table 64: **Metrics for UCEC data on the TSN network constructed with co-expression ratio threshold 0.0, $X_1$ as the control group and $t = 20$ (Uterus tissue | 142 samples | 1322 CGC-CGC pairs)**

| Method | Precision | Sensitivity | F1 Score | $\text{Precision}_{\text{strict}}$ | $\text{Sensitivity}_{\text{strict}}$ | $\text{F1 Score}_{\text{strict}}$ |
|---|---|---|---|---|---|---|
| DISCOVER | 0.653 | 0.180 | 0.282 | 0.719 | 0.146 | 0.243 |
| Fisher's Exact Test | 0.769 | 0.008 | 0.015 | 0.769 | 0.008 | 0.016 |
| MEGSA | 0.786 | 0.008 | 0.016 | 0.786 | 0.008 | 0.016 |
| WExT | 0.619 | 0.282 | 0.388 | 0.667 | 0.230 | 0.342 |

Table 65: **Metrics for UCEC data on the TSN network constructed with co-expression ratio threshold 0.5, $X_1$ as the control group and $t = 20$ (Uterus tissue | 142 samples | 1224 CGC-CGC pairs)**

| Method | Precision | Sensitivity | F1 Score | $\text{Precision}_{\text{strict}}$ | $\text{Sensitivity}_{\text{strict}}$ | $\text{F1 Score}_{\text{strict}}$ |
|---|---|---|---|---|---|---|
| DISCOVER | 0.658 | 0.186 | 0.290 | 0.717 | 0.150 | 0.248 |
| Fisher's Exact Test | 0.833 | 0.008 | 0.016 | 0.833 | 0.008 | 0.016 |
| MEGSA | 0.786 | 0.009 | 0.018 | 0.786 | 0.009 | 0.018 |
| WExT | 0.614 | 0.286 | 0.390 | 0.665 | 0.233 | 0.345 |

### Results of network-centric ME evaluation framework run on the tissue-specific network (TSN) with control group $X_2$ and $t = 20$

Table 66: **Metrics for BLCA data on the TSN network constructed with co-expression ratio threshold 0.0, $X_2$ as the control group and $t = 20$ (Bladder tissue | 21 samples | 24 CGC-CGC pairs)**

| Method | Precision | Sensitivity | F1 Score | $\text{Precision}_{\text{strict}}$ | $\text{Sensitivity}_{\text{strict}}$ | $\text{F1 Score}_{\text{strict}}$ |
|---|---|---|---|---|---|---|
| DISCOVER | 0.537 | 0.276 | 0.365 | 0.579 | 0.210 | 0.308 |
| DISCOVER Strat | 0.455 | 0.048 | 0.086 | 0.400 | 0.038 | 0.069 |
| Fisher's Exact Test | 0.444 | 0.038 | 0.069 | 0.375 | 0.028 | 0.052 |
| MEGSA | 0.571 | 0.075 | 0.133 | 0.538 | 0.066 | 0.118 |
| MEMO | 0.566 | 0.388 | 0.460 | 0.495 | 0.215 | 0.300 |
| WExT | 0.575 | 0.438 | 0.497 | 0.596 | 0.295 | 0.395 |

Table 67: **Metrics for BLCA data on the TSN network constructed with co-expression ratio threshold 0.5, $X_2$ as the control group and $t = 20$ (Bladder tissue | 21 samples | 21 CGC-CGC pairs)**

| Method | Precision | Sensitivity | F1 Score | $\text{Precision}_{\text{strict}}$ | $\text{Sensitivity}_{\text{strict}}$ | $\text{F1 Score}_{\text{strict}}$ |
|---|---|---|---|---|---|---|
| DISCOVER | 0.537 | 0.276 | 0.365 | 0.579 | 0.210 | 0.308 |
| DISCOVER Strat | 0.455 | 0.048 | 0.086 | 0.400 | 0.038 | 0.069 |
| Fisher's Exact Test | 0.444 | 0.038 | 0.069 | 0.375 | 0.028 | 0.052 |
| MEGSA | 0.571 | 0.075 | 0.133 | 0.538 | 0.066 | 0.118 |
| MEMO | 0.566 | 0.388 | 0.460 | 0.495 | 0.215 | 0.300 |
| WExT | 0.575 | 0.438 | 0.497 | 0.596 | 0.295 | 0.395 |

Table 68: **Metrics for BRCA data on the TSN network constructed with co-expression ratio threshold 0.0, $X_2$ as the control group and $t = 20$ (Breast tissue | 459 samples | 9 CGC-CGC pairs)**

| Method | Precision | Sensitivity | F1 Score | $\text{Precision}_{\text{strict}}$ | $\text{Sensitivity}_{\text{strict}}$ | $\text{F1 Score}_{\text{strict}}$ |
|---|---|---|---|---|---|---|
| DISCOVER | 0.537 | 0.276 | 0.365 | 0.579 | 0.210 | 0.308 |
| DISCOVER Strat | 0.455 | 0.048 | 0.086 | 0.400 | 0.038 | 0.069 |
| Fisher's Exact Test | 0.444 | 0.038 | 0.069 | 0.375 | 0.028 | 0.052 |
| MEGSA | 0.571 | 0.075 | 0.133 | 0.538 | 0.066 | 0.118 |
| MEMO | 0.566 | 0.388 | 0.460 | 0.495 | 0.215 | 0.300 |
| WExT | 0.575 | 0.438 | 0.497 | 0.596 | 0.295 | 0.395 |

Table 69: **Metrics for BRCA data on the TSN network constructed with co-expression ratio threshold 0.5, $X_2$ as the control group and $t = 20$ (Breast tissue | 459 samples | 5 CGC-CGC pairs)**

| Method | Precision | Sensitivity | F1 Score | $\text{Precision}_{\text{strict}}$ | $\text{Sensitivity}_{\text{strict}}$ | $\text{F1 Score}_{\text{strict}}$ |
|---|---|---|---|---|---|---|
| DISCOVER | 0.537 | 0.276 | 0.365 | 0.579 | 0.210 | 0.308 |
| DISCOVER Strat | 0.455 | 0.048 | 0.086 | 0.400 | 0.038 | 0.069 |
| Fisher's Exact Test | 0.444 | 0.038 | 0.069 | 0.375 | 0.028 | 0.052 |
| MEGSA | 0.571 | 0.075 | 0.133 | 0.538 | 0.066 | 0.118 |
| MEMO | 0.566 | 0.388 | 0.460 | 0.495 | 0.215 | 0.300 |
| WExT | 0.575 | 0.438 | 0.497 | 0.596 | 0.295 | 0.395 |

Table 70: **Metrics for COADREAD data on the TSN network constructed with co-expression ratio threshold 0.0, $X_2$ as the control group and $t = 20$ (Colon tissue | 779 samples | 105 CGC-CGC pairs)**

| Method | Precision | Sensitivity | F1 Score | $\text{Precision}_{\text{strict}}$ | $\text{Sensitivity}_{\text{strict}}$ | $\text{F1 Score}_{\text{strict}}$ |
|---|---|---|---|---|---|---|
| DISCOVER | 0.537 | 0.276 | 0.365 | 0.579 | 0.210 | 0.308 |
| DISCOVER Strat | 0.455 | 0.048 | 0.086 | 0.400 | 0.038 | 0.069 |
| Fisher's Exact Test | 0.444 | 0.038 | 0.069 | 0.375 | 0.028 | 0.052 |
| MEGSA | 0.571 | 0.075 | 0.133 | 0.538 | 0.066 | 0.118 |
| MEMO | 0.566 | 0.388 | 0.460 | 0.495 | 0.215 | 0.300 |
| WExT | 0.575 | 0.438 | 0.497 | 0.596 | 0.295 | 0.395 |

Table 71: **Metrics for COADREAD data on the TSN network constructed with co-expression ratio threshold 0.5, $X_2$ as the control group and $t = 20$ (Colon tissue | 779 samples | 105 CGC-CGC pairs)**

| Method | Precision | Sensitivity | F1 Score | $\text{Precision}_{\text{strict}}$ | $\text{Sensitivity}_{\text{strict}}$ | $\text{F1 Score}_{\text{strict}}$ |
|---|---|---|---|---|---|---|
| DISCOVER | 0.537 | 0.276 | 0.365 | 0.579 | 0.210 | 0.308 |
| DISCOVER Strat | 0.455 | 0.048 | 0.086 | 0.400 | 0.038 | 0.069 |
| Fisher's Exact Test | 0.444 | 0.038 | 0.069 | 0.375 | 0.028 | 0.052 |
| MEGSA | 0.571 | 0.075 | 0.133 | 0.538 | 0.066 | 0.118 |
| MEMO | 0.566 | 0.388 | 0.460 | 0.495 | 0.215 | 0.300 |
| WExT | 0.575 | 0.438 | 0.497 | 0.596 | 0.295 | 0.395 |

Table 72: **Metrics for LUAD data on the TSN network constructed with co-expression ratio threshold 0.0, $X_2$ as the control group and $t = 20$ (Lung tissue | 578 samples | 53 CGC-CGC pairs)**

| Method | Precision | Sensitivity | F1 Score | $\text{Precision}_{\text{strict}}$ | $\text{Sensitivity}_{\text{strict}}$ | $\text{F1 Score}_{\text{strict}}$ |
|---|---|---|---|---|---|---|
| DISCOVER | 0.537 | 0.276 | 0.365 | 0.579 | 0.210 | 0.308 |
| DISCOVER Strat | 0.455 | 0.048 | 0.086 | 0.400 | 0.038 | 0.069 |
| Fisher's Exact Test | 0.444 | 0.038 | 0.069 | 0.375 | 0.028 | 0.052 |
| MEGSA | 0.571 | 0.075 | 0.133 | 0.538 | 0.066 | 0.118 |
| MEMO | 0.566 | 0.388 | 0.460 | 0.495 | 0.215 | 0.300 |
| WExT | 0.575 | 0.438 | 0.497 | 0.596 | 0.295 | 0.395 |

Table 73: **Metrics for LUAD data on the TSN network constructed with co-expression ratio threshold 0.5, $X_2$ as the control group and $t = 20$ (Lung tissue | 578 samples | 34 CGC-CGC pairs)**

| Method | Precision | Sensitivity | F1 Score | $\text{Precision}_{\text{strict}}$ | $\text{Sensitivity}_{\text{strict}}$ | $\text{F1 Score}_{\text{strict}}$ |
|---|---|---|---|---|---|---|
| DISCOVER | 0.537 | 0.276 | 0.365 | 0.579 | 0.210 | 0.308 |
| DISCOVER Strat | 0.455 | 0.048 | 0.086 | 0.400 | 0.038 | 0.069 |
| Fisher's Exact Test | 0.444 | 0.038 | 0.069 | 0.375 | 0.028 | 0.052 |
| MEGSA | 0.571 | 0.075 | 0.133 | 0.538 | 0.066 | 0.118 |
| MEMO | 0.566 | 0.388 | 0.460 | 0.495 | 0.215 | 0.300 |
| WExT | 0.575 | 0.438 | 0.497 | 0.596 | 0.295 | 0.395 |

Table 74: **Metrics for LUSC data on the TSN network constructed with co-expression ratio threshold 0.0, $X_2$ as the control group and $t = 20$ (Lung tissue | 578 samples | 21 CGC-CGC pairs)**

| Method | Precision | Sensitivity | F1 Score | $\text{Precision}_{\text{strict}}$ | $\text{Sensitivity}_{\text{strict}}$ | $\text{F1 Score}_{\text{strict}}$ |
|---|---|---|---|---|---|---|
| DISCOVER | 0.537 | 0.276 | 0.365 | 0.579 | 0.210 | 0.308 |
| DISCOVER Strat | 0.455 | 0.048 | 0.086 | 0.400 | 0.038 | 0.069 |
| Fisher's Exact Test | 0.444 | 0.038 | 0.069 | 0.375 | 0.028 | 0.052 |
| MEGSA | 0.571 | 0.075 | 0.133 | 0.538 | 0.066 | 0.118 |
| MEMO | 0.566 | 0.388 | 0.460 | 0.495 | 0.215 | 0.300 |
| WExT | 0.575 | 0.438 | 0.497 | 0.596 | 0.295 | 0.395 |

Table 75: **Metrics for LUSC data on the TSN network constructed with co-expression ratio threshold 0.5, $X_2$ as the control group and $t = 20$ (Lung tissue | 578 samples | 20 CGC-CGC pairs)**

| Method | Precision | Sensitivity | F1 Score | $\text{Precision}_{\text{strict}}$ | $\text{Sensitivity}_{\text{strict}}$ | $\text{F1 Score}_{\text{strict}}$ |
|---|---|---|---|---|---|---|
| DISCOVER | 0.537 | 0.276 | 0.365 | 0.579 | 0.210 | 0.308 |
| DISCOVER Strat | 0.455 | 0.048 | 0.086 | 0.400 | 0.038 | 0.069 |
| Fisher's Exact Test | 0.444 | 0.038 | 0.069 | 0.375 | 0.028 | 0.052 |
| MEGSA | 0.571 | 0.075 | 0.133 | 0.538 | 0.066 | 0.118 |
| MEMO | 0.566 | 0.388 | 0.460 | 0.495 | 0.215 | 0.300 |
| WExT | 0.575 | 0.438 | 0.497 | 0.596 | 0.295 | 0.395 |

Table 76: **Metrics for SKCM data on the TSN network constructed with co-expression ratio threshold 0.0, $X_2$ as the control group and $t = 20$ (Skin tissue | 1809 samples | 312 CGC-CGC pairs)**

| Method | Precision | Sensitivity | F1 Score | $\text{Precision}_{\text{strict}}$ | $\text{Sensitivity}_{\text{strict}}$ | $\text{F1 Score}_{\text{strict}}$ |
|---|---|---|---|---|---|---|
| DISCOVER | 0.537 | 0.276 | 0.365 | 0.579 | 0.210 | 0.308 |
| DISCOVER Strat | 0.455 | 0.048 | 0.086 | 0.400 | 0.038 | 0.069 |
| Fisher's Exact Test | 0.444 | 0.038 | 0.069 | 0.375 | 0.028 | 0.052 |
| MEGSA | 0.571 | 0.075 | 0.133 | 0.538 | 0.066 | 0.118 |
| MEMO | 0.566 | 0.388 | 0.460 | 0.495 | 0.215 | 0.300 |
| WExT | 0.575 | 0.438 | 0.497 | 0.596 | 0.295 | 0.395 |

Table 77: **Metrics for SKCM data on the TSN network constructed with co-expression ratio threshold 0.5, $X_2$ as the control group and $t = 20$ (Skin tissue | 1809 samples | 263 CGC-CGC pairs)**

| Method | Precision | Sensitivity | F1 Score | $\text{Precision}_{\text{strict}}$ | $\text{Sensitivity}_{\text{strict}}$ | $\text{F1 Score}_{\text{strict}}$ |
|---|---|---|---|---|---|---|
| DISCOVER | 0.537 | 0.276 | 0.365 | 0.579 | 0.210 | 0.308 |
| DISCOVER Strat | 0.455 | 0.048 | 0.086 | 0.400 | 0.038 | 0.069 |
| Fisher's Exact Test | 0.444 | 0.038 | 0.069 | 0.375 | 0.028 | 0.052 |
| MEGSA | 0.571 | 0.075 | 0.133 | 0.538 | 0.066 | 0.118 |
| MEMO | 0.566 | 0.388 | 0.460 | 0.495 | 0.215 | 0.300 |
| WExT | 0.575 | 0.438 | 0.497 | 0.596 | 0.295 | 0.395 |

Table 78: **Metrics for STAD data on the TSN network constructed with co-expression ratio threshold 0.0, $X_2$ as the control group and $t = 20$ (Stomach tissue | 359 samples | 70 CGC-CGC pairs)**

| Method | Precision | Sensitivity | F1 Score | $\text{Precision}_{\text{strict}}$ | $\text{Sensitivity}_{\text{strict}}$ | $\text{F1 Score}_{\text{strict}}$ |
|---|---|---|---|---|---|---|
| DISCOVER | 0.537 | 0.276 | 0.365 | 0.579 | 0.210 | 0.308 |
| DISCOVER Strat | 0.455 | 0.048 | 0.086 | 0.400 | 0.038 | 0.069 |
| Fisher's Exact Test | 0.444 | 0.038 | 0.069 | 0.375 | 0.028 | 0.052 |
| MEGSA | 0.571 | 0.075 | 0.133 | 0.538 | 0.066 | 0.118 |
| MEMO | 0.566 | 0.388 | 0.460 | 0.495 | 0.215 | 0.300 |
| WExT | 0.575 | 0.438 | 0.497 | 0.596 | 0.295 | 0.395 |

Table 79: **Metrics for STAD data on the TSN network constructed with co-expression ratio threshold 0.5, $X_2$ as the control group and $t = 20$ (Stomach tissue | 359 samples | 53 CGC-CGC pairs)**

| Method | Precision | Sensitivity | F1 Score | $\text{Precision}_{\text{strict}}$ | $\text{Sensitivity}_{\text{strict}}$ | $\text{F1 Score}_{\text{strict}}$ |
|---|---|---|---|---|---|---|
| DISCOVER | 0.537 | 0.276 | 0.365 | 0.579 | 0.210 | 0.308 |
| DISCOVER Strat | 0.455 | 0.048 | 0.086 | 0.400 | 0.038 | 0.069 |
| Fisher's Exact Test | 0.444 | 0.038 | 0.069 | 0.375 | 0.028 | 0.052 |
| MEGSA | 0.571 | 0.075 | 0.133 | 0.538 | 0.066 | 0.118 |
| MEMO | 0.566 | 0.388 | 0.460 | 0.495 | 0.215 | 0.300 |
| WExT | 0.575 | 0.438 | 0.497 | 0.596 | 0.295 | 0.395 |

Table 80: **Metrics for UCEC data on the TSN network constructed with co-expression ratio threshold 0.0, $X_2$ as the control group and $t = 20$ (Uterus tissue | 142 samples | 1159 CGC-CGC pairs)**

| Method | Precision | Sensitivity | F1 Score | $\text{Precision}_{\text{strict}}$ | $\text{Sensitivity}_{\text{strict}}$ | $\text{F1 Score}_{\text{strict}}$ |
|---|---|---|---|---|---|---|
| DISCOVER | 0.537 | 0.276 | 0.365 | 0.579 | 0.210 | 0.308 |
| DISCOVER Strat | 0.455 | 0.048 | 0.086 | 0.400 | 0.038 | 0.069 |
| Fisher's Exact Test | 0.444 | 0.038 | 0.069 | 0.375 | 0.028 | 0.052 |
| MEGSA | 0.571 | 0.075 | 0.133 | 0.538 | 0.066 | 0.118 |
| MEMO | 0.566 | 0.388 | 0.460 | 0.495 | 0.215 | 0.300 |
| WExT | 0.575 | 0.438 | 0.497 | 0.596 | 0.295 | 0.395 |

Table 81: **Metrics for UCEC data on the TSN network constructed with co-expression ratio threshold 0.5, $X_2$ as the control group and $t = 20$ (Uterus tissue | 142 samples | 1078 CGC-CGC pairs)**

| Method | Precision | Sensitivity | F1 Score | $\text{Precision}_{\text{strict}}$ | $\text{Sensitivity}_{\text{strict}}$ | $\text{F1 Score}_{\text{strict}}$ |
|---|---|---|---|---|---|---|
| DISCOVER | 0.537 | 0.276 | 0.365 | 0.579 | 0.210 | 0.308 |
| DISCOVER Strat | 0.455 | 0.048 | 0.086 | 0.400 | 0.038 | 0.069 |
| Fisher's Exact Test | 0.444 | 0.038 | 0.069 | 0.375 | 0.028 | 0.052 |
| MEGSA | 0.571 | 0.075 | 0.133 | 0.538 | 0.066 | 0.118 |
| MEMO | 0.566 | 0.388 | 0.460 | 0.495 | 0.215 | 0.300 |
| WExT | 0.575 | 0.438 | 0.497 | 0.596 | 0.295 | 0.395 |

Mutual exclusivities of tissue-specific and non-tissue-specific gene pairs

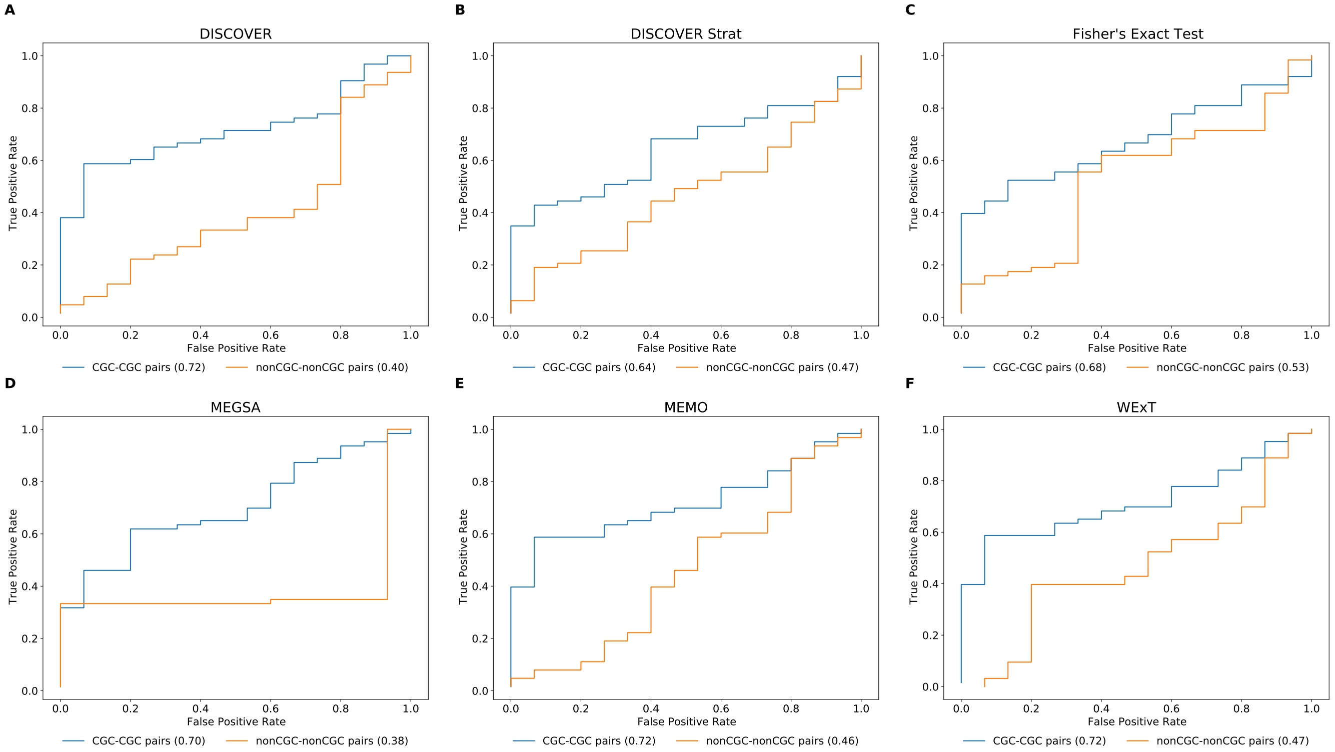


Fig 33: ROC curves for comparing the mutual exclusivities of tissue-specific and non-tissue-specific CGC-CGC gene pairs and non-CGC-non-CGC gene pairs on COADREAD data with $t = 20$ setting. (positive set: 63 edges, negative set: 15 edges)

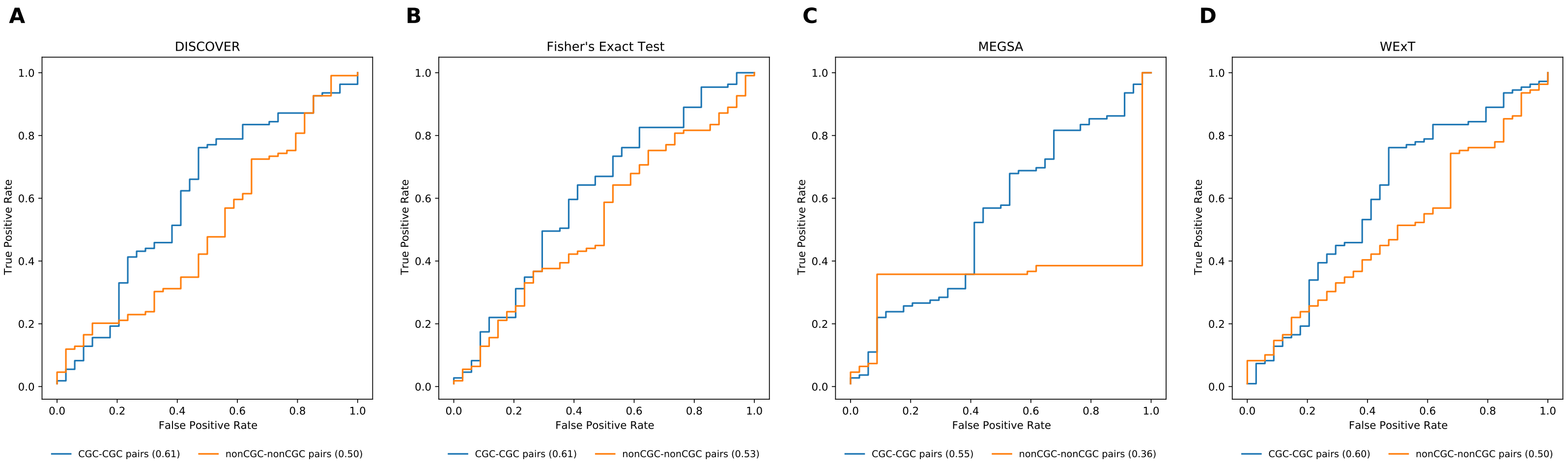


Fig 34: ROC curves for comparing the mutual exclusivities of tissue-specific and non-tissue-specific CGC-CGC gene pairs and non-CGC-non-CGC gene pairs on SKCM data with $t = 20$ setting. (positive set: 109 edges, negative set: 34 edges)

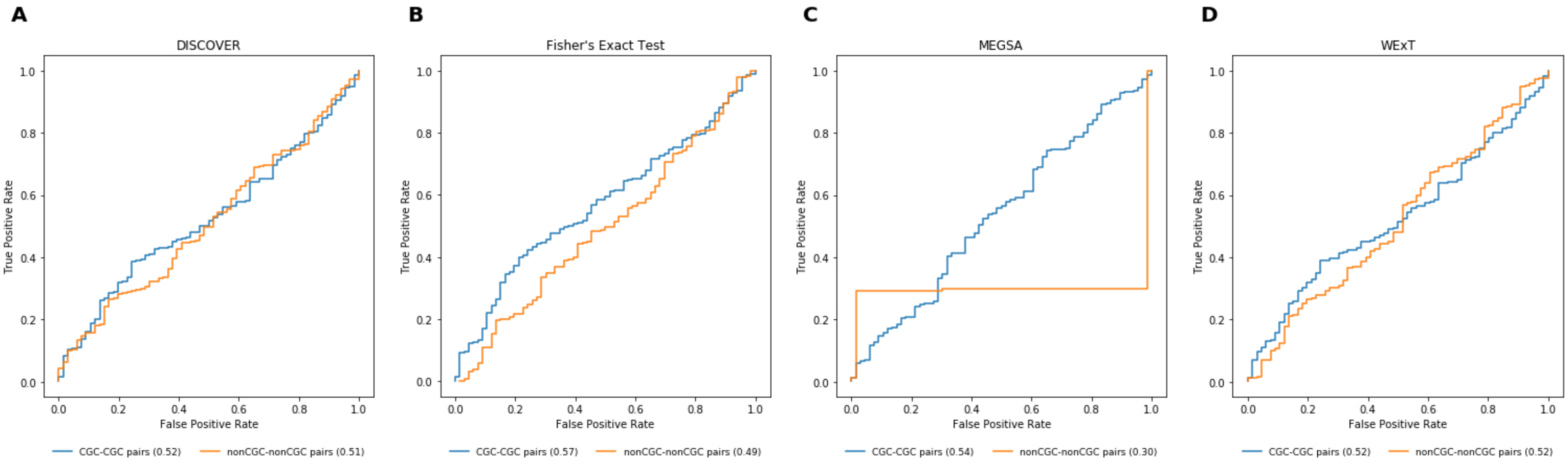


Fig 35: ROC curves for comparing the mutual exclusivities of tissue-specific and non-tissue-specific CGC-CGC gene pairs and non-CGC-non-CGC gene pairs on UCEC data with $t = 20$ setting. (positive set: 519 edges, negative set: 66 edges)

### Effect of estimating ME with DISCOVER, Fisher's Exact Test and WExT within the MEXCOwalk model

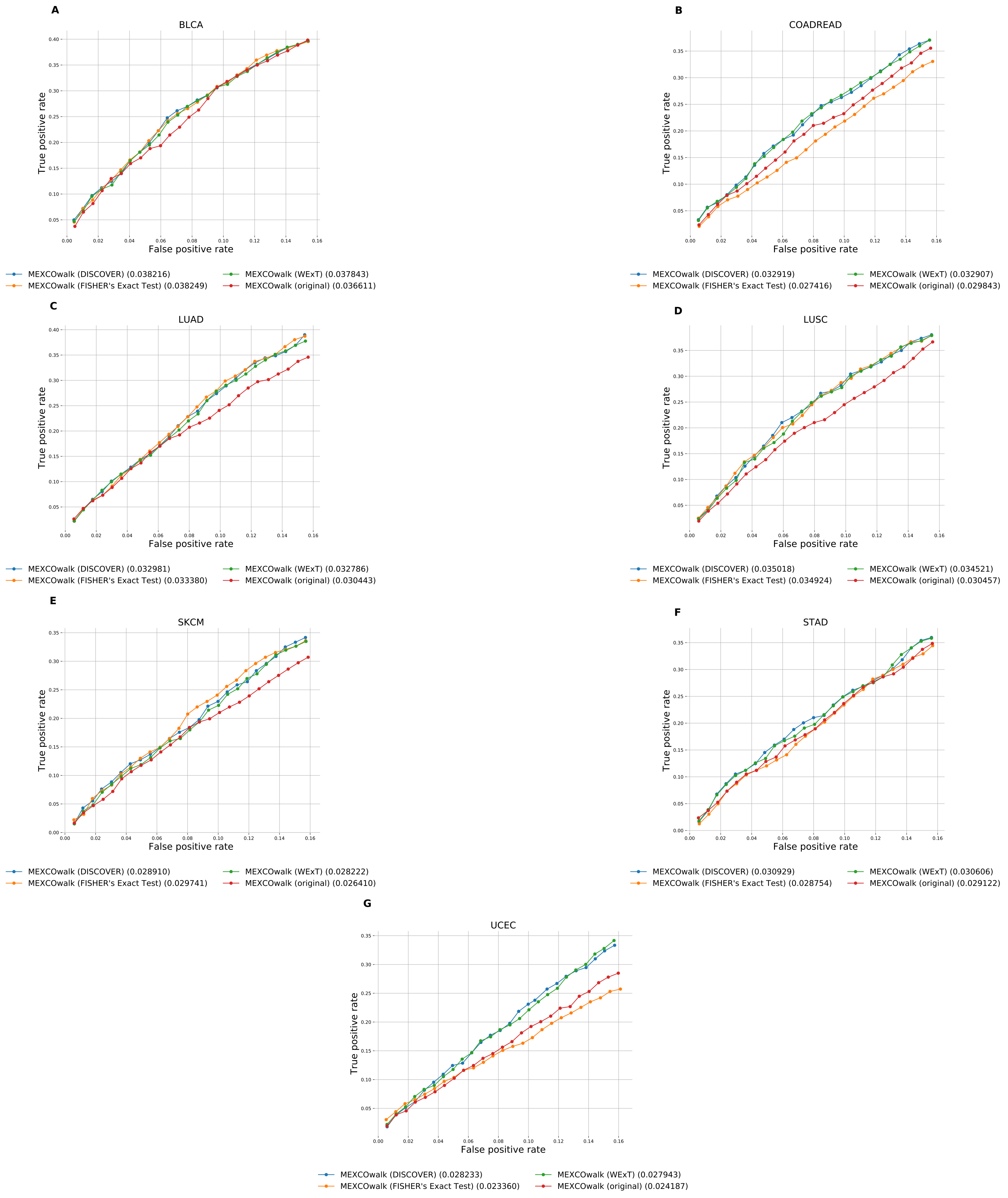
